# A robust metatranscriptomic technology for population-scale studies of diet, gut microbiome, and human health

**DOI:** 10.1101/659615

**Authors:** Andrew Hatch, James Horne, Ryan Toma, Brittany L. Twibell, Kalie M. Somerville, Benjamin Pelle, Kinga P. Canfield, Matvey Genkin, Guruduth Banavar, Ally Perlina, Helen Messier, Niels Klitgord, Momchilo Vuyisich

## Abstract

A functional readout of the gut microbiome is necessary to enable precise control of the gut microbiome’s functions, which support human health and prevent or minimize a wide range of chronic diseases. Stool metatranscriptomic analysis offers a comprehensive functional view of the gut microbiome, but despite its usefulness, it has rarely been used in clinical studies due to its complexity, cost, and bioinformatic challenges. This method has also received criticism due to potential intra-sample variability, rapid changes, and RNA degradation. Here, we describe a robust and automated stool metatranscriptomic method, called Viomega, which was specifically developed for population-scale studies. Viomega includes sample collection, ambient temperature sample preservation, total RNA extraction, physical removal of ribosomal RNAs (rRNAs), preparation of directional Illumina libraries, Illumina sequencing, taxonomic classification based on a database of >110,000 microbial genomes, and quantitative microbial gene expression analysis using a database of ~100 million microbial genes. We applied this method to 10,000 human stool samples, and performed several small-scale studies to demonstrate sample stability and consistency. In summary, Viomega is an inexpensive, high throughput, automated, and accurate sample-to-result stool metatranscriptomic technology platform for large-scale studies and a wide range of applications.

## INTRODUCTION

The human gut contains a vast number of commensal microorganisms performing a wide variety of metabolic functions. Metabolites produced by these microorganisms can have profound effects on human physiology, with direct links to health and disease status [1–4]. Gut dysbiosis likely contributes to the development and progression of many diseases and disorders, such as cardiovascular disease, hypertension, obesity, diabetes, and autoimmune diseases [5–9]. There is also strong evidence that the gut microorganisms directly interact with the nervous system, establishing the gut-brain axis [10]. The gut-brain axis has been shown to modulate the development of neurodegenerative diseases such as Alzheimer’s disease, Autism Spectrum Disorder (ASD) and Parkinson’s disease [11–14].

The gut microbiome plays a critical role in physiological homeostasis, resulting in increasing scientific investigation into the extent of the gut microbiome’s role in human health and disease. Humans have co-evolved with the microbiome and have become dependent on its biochemical output, such as certain vitamins and short-chain fatty acids [15,16]. The gut microbiome can also produce harmful biochemicals that have been implicated in various disease states [15]. To fully understand the relationships between the gut microbiome and human health status, biochemical functions of the microorganisms must be identified and quantified. Several next generation sequencing-based methods have been used for analyzing the gut microbiome, each with clear advantages and disadvantages. The simplest, least expensive, and most common method is 16S rRNA gene sequencing [17], which sequences a small portion of the highly conserved prokaryotic 16S ribosomal RNA gene [18]. This method can provide taxonomic resolution to the genus level [19,20], but does not measure the biochemical functions of the microorganisms [18] or distinguish living from dead organisms. In addition, traditional 16S rRNA sequencing excludes some bacteria, most archaea, and all eukaryotic organisms and viruses [21], resulting in a limited view of the gut microbiome ecosystem.

Metagenomic (shotgun DNA) sequencing provides strain-level resolution of all DNA-based microorganisms [18] (it does not detect RNA viruses or RNA bacteriophages). However, it can only identify the *potential* biochemical functions of the microbiome and can neither identify nor quantify the active biochemical pathways. This is a disadvantage for studying dysbiosis-related disease states, such as inflammatory bowel disease (IBD), which has been shown to have a disparity between metagenomic potential pathways and actual biochemical pathways expressed in disease and control populations [22].

Metatranscriptomic analysis (metatranscriptomics, RNA sequencing, RNAseq) offers insights into the biochemical activities of the gut microbiome by quantifying expression levels of active microbial genes, allowing for assessment of pathway activities, while also providing strain-level taxonomic resolution for all metabolically active organisms and viruses [23,24]. To date, metatranscriptomic analyses of stool samples have been limited due to the cost and complexity of both laboratory and bioinformatic methods [25]. By removing less informative rRNA, more valuable transcriptome data can be generated with less sequencing depth [26,24], resulting in reduced per-sample sequencing costs.

An automated technology has been developed for metatranscriptomic analysis of human clinical samples, called Viomega. In this study, Viomega was applied to 10,000 human stool samples to gain a better understanding of the strain-level taxonomies and microbial functions. Several small-scale studies were performed to quantify the metatranscriptomic stability in the lower colon and measure the intra-sample variability of metatranscriptomic analyses.

## MATERIALS AND METHODS

### Study participants, ethics, and sample collection and transportation

For this study, Viome used data from 10,000 participants. All study participants gave consent to being in the study, and all study procedures were approved by a federally accredited Institutional Review Board (IRB). Participants were recruited from any age, gender, and ethnic group.

Stool samples were collected using Viome’s Gut Intelligence kit by each study participant at their own residences. The kit included a sample collection tube with an integrated scoop, a proprietary RNA preservative, and sterile glass beads. A pea-sized stool sample was collected and placed inside the tube and vigorously shaken to homogenize the sample, exposing it to the RNA preservative. The sample was then shipped at room temperature using a common courier to Viome labs for analysis. Shipping times ranged from one to twelve days. Each participant completed a questionnaire with general lifestyle and health information.

### Metatranscriptomic analysis of stool samples

For the metatranscriptomic analysis of 10,000 stool samples, a proprietary sample-to-result automated platform called Viomega was used. Stool samples were lysed using bead-beating in a strong chemical denaturant and then placed on an automated liquid handler, which performed all downstream laboratory methods. Samples were processed in batches in a 96-well microplate; each batch consisted of ninety-four human stool samples, a negative process control (NPC, water), and a positive process control (PPC, custom synthetic RNA). RNA was extracted using a proprietary method. Briefly, silica-coated beads and a series of washes were used to purify RNA after lysis and RNA was eluted in water. DNA was degraded using RNase-free DNase.

The majority of prokaryotic ribosomal RNAs (rRNAs: 16S and 23S) were removed using a custom subtractive hybridization method. Biotinylated DNA probes with sequences complementary to rRNAs were added to total RNA, the mixture was heated and cooled, and the probe-rRNA complexes were removed using magnetic streptavidin beads. The remaining RNAs were converted to directional sequencing libraries with unique dual-barcoded adapters and ultra-pure reagents. Libraries were pooled and quality controlled with dsDNA Qubit (ThermoFisher) and Fragment Analyzer (Advanced Analytical). Library pools were sequenced on Illumina NextSeq or NovaSeq instruments using 300 cycle kits.

Viomega’s bioinformatics module operates on Amazon Web Services and includes quality control, taxonomic profiling and functional analysis. Quality control tools trim and filter the raw reads and quantify the amounts of sample-to-sample cross-talk and background contamination by microbial taxa. Viomega generates read-based taxonomy assignments using a multi-step process. The sequencing reads are aligned to a proprietary database of pre-computed genomic signatures at three taxonomic levels: strain, species, and genus. The unique signatures are computed from full-length genomes by removing short subsequences of a defined length, k, (k-mers) shared among more than one genome and keeping unique k-mers that make up the signature [27]. The Viomega taxonomy database was generated from a large RefSeq database containing more than 110,000 microbial genomes. After the initial taxonomic assignments were generated, potential false-positives were removed using an auto-blast algorithm that uses an even larger database of organisms.

Identity and relative activity of microbial genes and enzymatic functions in the stool samples were assessed using a proprietary algorithm. At a high level, this involves a multi-tiered approach to align the sample reads to the integrated gene catalog (IGC) [28] library of genes to first identify and then quantify the genes in the sample. Informative genes (i.e. non-rRNA) were quantified in units of transcripts per million (TPM) to allow for cross-sample comparisons. Using the Kyoto Encyclopedia of Genes and Genomes (KEGG) [29] annotation mapping of IGC genes to KEGG orthologies (KOs), the enzymic functions and activity were quantified in these samples as the aggregate TPM. The KEGG mapping also allows for functional modules and pathway analysis.

### Small-scale studies

For the validation of sample lysis in the Viomega pipeline the following organisms were grown in nutrient broth at 37°C and 450 rpm in a VWR incubating mini shaker: *Bacillus subtilis* Marburg strain (ATCC 6051-U), *Corynebacterium stationis* strain NCTC 2399 (ATCC 6872), *Citrobacter freundii* strain ATCC 13316, NCTC 9750 (ATCC 8090), and *Serratia liquefaciens* strain CDC 1284-57 ATCC 12926 (ATCC 27592). In addition, the following organisms were grown in yeast mold broth at 37°C and 450 rpm in a VWR incubating mini shaker: *Saccharomyces cerevisiae* strain S288C (ATCC 204508) and *Candida dubliniensis* strain CBS 7987 (ATCC MYA-646).

To illustrate the accuracy of taxonomic classification at the species level of the Viomega technology the 10 Strain Even Mix Whole Cell Material (ATCC^®^ MSA-2003™) product was utilized. As stated by the manufacturer, this product is comprised of an even mixture of the following organisms: *Bacillus cereus* (ATCC 10987), *Bifidobacterium adolescentis* (ATCC 15703), *Clostridium beijerinckii* (ATCC 35702), *Deinococcus radiodurans* (ATCC BAA816), *Enterococcus faecalis* (ATCC 47077), *Escherichia coli* (ATCC 700926), *Lactobacillus gasseri* (ATCC 33323), *Rhodobacter sphaeroides* (ATCC 17029), *Staphylococcus epidermidis* (ATCC 12228), and *Streptococcus mutans* (ATCC 700610).

## RESULTS AND DISCUSSION

### Validation of Viomega technology Sample lysis

Uneven sample lysis can introduce major errors in any method since sample composition can vary widely in terms of easy-to-lyse microorganisms (viruses and Gram(-) bacteria) and difficult to very-difficult-to-lyse Gram(+) bacteria and yeast. Viomega utilizes a combination of chemical (denaturant) and physical (bead beating) sample lysis, which has been shown to have the best efficiency. To test this method, two strains of Gram(−), two strains of Gram(+), and two strains of yeast were grown to an optical density range of 0.4-0.8 AU. Equal amounts of each organism in triplicate then underwent chemical and physical sample lysis and total RNA was extracted from each sample. RNA yields obtained were consistent and show no bias against Gram(+) bacteria or yeast (average yield: G(−) = 93.3 ng/*µ*L, 83.6 ng/*µ*L; G(+) = 131.9 ng/*µ*L, 152.3 ng/*µ*L; yeast = 113.1 ng/*µ*L, 100.8 ng/*µ*L) (Figure 1). The process was also reproducible, with very small variability across technical replicates (standard deviation range = 3.4 - 11.0 ng/*µ*L).

**Figure 1.**
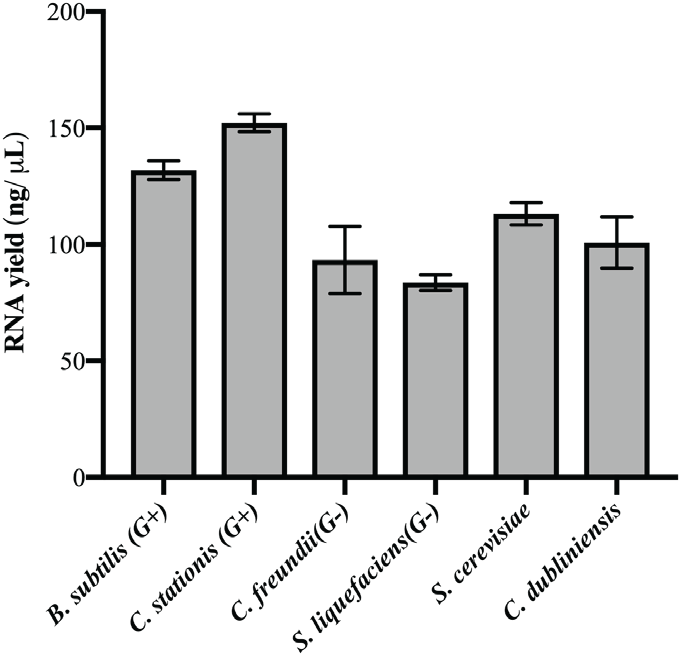
Average RNA yields (ng/*μ*L) of model organisms after sample lysis and RNA extraction (*B*. *subtilis*: 131.9 +/− 4.0 ng/μL; *C. stationis*: 152.3 +/− 3.9 ng/*μ*L; *C. fruendii*: 93.3 +/− 14.4 ng/*μ*L; *S. liquefaciens*: 83.6 +/− 3.4 ng/*μ*L; *S. cerevisiae*: 113.1 +/− 4.8 ng/*μ*L; *C. dubliniensis*: 100.8 +/− 11.0 ng/*μ*L; n = 3)

### Ambient temperature sample transportation

A noted shortcoming of metatranscriptomics is that it analyzes labile RNA molecules. This is most apparent in the case of dead organisms, as the existing RNA is rapidly degraded, while no new transcripts are made. In living organisms, however, RNA is continuously made and degraded. By exposing living organisms to appropriate reagents, this dynamic equilibrium of gene expression can be “frozen in time” at the time of sample collection and quantitatively analyzed later. To achieve this, Viomega uses a chemical denaturant/RNA stabilizing solution that ensures the preservation of RNA integrity during sample transport at ambient temperatures. Fourteen aliquots were made from a single donor sample; four aliquots were processed using Viomega immediately, while three samples were stored at room temperature (RT) for four weeks prior to processing. Seven aliquots were shipped through a standard courier and held on-site at the laboratory for a total time of four weeks prior to processing. All comparisons show very strong correlation with a Spearman correlation value of 0.8 or greater (Figure 2) [30]. No difference was found in taxonomic profiling or functional composition between time to processing or shipping conditions prior to processing (Figure 2).

**Figure 2.**
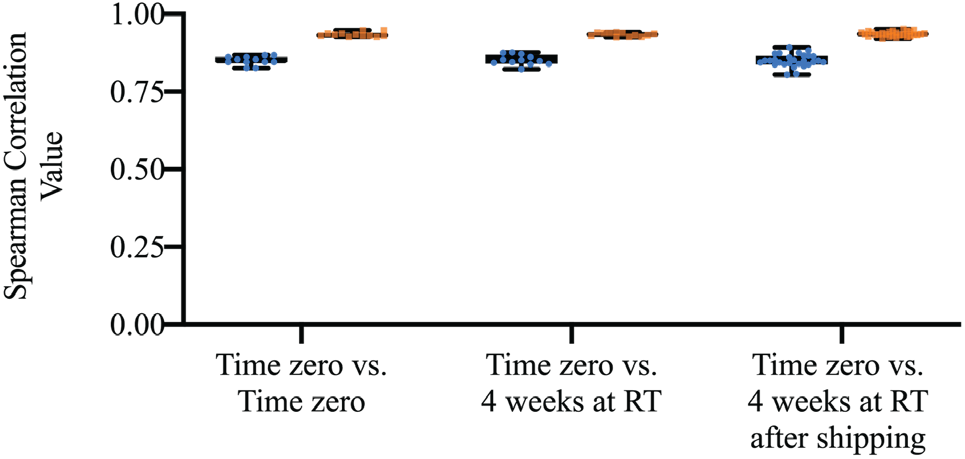
Spearman Correlation of taxonomic data (blue) and functional data (orange) between and within storage conditions. Median correlation of taxonomic data: T_0_ vs T_0_ = 0.8493; T_0_ vs T_4weeksRT_ = 0.8503; T_0_ vs T_4weeksSHIP_ = 0.8493. Median correlation values of functional data: T_0_ vs T_0_ = 0.932; T_0_ vs T_4weeksRT_ = 0.9341; T_0_ vs T_4weeksSHIP_ = 0.9349.

### Sample-to-sample cross-talk (STSC)

STSC (also known as barcode switching, barcode hopping, or read misassignment) can cause significant errors when sequencing many samples on a single sequencing run. Standard library preparation methods for sample barcoding have high error rates, from 0.2-5%, especially on the newest generation of Illumina platforms that use ExAmp technology [31,32]. This phenomenon can cause errors in the reported taxonomies, *e.g.* abundant taxa in a sequencing run being assigned to samples in which those taxa did not exist. Viomega minimizes STSC by a combination of specially produced barcode oligos, dual unique barcode sequences of 11 bps each, and only reporting the taxa that did not exceed the rate of measured STSC on each batch of 96 samples. STSC was quantified by introducing a synthetic, non-natural RNA sample (PPC) in each microplate and measuring its quantity in each of the other samples. The PPC sample was randomly positioned in each plate. STSC in Viomega mostly falls under 1 read per million reads (0.0001%) (Figure 3) on NovaSeq platform (S1 flow cell, 300 cycle kits), which is more than 1,000-fold lower than in data obtained using commercial library preparation kits [31–33].

**Figure 3.**
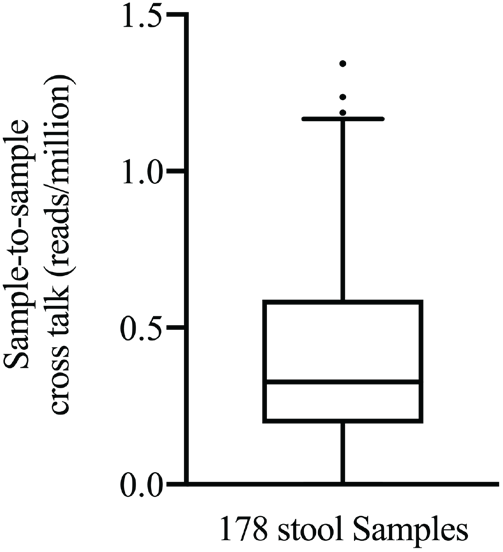
Sample-to-sample cross-talk (reads per million) for 178 stool samples sequenced on the Illumina NovaSeq platform using Viomega dual unique barcode sequences. Sample-to-sample cross-talk determined by measuring the occurrence of non-natural PPC reads in each stool sample. Range = 0 - 1.34 reads per million; median = 0.33 reads per million.

### Background contamination of samples

Since any metagenomic or metatranscriptomic analysis identifies all taxa in a sample, nucleic acid contamination of the reagents (especially purification kits and enzymes), instruments, and poor laboratory practices can lead to the inclusion of contaminating taxa into scientific results [34–36]. To minimize this, Viomega uses ultra-pure reagents, good laboratory practices, and fully automated liquid handling systems. Every plate of ninety-six samples contained a positive process control (PPC) sample, which is a synthetic RNA that was subjected to the same process as the rest of the samples (from kit manufacturing to bioinformatics). This sample was sequenced and analyzed like all other samples on the plate, allowing any microbial contamination to be detected. Over the course of twenty consecutive batches (1,880 stool samples, 20 PPC samples) the level of back-ground contamination observed in PPC samples was extraordinarily low, with an average of 1.4 contaminating reads (std dev. 2.6; n=20) out of 5-15 million sequencing reads and 0.3 contaminating taxa (std. dev. 0.5; n = 20). Across twenty batches, the number of contaminating taxa was either zero or one, with a maximum of ten sequencing reads (out of an average of ~10 million) assigned to the taxon. These values were below the threshold for reporting any microorganism from the Viomega analysis and therefore do not cause any false positives to the results.

### Depletion of ribosomal RNAs

The vast majority of RNA molecules in any biological sample are ribosomal RNAs (rRNA). Approximately 96% of all reads from stool samples align to microbial rRNAs (Table S1), leaving only ~4% of sequencing data aligning to microbial messenger RNAs. Since rRNA sequences are not very informative (housekeeping functions, poor taxonomic resolution), and a key goal of Viomega is to deeply probe the functional landscape of the gut microbiome (*i.e.* quantify the messenger RNAs), a subtractive hybridization method for rRNA depletion has been implemented in the Viomega process. This fully automated method reduces rRNA to 60.4 +/- 14.9% (n = 90), thus providing an average enrichment of microbial messenger RNAs of ~10 fold by increasing sequencing data aligning to microbial messengers RNAs to ~40%.

### Viomega: Accuracy of taxonomic classification

Given the large amounts of metatranscriptomic data obtained from each sample (over one giga-base pairs) and a very large database (more than 110,000 genomes), it is extremely challenging to have a high throughput, fully automated, cost-effective, cloud-based, and highly accurate bioinformatic pipeline. Viomega is a fully automated cloud application whose efficiency comes from using a pre-computed database of microbial signatures. This approach reduces the amount of searchable sequence space by roughly two orders of magnitude and largely eliminates false positive results. Viomega technology was used to analyze a commercially available mock community (10 Strain Even Mix Whole Cell Material, ATCC® MSA-2003™). For identification at the species level, Viomega shows 100% accuracy consistent with the mock community, with no false positive or false negative calls (Figure 4). The whole cell relative abundance of these microorganisms was reported by the supplier as identical. However, the relative RNA amounts (measured as relative activity by Viomega) may not be the same due to potential differences in how each monoculture was grown, processed, and stored prior to the preparation of the mock community. It is also possible that *Deinococcus radiodurans* contains more RNA per cell than other bacteria, due to its diploid genome. The somewhat higher relative abundance cannot be explained with facile lysis (it is a Gram-positive organism) or low GC content (67%).

**Figure 4.**
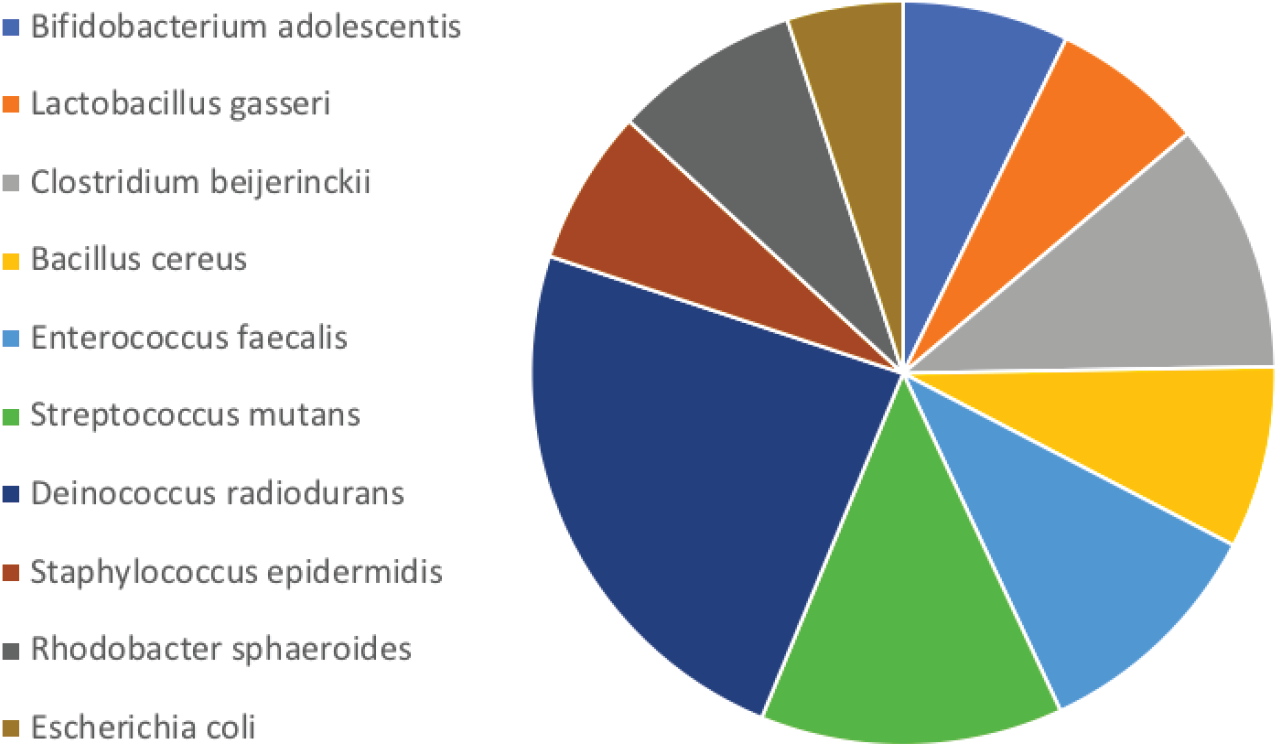
Accuracy and relative abundance of Viomega analysis on a commercial mock community. Species contents of the mock community as listed by the supplier are shown (left); supplier lists relative abundance of whole cells as equal among all species. Viomega achieves 100% accuracy at species level identification.

### Findings from the Viomega taxonomic classification

Using the Viomega taxonomy classification pipeline, a total of 2,723 microbial strains, 1,946 microbial species, and 528 microbial genera have been identified in 10,000 human stool samples. The identified microorganisms include bacteria, archaea, viruses, bacteriophages, and eukaryotes (Table 1).

**Table 1.** Top ten strains, species, and genera identified in 10,000 human stool samples, based on their prevalence. See supplementary materials for all taxa identified in 10,000 human stool samples (Table S2 for strain, S3 for species, S4 for genera).

| TOP 10 GENERA |  | TOP 10 SPECIES |  | TOP 10 STRAINS |  |
| --- | --- | --- | --- | --- | --- |
| Genus | Prevalence, % | Species | Prevalence, % | Strains | Prevalence, % |
| Clostridium | 99.7 | Bacteroides vulgatus | 97.1 | Eggerthella lenta 1_1_60AFAA | 97.1 |
| Bacteroides | 99.6 | Acinetobacter baumannii | 96.8 | [Eubacterium] hallii DSM 3353 | 93.9 |
| Blautia | 97.6 | Faecalibacterium prausnitzii | 96.4 | Veillonella dispar ATCC 17748 | 92.3 |
| Acinetobacter | 97.5 | Bacteroides uniformis | 95.4 | Anaerotruncus colihominis DSM 17241 | 91.4 |
| Eubacterium | 97.2 | Eggerthella lenta | 91.8 | Clostridium phoceensis strain GD3 | 90.8 |
| Parabacteroides | 96.9 | [Eubacterium] hallii | 91.5 | Blautia obeum ATCC 29174 | 89.7 |
| Lactococcus | 96.9 | Anaerotruncus colihominis | 91.4 | [Eubacterium] eligens strain 2789STDY5834875 | 88.9 |
| Faecalibacterium | 96.4 | Clostridium phoceensis | 90.8 | Faecalibacterium cf. prausnitzii KLE1255 | 88.3 |
| Roseburia | 95.7 | Veillonella dispar | 89.0 | Faecalibacterium prausnitzii A2-165 | 86.0 |
| Alistipes | 95.1 | Fusicatenibacter saccharivorans | 87.3 | Roseburia hominis A2-183 | 83.4 |

### Viomega: Quantification of microbial biochemical functions

Using Viomega’s functional analysis tools, the expression of >100,000 microbial open reading frames (ORFs), which were grouped into 6,879 KEGG functions, were identified and quantified from 10,000 human stool samples. Top ten KEGG functions are shown in Table 2.

**Table 2.** Top ten KEGG functions identified in 10,000 human stool samples. See supplementary material for top 100 KEGG functions (Table S5).

| KO ID | Name | 100.00 |
| --- | --- | --- |
| K00936 | pdtaS | 100.00 |
| K01190 | lacZ | 99.99 |
| K03046 | rpoC | 99.99 |
| K02355 | fusA, GFM, EFG | 99.99 |
| K00540 | fqr | 99.99 |
| K03695 | clpB | 99.99 |
| K03296 | TC.HAE1 | 99.99 |
| K01362 | OVCH | 99.98 |
| K02358 | tuf, TUFM | 99.98 |
| K06950 | K06950 | 99.98 |

### Intra-sample variability of metatranscriptomic analyses

Because large scale studies would preferably analyze a single stool sample (instead of an average of multiples), it was important to understand the variability of microbial taxonomy and functions across individual stool samples. To understand this variability, three volunteers (P11, P12, and P13) collected samples from three parts of their stool samples: 1) one end, 2) the opposite end, and 3) the middle. Each biological sample was split into three technical replicates (a, b, and c). All samples were analyzed using Viomega, followed by unsupervised clustering analysis (Kendall’s correlation). All biological and technical replicates from the same stool sample (in-group) clustered by participant with very high similarity, and were different from the outgroup samples, especially at the strain level taxonomy (Figure 5). This mini-study shows high uniformity of metatranscriptomic data across stool samples. While there have been claims of large intra-sample variability [37], these were likely based on biased methods, and not real differences in microbial taxonomy [38]. For large scale studies, it is cost-prohibitive to collect and analyze multiple samples per collection time; Viomega metatranscriptomic analysis provides reproducible results across stool samples.

**Figure 5.**
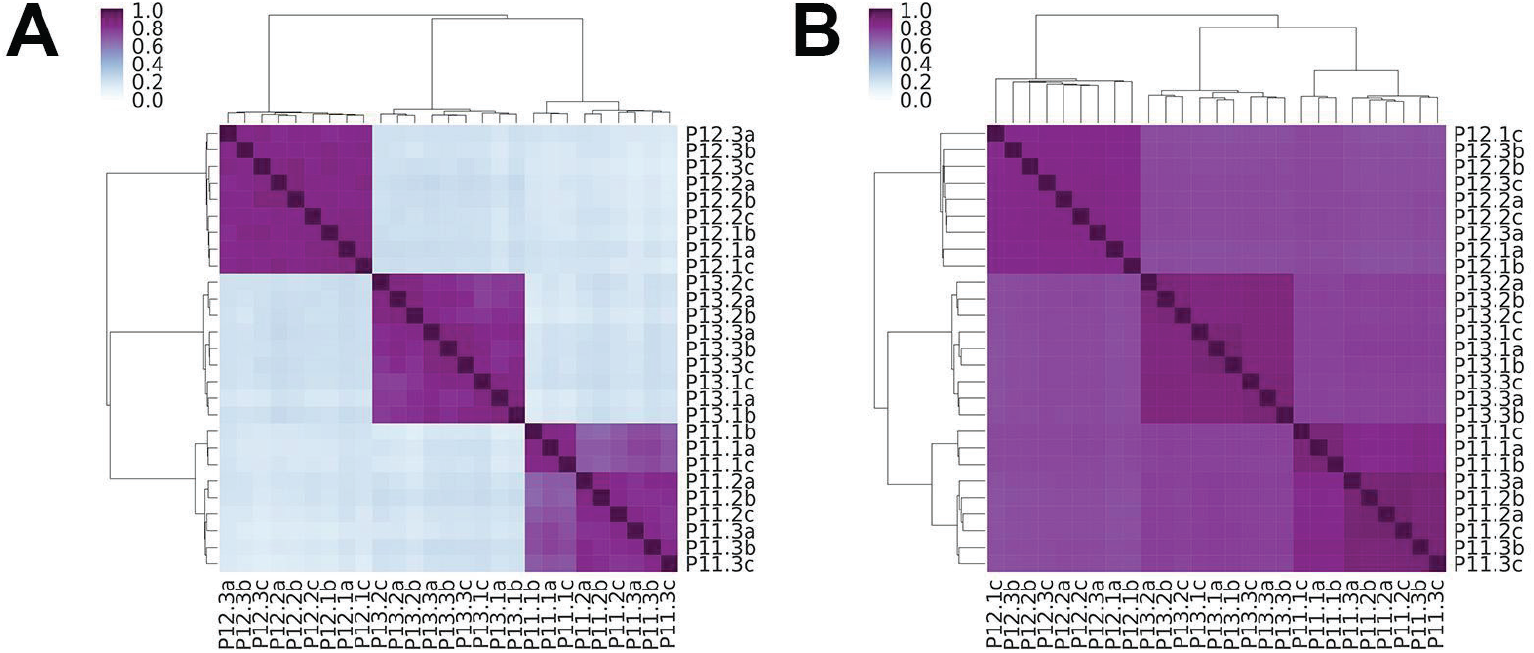
Intra-sample variability of microbial taxonomy and functions using Viomega analysis. Three participants (P11, P12, and P13) provided three stool samples each, from three different parts of the stool (1, 2, and 3). Each biological sample was analyzed as three technical replicates (a, b, and c). Following Viomega analysis, unsupervised clustering analysis (Kendall’s correlation) was performed on microbial taxonomy (at the strain level) (A) and biochemical functions at KEGG level (B).

### Short term (minutes) stability of stool metatranscriptomes

To identify any changes in the measured microbial taxonomy and functions in the first few minutes after a stool sample was produced, three participants (P12, P13, and P14) were asked to collect samples from the same stool (a) immediately, (b) three minutes later, and (c) ten minutes later. Unsupervised clustering analysis (Kendall’s correlation) was performed on the nine samples, and all samples clustered with high similarity based on the sample, and not the time of collection (Figure 6).

**Figure 6.**
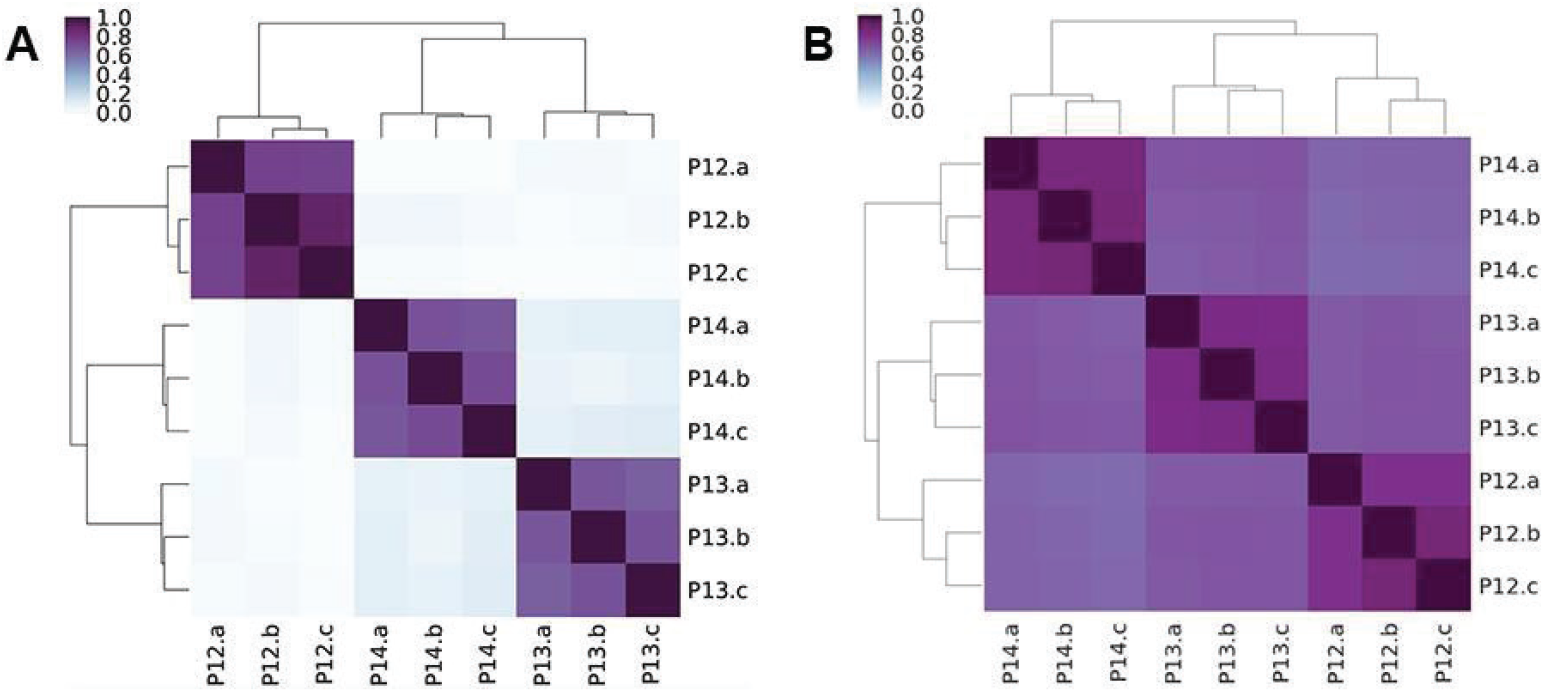
Stool samples collected immediately and at three and ten minutes (a, b, and c, respectively) after the stool was produced by three participants (P12, P13, and P14). Unsupervised clustering analysis (Kendall’s correlation) of samples shows a high similarity of strain level taxonomy (A) and KEGG-based microbial functions (B) based on the donor, and not based on the time of collection.

### Long-term (weeks) stability of stool metatranscriptomes

Because gene expression can change rapidly due to environmental changes, a mini-study was performed to look for metatranscriptome changes in stool microbiome over time. Seven volunteers were asked to maintain their normal diet and lifestyle for two weeks. During this period, three stool samples were collected from each participant: time zero, one week later, and two weeks later. The twenty-one samples were analyzed with Viomega and unsupervised clustering analysis (Kendall’s correlation) was performed based on taxonomy and KEGG functions (Figure 7). For both taxonomy (panel A) and KEGG functions (panel B), the samples clustered by the participant, confirming that both gut microbiome composition and biochemical functions were stable over the course of the study while maintaining a consistent diet. These data clearly demonstrate the utility of Viomega technology, as the microbial metatranscriptome was maintained with a consistent diet over a period of weeks.

**Figure 7.**
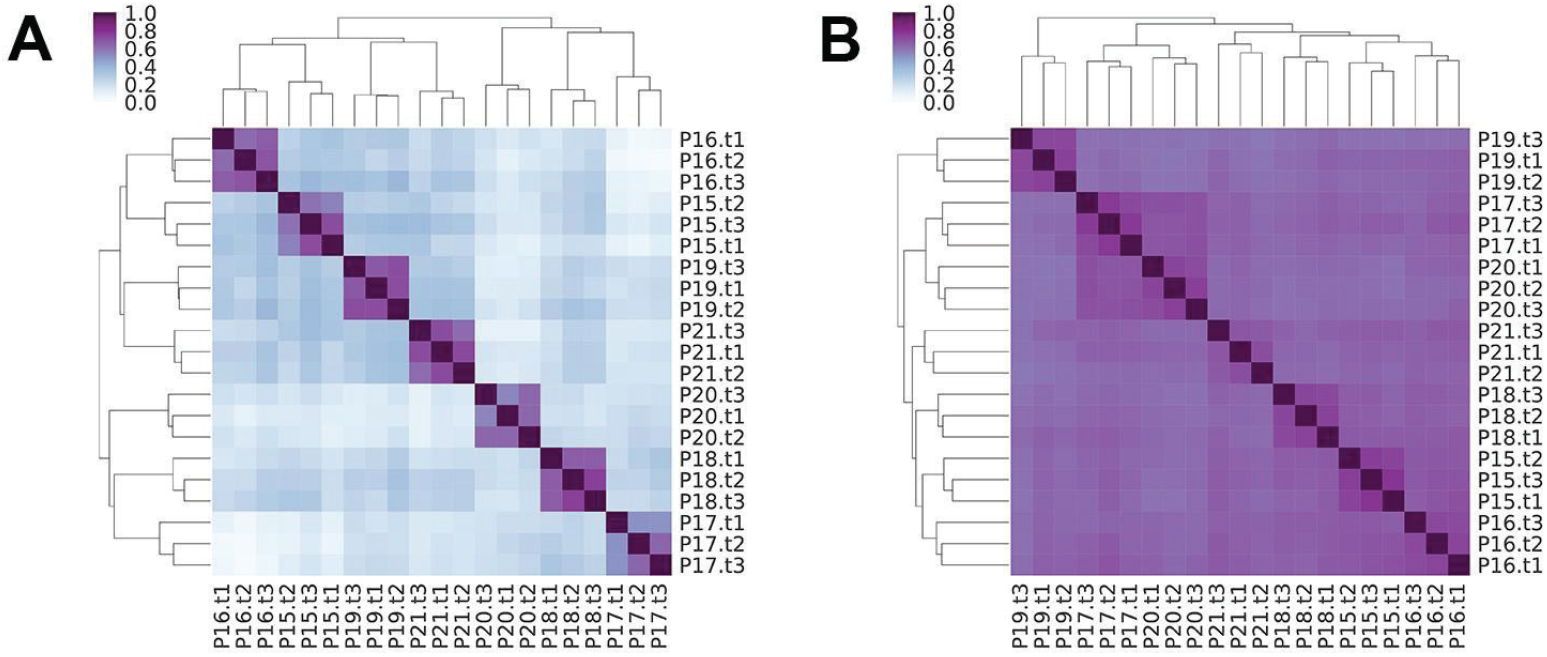
Unsupervised clustering analysis (Kendall’s correlation) of gut microbiome samples collected from seven participants (P15 - P21) at three time points two weeks apart (week 0, 1, and 2), while not changing the diet. Panel A shows a high degree of clustering of microbial taxonomy (at the strain level) by person, longitudinally, with high in-group similarity. Panel B shows a high degree of clustering of microbial functions (KEGG) by person, longitudinally, with high in-group similarity.

## CONCLUSIONS

In this report, Viomega, a sample-to-result, automated and robust stool metatranscriptomic analysis technology is described. Viomega includes at-home sample collection, stability at ambient temperatures during transport (for up to twenty-eight days), complete sample lysis, RNA extraction, physical removal of non-informative (ribosomal) RNAs, sequencing library preparation, Illumina sequencing, and a quantitative bioinformatic analysis platform that includes taxonomic classification and functional analysis. Almost all laboratory steps are performed in a 96-well format using automated liquid handlers. All bioinformatic analyses are automatically performed on cloud servers. Viomega includes several critically important quality control steps, both per-sample (number of base pairs generated for microbial messenger RNAs, % rRNA, and sample-to-sample cross-talk) and per-batch of ninety-six samples (background contamination, process control samples, RNA yields, etc.).

Using a commercial mock community, Viomega shows 100% accuracy (no false positives or negatives) at the species level. Since the ground truth for RNA content of each member of the mock community cannot be obtained from the manufacturer, it is unclear whether the small differences in the relative abundance of the ten microorganisms provided are an artifact of the sample or the method of producing the mock community.

Viomega was applied to 10,000 human stool samples and identified several thousand taxa at the strain, species, and genus ranks. More than 100,000 open reading frames (ORFs) were identified, quantified, and grouped into thousands of KEGG functions. The large bioinformatic data outputs of Viomega are being used to learn how gut microbiome taxonomy and functions are affected by the diet, develop improved models of how to precisely control the gut microbiome using diet, and how the gut microbiome correlates with human health and diseases. These analyses will be described in upcoming publications. While Viomega was specifically designed for stool sample analysis, modifications may be made for alternative pipelines for other types of human clinical applications in the future.

Viomega was used to perform several small-scale studies to demonstrate the robustness of stool metatranscriptomic analysis when the methods introduce minimal biases. These studies show that it is possible to collect a single stool sample as representative of the entire colonic microbiome. The studies also establish that the gut metatranscriptome exhibits weeks-long stability without a diet change both compositionally and functionally. It should be noted that all participants involved in the above mini-studies were self-reported healthy individuals. In each study, clustering by taxonomy showed much lower out-group similarity than clustering by function. While taxonomy has been shown to vary from person to person among healthy individuals [39], the observed clustering patterns (Figures 5, 6, 7) suggest similar functionality between healthy individuals; although different organisms are present, they are performing similar biochemical functions.

Viomega is a robust technology that offers a rapid and comprehensive taxonomic *and* functional readout of the gut microbiome. In addition, the cost to process a human stool sample through the Viomega pipeline ($199 for the Viome Gut Intelligence™ Test at the time of submission) is inexpensive compared to similar services ($15,000 for up to five samples through The Human Microbiome Project -“What are they actually doing” service) [40] largely due to batched processing, removal of rRNAs, and the unique Viomega taxonomy database. This technology will increase overall understanding of the interplay among diet, gut microbiome, and human health, and is enabling gut microbiome-based personalized nutrition as an emerging field. These advances may fuel mitigation and treatment for a variety of human health conditions, such as cardiovascular disease, obesity, autoimmune disease, ASD, and Parkinson’s disease.

## Conflicts of Interest

All authors are also employees of Viome, Inc, which sponsored the study.

## Funding Statement

Viome, Inc. funded these studies.

## Supplementary Materials

Supplementary Table S1 has data for the percent ribosomal RNA in stool samples that are processed through Viomega without the custom rRNA depletion method. Supplementary Tables S2-S4 show all taxa identified by Viomega in 10,000 human stool samples (strains, species, and genera, respectively). Supplementary Table S5 shows the top 100 KEGG functions identified by Viomega in 10,000 human stool samples.

## References

[1] Nasca C, Bigio B, Lee FS, Young SP, Kautz MM, Albright A, Beasley J, Millington DS, Mathé AA, Kocsis JH, Murrough JW, McEwen BS, Rasgon N. 2018. Acetyl-l-carnitine deficiency in patients with major depressive disorder. Proceedings of the National Academy of Sciences 115:8627–8632. DOI: 10.1073/pnas.1801609115.

[2] Flint HJ. 2016. Gut microbial metabolites in health and disease. Gut Microbes 7:187–188. DOI: 10.1080/19490976.2016.1182295.

[3] Postler TS, Ghosh S. 2017. Understanding the Holobiont: How Microbial Metabolites Affect Human Health and Shape the Immune System. Cell Metabolism 26:110–130. DOI: 10.1016/j.cmet.2017.05.008.

[4] Clemente JC, Ursell LK, Parfrey LW, Knight R. 2012. The Impact of the Gut Microbiota on Human Health: An Integrative View. Cell 148:1258–1270. DOI: 10.1016/j.cell.2012.01.035.

[5] Li J, Zhao F, Wang Y, Chen J, Tao J, Tian G, Wu S, Liu W, Cui Q, Geng B, Zhang W, Weldon R, Auguste K, Yang L, Liu X, Chen L, Yang X, Zhu B, Cai J. 2017. Gut microbiota dysbiosis contributes to the development of hypertension. Microbiome 5. DOI: 10.1186/s40168-016-0222-x.

[6] Parekh PJ, Balart LA, Johnson DA. 2015. The Influence of the Gut Microbiome on Obesity, Metabolic Syndrome and Gastrointestinal Disease: Clinical and Translational Gastroenterology 6:e91. DOI: 10.1038/ctg.2015.16.

[7] Zhang Y, Zhang H. 2013. Microbiota associated with type 2 diabetes and its related complications. Food Science and Human Wellness 2:167–172. DOI: 10.1016/j.fshw.2013.09.002.

[8] Karlsson FH, Fåk F, Nookaew I, Tremaroli V, Fagerberg B, Petranovic D, Bäckhed F, Nielsen J. 2012. Symptomatic atherosclerosis is associated with an altered gut metagenome. Nature Communications 3. DOI: 10.1038/ncomms2266.

[9] Chervonsky AV. 2013. Microbiota and Autoimmunity. Cold Spring Harbor Perspectives in Biology 5:a007294–a007294. DOI: 10.1101/cshperspect.a007294.

[10] Mayer EA, Tillisch K, Gupta A. 2015. Gut/brain axis and the microbiota. Journal of Clinical Investigation 125:926–938. DOI: 10.1172/JCI76304.

[11] Wong H, Hoeffer C. 2018. Maternal IL-17A in autism. Experimental Neurology 299:228–240. DOI: 10.1016/j.expneurol.2017.04.010.

[12] Yano JM, Yu K, Donaldson GP, Shastri GG, Ann P, Ma L, Nagler CR, Ismagilov RF, Mazmanian SK, Hsiao EY. 2015. Indigenous Bacteria from the Gut Microbiota Regulate Host Serotonin Biosynthesis. Cell 161:264–276. DOI: 10.1016/j.cell.2015.02.047.

[13] Harach T, Marungruang N, Duthilleul N, Cheatham V, Mc Coy KD, Frisoni G, Neher JJ, Fåk F, Jucker M, Lasser T, Bolmont T. 2017. Reduction of Abeta amyloid pathology in APPPS1 transgenic mice in the absence of gut microbiota. Scientific Reports 7. DOI: 10.1038/srep41802.

[14] Scheperjans F, Aho V, Pereira PAB, Koskinen K, Paulin L, Pekkonen E, Haapaniemi E, Kaakkola S, Eerola Rautio J, Pohja M, Kinnunen E, Murros K, Auvinen P. 2015. Gut microbiota are related to Parkinson’s disease and clinical phenotype. Movement Disorders 30:350–358. DOI: 10.1002/mds.26069.

[15] Wang B, Yao M, Lv L, Ling Z, Li L. 2017. The Human Microbiota in Health and Disease. Engineering 3:71–82. DOI: 10.1016/J.ENG.2017.01.008.

[16] Lloyd-Price J, Abu-Ali G, Huttenhower C. 2016. The healthy human microbiome. Genome Medicine 8. DOI: 10.1186/s13073-016-0307-y.

[17] Woo PCY, Lau SKP, Teng JLL, Tse H, Yuen K-Y. 2008. Then and now: use of 16S rDNA gene sequencing for bacterial identification and discovery of novel bacteria in clinical microbiology laboratories. Clinical Microbiology and Infection 14:908–934. DOI: 10.1111/j.1469-0691.2008.02070.x.

[18] Langille MGI, Zaneveld J, Caporaso JG, McDonald D, Knights D, Reyes JA, Clemente JC, Burkepile DE, Vega Thurber RL, Knight R, Beiko RG, Huttenhower C. 2013. Predictive functional profiling of microbial communities using 16S rRNA marker gene sequences. Nature Biotechnology 31:814–821. DOI: 10.1038/nbt.2676.

[19] Knight R, Vrbanac A, Taylor BC, Aksenov A, Callewaert C, Debelius J, Gonzalez A, Kosciolek T, McCall L-I, McDonald D, Melnik AV, Morton JT, Navas J, Quinn RA, Sanders JG, Swafford AD, Thompson LR, Tripathi A, Xu ZZ, Zaneveld JR, Zhu Q, Caporaso JG, Dorrestein PC. 2018. Best practices for analysing microbiomes. Nature Reviews Microbiology 16:410–422. DOI: 10.1038/s41579-018-0029-9.

[20] Poretsky R, Rodriguez-R LM, Luo C, Tsementzi D, Konstantinidis KT. 2014. Strengths and Limitations of 16S rRNA Gene Amplicon Sequencing in Revealing Temporal Microbial Community Dynamics. PLoS ONE 9:e93827. DOI: 10.1371/journal.pone.0093827.

[21] Raymann K, Moeller AH, Goodman AL, Ochman H. 2017. Unexplored Archaeal Diversity in the Great Ape Gut Microbiome. mSphere 2:e00026–17. DOI: 10.1128/mSphere.00026-17.

[22] Schirmer M, Franzosa EA, Lloyd-Price J, McIver LJ, Schwager R, Poon TW, Ananthakrishnan AN, Andrews E, Barron G, Lake K, Prasad M, Sauk J, Stevens B, Wilson RG, Braun J, Denson LA, Kugathasan S, McGovern DPB, Vlamakis H, Xavier RJ, Huttenhower C. 2018. Dynamics of metatranscription in the inflammatory bowel disease gut microbiome. Nature Microbiology 3:337–346. DOI: 10.1038/s41564-017-0089-z.

[23] Gosalbes MJ, Durbán A, Pignatelli M, Abellan JJ, Jiménez-Hernández N, Pérez-Cobas AE, Latorre A, Moya A. 2011. Metatranscriptomic Approach to Analyze the Functional Human Gut Microbiota. PLoS ONE 6:e17447. DOI: 10.1371/journal.pone.0017447.

[24] Bashiardes S, Zilberman-Schapira G, Elinav E. 2016. Use of Metatranscriptomics in Microbiome Research. Bioinformatics and Biology Insights 10:BBI.S34610. DOI: 10.4137/BBI.S34610.

[25] Knight R, Jansson J, Field D, Fierer N, Desai N, Fuhrman JA, Hugenholtz P, van der Lelie D, Meyer F, Stevens R, Bailey MJ, Gordon JI. Kowalchuk GA, Gilbert JA. 2012. Unlocking the potential of metagenomics through replicated experimental design. Nature Biotechnology 30:513–520. DOI: 10.1038/nbt.2235.

[26] He S, Wurtzel O, Singh K, Froula JL, Yilmaz S, Tringe SG, Wang Z, Chen F, Lindquist EA, Sorek R, Hugenholtz P. 2010. Validation of two ribosomal RNA removal methods for microbial metatranscriptomics. Nature Methods 7:807–812. DOI: 10.1038/nmeth.1507.

[27] Freitas TAK, Li P-E, Scholz MB, Chain PSG. 2015. Accurate read-based metagenome characterization using a hierarchical suite of unique signatures. Nucleic Acids Research 43:e69. DOI: 10.1093/nar/gkv180.

[28] MetaHIT Consortium, Li J, Jia H, Cai X, Zhong H, Feng Q, Sunagawa S, Arumugam M, Kultima JR, Prifti E, Nielsen T, Juncker AS, Manichanh C, Chen B, Zhang W, Levenez F, Wang J, Xu X, Xiao L, Liang S, Zhang D, Zhang Z, Chen W, Zhao H, Al-Aama JY, Edris S, Yang H, Wang J, Hansen T, Nielsen HB, Brunak S, Kristiansen K, Guarner F, Pedersen O, Doré J, Ehrlich SD, Bork P, Wang J. 2014. An integrated catalog of reference genes in the human gut microbiome. Nature Biotechnology 32:834–841. DOI: 10.1038/nbt.2942.

[29] Kanehisa M, Sato Y, Kawashima M, Furumichi M, Tanabe M. 2016. KEGG as a reference resource for gene and protein annotation. Nucleic Acids Research 44:D457–D462. DOI: 10.1093/nar/gkv1070.

[30] Akoglu H. 2018. User’s guide to correlation coefficients. Turkish Journal of Emergency Medicine 18:91–93. DOI: 10.1016/j.tjem.2018.08.001.

[31] D’Amore R, Ijaz UZ, Schirmer M, Kenny JG, Gregory R, Darby AC, Shakya M, Podar M, Quince C, Hall N. 2016. A comprehensive benchmarking study of protocols and sequencing platforms for 16S rRNA community profiling. BMC Genomics 17. DOI: 10.1186/s12864-015-2194-9.

[32] Sinha R, Stanley G, Gulati GS, Ezran C, Travaglini KJ, Wei E, Chan CKF, Nabhan AN, Su T, Morganti RM, Conley SD, Chaib H, Red-Horse K, Longaker MT, Snyder MP, Krasnow MA, Weissman IL. 2017. Index Switching Causes “Spreading-Of-Signal” Among Multiplexed Samples In Illumina HiSeq 4000 DNA Sequencing. bioRxiv:125724. DOI: 10.1101/125724.

[33] Illumina. 2017. Effects of index misassignment on multiplexing and downstream analysis.

[34] Salter SJ, Cox MJ, Turek EM, Calus ST, Cookson WO, Moffatt MF, Turner P, Parkhill J, Loman NJ, Walker AW. 2014. Reagent and laboratory contamination can critically impact sequence-based microbiome analyses. BMC Biology 12. DOI: 10.1186/s12915-014-0087-z.

[35] Glassing A, Dowd SE, Galandiuk S, Davis B, Chiodini RJ. 2016. Inherent bacterial DNA contamination of extraction and sequencing reagents may affect interpretation of microbiota in low bacterial biomass samples. Gut Pathogens 8. DOI: 10.1186/s13099-016-0103-7.

[36] Lusk RW. 2014. Diverse and Widespread Contamination Evident in the Unmapped Depths of High Throughput Sequencing Data. PLOS ONE 9:e110808. DOI: 10.1371/journal.pone.0110808.

[37] Eckburg PB, Bik EM, Bernstein CN, Purdom E, Dethlefsen L, Sargent M, Gill SR, Nelson KE, Relman DA. 2005. Diversity of the Human Intestinal Microbial Flora. Science 308:1635–1638. DOI: 10.1126/science.1110591.

[38] Wu GD, Lewis JD, Hoffmann C, Chen Y-Y, Knight R, Bittinger K, Hwang J, Chen J, Berkowsky R, Nessel L, Li H, Bushman FD. 2010. Sampling and pyrosequencing methods for characterizing bacterial communities in the human gut using 16S sequence tags. BMC Microbiology 10:206. DOI: 10.1186/1471-2180-10-206.

[39] The Human Microbiome Project Consortium, Huttenhower C, Gevers D, Knight R, Abubucker S, Badger JH, Chinwalla AT, Creasy HH, Earl AM, FitzGerald MG, Fulton RS, Giglio MG, Hallsworth-Pepin K, Lobos EA, Madupu R, Magrini V, Martin JC, Mitreva M, Muzny DM, Sodergren EJ, Versalovic J, Wollam AM, Worley KC, Wortman JR, Young SK, Zeng Q, Aagaard KM, Abolude OO, Allen-Vercoe E, Alm EJ, Alvarado L, Andersen GL, Anderson S, Appelbaum E, Arachchi HM, Armitage G, Arze CA, Ayvaz T, Baker CC, Begg L, Belachew T, Bhonagiri V, Bihan M, Blaser MJ, Bloom T, Bonazzi V, Paul Brooks J, Buck GA, Buhay CJ, Busam DA, Campbell JL, Canon SR, Cantarel BL, Chain PSG, Chen I-MA, Chen L, Chhibba S, Chu K, Ciulla DM, Clemente JC, Clifton SW, Conlan S, Crabtree J, Cutting MA, Davidovics NJ, Davis CC, DeSantis TZ, Deal C, Delehaunty KD, Dewhirst FE, Deych E, Ding Y, Dooling DJ, Dugan SP, Michael Dunne W, Scott Durkin A, Edgar RC, Erlich RL, Farmer CN, Farrell RM, Faust K, Feldgarden M, Felix VM, Fisher S, Fodor AA, Forney LJ, Foster L, Di Francesco V, Friedman J, Friedrich DC, Fronick CC, Fulton LL, Gao H, Garcia N, Giannoukos G, Giblin C, Giovanni MY, Goldberg JM, Goll J, Gonzalez A, Griggs A, Gujja S, Kinder Haake S, Haas BJ, Hamilton HA, Harris EL, Hepburn TA, Herter B, Hoffmann DE, Holder ME, Howarth C, Huang KH, Huse SM, Izard J, Jansson JK, Jiang H, Jordan C, Joshi V, Katancik JA, Keitel WA, Kelley ST, Kells C, King NB, Knights D, Kong HH, Koren O, Koren S, Kota KC, Kovar CL, Kyrpides NC, La Rosa PS, Lee SL, Lemon KP, Lennon N, Lewis CM, Lewis L, Ley RE, Li K, Liolios K, Liu B, Liu Y, Lo C-C, Lozupone CA, Dwayne Lunsford R, Madden T, Mahurkar AA, Mannon PJ, Mardis ER, Markowitz VM, Mavromatis K, McCorrison JM, McDonald D, McEwen J, McGuire AL, McInnes P, Mehta T, Mihindukulasuriya KA, Miller JR, Minx PJ, Newsham I, Nusbaum C, O’Laughlin M, Orvis J, Pagani I, Palaniappan K, Patel SM, Pearson M, Peterson J, Podar M, Pohl C, Pollard KS, Pop M, Priest ME, Proctor LM, Qin X, Raes J, Ravel J, Reid JG, Rho M, Rhodes R, Riehle KP, Rivera MC, Rodriguez-Mueller B, Rogers Y-H, Ross MC, Russ C, Sanka RK, Sankar P, Fah Sathirapongsasuti J, Schloss JA, Schloss PD, Schmidt TM, Scholz M, Schriml L, Schubert AM, Segata N, Segre JA, Shannon WD, Sharp RR, Sharpton TJ, Shenoy N, Sheth NU, Simone GA, Singh I, Smillie CS, Sobel JD, Sommer DD, Spicer P, Sutton GG, Sykes SM, Tabbaa DG, Thiagarajan M, Tomlinson CM, Torralba M, Treangen TJ, Truty RM, Vishnivetskaya TA, Walker J, Wang L, Wang Z, Ward DV, Warren W, Watson MA, Wellington C, Wetterstrand KA, White JR, Wilczek-Boney K, Wu Y, Wylie KM, Wylie T, Yandava C, Ye L, Ye Y, Yooseph S, Youmans BP, Zhang L, Zhou Y, Zhu Y, Zoloth L, Zucker JD, Birren BW, Gibbs RA, Highlander SK, Methé BA, Nelson KE, Petrosino JF, Weinstock GM, Wilson RK, White O. 2012. Structure, function and diversity of the healthy human microbiome. Nature 486:207–214. DOI: 10.1038/nature11234.

[40] American Gut. Available at http://humanfoodproject.com/americangut/ (accessed May 29, 2019).

