## Supplemental data for "A robust metatranscriptomic technology for population-scale studies of diet, gut microbiome, and human health"

Table S1: Percent of sequencing reads assigned to rRNA for stool samples processed through Viomega without the custom rRNA depletion method. Average percent rRNA = 95.7 +/- 1.8%.

| **Sample** | **Human Stool RNA pool** | **# of stool RNA samples in pool** | **rRNA, %** |
|---|---|---|---|
| 1 | A | 37 | 97.26 |
| 2 | | | 97.51 |
| 3 | | | 97.31 |
| 4 | B | 42 | 95.3 |
| 5 | | | 94.29 |
| 6 | | | 94.4 |
| 7 | C | 38 | 93.22 |
| 8 | | | 92.76 |
| 9 | | | 93.41 |
| 10 | D | 12 | 98.09 |
| 11 | | | 97.85 |
| 12 | | | 97.89 |
| 13 | E | 12 | 96.97 |
| 14 | | | 96.3 |
| 15 | | | 96.69 |
| 16 | F | 12 | 94.71 |
| 17 | | | 94.26 |
| 18 | | | 93.97 |

Table S2: All strains identified in 10,000 human stool samples

| | Taxonomy ID | Strain name | SuperKingdom | Prevalence in 10,000 samples, % |
|---|---|---|---|---|
| 1 | 742768 | Eggerthella lenta 1_1_60AFAA | Bacteria | 97.08 |
| 2 | 411469 | [Eubacterium] hallii DSM 3353 | Bacteria | 93.93 |
| 3 | 546273 | Veillonella dispar ATCC 17748 | Bacteria | 92.34 |
| 4 | 445972 | Anaerotruncus colihominis DSM 17241 | Bacteria | 91.44 |
| 5 | 1650661.1 | Clostridium phoceensis strain GD3 | Bacteria | 90.83 |
| 6 | 411459 | Blautia obeum ATCC 29174 | Bacteria | 89.67 |
| 7 | 39485.2 | [Eubacterium] eligens strain 2789STDY5834875 | Bacteria | 88.93 |
| 8 | 748224 | Faecalibacterium cf. prausnitzii KLE1255 | Bacteria | 88.31 |
| 9 | 411483 | Faecalibacterium prausnitzii A2-165 | Bacteria | 86.05 |
| 10 | 585394 | Roseburia hominis A2-183 | Bacteria | 83.35 |
| 11 | 445970 | Alistipes putredinis DSM 17216 | Bacteria | 79.92 |
| 12 | 1408428 | Bilophila wadsworthia ATCC 49260 | Bacteria | 79.64 |
| 13 | 1737424.1 | Blautia massiliensis sp. GD8 | Bacteria | 78.91 |
| 14 | 709991 | Odoribacter splanchnicus DSM 20712 | Bacteria | 78.15 |
| 15 | 411485 | Faecalibacterium prausnitzii M21/2 | Bacteria | 77.79 |
| 16 | 853.1 | Faecalibacterium prausnitzii strain 2789STDY5834970 | Bacteria | 76.83 |
| 17 | 1715004.1 | Clostridiales bacterium KLE1615 | Bacteria | 76.44 |
| 18 | 717959 | Alistipes shahii WAL 8301 | Bacteria | 76.40 |
| 19 | 1519439.1 | Oscillibacter sp. ER4 | Bacteria | 73.98 |
| 20 | 39491.1 | [Eubacterium] rectale strain T1-815 | Bacteria | 73.62 |
| 21 | 39488.1 | [Eubacterium] hallii strain 2789STDY5834835 | Bacteria | 73.28 |
| 22 | 411477 | Parabacteroides merdae ATCC 43184 | Bacteria | 73.10 |
| 23 | 411471 | Subdoligranulum variabile DSM 15176 | Bacteria | 72.31 |
| 24 | 39490.1 | Eubacterium ramulus strain 2789STDY5608891 | Bacteria | 71.84 |
| 25 | 1384484 | Adlercreutzia equolifaciens DSM 19450 | Bacteria | 70.35 |
| 26 | 999413 | [Clostridium] innocuum 2959 | Bacteria | 70.25 |
| 27 | 428125 | [Clostridium] leptum DSM 753 | Bacteria | 70.16 |
| 28 | 657308 | Gordonibacter pamelaeae 7-10-1-b | Bacteria | 64.96 |
| 29 | 1697794.1 | Clostridia bacterium UC5.1-1D1 | Bacteria | 63.20 |
| 30 | 1310949 | Acinetobacter baumannii 24975_5 | Bacteria | 62.98 |
| 31 | 1118061.1 | Alistipes obesi | Bacteria | 62.42 |
| 32 | 1871035.1 | Ruminococcus sp. Marseille-P3213 sp. Marseille-P3213 | Bacteria | 61.52 |
| 33 | 214856.1 | Alistipes finegoldii strain 2789STDY5608890 | Bacteria | 61.03 |
| 34 | 853.3 | Faecalibacterium prausnitzii strain 2789STDY5608869 | Bacteria | 59.87 |
| 35 | 1033732 | Alistipes senegalensis JC50 | Bacteria | 59.30 |
| 36 | 1561.2 | Clostridium baratii strain 2789STDY5834907 | Bacteria | 57.98 |
| 37 | 1203611 | Alistipes onderdonkii WAL 8169 = DSM 19147 | Bacteria | 57.58 |
| 38 | 1121130 | Butyricimonas virosa DSM 23226 | Bacteria | 57.46 |
| 39 | 679935 | Alistipes finegoldii DSM 17242 | Bacteria | 56.98 |
| 40 | 166486.1 | Roseburia intestinalis strain 2789STDY5834960 | Bacteria | 56.66 |
| 41 | 411460 | Ruminococcus torques ATCC 27756 | Bacteria | 55.74 |

| | | | | |
|---|---|---|---|---|
| 42 | 1917876.1 | Blautia sp. Marseille-P3087 sp. Marseille-P3087 | Bacteria | 54.52 |
| 43 | 1673721.1 | Intestinimonas massiliensis sp. GD2 | Bacteria | 54.41 |
| 44 | 552398.1 | Ruminococcaceae bacterium D16 | Bacteria | 53.64 |
| 45 | 1841867.1 | Phocea massiliensis strain Marseille-P2769 | Bacteria | 53.62 |
| 46 | 999420 | Parabacteroides merdae CL03T12C32 | Bacteria | 53.57 |
| 47 | 360807.3 | Roseburia inulinivorans strain 2789STDY5608887 | Bacteria | 53.37 |
| 48 | 742725 | Alistipes indistinctus YIT 12060 | Bacteria | 52.74 |
| 49 | 1232459.1 | Oscillospiraceae bacterium VE202-24 | Bacteria | 52.59 |
| 50 | 1160721.1 | Ruminococcus bicirculans | Bacteria | 52.44 |
| 51 | 411903 | Collinsella aerofaciens ATCC 25986 | Bacteria | 52.05 |
| 52 | 88431.4 | Dorea longicatena strain 2789STDY5834914 | Bacteria | 52.01 |
| 53 | 742726 | Barnesiella intestinihominis YIT 11860 | Bacteria | 50.96 |
| 54 | 46506.1 | Bacteroides stercoris strain CL09T03C01 | Bacteria | 50.29 |
| 55 | 469610.1 | Burkholderiales bacterium 1_1_47 | Bacteria | 50.25 |
| 56 | 301302.1 | Roseburia faecis | Bacteria | 50.11 |
| 57 | 1121098 | Bacteroides massiliensis B84634 = Timone 84634 = DSM 17679 = JCM 13223 | Bacteria | 49.60 |
| 58 | 1339345 | Parabacteroides distasonis str. 3999B T(B) 6 | Bacteria | 49.53 |
| 59 | 428126 | [Clostridium] spiroforme DSM 1552 | Bacteria | 49.02 |
| 60 | 1235786 | Bacteroides vulgatus dnLKV7 | Bacteria | 48.68 |
| 61 | 411463 | Eubacterium ventriosum ATCC 27560 | Bacteria | 47.90 |
| 62 | 762966 | Parasutterella excrementihominis YIT 11859 | Bacteria | 47.89 |
| 63 | 1095771.1 | Ruminococcus sp. JC304 | Bacteria | 47.14 |
| 64 | 1235787 | Bacteroides uniformis dnLKV2 | Bacteria | 46.81 |
| 65 | 536231 | Roseburia intestinalis L1-82 | Bacteria | 46.72 |
| 66 | 665949.1 | Tannerella sp. 6_1_58FAA_CT1 | Bacteria | 45.53 |
| 67 | 301302.2 | Roseburia faecis strain 2789STDY5608863 | Bacteria | 45.20 |
| 68 | 411470 | Ruminococcus gnavus ATCC 29149 | Bacteria | 44.60 |
| 69 | 411461 | Dorea formicigenerans ATCC 27755 | Bacteria | 43.73 |
| 70 | 853.2 | Faecalibacterium prausnitzii strain 2789STDY5834930 | Bacteria | 43.69 |
| 71 | 997877 | Bacteroides dorei CL03T12C01 | Bacteria | 43.66 |
| 72 | 665956.1 | Subdoligranulum sp. 4_3_54A2FAA | Bacteria | 43.50 |
| 73 | 1211813 | Alistipes ihumii AP11 | Bacteria | 43.27 |
| 74 | 1703332.1 | Lachnospiraceae bacterium TF01-11 | Bacteria | 43.12 |
| 75 | 39492.1 | [Eubacterium] siraeum strain 2789STDY5834928 | Bacteria | 42.96 |
| 76 | 908612.1 | Alistipes sp. HGB5 | Bacteria | 42.46 |
| 77 | 1504823.1 | bacterium LF-3 | Bacteria | 42.43 |
| 78 | 411467 | Pseudoflavonifractor capillosus ATCC 29799 | Bacteria | 41.18 |
| 79 | 1073351 | Bacteroides stercoris CC31F | Bacteria | 40.91 |
| 80 | 214856.2 | Alistipes finegoldii strain 2789STDY5834947 | Bacteria | 40.89 |
| 81 | 1150298.2 | Fusicatenibacter saccharivorans strain 2789STDY5834885 | Bacteria | 40.87 |
| 82 | 1150298.1 | Fusicatenibacter saccharivorans strain 2789STDY5608849 | Bacteria | 40.83 |
| 83 | 450746.1 | Coprobacillus sp. 8_1_38FAA | Bacteria | 40.46 |
| 84 | 471875 | Ruminococcus lactaris ATCC 29176 | Bacteria | 40.21 |
| 85 | 39485.1 | [Eubacterium] eligens strain 2789STDY5834878 | Bacteria | 39.69 |

| | | | | |
|---|---|---|---|---|
| 86 | 658087.1 | Lachnospiraceae bacterium 7_1_58FAA | Bacteria | 39.14 |
| 87 | 39491.2 | [Eubacterium] rectale strain 2789STDY5834968 | Bacteria | 39.06 |
| 88 | 483215 | Bacteroides finegoldii DSM 17565 | Bacteria | 38.99 |
| 89 | 1297617.1 | Intestinimonas butyriciproducens strain AF211 | Bacteria | 38.89 |
| 90 | 545696 | Holdemania filiformis DSM 12042 | Bacteria | 38.37 |
| 91 | 1720200.1 | Anaerotruncus rubiinfantis sp. MT15 | Bacteria | 38.13 |
| 92 | 562.665 | Escherichia coli isolate 15 | Bacteria | 37.84 |
| 93 | 411486.1 | Clostridium sp. M62/1 | Bacteria | 37.66 |
| 94 | 1499682.1 | Alistipes sp. AL-1 | Bacteria | 37.07 |
| 95 | 585543.1 | Bacteroides sp. D20 | Bacteria | 36.99 |
| 96 | 820.3 | Bacteroides uniformis strain 2789STDY5834847 | Bacteria | 36.89 |
| 97 | 997891 | Bacteroides vulgatus CL09T03C04 | Bacteria | 36.83 |
| 98 | 1232439.1 | Clostridiales bacterium VE202-03 | Bacteria | 36.69 |
| 99 | 820.5 | Bacteroides uniformis strain 2789STDY5608791 | Bacteria | 36.47 |
| 100 | 1121115 | Blautia wexlerae DSM 19850 | Bacteria | 36.26 |
| 101 | 1150298.3 | Fusicatenibacter saccharivorans strain 2789STDY5834923 | Bacteria | 36.13 |
| 102 | 1720194.1 | Clostridium sp. AT4 sp. AT5 | Bacteria | 35.76 |
| 103 | 47678.2 | Bacteroides caccae strain 2789STDY5834880 | Bacteria | 35.52 |
| 104 | 820.9 | Bacteroides uniformis strain KLE1607 | Bacteria | 35.44 |
| 105 | 515619 | [Eubacterium rectale] ATCC 33656 | Bacteria | 34.82 |
| 106 | 435590 | Bacteroides vulgatus ATCC 8482 | Bacteria | 34.72 |
| 107 | 418240.2 | Blautia wexlerae strain 2789STDY5834911 | Bacteria | 34.25 |
| 108 | 1750560.1 | Parabacteroides sp. SN4 strain SN4, sp. SB4 | Bacteria | 34.16 |
| 109 | 1739298.1 | Bacteroides sp. HMSC067B03 | Bacteria | 33.55 |
| 110 | 1232453.1 | Clostridiales bacterium VE202-21 | Bacteria | 33.12 |
| 111 | 428128 | [Eubacterium] siraeum DSM 15702 | Bacteria | 33.09 |
| 112 | 1232438.1 | Clostridiales bacterium VE202-01 | Bacteria | 33.00 |
| 113 | 649724.1 | Clostridium sp. ATCC BAA-442 | Bacteria | 32.75 |
| 114 | 762984 | Bacteroides clarus YIT 12056 | Bacteria | 32.70 |
| 115 | 329854.2 | Bacteroides intestinalis strain KLE1704 | Bacteria | 32.24 |
| 116 | 411901 | Bacteroides caccae ATCC 43185 | Bacteria | 32.09 |
| 117 | 820.1 | Bacteroides uniformis | Bacteria | 31.96 |
| 118 | 411462 | Dorea longicatena DSM 13814 | Bacteria | 31.84 |
| 119 | 1697793.1 | Clostridia bacterium UC5.1-1E11 | Bacteria | 31.65 |
| 120 | 742722.1 | Collinsella sp. 4_8_47FAA | Bacteria | 31.64 |
| 121 | 821.2 | Bacteroides vulgatus strain 2789STDY5834842 | Bacteria | 31.63 |
| 122 | 40520.1 | Blautia obeum strain 2789STDY5834921 | Bacteria | 31.62 |
| 123 | 1432052.6 | Eisenbergiella tayi strain NML150140-1 | Bacteria | 31.17 |
| 124 | 537012 | Bacteroides cellulosilyticus DSM 14838 | Bacteria | 30.84 |
| 125 | 562983 | Gemella sanguinis M325 | Bacteria | 30.79 |
| 126 | 1280698 | Dorea longicatena AGR2136 | Bacteria | 30.28 |
| 127 | 457412.1 | Ruminococcus sp. 5_1_39BFAA | Bacteria | 29.93 |
| 128 | 762968 | Paraprevotella clara YIT 11840 | Bacteria | 29.81 |
| 129 | 39488.2 | [Eubacterium] hallii strain 2789STDY5834966 | Bacteria | 29.13 |
| 130 | 470146 | Coprococcus comes ATCC 27758 | Bacteria | 28.79 |
| 131 | 702450 | Turicibacter sanguinis PC909 | Bacteria | 28.77 |

| | | | | |
|---|---|---|---|---|
| 132 | 1871020.1 | Clostridium sp. Marseille-P3244 sp. Marseille-P3244 | Bacteria | 28.52 |
| 133 | 515620 | [Eubacterium] eligens ATCC 27750 | Bacteria | 28.51 |
| 134 | 1871018.1 | Angelakisella massiliensis strain Marseille-P3217 | Bacteria | 28.49 |
| 135 | 1852384.1 | Ruminococcaceae bacterium Marseille-P2963 | Bacteria | 28.31 |
| 136 | 742821 | Sutterella wadsworthensis 3_1_45B | Bacteria | 27.91 |
| 137 | 40520.5 | Blautia obeum strain 2789STDY5834957 | Bacteria | 27.87 |
| 138 | 762982 | Paraprevotella xylaniphila YIT 11841 | Bacteria | 27.66 |
| 139 | 1776382.1 | Neglecta timonensis strain SN17 | Bacteria | 27.37 |
| 140 | 622312 | Roseburia inulinivorans DSM 16841 | Bacteria | 27.10 |
| 141 | 997873 | Bacteroides caccae CL03T12C61 | Bacteria | 27.05 |
| 142 | 411473 | Ruminococcus callidus ATCC 27760 | Bacteria | 27.03 |
| 143 | 33039.3 | [Ruminococcus] torques strain 2789STDY5608867 | Bacteria | 26.72 |
| 144 | 1211417.1 | uncultured phage crAssphage | Viruses | 26.64 |
| 145 | 360807.2 | Roseburia inulinivorans strain 2789STDY5608835 | Bacteria | 26.64 |
| 146 | 1310661 | Acinetobacter baumannii 855125 | Bacteria | 26.25 |
| 147 | 1073376 | Ruminococcus lactaris CC59_002D | Bacteria | 26.25 |
| 148 | 483217 | Bacteroides dorei DSM 17855 | Bacteria | 26.24 |
| 149 | 33043.4 | Coprococcus eutactus strain 2789STDY5608829 | Bacteria | 26.20 |
| 150 | 1870991.1 | Massilioclostridium coli strain Marseille-P2976 | Bacteria | 26.16 |
| 151 | 33035.1 | Blautia producta strain ER3 | Bacteria | 26.09 |
| 152 | 1776384.1 | Emergencia timonensis strain SN18 | Bacteria | 26.09 |
| 153 | 1203465 | Bacteroides timonensis AP1 | Bacteria | 25.91 |
| 154 | 445971 | Anaerofustis stercorihominis DSM 17244 | Bacteria | 25.70 |
| 155 | 1352.143 | Enterococcus faecium isolate Hp_74-d6 | Bacteria | 25.50 |
| 156 | 537006 | Parabacteroides johnsonii DSM 18315 | Bacteria | 25.49 |
| 157 | 457389.1 | Bacteroides sp. 3_1_13 | Bacteria | 25.47 |
| 158 | 1816676.1 | Alistipes sp. Marseille-P2431 sp. Marseille-P2431 | Bacteria | 25.09 |
| 159 | 1163670.1 | Bacteroides sp. 14(A) | Bacteria | 25.06 |
| 160 | 39491.3 | [Eubacterium] rectale strain 2789STDY5608860 | Bacteria | 25.02 |
| 161 | 1627893.1 | Ruminococcaceae bacterium cv2 | Bacteria | 24.98 |
| 162 | 470145 | Bacteroides coprocola DSM 17136 | Bacteria | 24.89 |
| 163 | 469592.1 | Bacteroides sp. 3_1_19 | Bacteria | 24.80 |
| 164 | 821.3 | Bacteroides vulgatus strain 2789STDY5834944 | Bacteria | 24.79 |
| 165 | 1408437 | Butyricicoccus desmolans ATCC 43058 | Bacteria | 24.66 |
| 166 | 100886.1 | Catenibacterium mitsuokai strain 2789STDY5608825 | Bacteria | 24.64 |
| 167 | 360807.1 | Roseburia inulinivorans | Bacteria | 24.53 |
| 168 | 537011 | Prevotella copri DSM 18205 | Bacteria | 24.34 |
| 169 | 658662.1 | Parabacteroides sp. D26 | Bacteria | 24.33 |
| 170 | 665953 | Bacteroides eggerthii 1_2_48FAA | Bacteria | 24.29 |
| 171 | 999421 | Parabacteroides merdae CL09T00C40 | Bacteria | 24.19 |
| 172 | 997888 | Bacteroides finegoldii CL09T03C10 | Bacteria | 24.06 |
| 173 | 511680 | Butyrivibrio crossotus DSM 2876 | Bacteria | 23.93 |
| 174 | 563193.1 | Parabacteroides sp. D13 | Bacteria | 23.81 |
| 175 | 411468 | [Clostridium] scindens ATCC 35704 | Bacteria | 23.75 |
| 176 | 818.7 | Bacteroides thetaiotaomicron strain 14-106904-2 | Bacteria | 23.72 |

| | | | | |
|---|---|---|---|---|
| 177 | 649756.2 | Anaerostipes hadrus strain 2789STDY5608830 | Bacteria | 23.70 |
| 178 | 1852366.1 | Holdemania sp. Marseille-P2844 sp. Marseille-P2844 | Bacteria | 23.65 |
| 179 | 1697787.1 | Clostridia bacterium UC5.1-1D10 | Bacteria | 23.58 |
| 180 | 626939 | Phascolarctobacterium succinatutens YIT 12067 | Bacteria | 23.55 |
| 181 | 1352.158 | Enterococcus faecium isolate Hp_23-14 | Bacteria | 23.48 |
| 182 | 1121323 | [Clostridium] lactatifermentans DSM 14214 | Bacteria | 23.19 |
| 183 | 74426.1 | Collinsella aerofaciens strain 2789STDY5834902 | Bacteria | 23.12 |
| 184 | 329854.1 | Bacteroides intestinalis | Bacteria | 22.63 |
| 185 | 1122155 | Lactonifactor longoviformis DSM 17459 | Bacteria | 22.62 |
| 186 | 820.2 | Bacteroides uniformis strain 2789STDY5834942 | Bacteria | 22.50 |
| 187 | 471870 | Bacteroides intestinalis DSM 17393 | Bacteria | 22.48 |
| 188 | 1203554 | Sutterella wadsworthensis HGA0223 | Bacteria | 21.95 |
| 189 | 518637 | Holdemanella biformis DSM 3989 | Bacteria | 21.77 |
| 190 | 1339352 | Bacteroides vulgatus str. 3975 RP4 | Bacteria | 21.76 |
| 191 | 410072.2 | Coprococcus comes strain 2789STDY5834962 | Bacteria | 21.75 |
| 192 | 556260 | Bacteroides dorei 5_1_36/D4 | Bacteria | 21.34 |
| 193 | 556259.1 | Bacteroides sp. D2 | Bacteria | 21.34 |
| 194 | 40520.2 | Blautia obeum strain 2789STDY5834861 | Bacteria | 21.30 |
| 195 | 821.5 | Bacteroides vulgatus strain mpk | Bacteria | 21.03 |
| 196 | 936548.1 | Actinomyces sp. ICM47 | Bacteria | 20.98 |
| 197 | 820.6 | Bacteroides uniformis strain 2789STDY5834898 | Bacteria | 20.79 |
| 198 | 28052.2 | Lachnospira pectinoschiza strain 2789STDY5834836 | Bacteria | 20.70 |
| 199 | 1339335 | Bacteroides fragilis str. 3-F-2 #6 | Bacteria | 20.57 |
| 200 | 1226324.1 | Blautia sp. KLE 1732 | Bacteria | 20.41 |
| 201 | 566550 | Hungatella hathewayi DSM 13479 | Bacteria | 20.30 |
| 202 | 338188.1 | Bacteroides finegoldii strain 2789STDY5608840 | Bacteria | 20.26 |
| 203 | 457393.1 | Bacteroides sp. 4_1_36 | Bacteria | 20.16 |
| 204 | 592028 | Dialister invisus DSM 15470 | Bacteria | 20.07 |
| 205 | 40520.4 | Blautia obeum strain 2789STDY5608837 | Bacteria | 19.99 |
| 206 | 821.1 | Bacteroides vulgatus strain 2789STDY5834897 | Bacteria | 19.95 |
| 207 | 1034345 | Senegalimassilia anaerobia JC110 | Bacteria | 19.53 |
| 208 | 239935.2 | Akkermansia muciniphila strain YL44 | Bacteria | 19.20 |
| 209 | 823.1 | Parabacteroides distasonis strain 2789STDY5608872 | Bacteria | 19.04 |
| 210 | 457395.1 | Bacteroides sp. 9_1_42FAA | Bacteria | 19.00 |
| 211 | 28052.1 | Lachnospira pectinoschiza strain 2789STDY5834886 | Bacteria | 18.95 |
| 212 | 1211819 | Holdemania massiliensis AP2 | Bacteria | 18.88 |
| 213 | 1033731 | Alistipes timonensis JC136 | Bacteria | 18.86 |
| 214 | 411489.1 | Clostridium sp. L2-50 | Bacteria | 18.85 |
| 215 | 999419 | Parabacteroides johnsonii CL02T12C29 | Bacteria | 18.81 |
| 216 | 742823 | Sutterella wadsworthensis 2_1_59BFAA | Bacteria | 18.73 |
| 217 | 40520.3 | Blautia obeum strain 2789STDY5608838 | Bacteria | 18.70 |
| 218 | 33039.4 | [Ruminococcus] torques strain 2789STDY5834841 | Bacteria | 18.67 |
| 219 | 74426.3 | Collinsella aerofaciens strain 2789STDY5608823 | Bacteria | 18.63 |

| | | | | |
|---|---|---|---|---|
| 220 | 702443 | Bacteroides ovatus SD CMC 3f | Bacteria | 18.43 |
| 221 | 1550024.2 | Ruthenibacterium lactatiformans strain 585-1 | Bacteria | 18.36 |
| 222 | 1720300.1 | Ruminococcus sp. AT10 | Bacteria | 18.33 |
| 223 | 1121129 | Butyricimonas synergistica DSM 23225 | Bacteria | 18.19 |
| 224 | 457394.1 | Bacteroides sp. 4_3_47FAA | Bacteria | 18.14 |
| 225 | 823.4 | Parabacteroides distasonis strain 2789STDY5834901 | Bacteria | 17.92 |
| 226 | 693988.1 | Bilophila sp. 4_1_30 | Bacteria | 17.92 |
| 227 | 1280691 | Blautia wexlerae AGR2146 | Bacteria | 17.69 |
| 228 | 1226325.1 | Clostridium sp. KLE 1755 | Bacteria | 17.61 |
| 229 | 39491.4 | [Eubacterium] rectale strain 2789STDY5834884 | Bacteria | 17.59 |
| 230 | 1121096 | Bacteroides gallinarum DSM 18171 = JCM 13658 | Bacteria | 17.53 |
| 231 | 537013 | [Clostridium] methylpentosum DSM 5476 | Bacteria | 17.24 |
| 232 | 1574262.1 | Sutterella sp. KLE1602 | Bacteria | 17.22 |
| 233 | 820.7 | Bacteroides uniformis strain 2789STDY5608864 | Bacteria | 17.11 |
| 234 | 411490 | Anaerostipes caccae DSM 14662 | Bacteria | 17.09 |
| 235 | 1870993.1 | Tyzzerella sp. Marseille-P3062 sp. Marseille-P3062 | Bacteria | 16.92 |
| 236 | 483216 | Bacteroides eggerthii DSM 20697 | Bacteria | 16.85 |
| 237 | 1739319.1 | Bacteroides sp. HMSC068A09 | Bacteria | 16.67 |
| 238 | 563192 | Bilophila wadsworthia 3_1_6 | Bacteria | 16.53 |
| 239 | 1123075 | Ruminococcus gauvreauii DSM 19829 | Bacteria | 16.49 |
| 240 | 1078089.1 | Bacteroides sp. HPS0048 | Bacteria | 16.46 |
| 241 | 1280669.1 | Dorea sp. AGR2135 | Bacteria | 16.34 |
| 242 | 1203608 | Bacteroides nordii WAL 11050 = JCM 12987 | Bacteria | 16.27 |
| 243 | 371601.1 | Bacteroides xylanisolvens strain 2789STDY5608839 | Bacteria | 16.25 |
| 244 | 665954 | Bacteroides ovatus 3_8_47FAA | Bacteria | 15.92 |
| 245 | 33039.2 | [Ruminococcus] torques strain 2789STDY5608833 | Bacteria | 15.88 |
| 246 | 742765 | Dorea formicigenerans 4_6_53AFAA | Bacteria | 15.76 |
| 247 | 457390.1 | Bacteroides sp. 3_1_23 | Bacteria | 15.65 |
| 248 | 418240.1 | Blautia wexlerae strain 2789STDY5834863 | Bacteria | 15.65 |
| 249 | 47678.1 | Bacteroides caccae strain 2789STDY5834946 | Bacteria | 15.60 |
| 250 | 1432052.1 | Eisenbergiella tayi strain NML | Bacteria | 15.53 |
| 251 | 1232443.1 | Clostridiales bacterium VE202-13 | Bacteria | 15.53 |
| 252 | 1232457.1 | Clostridiales bacterium VE202-27 | Bacteria | 15.39 |
| 253 | 1653435.1 | Clostridium sp. BR31 | Bacteria | 15.29 |
| 254 | 818.2 | Bacteroides thetaiotaomicron strain 19_BTHE | Bacteria | 15.29 |
| 255 | 1256908 | Eubacterium ramulus ATCC 29099 | Bacteria | 15.25 |
| 256 | 823.3 | Parabacteroides distasonis strain 2789STDY5608822 | Bacteria | 15.17 |
| 257 | 1235785 | Bacteroides thetaiotaomicron dnLKV9 | Bacteria | 15.12 |
| 258 | 411474 | Coprococcus eutactus ATCC 27759 | Bacteria | 15.07 |
| 259 | 457391.1 | Bacteroides sp. 3_1_33FAA | Bacteria | 14.96 |
| 260 | 823.2 | Parabacteroides distasonis strain 2789STDY5834948 | Bacteria | 14.87 |
| 261 | 457415.1 | Synergistes sp. 3_1_syn1 | Bacteria | 14.80 |

| | | | | |
|---|---|---|---|---|
| 262 | 349741 | Akkermansia muciniphila ATCC BAA-835 | Bacteria | 14.79 |
| 263 | 1235788 | Bacteroides massiliensis dnLKV3 | Bacteria | 14.78 |
| 264 | 997892 | Bacteroides xylanisolvens CL03T12C04 | Bacteria | 14.76 |
| 265 | 818.5 | Bacteroides thetaiotaomicron strain 2789STDY5834945 | Bacteria | 14.69 |
| 266 | 1339314 | Bacteroides fragilis str. 3976T8 | Bacteria | 14.56 |
| 267 | 246787.1 | Bacteroides cellulosilyticus strain CL09T06C25 | Bacteria | 14.53 |
| 268 | 500632 | Tyzzerella nexilis DSM 1787 | Bacteria | 14.51 |
| 269 | 911128 | Methanobrevibacter smithii TS94C | Archaea | 14.47 |
| 270 | 658655.1 | Lachnospiraceae bacterium 1_4_56FAA | Bacteria | 14.41 |
| 271 | 818.1 | Bacteroides thetaiotaomicron strain 7330 | Bacteria | 14.38 |
| 272 | 469590.1 | Bacteroides sp. 2_2_4 | Bacteria | 14.28 |
| 273 | 484018 | Bacteroides plebeius DSM 17135 | Bacteria | 14.17 |
| 274 | 1077285 | Bacteroides faecis MAJ27 | Bacteria | 14.11 |
| 275 | 1650663.1 | Fournierella massiliensis strain AM2 | Bacteria | 14.03 |
| 276 | 649756.3 | Anaerostipes hadrus strain 2789STDY5834860 | Bacteria | 14.03 |
| 277 | 1297617.2 | Intestinimonas butyriciproducens strain 27-5-10 | Bacteria | 14.02 |
| 278 | 88431.3 | Dorea longicatena strain 2789STDY5608866 | Bacteria | 13.99 |
| 279 | 997884 | Bacteroides nordii CL02T12C05 | Bacteria | 13.95 |
| 280 | 910311.1 | Eggerthella sp. HGA1 | Bacteria | 13.94 |
| 281 | 483218 | [Bacteroides] pectinophilus ATCC 43243 | Bacteria | 13.93 |
| 282 | 1234889 | Leuconostoc mesenteroides subsp. cremoris TIFN8 | Bacteria | 13.93 |
| 283 | 1207542 | Bifidobacterium bifidum LMG 13195 | Bacteria | 13.92 |
| 284 | 997874 | Bacteroides cellulosilyticus CL02T12C19 | Bacteria | 13.65 |
| 285 | 476272 | Blautia hydrogenotrophica DSM 10507 | Bacteria | 13.34 |
| 286 | 818.1 | Bacteroides thetaiotaomicron | Bacteria | 13.27 |
| 287 | 1002367 | Prevotella stercorea DSM 18206 | Bacteria | 13.15 |
| 288 | 1841855.1 | Bacteroides sp. Marseille-P2653 sp. Marseille-P2653 | Bacteria | 12.94 |
| 289 | 1298596 | Ruminococcus faecis JCM 15917 | Bacteria | 12.86 |
| 290 | 1574263.1 | Candidatus Stoquefichus sp. KLE1796 | Bacteria | 12.66 |
| 291 | 1745713.1 | Bariatricus massiliensis strain AT12 | Bacteria | 12.58 |
| 292 | 1157708 | Variovorax paradoxus 110B | Bacteria | 12.53 |
| 293 | 1310866 | Acinetobacter baumannii 25307_2 | Bacteria | 12.52 |
| 294 | 999416 | Parabacteroides distasonis CL03T12C09 | Bacteria | 12.45 |
| 295 | 1349822 | Coprobacter fastidiosus NSB1 | Bacteria | 12.41 |
| 296 | 469589.1 | Bacteroides sp. 2_1_33B | Bacteria | 12.40 |
| 297 | 1121101 | Bacteroides salyersiae WAL 10018 = DSM 18765 = JCM 12988 | Bacteria | 12.14 |
| 298 | 435591 | Parabacteroides distasonis ATCC 8503 | Bacteria | 11.85 |
| 299 | 411475 | Flavonifractor plautii ATCC 29863 | Bacteria | 11.78 |
| 300 | 818.6 | Bacteroides thetaiotaomicron strain 2789STDY5834899 | Bacteria | 11.77 |
| 301 | 1907659.1 | Blautia sp. Marseille-P3201T strain Marseille-P3201 | Bacteria | 11.75 |
| 302 | 28116.7 | Bacteroides ovatus strain KLE1656 | Bacteria | 11.74 |
| 303 | 1550024.1 | Ruthenibacterium lactatiformans strain 668 | Bacteria | 11.72 |

| | | | | |
|---|---|---|---|---|
| 304 | 1852361.1 | Actinomyces sp. Marseille-P2825 sp. Marseille-P2825 | Bacteria | 11.69 |
| 305 | 1034346.1 | Dielma fastidiosa | Bacteria | 11.66 |
| 306 | 1805476.1 | Blautia sp. Marseille-P2398 | Bacteria | 11.64 |
| 307 | 821.4 | Bacteroides vulgatus strain NLAE-zl-G202 | Bacteria | 11.61 |
| 308 | 1796613.1 | Bacteroides caecimuris strain I48 | Bacteria | 11.61 |
| 309 | 371601.3 | Bacteroides xylanisolvens strain NLAE-zl-G339 | Bacteria | 11.60 |
| 310 | 169435.1 | Anaerotruncus colihominis strain 2789STDY5834939 | Bacteria | 11.44 |
| 311 | 1151410 | Clostridioides difficile P28 | Bacteria | 11.42 |
| 312 | 154046.2 | Hungatella hathewayi strain 2789STDY5834916 | Bacteria | 11.41 |
| 313 | 818.9 | Bacteroides thetaiotaomicron strain KPPR-3 | Bacteria | 11.36 |
| 314 | 818.4 | Bacteroides thetaiotaomicron strain 2789STDY5608873 | Bacteria | 11.27 |
| 315 | 518636 | [Clostridium asparagiforme] DSM 15981 | Bacteria | 11.22 |
| 316 | 999418 | Parabacteroides goldsteinii CL02T12C30 | Bacteria | 11.15 |
| 317 | 556269 | Oxalobacter formigenes OXCC13 | Bacteria | 11.15 |
| 318 | 702446 | Bacteroides vulgatus PC510 | Bacteria | 11.15 |
| 319 | 1352.147 | Enterococcus faecium isolate Hp_7-6 | Bacteria | 10.88 |
| 320 | 469586.1 | Bacteroides sp. 1_1_6 | Bacteria | 10.87 |
| 321 | 1227261 | Actinobaculum sp. oral taxon 183 str. F0552 | Bacteria | 10.87 |
| 322 | 28116.5 | Bacteroides ovatus strain NLAE-zl-C500 | Bacteria | 10.82 |
| 323 | 47770.1 | Lactobacillus crispatus strain C25 | Bacteria | 10.74 |
| 324 | 658659.1 | Erysipelotrichaceae bacterium 3_1_53 | Bacteria | 10.61 |
| 325 | 154046.1 | Hungatella hathewayi strain 2789STDY5608850 | Bacteria | 10.54 |
| 326 | 820.4 | Bacteroides uniformis strain 2789STDY5834844 | Bacteria | 10.53 |
| 327 | 1121333 | [Clostridium] saccharogumia DSM 17460 | Bacteria | 10.46 |
| 328 | 213810 | Ruminococcus champanellensis 18P13 = JCM 17042 | Bacteria | 10.40 |
| 329 | 33043.2 | Coprococcus eutactus strain 2789STDY5608888 | Bacteria | 10.38 |
| 330 | 521003 | Collinsella intestinalis DSM 13280 | Bacteria | 10.38 |
| 331 | 292800.2 | Flavonifractor plautii strain 2789STDY5834932 | Bacteria | 10.31 |
| 332 | 742738 | Clostridium orbiscindens 1_3_50AFAA | Bacteria | 10.24 |
| 333 | 469614.1 | Erysipelotrichaceae bacterium 6_1_45 | Bacteria | 10.20 |
| 334 | 626937.1 | Christensenella minuta strain DSM | Bacteria | 10.17 |
| 335 | 469593.1 | Bacteroides sp. 3_1_40A | Bacteria | 10.11 |
| 336 | 1076696 | Entamoeba nuttalli P19 | Eukaryota | 9.85 |
| 337 | 649742 | Actinomyces odontolyticus F0309 | Bacteria | 9.84 |
| 338 | 823.5 | Parabacteroides distasonis | Bacteria | 9.80 |
| 339 | 33043.3 | Coprococcus eutactus strain 2789STDY5608843 | Bacteria | 9.73 |
| 340 | 33038.1 | [Ruminococcus] gnavus | Bacteria | 9.69 |
| 341 | 457387.1 | Bacteroides sp. 1_1_30 | Bacteria | 9.63 |
| 342 | 562.2335 | Escherichia coli strain LS5218 | Bacteria | 9.53 |
| 343 | 469591.1 | Parabacteroides sp. 20_3 | Bacteria | 9.47 |
| 344 | 88431.1 | Dorea longicatena strain 2789STDY5834961 | Bacteria | 9.43 |
| 345 | 1297617.3 | Intestinimonas butyriciproducens strain ER1 | Bacteria | 9.30 |
| 346 | 291644.1 | Bacteroides salyersiae strain 2789STDY5608871 | Bacteria | 9.22 |
| 347 | 74426.2 | Collinsella aerofaciens strain 2789STDY5608842 | Bacteria | 9.19 |

| | | | | |
|---|---|---|---|---|
| 348 | 763034 | Bacteroides fluxus YIT 12057 | Bacteria | 9.15 |
| 349 | 88431.2 | Dorea longicatena strain 2789STDY5608851 | Bacteria | 9.03 |
| 350 | 1188792.1 | Phaseolus vulgaris endornavirus 1 | Viruses | 8.91 |
| 351 | 1852370.1 | Prevotellamassilia timonensis strain Marseille-P2831 | Bacteria | 8.82 |
| 352 | 33039.1 | [Ruminococcus] torques strain 2789STDY5834889 | Bacteria | 8.75 |
| 353 | 649756.6 | Anaerostipes hadrus strain 2789STDY5834959 | Bacteria | 8.74 |
| 354 | 1871006.1 | Bacteroides sp. Marseille-P3132 sp. Marseille-P3132 | Bacteria | 8.70 |
| 355 | 1776379.1 | Prevotella sp. KHD1 sp. KHD1 | Bacteria | 8.67 |
| 356 | 817.2 | Bacteroides fragilis strain 14-106904-1 | Bacteria | 8.57 |
| 357 | 818.11 | Bacteroides thetaiotaomicron strain KLE1254 | Bacteria | 8.53 |
| 358 | 553973 | [Clostridium] hylemonae DSM 15053 | Bacteria | 8.43 |
| 359 | 818.8 | Bacteroides thetaiotaomicron isolate 3731 | Bacteria | 8.36 |
| 360 | 1871021.1 | Lachnoclostridium phocaeense strain Marseille-P3177T sp. Marseille-P3177 | Bacteria | 8.22 |
| 361 | 658089.1 | Lachnospiraceae bacterium 5_1_63FAA | Bacteria | 8.21 |
| 362 | 410072.1 | Coprococcus comes strain 2789STDY5608832 | Bacteria | 8.15 |
| 363 | 649756.1 | Anaerostipes hadrus strain PEL | Bacteria | 7.98 |
| 364 | 1506471.1 | Sutterellaceae bacterium ND3 | Bacteria | 7.96 |
| 365 | 226186 | Bacteroides thetaiotaomicron VPI-5482 | Bacteria | 7.91 |
| 366 | 742727 | Bacteroides oleiciplenus YIT 12058 | Bacteria | 7.82 |
| 367 | 658086.1 | Lachnospiraceae bacterium 3_1_57FAA_CT1 | Bacteria | 7.80 |
| 368 | 888727 | Eubacterium sulci ATCC 35585 | Bacteria | 7.79 |
| 369 | 649756.7 | Anaerostipes hadrus strain BPB5 | Bacteria | 7.78 |
| 370 | 411464 | Desulfovibrio piger ATCC 29098 | Bacteria | 7.76 |
| 371 | 1392862 | Escherichia coli M17 | Bacteria | 7.73 |
| 372 | 1841856.1 | Bacteroides mediterraneensis strain Marseille-P2644 | Bacteria | 7.72 |
| 373 | 1232444.1 | Clostridiales bacterium VE202-15 | Bacteria | 7.62 |
| 374 | 876.1 | Desulfovibrio desulfuricans strain DSM | Bacteria | 7.57 |
| 375 | 292800.1 | Flavonifractor plautii strain 2789STDY5834892 | Bacteria | 7.54 |
| 376 | 428127 | [Eubacterium] dolichum DSM 3991 | Bacteria | 7.51 |
| 377 | 665950.1 | Lachnospiraceae bacterium 3_1_46FAA | Bacteria | 7.47 |
| 378 | 1852381.1 | Sutterellaceae bacterium Marseille-P2968 | Bacteria | 7.46 |
| 379 | 545697 | Clostridium celatum DSM 1785 | Bacteria | 7.43 |
| 380 | 457421.1 | Clostridiales bacterium 1_7_47FAA | Bacteria | 7.39 |
| 381 | 46503.1 | Parabacteroides merdae strain 2789STDY5834848 | Bacteria | 7.28 |
| 382 | 1339276 | Bacteroides fragilis str. DS-208 | Bacteria | 7.26 |
| 383 | 1903263.1 | Traorella massiliensis strain Marseille-P3110 | Bacteria | 7.21 |
| 384 | 85831.1 | Bacteroides acidifaciens | Bacteria | 7.15 |
| 385 | 12239.2 | Pepper mild mottle virus | Viruses | 7.14 |
| 386 | 997887 | Bacteroides salyersiae CL02T12C01 | Bacteria | 7.12 |
| 387 | 649756.4 | Anaerostipes hadrus strain 2789STDY5834908 | Bacteria | 7.12 |
| 388 | 1917883.1 | Bacteroides sp. Marseille-P3166 sp. Marseille-P3166 | Bacteria | 7.09 |

| | | | | |
|---|---|---|---|---|
| 389 | 410072.3 | Coprococcus comes strain 2789STDY5834866 | Bacteria | 7.05 |
| 390 | 999417 | Parabacteroides distasonis CL09T03C24 | Bacteria | 6.99 |
| 391 | 1121445 | Desulfovibrio desulfuricans subsp. desulfuricans DSM 642 | Bacteria | 6.92 |
| 392 | 1834205.1 | Burkholderiales bacterium YL45 | Bacteria | 6.80 |
| 393 | 742818 | Slackia piriformis YIT 12062 | Bacteria | 6.79 |
| 394 | 1339283 | Bacteroides fragilis str. 3996 N(B) 6 | Bacteria | 6.64 |
| 395 | 1547597.1 | Sanguibacteroides justesenii strain OUH | Bacteria | 6.58 |
| 396 | 938289.1 | Levyella massiliensis | Bacteria | 6.55 |
| 397 | 1871013.1 | Parabacteroides sp. Marseille-P3236 strain Marseille-P3236, sp. Marseille-P3136 | Bacteria | 6.54 |
| 398 | 457397.1 | Clostridium sp. 1_1_41A1FAA | Bacteria | 6.52 |
| 399 | 1339271 | Bacteroides fragilis str. J-143-4 | Bacteria | 6.47 |
| 400 | 457402.1 | Eubacterium sp. 3_1_31 | Bacteria | 6.46 |
| 401 | 1115692.1 | Cannabis cryptic virus isolate hemp09 | Viruses | 6.43 |
| 402 | 1339298 | Bacteroides fragilis str. 3725 D9(v) | Bacteria | 6.42 |
| 403 | 742817 | Odoribacter laneus YIT 12061 | Bacteria | 6.38 |
| 404 | 927665 | Parabacteroides goldsteinii DSM 19448 = WAL 12034 | Bacteria | 6.38 |
| 405 | 12968.1 | Blastocystis hominis isolate B | Eukaryota | 6.35 |
| 406 | 1676614.1 | Prevotella sp. 109 | Bacteria | 6.31 |
| 407 | 1291051 | [Clostridium] glycyrrhizinilyticum JCM 13369 | Bacteria | 6.23 |
| 408 | 585544.1 | Bacteroides sp. D22 | Bacteria | 6.21 |
| 409 | 1499681.1 | Collinsella sp. MS5 | Bacteria | 6.13 |
| 410 | 12172.2 | Shallot latent virus isolate MS/SW/Aus2 | Viruses | 6.12 |
| 411 | 1602172.1 | Prevotella sp. P5-125 | Bacteria | 6.11 |
| 412 | 1697795.1 | Clostridia bacterium UC5.1-2H11 | Bacteria | 6.11 |
| 413 | 817.3 | Bacteroides fragilis strain DCMOUH0085B | Bacteria | 6.09 |
| 414 | 410072.4 | Coprococcus comes strain 2789STDY5834913 | Bacteria | 6.08 |
| 415 | 649756.5 | Anaerostipes hadrus strain 2789STDY5608868 | Bacteria | 6.08 |
| 416 | 186772.1 | Saccharomyces 20S RNA narnavirus | Viruses | 6.07 |
| 417 | 28116.4 | Bacteroides ovatus strain NLAE-zl-C57 | Bacteria | 5.92 |
| 418 | 360107 | Campylobacter hominis ATCC BAA-381 | Bacteria | 5.90 |
| 419 | 1284708.1 | Tissierellia bacterium S7-1-4 | Bacteria | 5.87 |
| 420 | 328812.1 | Parabacteroides goldsteinii strain 910340 | Bacteria | 5.77 |
| 421 | 154288.3 | Turicibacter sanguinis strain 2789STDY5608865 | Bacteria | 5.70 |
| 422 | 1579343.1 | Streptococcus sp. 263_SSPC | Bacteria | 5.56 |
| 423 | 1078087.1 | Parabacteroides sp. HGS0025 | Bacteria | 5.56 |
| 424 | 362693.1 | Oryza sativa endornavirus | Viruses | 5.53 |
| 425 | 1816678.1 | Christensenella timonensis strain Marseille-P2437 | Bacteria | 5.40 |
| 426 | 997885 | Bacteroides ovatus CL02T12C04 | Bacteria | 5.38 |
| 427 | 817.11 | Bacteroides fragilis strain 20656-2-1 | Bacteria | 5.38 |
| 428 | 33043.1 | Coprococcus eutactus strain 2789STDY5834963 | Bacteria | 5.37 |
| 429 | 658085.1 | Lachnospiraceae bacterium 5_1_57FAA | Bacteria | 5.34 |
| 430 | 901.1 | Desulfovibrio piger isolate DESPIGER1 | Bacteria | 5.34 |
| 431 | 435830 | Actinomyces graevenitzii C83 | Bacteria | 5.24 |
| 432 | 665940.1 | Clostridium sp. 7_3_54FAA | Bacteria | 5.22 |
| 433 | 479437 | Eggerthella lenta DSM 2243 | Bacteria | 5.14 |

| | | | | |
|---|---|---|---|---|
| 434 | 1295009 | Candidatus Methanomassiliicoccus intestinalis Issoire-Mx1 | Archaea | 5.14 |
| 435 | 999408 | [Clostridium] clostridioforme 90A8 | Bacteria | 5.13 |
| 436 | 1227262 | Actinomyces johnsonii F0510 | Bacteria | 5.13 |
| 437 | 944170.2 | Blastocystis sp. subtype 4 strain WR1 | Eukaryota | 5.06 |
| 438 | 451640 | Catenibacterium mitsuokai DSM 15897 | Bacteria | 5.01 |
| 439 | 1459804 | Finegoldia magna ALB8 | Bacteria | 4.91 |
| 440 | 1121114 | Blautia producta ATCC 27340 = DSM 2950 | Bacteria | 4.83 |
| 441 | 1658112.1 | Eubacterium sp. SB2 | Bacteria | 4.79 |
| 442 | 1339348 | Bacteroides uniformis str. 3978 T3 i | Bacteria | 4.79 |
| 443 | 997886 | Bacteroides ovatus CL03T12C18 | Bacteria | 4.68 |
| 444 | 742743 | Dialister succinatiphilus YIT 11850 | Bacteria | 4.65 |
| 445 | 1123263 | Solobacterium moorei DSM 22971 | Bacteria | 4.61 |
| 446 | 28116.2 | Bacteroides ovatus strain 2789STDY5834943 | Bacteria | 4.55 |
| 447 | 1339344 | Parabacteroides distasonis str. 3999B T(B) 4 | Bacteria | 4.45 |
| 448 | 290052.1 | Acetivibrio ethanolgignens strain ACET-33324 | Bacteria | 4.44 |
| 449 | 1432052.7 | Eisenbergiella tayi strain DSM | Bacteria | 4.34 |
| 450 | 592026 | Catonella morbi ATCC 51271 | Bacteria | 4.26 |
| 451 | 1120941 | Actinomyces dentalis DSM 19115 | Bacteria | 4.24 |
| 452 | 1519438.1 | Eubacterium sp. ER2 | Bacteria | 4.21 |
| 453 | 327387.1 | Tropical soda apple mosaic virus isolate Okeechobee | Viruses | 4.17 |
| 454 | 445973 | Intestinibacter bartlettii DSM 16795 | Bacteria | 4.14 |
| 455 | 239935.1 | Akkermansia muciniphila isolate Urmite | Bacteria | 4.13 |
| 456 | 817.19 | Bacteroides fragilis strain DCMOUH0042B | Bacteria | 4.10 |
| 457 | 658656.1 | Lachnospiraceae bacterium 6_1_37FAA | Bacteria | 4.07 |
| 458 | 1841865.1 | Mediterranea massiliensis strain Marseille-P2645 | Bacteria | 4.05 |
| 459 | 292800.3 | Flavonifractor plautii strain YL31 | Bacteria | 4.04 |
| 460 | 1761477.1 | Tomato brown rugose fruit virus isolate Tom1-Jo | Viruses | 4.03 |
| 461 | 1871003.1 | Tidjanibacter massiliensis strain Marseille-P3084 | Bacteria | 4.02 |
| 462 | 1349763 | Curvibacter delicatus NBRC 14919 | Bacteria | 4.02 |
| 463 | 547042 | Bacteroides coprophilus DSM 18228 = JCM 13818 | Bacteria | 4.00 |
| 464 | 1671366.1 | Ruminococcus sp. DSM 100440 | Bacteria | 3.97 |
| 465 | 371601.2 | Bacteroides xylanisolvens strain NLAE-zl-C202 | Bacteria | 3.94 |
| 466 | 936595.1 | Lachnoanaerobaculum sp. OBRC5-5 | Bacteria | 3.91 |
| 467 | 1723384.1 | Fenollaria timonensis | Bacteria | 3.91 |
| 468 | 667015 | Bacteroides salanitronis DSM 18170 | Bacteria | 3.86 |
| 469 | 658657.1 | Erysipelotrichaceae bacterium 21_3 | Bacteria | 3.86 |
| 470 | 1531.1 | [Clostridium] clostridioforme strain 2789STDY5834865 | Bacteria | 3.85 |
| 471 | 457424 | Bacteroides fragilis 3_1_12 | Bacteria | 3.83 |
| 472 | 1871030.1 | Merdibacter massiliensis strain Marseille-P3254 | Bacteria | 3.80 |
| 473 | 742742 | Collinsella tanakaei YIT 12063 | Bacteria | 3.79 |
| 474 | 1423800 | Lactobacillus sanfranciscensis DSM 20451 | Bacteria | 3.75 |
| 475 | 1501392.1 | Coprobacter secundus strain 177 | Bacteria | 3.65 |
| 476 | 712411.1 | Olsenella sp. oral taxon 807 strain F0089 | Bacteria | 3.53 |
| 477 | 1133596 | Pediococcus pentosaceus IE-3 | Bacteria | 3.50 |

| | | | | |
|---|---|---|---|---|
| 478 | 1816694.1 | Clostridium sp. Marseille-P2538 sp. Marseille-P2538 | Bacteria | 3.48 |
| 479 | 261299.1 | Intestinibacter bartlettii strain 2789STDY5834879 | Bacteria | 3.48 |
| 480 | 880074 | Barnesiella viscericola DSM 18177 | Bacteria | 3.46 |
| 481 | 1197717.1 | Cloacibacillus porcorum strain CL-84 | Bacteria | 3.44 |
| 482 | 1602168.1 | Prevotella sp. P4-65 | Bacteria | 3.42 |
| 483 | 28116.1 | Bacteroides ovatus strain CL09T03C03 | Bacteria | 3.38 |
| 484 | 999407 | [Clostridium] clostridioforme 90A7 | Bacteria | 3.32 |
| 485 | 1261635.1 | Roseburia sp. 831b | Bacteria | 3.28 |
| 486 | 717960 | Faecalitalea cylindroides T2-87 | Bacteria | 3.25 |
| 487 | 1121094 | Bacteroides barnesiae DSM 18169 = JCM 13652 | Bacteria | 3.25 |
| 488 | 1121866 | Enterorhabdus mucosicola DSM 19490 | Bacteria | 3.23 |
| 489 | 368735.1 | Bell pepper mottle virus | Viruses | 3.23 |
| 490 | 1903262.1 | Bacteroides sp. Marseille-P3108 sp. Marseille-P3108 | Bacteria | 3.23 |
| 491 | 1686296.1 | Gabonia massiliensis strain GM3 | Bacteria | 3.22 |
| 492 | 742737 | Hungatella hathewayi WAL-18680 | Bacteria | 3.20 |
| 493 | 1261634.1 | Roseburia sp. 499 | Bacteria | 3.19 |
| 494 | 1609975.1 | Clostridium sp. FS41 | Bacteria | 3.19 |
| 495 | 44742.1 | Desulfovibrio fairfieldensis strain CCUG | Bacteria | 3.16 |
| 496 | 562.2194 | Escherichia coli strain UPEC_014 | Bacteria | 3.12 |
| 497 | 1339307 | Bacteroides fragilis str. 3719 A10 | Bacteria | 3.10 |
| 498 | 1121344 | [Clostridium] viride DSM 6836 | Bacteria | 3.08 |
| 499 | 157777.1 | Pea streak virus isolate VRS-541 | Viruses | 3.06 |
| 500 | 1522.3 | [Clostridium] innocuum strain NLAE-zl-C381 | Bacteria | 3.05 |
| 501 | 33038.2 | 2789STDY5608852 | Bacteria | 3.02 |
| 502 | 537007 | Blautia hansenii DSM 20583 | Bacteria | 3.00 |
| 503 | 665938.1 | Bacteroides sp. 2_1_56FAA | Bacteria | 2.97 |
| 504 | 480391.1 | Pediococcus argentinicus strain DSM | Bacteria | 2.96 |
| 505 | 1907658.1 | Bacteroides sp. Marseille-P3208T strain Marseille-P3208 | Bacteria | 2.93 |
| 506 | 888056 | Actinomyces sp. oral taxon 448 str. F0400 | Bacteria | 2.92 |
| 507 | 1736.1 | Eubacterium limosum strain 32_A2 | Bacteria | 2.91 |
| 508 | 53443.1 | Blautia hydrogenotrophica strain 2789STDY5608857 | Bacteria | 2.91 |
| 509 | 1232460.1 | Clostridiales bacterium VE202-28 | Bacteria | 2.91 |
| 510 | 999410 | [Clostridium] clostridioforme CM201 | Bacteria | 2.88 |
| 511 | 665937.1 | Anaerostipes sp. 3_2_56FAA | Bacteria | 2.86 |
| 512 | 39482.1 | [Eubacterium] contortum strain 2789STDY5834876 | Bacteria | 2.85 |
| 513 | 1496.4 | Clostridioides difficile isolate VL_0437 | Bacteria | 2.85 |
| 514 | 1384063 | Ruminococcus gnavus AGR2154 | Bacteria | 2.83 |
| 515 | 1232447.1 | Clostridiales bacterium VE202-09 | Bacteria | 2.82 |
| 516 | 339860 | Methanosphaera stadtmanae DSM 3091 | Archaea | 2.80 |
| 517 | 880526 | Rikenella microfusus DSM 15922 | Bacteria | 2.77 |
| 518 | 1496.59 | Clostridioides difficile strain CD105KSE11 | Bacteria | 2.77 |
| 519 | 866499 | Cloacibacillus evryensis DSM 19522 | Bacteria | 2.76 |
| 520 | 1232442.1 | Clostridiales bacterium VE202-06 | Bacteria | 2.74 |

| | | | | |
|---|---|---|---|---|
| 521 | 1870994.1 | Urmitella timonensis sp. Marseille-P2918 | Bacteria | 2.73 |
| 522 | 445975 | Collinsella stercoris DSM 13279 | Bacteria | 2.71 |
| 523 | 457422.1 | Erysipelotrichaceae bacterium 2_2_44A | Bacteria | 2.70 |
| 524 | 469597.1 | Coprobacillus sp. 8_2_54BFAA | Bacteria | 2.70 |
| 525 | 658083.1 | Lachnospiraceae bacterium 6_1_63FAA | Bacteria | 2.70 |
| 526 | 1321817 | Actinomyces graevenitzii F0530 | Bacteria | 2.59 |
| 527 | 1120979 | Alloscardovia omnicolens DSM 21503 | Bacteria | 2.56 |
| 528 | 1423794 | Lactobacillus pontis DSM 8475 | Bacteria | 2.55 |
| 529 | 1816677.1 | Butyricimonas sp. Marseille-P2440 sp. Marseille-P2440 | Bacteria | 2.54 |
| 530 | 658088.1 | Lachnospiraceae bacterium 9_1_43BFAA | Bacteria | 2.53 |
| 531 | 944168.1 | Blastocystis sp. subtype 3 | Eukaryota | 2.51 |
| 532 | 665941.1 | Coprobacillus sp. 3_3_56FAA | Bacteria | 2.50 |
| 533 | 1122217 | Megamonas rupellensis DSM 19944 | Bacteria | 2.48 |
| 534 | 84024.3 | Clostridium disporicum strain 2789STDY5834856 | Bacteria | 2.48 |
| 535 | 817.22 | Bacteroides fragilis strain DCMOUH0018B | Bacteria | 2.44 |
| 536 | 1118060 | Enorma massiliensis phI | Bacteria | 2.40 |
| 537 | 47900.2 | Garlic common latent virus isolate SW3.2 | Viruses | 2.39 |
| 538 | 187979.1 | Mitsuokella jalaludinii strain 2789STDY5608828 | Bacteria | 2.38 |
| 539 | 1574264.1 | Akkermansia sp. KLE1797 | Bacteria | 2.37 |
| 540 | 1602171.1 | Prevotella sp. P5-119 | Bacteria | 2.35 |
| 541 | 518634 | Bifidobacterium breve DSM 20213 = JCM 1192 | Bacteria | 2.35 |
| 542 | 59201.96 | Salmonella enterica subsp. enterica strain SE696A | Bacteria | 2.30 |
| 543 | 556268 | Oxalobacter formigenes HOxBLS | Bacteria | 2.29 |
| 544 | 1852383.1 | Ruminococcaceae bacterium Marseille-P2935 | Bacteria | 2.28 |
| 545 | 59201.11 | Salmonella enterica subsp. enterica strain ADRDL-LA-5-2013 | Bacteria | 2.28 |
| 546 | 649755 | Faecalitalea cylindroides ATCC 27803 | Bacteria | 2.26 |
| 547 | 478820 | Blastocystis sp. ATCC 50177/Nand II | Eukaryota | 2.26 |
| 548 | 1235797.1 | Oscillibacter sp. 1-3 | Bacteria | 2.26 |
| 549 | 908937 | Prevotella dentalis DSM 3688 | Bacteria | 2.25 |
| 550 | 1736.2 | Eubacterium limosum strain ATCC | Bacteria | 2.21 |
| 551 | 1122971 | Porphyromonas bennonis DSM 23058 = JCM 16335 | Bacteria | 2.21 |
| 552 | 742734 | [Clostridium] citroniae WAL-19142 | Bacteria | 2.21 |
| 553 | 649757 | Anaerostipes hadrus DSM 3319 | Bacteria | 2.18 |
| 554 | 39778.1 | Veillonella dispar strain DNF00926 | Bacteria | 2.17 |
| 555 | 1496.33 | Clostridioides difficile strain 7.10492 | Bacteria | 2.16 |
| 556 | 693979 | Bacteroides helcogenes P 36-108 | Bacteria | 2.15 |
| 557 | 1739394.1 | Porphyromonas sp. HMSC065F10 | Bacteria | 2.12 |
| 558 | 1796616.1 | Blautia sp. YL58 sp. YL58 | Bacteria | 2.09 |
| 559 | 999403 | [Clostridium] clostridioforme 90A1 | Bacteria | 2.07 |
| 560 | 742733 | [Clostridium] citroniae WAL-17108 | Bacteria | 2.04 |
| 561 | 28125.4 | Prevotella bivia strain GED7880 | Bacteria | 2.03 |
| 562 | 469587.1 | Bacteroides sp. 2_1_16 | Bacteria | 2.03 |
| 563 | 1230734.1 | Clostridiales bacterium S5-A14a | Bacteria | 2.03 |
| 564 | 412133 | Trichomonas vaginalis G3 | Eukaryota | 2.02 |
| 565 | 502558.1 | Eggerthella sp. YY7918 | Bacteria | 2.02 |

| | | | | |
|---|---|---|---|---|
| 566 | 1658109.1 | Candidatus Stoquefichus sp. SB1 | Bacteria | 2.01 |
| 567 | 1235794 | Enterorhabdus caecimuris B7 | Bacteria | 1.98 |
| 568 | 817.5 | Bacteroides fragilis | Bacteria | 1.98 |
| 569 | 1491.27 | Clostridium botulinum strain SU0801 | Bacteria | 1.98 |
| 570 | 997897 | [Clostridium] bolteae 90B8 | Bacteria | 1.97 |
| 571 | 665942.1 | Desulfovibrio sp. 6_1_46AFAA | Bacteria | 1.95 |
| 572 | 1585974.1 | Beduini massiliensis strain GM1 | Bacteria | 1.95 |
| 573 | 442302.1 | Porcine picobirnavirus strain 221/04-16/ITA/2004 | Viruses | 1.95 |
| 574 | 817.2 | Bacteroides fragilis strain DCMOUH0067B | Bacteria | 1.92 |
| 575 | 883077 | Actinomyces turicensis ACS-279-V-Col4 | Bacteria | 1.91 |
| 576 | 47900.1 | Garlic common latent virus | Viruses | 1.90 |
| 577 | 747056.1 | Blueberry shock virus isolate Berkely | Viruses | 1.89 |
| 578 | 706434 | Megasphaera micronuciformis F0359 | Bacteria | 1.89 |
| 579 | 665943.1 | Eggerthella sp. 1_3_56FAA | Bacteria | 1.87 |
| 580 | 817.8 | Bacteroides fragilis strain JIM10 | Bacteria | 1.87 |
| 581 | 1235789 | Parabacteroides goldsteinii dnLKV18 | Bacteria | 1.85 |
| 582 | 1395125 | Prevotella salivae F0493 | Bacteria | 1.84 |
| 583 | 1564113.1 | Sphingomonas sp. Ant H11 | Bacteria | 1.83 |
| 584 | 1123313 | Faecalicoccus pleomorphus DSM 20574 | Bacteria | 1.82 |
| 585 | 252598.1 | Saccharomyces sp. 'boulardii' strain unique28 | Eukaryota | 1.82 |
| 586 | 509923.1 | Beet cryptic virus 1 | Viruses | 1.80 |
| 587 | 1603888.1 | Megasphaera sp. MJR8396C | Bacteria | 1.80 |
| 588 | 154288.4 | Turicibacter sanguinis strain 2789STDY5834949 | Bacteria | 1.78 |
| 589 | 1632013.1 | Drancourtella massiliensis strain GD1 | Bacteria | 1.78 |
| 590 | 1611875.1 | RNA | Viruses | 1.78 |
| 591 | 1903261.1 | Desulfovibrio sp. Marseille-P3199 sp. Marseille-P3199 | Bacteria | 1.77 |
| 592 | 1602169.1 | Prevotella sp. P4-76 | Bacteria | 1.75 |
| 593 | 1871016.1 | Collinsella sp. Marseille-P3245 sp. Marseille-P3245 | Bacteria | 1.73 |
| 594 | 1078090.1 | Coprococcus sp. HPP0074 | Bacteria | 1.72 |
| 595 | 706436 | Capnocytophaga sp. oral taxon 329 str. F0087 | Bacteria | 1.72 |
| 596 | 1073388 | Bacteroides fragilis HMW 616 | Bacteria | 1.71 |
| 597 | 1030842 | Ruminococcus flavefaciens 17 | Bacteria | 1.69 |
| 598 | 470.218 | Acinetobacter baumannii strain ABBL027 | Bacteria | 1.69 |
| 599 | 999412 | Hungatella hathewayi 12489931 | Bacteria | 1.67 |
| 600 | 1720313.1 | Bittarella massiliensis strain GD6 | Bacteria | 1.67 |
| 601 | 1871022.1 | Libanicoccus massiliensis strain Marseille-P3237T sp. Marseille-P3237 | Bacteria | 1.65 |
| 602 | 747056.2 | Blueberry shock virus | Viruses | 1.65 |
| 603 | 1339275 | Bacteroides fragilis str. Ds-233 | Bacteria | 1.64 |
| 604 | 742735 | [Clostridium] clostridioforme 2_1_49FAA | Bacteria | 1.62 |
| 605 | 857135 | Streptococcus mutans U138 | Bacteria | 1.62 |
| 606 | 39950.1 | Dialister pneumosintes strain F0677 | Bacteria | 1.62 |
| 607 | 1280695 | [Clostridium] clostridioforme AGR2157 | Bacteria | 1.57 |
| 608 | 1437603 | Bifidobacterium mongoliense DSM 21395 | Bacteria | 1.56 |
| 609 | 1069534 | Lactobacillus ruminis ATCC 27782 | Bacteria | 1.56 |
| 610 | 742723.1 | Lachnospiraceae bacterium 2_1_46FAA | Bacteria | 1.56 |

| | | | | |
|---|---|---|---|---|
| 611 | 1352.159 | Enterococcus faecium isolate Hp_6-9 | Bacteria | 1.55 |
| 612 | 1232452.1 | Clostridiales bacterium VE202-14 | Bacteria | 1.54 |
| 613 | 1323529.1 | Dill cryptic virus 2 isolate IPP_hortorum | Viruses | 1.53 |
| 614 | 1499684.1 | Clostridium sp. CL-2 | Bacteria | 1.53 |
| 615 | 1907654.1 | Collinsella sp. Marseille-P3296T strain Marseille-P3296 | Bacteria | 1.52 |
| 616 | 1834196.1 | Lachnoclostridium sp. YL32 sp. YL32 | Bacteria | 1.52 |
| 617 | 90371.26 | Salmonella enterica subsp. enterica serovar Typhimurium strain CFSAN033887 | Bacteria | 1.52 |
| 618 | 658665.1 | Dorea sp. D27 | Bacteria | 1.51 |
| 619 | 552396.1 | Erysipelotrichaceae bacterium 5_2_54FAA | Bacteria | 1.51 |
| 620 | 755172.1 | Peptoniphilus coxii strain DNF00729 | Bacteria | 1.50 |
| 621 | 1602170.1 | Prevotella sp. P5-60 | Bacteria | 1.49 |
| 622 | 1531.2 | [Clostridium] clostridioforme strain ATCC | Bacteria | 1.49 |
| 623 | 679190 | Prevotella buccalis ATCC 35310 | Bacteria | 1.49 |
| 624 | 33760.1 | Prune dwarf virus | Viruses | 1.49 |
| 625 | 1714570.1 | Blueberry shoestring virus | Viruses | 1.48 |
| 626 | 165432.1 | Cucumber leaf spot virus | Viruses | 1.47 |
| 627 | 1111120.1 | Acidaminococcus sp. BV3L6 | Bacteria | 1.46 |
| 628 | 818.3 | Bacteroides thetaiotaomicron strain 2789STDY5834846 | Bacteria | 1.46 |
| 629 | 1122989 | Prevotella oris DSM 18711 = JCM 12252 | Bacteria | 1.45 |
| 630 | 1352.16 | Enterococcus faecium isolate Hp_5-10 | Bacteria | 1.45 |
| 631 | 626522 | Alloprevotella tannerae ATCC 51259 | Bacteria | 1.43 |
| 632 | 1841857.1 | Culturomica massiliensis strain Marseille-P2698 | Bacteria | 1.40 |
| 633 | 1352.152 | Enterococcus faecium isolate Hp_76-7 | Bacteria | 1.39 |
| 634 | 28901.45 | Salmonella enterica strain NGUA07 | Bacteria | 1.39 |
| 635 | 742816 | Megamonas funiformis YIT 11815 | Bacteria | 1.36 |
| 636 | 1797112.1 | Olsenella sp. kh2p3 sp. kh2p3 | Bacteria | 1.34 |
| 637 | 1339274 | Bacteroides fragilis str. A7 (UDC12-2) | Bacteria | 1.32 |
| 638 | 1120921 | Acidaminococcus intestini DSM 21505 | Bacteria | 1.32 |
| 639 | 1917878.1 | Prevotella ihumii sp. Marseille-P3385 | Bacteria | 1.30 |
| 640 | 28901.37 | Salmonella enterica strain NGUA04 | Bacteria | 1.30 |
| 641 | 1739517.1 | Bacteroides sp. HMSC073E02 | Bacteria | 1.30 |
| 642 | 1432052.4 | Eisenbergiella tayi strain NML140904 | Bacteria | 1.29 |
| 643 | 999406 | [Clostridium] clostridioforme 90A6 | Bacteria | 1.29 |
| 644 | 641112 | Ruminococcus flavefaciens FD-1 | Bacteria | 1.29 |
| 645 | 1105029.1 | Actinomyces sp. ICM39 | Bacteria | 1.29 |
| 646 | 1280705 | Prevotella bryantii C21a | Bacteria | 1.27 |
| 647 | 84024.1 | Clostridium disporicum strain 2789STDY5834855 | Bacteria | 1.27 |
| 648 | 1284775.1 | Prevotella sp. S7-1-8 | Bacteria | 1.26 |
| 649 | 1581179.1 | Clostridium sp. HMSC19A11 | Bacteria | 1.25 |
| 650 | 1339308 | Bacteroides fragilis str. 3774 T13 | Bacteria | 1.25 |
| 651 | 1423719 | Lactobacillus algidus DSM 15638 | Bacteria | 1.24 |
| 652 | 1078091.1 | Coprococcus sp. HPP0048 | Bacteria | 1.24 |
| 653 | 1211843 | Candidatus Soleaferrea massiliensis AP7 | Bacteria | 1.24 |
| 654 | 1739408.1 | Streptococcus sp. HMSC072G04 | Bacteria | 1.23 |
| 655 | 469616 | Fusobacterium mortiferum ATCC 9817 | Bacteria | 1.22 |

| | | | | |
|---|---|---|---|---|
| 656 | 904144 | Gardnerella vaginalis 101 | Bacteria | 1.21 |
| 657 | 596315 | Peptostreptococcus stomatis DSM 17678 | Bacteria | 1.21 |
| 658 | 1739304.1 | Anaerosphaera sp. HMSC064C01 | Bacteria | 1.20 |
| 659 | 1870997.1 | Mogibacterium sp. Marseille-P3115 sp. Marseille-P3115 | Bacteria | 1.20 |
| 660 | 1232428.1 | Megasphaera massiliensis strain NP3 | Bacteria | 1.19 |
| 661 | 658082.1 | Lachnospiraceae bacterium 2_1_58FAA | Bacteria | 1.19 |
| 662 | 1042156.1 | Clostridium sp. SY8519 | Bacteria | 1.19 |
| 663 | 42817.1 | Corynebacterium argentoratense strain CNM | Bacteria | 1.19 |
| 664 | 1261636.1 | Anaerostipes sp. 494a | Bacteria | 1.17 |
| 665 | 1697788.1 | Clostridia bacterium UC5.1-2H6 | Bacteria | 1.16 |
| 666 | 817.14 | Bacteroides fragilis strain 2-078382-3 | Bacteria | 1.16 |
| 667 | 1120943 | Actinomyces gerencseriae DSM 6844 | Bacteria | 1.16 |
| 668 | 1260.1 | Finegoldia magna strain GED7760A | Bacteria | 1.16 |
| 669 | 411466 | Actinomyces odontolyticus ATCC 17982 | Bacteria | 1.16 |
| 670 | 1122975 | Porphyromonas somerae DSM 23386 | Bacteria | 1.16 |
| 671 | 500635 | Mitsuokella multacida DSM 20544 | Bacteria | 1.16 |
| 672 | 1401073 | Prevotella melaninogenica DNF00666 | Bacteria | 1.11 |
| 673 | 12175.6 | Apple chlorotic leaf spot virus | Viruses | 1.11 |
| 674 | 857290 | Scardovia wiggsiae F0424 | Bacteria | 1.08 |
| 675 | 585502 | Prevotella bergensis DSM 17361 | Bacteria | 1.08 |
| 676 | 1851429.1 | Christensenella sp. AF73-05CM02 | Bacteria | 1.07 |
| 677 | 1491.75 | Clostridium botulinum strain KAPB-3 | Bacteria | 1.07 |
| 678 | 137838.1 | Clostridium neonatale | Bacteria | 1.06 |
| 679 | 11613 | Tomato spotted wilt virus | Viruses | 1.05 |
| 680 | 1408895.1 | Dill cryptic virus 1 isolate IPP_hortorum | Viruses | 1.05 |
| 681 | 411472 | [Clostridium] symbiosum ATCC 14940 | Bacteria | 1.05 |
| 682 | 1410649 | Blautia schinkii DSM 10518 | Bacteria | 1.05 |
| 683 | 568816 | Acidaminococcus intestini RyC-MR95 | Bacteria | 1.03 |
| 684 | 1121334 | [Clostridium] sporosphaeroides DSM 1294 | Bacteria | 1.03 |
| 685 | 1680.2 | Bifidobacterium adolescentis strain Km | Bacteria | 1.03 |
| 686 | 243563.3 | Strawberry necrotic shock virus | Viruses | 1.01 |
| 687 | 198589.1 | Beet western yellows ST9 associated virus | Viruses | 1.01 |
| 688 | 1122976 | Porphyromonas uenonis DSM 23387 = JCM 13868 | Bacteria | 1.01 |
| 689 | 862962 | Bacteroides fragilis 638R | Bacteria | 1.01 |
| 690 | 679189 | Prevotella timonensis CRIS 5C-B1 | Bacteria | 1.00 |
| 691 | 297352 | Lactococcus piscium MKFS47 | Bacteria | 0.99 |
| 692 | 1834207.1 | Erysipelotrichaceae bacterium I46 | Bacteria | 0.99 |
| 693 | 666.83 | Vibrio cholerae strain MZO-2 | Bacteria | 0.98 |
| 694 | 791161 | Enterococcus faecium PC4.1 | Bacteria | 0.98 |
| 695 | 562982 | Gemella morbillorum M424 | Bacteria | 0.97 |
| 696 | 1280685.1 | Butyrivibrio sp. NC3005 | Bacteria | 0.97 |
| 697 | 562.188 | Escherichia coli strain USVAST406 | Bacteria | 0.96 |
| 698 | 457398.1 | Desulfovibrio sp. 3_1_syn3 | Bacteria | 0.95 |
| 699 | 1122982 | Prevotella denticola DSM 20614 = JCM 13449 | Bacteria | 0.95 |
| 700 | 295405 | Bacteroides fragilis YCH46 | Bacteria | 0.94 |
| 701 | 246199 | Ruminococcus albus 8 | Bacteria | 0.94 |

| | | | | |
|---|---|---|---|---|
| 702 | 28128.1 | Prevotella corporis strain MJR7716 | Bacteria | 0.93 |
| 703 | 1491.58 | Clostridium botulinum strain DB-2 | Bacteria | 0.92 |
| 704 | 556261.1 | Clostridium sp. D5 | Bacteria | 0.92 |
| 705 | 1414720.1 | Clostridium saudiense strain JCC | Bacteria | 0.91 |
| 706 | 1401062 | Prevotella timonensis S9-PR14 | Bacteria | 0.91 |
| 707 | 1491.57 | Clostridium botulinum strain 713_CBOT | Bacteria | 0.89 |
| 708 | 1329795.1 | Clostridiaceae bacterium MS3 | Bacteria | 0.89 |
| 709 | 817.26 | Bacteroides fragilis strain S14 | Bacteria | 0.88 |
| 710 | 547043 | Bifidobacterium pseudocatenulatum DSM 20438 = JCM 1200 = LMG 10505 | Bacteria | 0.88 |
| 711 | 270498.2 | Catabacter hongkongensis strain HKU16 | Bacteria | 0.88 |
| 712 | 311413.2 | Lettuce big-vein associated virus isolate Ls302 | Viruses | 0.86 |
| 713 | 1528099.1 | Lawsonella clevelandensis | Bacteria | 0.84 |
| 714 | 1111134.1 | Peptoniphilus sp. BV3C26 | Bacteria | 0.83 |
| 715 | 525282 | Finegoldia magna ATCC 53516 | Bacteria | 0.83 |
| 716 | 1235792.1 | Lachnospiraceae bacterium M18-1 | Bacteria | 0.83 |
| 717 | 90371.43 | Salmonella enterica subsp. enterica serovar Typhimurium strain CFSAN033867 | Bacteria | 0.82 |
| 718 | 559292 | Saccharomyces cerevisiae S288C | Eukaryota | 0.80 |
| 719 | 752555 | Prevotella bryantii B14 | Bacteria | 0.79 |
| 720 | 1352.155 | Enterococcus faecium isolate Hp_6-10 | Bacteria | 0.79 |
| 721 | 1308.7 | Streptococcus thermophilus | Bacteria | 0.79 |
| 722 | 1778580.3 | Nectarine virus M isolate NeVM/12P42 | Viruses | 0.78 |
| 723 | 999425.1 | Streptococcus sp. F0442 | Bacteria | 0.78 |
| 724 | 562.2364 | Escherichia coli strain ICBECAM7 | Bacteria | 0.77 |
| 725 | 729.12 | Haemophilus parainfluenzae strain 146_HPAR | Bacteria | 0.77 |
| 726 | 1504822.1 | bacterium OL-1 | Bacteria | 0.77 |
| 727 | 1151426 | Clostridioides difficile P51 | Bacteria | 0.77 |
| 728 | 762967 | Sutterella parvirubra YIT 11816 | Bacteria | 0.76 |
| 729 | 199.1 | Campylobacter concisus strain RMIT-JF1 | Bacteria | 0.76 |
| 730 | 1232448.1 | Clostridiales bacterium VE202-07 | Bacteria | 0.75 |
| 731 | 378833.1 | Sowbane mosaic virus | Viruses | 0.75 |
| 732 | 877411.1 | Ruminococcus sp. NK3A76 | Bacteria | 0.74 |
| 733 | 370354 | Entamoeba dispar SAW760 | Eukaryota | 0.74 |
| 734 | 596327 | Porphyromonas uenonis 60-3 | Bacteria | 0.73 |
| 735 | 470.677 | Acinetobacter baumannii strain XH753 | Bacteria | 0.73 |
| 736 | 70177.2 | Grapevine leafroll-associated virus 4 | Viruses | 0.72 |
| 737 | 53442.1 | Eubacterium callanderi strain FD | Bacteria | 0.72 |
| 738 | 1378168.1 | Firmicutes bacterium ASF500 | Bacteria | 0.71 |
| 739 | 861450 | Anaeroglobus geminatus F0357 | Bacteria | 0.71 |
| 740 | 1496.426 | Clostridioides difficile strain VRECD0128 | Bacteria | 0.70 |
| 741 | 42680.1 | Spinach latent virus | Viruses | 0.70 |
| 742 | 1339346 | Bacteroides ovatus str. 3725 D1 iv | Bacteria | 0.70 |
| 743 | 592010 | Abiotrophia defectiva ATCC 49176 | Bacteria | 0.69 |
| 744 | 562.765 | Escherichia coli isolate 14 | Bacteria | 0.69 |
| 745 | 1236498 | Bacteroides paurosaccharolyticus JCM 15092 | Bacteria | 0.69 |
| 746 | 1583.2 | Weissella confusa strain MBF8-1 | Bacteria | 0.68 |
| 747 | 1235811 | Prevotella disiens JCM 6334 = ATCC 29426 | Bacteria | 0.68 |

| | | | | |
|---|---|---|---|---|
| 748 | 1110546.1 | Veillonella tobetsuensis strain ATCC | Bacteria | 0.67 |
| 749 | 1235813 | Bacteroides pyogenes JCM 10003 | Bacteria | 0.67 |
| 750 | 1437612 | Bifidobacterium stercoris JCM 15918 | Bacteria | 0.67 |
| 751 | 1410665 | Mitsuokella jalaludinii DSM 13811 | Bacteria | 0.66 |
| 752 | 1673717.1 | Anaeromassilibacillus senegalensis strain mt9 | Bacteria | 0.66 |
| 753 | 28126.1 | Prevotella buccae strain 1205_PDEN | Bacteria | 0.66 |
| 754 | 1352.149 | Enterococcus faecium isolate Hp_22-12 | Bacteria | 0.65 |
| 755 | 1203555.1 | Acidaminococcus sp. HPA0509 | Bacteria | 0.65 |
| 756 | 1512.1 | [Clostridium] symbiosum strain 2789STDY5834864 | Bacteria | 0.65 |
| 757 | 1264.1 | Ruminococcus albus strain AR67 | Bacteria | 0.65 |
| 758 | 1776081.1 | Megasphaera sp. DISK 18 | Bacteria | 0.65 |
| 759 | 666.79 | Vibrio cholerae strain 3272-78 | Bacteria | 0.64 |
| 760 | 1122983 | Prevotella falsenii DSM 22864 = JCM 15124 | Bacteria | 0.64 |
| 761 | 60920.1 | Sanguibacter keddieii strain 250_SKED | Bacteria | 0.64 |
| 762 | 1588755.1 | Parvimonas sp. KA00067 | Bacteria | 0.63 |
| 763 | 592031 | Eubacterium saphenum ATCC 49989 | Bacteria | 0.62 |
| 764 | 698950 | Gardnerella vaginalis 284V | Bacteria | 0.62 |
| 765 | 478749 | Marvinbryantia formatexigens DSM 14469 | Bacteria | 0.61 |
| 766 | 28901.31 | Salmonella enterica strain NGUA25 | Bacteria | 0.61 |
| 767 | 411481 | Bifidobacterium adolescentis L2-32 | Bacteria | 0.61 |
| 768 | 187101.1 | Sneathia amnii sp. Sn35 | Bacteria | 0.60 |
| 769 | 1173061.1 | Geotrichum candidum strain CLIB | Eukaryota | 0.60 |
| 770 | 1577792.1 | Terrisporobacter othiniensis strain 08-306576 | Bacteria | 0.60 |
| 771 | 997895 | [Clostridium] bolteae 90B3 | Bacteria | 0.60 |
| 772 | 575611 | Prevotella buccae D17 | Bacteria | 0.60 |
| 773 | 1352.154 | Enterococcus faecium isolate Hp_21-11 | Bacteria | 0.59 |
| 774 | 1339277 | Bacteroides fragilis str. DS-71 | Bacteria | 0.59 |
| 775 | 1073375 | Ruminococcus gnavus CC55_001C | Bacteria | 0.59 |
| 776 | 817.4 | Bacteroides fragilis strain DCMOUH0017B | Bacteria | 0.59 |
| 777 | 1339278 | Bacteroides fragilis str. DS-166 | Bacteria | 0.58 |
| 778 | 762963 | Actinomyces sp. oral taxon 170 str. F0386 | Bacteria | 0.58 |
| 779 | 1261637.1 | Anaerostipes sp. 992a | Bacteria | 0.57 |
| 780 | 1035195 | Corynebacterium durum F0235 | Bacteria | 0.57 |
| 781 | 679191 | Prevotella amnii CRIS 21A-A | Bacteria | 0.57 |
| 782 | 500633 | [Clostridium] hiranonis DSM 13275 | Bacteria | 0.57 |
| 783 | 28026.6 | B29 | Bacteria | 0.56 |
| 784 | 1679.14 | Bifidobacterium longum subsp. longum strain CCUG30698 | Bacteria | 0.56 |
| 785 | 12042.2 | Beet western yellows virus | Viruses | 0.56 |
| 786 | 1292034 | Caulobacter crescentus OR37 | Bacteria | 0.56 |
| 787 | 31722.2 | Blueberry scorch virus isolate BC-2 | Viruses | 0.55 |
| 788 | 1121097 | Bacteroides graminisolvens DSM 19988 = JCM 15093 | Bacteria | 0.55 |
| 789 | 1321775 | Actinomyces sp. oral taxon 172 str. F0311 | Bacteria | 0.55 |
| 790 | 1007096 | Oscillibacter ruminantium GH1 | Bacteria | 0.54 |
| 791 | 1871336.1 | Criibacterium bergeronii strain CCRI-22567 | Bacteria | 0.54 |
| 792 | 1105030.1 | Actinomyces sp. ICM58 | Bacteria | 0.54 |

| | | | | |
|---|---|---|---|---|
| 793 | 1313215 | Aichi virus 1 | Viruses | 0.54 |
| 794 | 1122992 | Prevotella timonensis 4401737 = DSM 22865 = JCM 15640 | Bacteria | 0.53 |
| 795 | 525362 | Lactobacillus ruminis ATCC 25644 | Bacteria | 0.53 |
| 796 | 12451.1 | Raspberry bushy dwarf virus | Viruses | 0.53 |
| 797 | 699246 | Mageeibacillus indolicus UPII9-5 | Bacteria | 0.53 |
| 798 | 1720195.1 | Gabonibacter massiliensis strain GM7 | Bacteria | 0.53 |
| 799 | 1393034.1 | Atopobium deltae strain DNF00019 | Bacteria | 0.51 |
| 800 | 77095.1 | Prevotella bryantii strain KHPX14 | Bacteria | 0.51 |
| 801 | 1697790.1 | Clostridia bacterium UC5.1-1C12 | Bacteria | 0.51 |
| 802 | 322505.1 | Sharpea azabuensis strain DSM | Bacteria | 0.50 |
| 803 | 84024.2 | Clostridium disporicum strain 2789STDY5608827 | Bacteria | 0.50 |
| 804 | 1352.15 | Enterococcus faecium isolate Hp_23-9 | Bacteria | 0.50 |
| 805 | 1265.2 | Ruminococcus flavefaciens strain Y1 | Bacteria | 0.50 |
| 806 | 1736.3 | Eubacterium limosum strain SA11 | Bacteria | 0.50 |
| 807 | 77095.2 | Prevotella bryantii strain FB3001 | Bacteria | 0.50 |
| 808 | 1051631.1 | Streptococcus phage YMC-2011 | Viruses | 0.50 |
| 809 | 1532180.1 | Penicillium roqueforti ssRNA mycovirus 1 strain PRG42-7 | Viruses | 0.50 |
| 810 | 1852386.1 | Olsenella sp. Marseille-P2912 sp. Marseille-P2912 | Bacteria | 0.49 |
| 811 | 1161902 | Eubacterium nodatum ATCC 33099 | Bacteria | 0.49 |
| 812 | 1111135.1 | Coriobacteriaceae bacterium BV3Ac1 | Bacteria | 0.49 |
| 813 | 556270.1 | Coprobacillus sp. D7 | Bacteria | 0.48 |
| 814 | 1531429.1 | Coriobacteriaceae bacterium 68-1-3 | Bacteria | 0.48 |
| 815 | 1339292 | Bacteroides fragilis str. 1007-1-F #7 | Bacteria | 0.48 |
| 816 | 1232446.1 | Clostridiales bacterium VE202-18 | Bacteria | 0.48 |
| 817 | 1739529.1 | Porphyromonas sp. HMSC077F02 | Bacteria | 0.48 |
| 818 | 11987.2 | Melon necrotic spot virus | Viruses | 0.47 |
| 819 | 1401074 | Prevotella buccalis DNF00853 | Bacteria | 0.47 |
| 820 | 1805478.1 | Olsenella sp. Marseille-P2300 | Bacteria | 0.47 |
| 821 | 817.15 | Bacteroides fragilis strain 86-5443-2-2 | Bacteria | 0.47 |
| 822 | 642478.1 | Lettuce chlorosis virus | Viruses | 0.47 |
| 823 | 1697792.1 | Clostridia bacterium UC5.1-2F7 | Bacteria | 0.47 |
| 824 | 1680.1 | Bifidobacterium adolescentis strain 2789STDY5608862 | Bacteria | 0.47 |
| 825 | 1852374.1 | Ezakiella massiliensis strain Marseille-P2951T sp. Marseille-P2951 | Bacteria | 0.47 |
| 826 | 66200.1 | Carrot red leaf virus | Viruses | 0.47 |
| 827 | 585501 | Oribacterium sinus F0268 | Bacteria | 0.46 |
| 828 | 888050 | Actinomyces cardiffensis F0333 | Bacteria | 0.46 |
| 829 | 608534 | Oribacterium sp. oral taxon 078 str. F0262 | Bacteria | 0.46 |
| 830 | 877414.1 | Clostridiales bacterium NK3B98 | Bacteria | 0.46 |
| 831 | 1852375.1 | Acidaminococcus massiliensis strain Marseille-P2828 | Bacteria | 0.45 |
| 832 | 28901.13 | Salmonella enterica strain NGUA10 | Bacteria | 0.45 |
| 833 | 28026.3 | Bifidobacterium pseudocatenulatum strain CA-K29a | Bacteria | 0.45 |
| 834 | 580254.1 | Raphanus sativus cryptic virus 3 | Viruses | 0.45 |

| | | | | |
|---|---|---|---|---|
| 835 | 483214 | Methanobrevibacter smithii DSM 2375 | Archaea | 0.44 |
| 836 | 742741 | [Clostridium] symbiosum WAL-14673 | Bacteria | 0.44 |
| 837 | 729.1 | Haemophilus parainfluenzae strain 215035-2-ISO5 | Bacteria | 0.44 |
| 838 | 1351.95 | Enterococcus faecalis isolate Hp_74-d1 | Bacteria | 0.44 |
| 839 | 1680.5 | Bifidobacterium adolescentis strain 22L | Bacteria | 0.44 |
| 840 | 1522.2 | 2789STDY5834853 | Bacteria | 0.44 |
| 841 | 1673723.1 | Murdochiella massiliensis strain SIT12 | Bacteria | 0.43 |
| 842 | 216816.7 | Bifidobacterium longum isolate Bifido_09 | Bacteria | 0.43 |
| 843 | 1392494.1 | Lachnospiraceae bacterium AC2012 | Bacteria | 0.43 |
| 844 | 1392836.1 | Lachnospiraceae bacterium TWA4 | Bacteria | 0.43 |
| 845 | 633697 | [Eubacterium] cellulosolvens 6 | Bacteria | 0.42 |
| 846 | 1226323.1 | Oscillibacter sp. KLE 1745 | Bacteria | 0.42 |
| 847 | 290055.1 | [Eubacterium] fissicatena strain KCTC | Bacteria | 0.42 |
| 848 | 431317.1 | Triticum mosaic virus | Viruses | 0.42 |
| 849 | 1211844 | Candidatus Stoquefichus massiliensis AP9 | Bacteria | 0.42 |
| 850 | 12433.1 | Garlic virus A | Viruses | 0.42 |
| 851 | 1321819 | Bacteroides pyogenes F0041 | Bacteria | 0.41 |
| 852 | 1232445.1 | Clostridiales bacterium VE202-16 | Bacteria | 0.41 |
| 853 | 1365967 | Bifidobacterium breve MCC 1454 | Bacteria | 0.40 |
| 854 | 997894 | [Clostridium] bolteae 90A9 | Bacteria | 0.40 |
| 855 | 575593 | Lachnospiraceae oral taxon 107 str. F0167 | Bacteria | 0.40 |
| 856 | 1048332 | Streptococcus salivarius CCHSS3 | Bacteria | 0.40 |
| 857 | 1352.156 | Enterococcus faecium isolate Hp_24-3 | Bacteria | 0.39 |
| 858 | 1336250 | Hallella seregens ATCC 51272 | Bacteria | 0.39 |
| 859 | 1033744 | Peptoniphilus senegalensis JC140 | Bacteria | 0.39 |
| 860 | 1035196 | Peptostreptococcus anaerobius VPI 4330 = DSM 2949 | Bacteria | 0.39 |
| 861 | 31722.3 | Blueberry scorch virus | Viruses | 0.39 |
| 862 | 457404 | Fusobacterium ulcerans 12-1B | Bacteria | 0.39 |
| 863 | 827.1 | Campylobacter ureolyticus strain CIT007 | Bacteria | 0.39 |
| 864 | 1776177.1 | Cucumis melo endornavirus isolate CL-01 | Viruses | 0.39 |
| 865 | 1215078 | Clostridioides difficile E7 | Bacteria | 0.38 |
| 866 | 457396.1 | Clostridium sp. 7_2_43FAA | Bacteria | 0.38 |
| 867 | 1489836.1 | Atlantic salmon calicivirus isolate Nordland/2011 | Viruses | 0.38 |
| 868 | 97478.5 | Lactobacillus mucosae strain DPC | Bacteria | 0.37 |
| 869 | 1581061.1 | Abiotrophia sp. HMSC24B09 | Bacteria | 0.37 |
| 870 | 12187.2 | Strawberry mild yellow edge virus | Viruses | 0.37 |
| 871 | 103722.1 | Grapevine fleck virus | Viruses | 0.37 |
| 872 | 1341157 | Ruminococcus flavefaciens 007c | Bacteria | 0.36 |
| 873 | 5007.1 | Brettanomyces bruxellensis | Eukaryota | 0.36 |
| 874 | 1676617.1 | Ralstonia sp. MD27 | Bacteria | 0.35 |
| 875 | 1499683.1 | Clostridium sp. CL-6 | Bacteria | 0.35 |
| 876 | 31722.1 | Blueberry scorch virus isolate BC-1 | Viruses | 0.35 |
| 877 | 469378 | Cryptobacterium curtum DSM 15641 | Bacteria | 0.35 |
| 878 | 883109 | Eubacterium infirmum F0142 | Bacteria | 0.35 |
| 879 | 1321814 | Eubacterium brachy ATCC 33089 | Bacteria | 0.34 |
| 880 | 154288.2 | Turicibacter sanguinis strain 2789STDY5608821 | Bacteria | 0.34 |
| 881 | 1522.1 | [Clostridium] innocuum strain AN88 | Bacteria | 0.34 |

| | | | | |
|---|---|---|---|---|
| 882 | 888833 | Streptococcus australis ATCC 700641 | Bacteria | 0.33 |
| 883 | 397544.4 | Squash vein yellowing virus isolate IL | Viruses | 0.33 |
| 884 | 352165 | Pyramidobacter piscolens W5455 | Bacteria | 0.33 |
| 885 | 908340.1 | Clostridium sp. HGF2 | Bacteria | 0.33 |
| 886 | 145856.1 | Human picobirnavirus | Viruses | 0.33 |
| 887 | 1561.1 | Clostridium baratii strain XCM | Bacteria | 0.32 |
| 888 | 887898 | Lautropia mirabilis ATCC 51599 | Bacteria | 0.32 |
| 889 | 1282887 | Lachnospira multipara ATCC 19207 | Bacteria | 0.32 |
| 890 | 642492 | Clostridium lentocellum DSM 5427 | Bacteria | 0.32 |
| 891 | 688246 | Prevotella multisaccharivorax DSM 17128 | Bacteria | 0.32 |
| 892 | 1680.7 | Bifidobacterium adolescentis strain 150 | Bacteria | 0.32 |
| 893 | 1391469 | Enterococcus faecalis BM4654 | Bacteria | 0.31 |
| 894 | 1284680.1 | Actinomyces sp. S6-Spd3 | Bacteria | 0.31 |
| 895 | 1203606 | Butyricicoccus pullicaecorum 1.2 | Bacteria | 0.31 |
| 896 | 1125712 | Olsenella profusa F0195 | Bacteria | 0.31 |
| 897 | 1776390.1 | Peptoniphilus sp. KHD5 sp. KHD5 | Bacteria | 0.30 |
| 898 | 1114972 | Lactobacillus rossiae DSM 15814 | Bacteria | 0.30 |
| 899 | 742767 | Dysgonomonas mossii DSM 22836 | Bacteria | 0.30 |
| 900 | 1219626.1 | Peptostreptococcus sp. MV1 | Bacteria | 0.30 |
| 901 | 1304.27 | Streptococcus salivarius strain KB005 | Bacteria | 0.30 |
| 902 | 1339270 | Bacteroides fragilis str. I1345 | Bacteria | 0.30 |
| 903 | 1280702 | Bifidobacterium longum AGR2137 | Bacteria | 0.30 |
| 904 | 1121485 | Dysgonomonas capnocytophagoides DSM 22835 | Bacteria | 0.30 |
| 905 | 329.3 | Ralstonia pickettii strain CW2 | Bacteria | 0.29 |
| 906 | 1304.11 | Streptococcus salivarius strain 37-08 | Bacteria | 0.29 |
| 907 | 1458465 | Megasphaera elsdenii 14-14 | Bacteria | 0.29 |
| 908 | 32625.1 | Mushroom bacilliform virus | Viruses | 0.29 |
| 909 | 796942.1 | Stomatobaculum longum | Bacteria | 0.29 |
| 910 | 216816.1 | Bifidobacterium longum isolate Bifido_03 | Bacteria | 0.29 |
| 911 | 12282.1 | Tobacco ringspot virus | Viruses | 0.29 |
| 912 | 768727 | Veillonella parvula ACS-068-V-Sch12 | Bacteria | 0.29 |
| 913 | 762983 | Succinatimonas hippei YIT 12066 | Bacteria | 0.29 |
| 914 | 28026.7 | Bifidobacterium pseudocatenulatum strain 2789STDY5834840 | Bacteria | 0.29 |
| 915 | 1125718 | Actinomyces massiliensis F0489 | Bacteria | 0.29 |
| 916 | 1871012.1 | Mobilibacterium timonense strain Marseille-P3194 | Bacteria | 0.29 |
| 917 | 1121864 | Enterococcus cecorum DSM 20682 = ATCC 43198 | Bacteria | 0.28 |
| 918 | 29466.2 | Veillonella parvula strain UTDB1-3 | Bacteria | 0.28 |
| 919 | 193118.1 | Lucerne transient streak virus satellite RNA | Viruses | 0.28 |
| 920 | 1689303.1 | Lagierella massiliensis strain SIT14 | Bacteria | 0.28 |
| 921 | 1410644 | Bifidobacterium adolescentis DSM 20087 | Bacteria | 0.28 |
| 922 | 39046.1 | Cassava common mosaic virus | Viruses | 0.28 |
| 923 | 595.15 | Salmonella enterica subsp. enterica serovar Infantis strain 120100 | Bacteria | 0.28 |
| 924 | 649761 | Prevotella veroralis F0319 | Bacteria | 0.28 |
| 925 | 1408439 | Fusobacterium perfoetens ATCC 29250 | Bacteria | 0.28 |

| | | | | |
|---|---|---|---|---|
| 926 | 1680.3 | Bifidobacterium adolescentis strain 2789STDY5834850 | Bacteria | 0.28 |
| 927 | 1739305.1 | Peptoniphilus sp. HMSC062D09 | Bacteria | 0.28 |
| 928 | 1095750 | Lachnoanaerobaculum saburreum F0468 | Bacteria | 0.28 |
| 929 | 354328.3 | Bell pepper endornavirus | Viruses | 0.28 |
| 930 | 1229758 | Leuconostoc carnosum JB16 | Bacteria | 0.28 |
| 931 | 12282.2 | Tobacco ringspot virus isolate SK | Viruses | 0.28 |
| 932 | 563031.1 | Prevotella sp. C561 | Bacteria | 0.28 |
| 933 | 1680.6 | Bifidobacterium adolescentis strain BBMN23 | Bacteria | 0.28 |
| 934 | 1122150 | Lactobacillus nagelii DSM 13675 | Bacteria | 0.28 |
| 935 | 1588754.1 | Veillonellaceae bacterium DNF00626 | Bacteria | 0.28 |
| 936 | 859.1 | Fusobacterium necrophorum strain ATCC | Bacteria | 0.27 |
| 937 | 566552 | Bifidobacterium catenulatum DSM 16992 = JCM 1194 = LMG 11043 | Bacteria | 0.27 |
| 938 | 1871017.1 | Peptoniphilus urinimassiliensis strain Marseille-P3195 | Bacteria | 0.27 |
| 939 | 1679.8 | Bifidobacterium longum subsp. longum strain LO-K29a | Bacteria | 0.27 |
| 940 | 198599.1 | Saccharomyces 23S RNA narnavirus | Viruses | 0.27 |
| 941 | 1339315 | Bacteroides fragilis str. 3988T(B)14 | Bacteria | 0.27 |
| 942 | 54062.1 | Pediococcus parvulus strain 2.6 | Bacteria | 0.27 |
| 943 | 1338.1 | Streptococcus intermedius strain 631_SCON | Bacteria | 0.27 |
| 944 | 905.1 | Acidaminococcus fermentans strain pGA-4 | Bacteria | 0.27 |
| 945 | 1161745 | Bifidobacterium longum subsp. longum 2-2B | Bacteria | 0.26 |
| 946 | 29833.1 | Hanseniaspora uvarum | Eukaryota | 0.26 |
| 947 | 1120944 | Actinomyces israelii DSM 43320 | Bacteria | 0.26 |
| 948 | 1122216 | Megamonas hypermegale DSM 1672 | Bacteria | 0.26 |
| 949 | 1852387.1 | Streptococcus timonensis strain Marseille-P2915 | Bacteria | 0.26 |
| 950 | 706433 | Solobacterium moorei F0204 | Bacteria | 0.26 |
| 951 | 244362.1 | Ruminococcus sp. YE71 sp. YE71 | Bacteria | 0.25 |
| 952 | 1778580.1 | Nectarine virus M isolate NeVM/SF04522E | Viruses | 0.25 |
| 953 | 883156 | Veillonella seminalis ACS-216-V-Col6b | Bacteria | 0.25 |
| 954 | 1339272 | Bacteroides fragilis str. J38-1 | Bacteria | 0.25 |
| 955 | 1301220.1 | Citrus vein enation virus isolate VE-1 | Viruses | 0.25 |
| 956 | 997896 | [Clostridium] bolteae 90B7 | Bacteria | 0.25 |
| 957 | 40543.1 | Sneathia sanguinegens strain CCUG41628 | Bacteria | 0.25 |
| 958 | 450749.1 | Veillonella sp. 6_1_27 | Bacteria | 0.25 |
| 959 | 1449897.1 | Uncultured phage WW-nAnB strain 3 strain 3 | Viruses | 0.25 |
| 960 | 12145.1 | Tomato bushy stunt virus | Viruses | 0.24 |
| 961 | 1805477.1 | Clostridium sp. Marseille-P299 | Bacteria | 0.24 |
| 962 | 521095 | Atopobium parvulum DSM 20469 | Bacteria | 0.24 |
| 963 | 729.13 | Haemophilus parainfluenzae strain 1209_HPAR | Bacteria | 0.24 |
| 964 | 1463935.1 | Streptomyces sp. NRRL WC-3744 | Bacteria | 0.24 |
| 965 | 28026.1 | D29 | Bacteria | 0.24 |
| 966 | 1415630.1 | Pseudomonas sp. TKP sp. TKP | Bacteria | 0.23 |
| 967 | 39029.1 | Megasphaera cerevisiae strain NSB1 | Bacteria | 0.23 |
| 968 | 216816.9 | Bifidobacterium longum isolate Bifido_12 | Bacteria | 0.23 |
| 969 | 112227.1 | Cactus virus X | Viruses | 0.23 |

| | | | | |
|---|---|---|---|---|
| 970 | 29466.1 | Veillonella parvula strain DNF00876 | Bacteria | 0.23 |
| 971 | 1123311 | Streptococcus orisratti DSM 15617 | Bacteria | 0.23 |
| 972 | 12175.8 | Apple chlorotic leaf spot virus isolate QD-13 | Viruses | 0.23 |
| 973 | 1348633 | Lactococcus raffinolactis NBRC 100932 | Bacteria | 0.23 |
| 974 | 1040964 | Lactobacillus ruminis SPM0211 | Bacteria | 0.23 |
| 975 | 1496.114 | Clostridioides difficile isolate VL_0181 | Bacteria | 0.23 |
| 976 | 1423814 | Lactobacillus vaginalis DSM 5837 = ATCC 49540 | Bacteria | 0.22 |
| 977 | 714313 | Lactobacillus sanfranciscensis TMW 1.1304 | Bacteria | 0.22 |
| 978 | 29397.1 | Lactobacillus delbrueckii subsp. lactis | Bacteria | 0.22 |
| 979 | 887325 | Lachnoanaerobaculum saburreum DSM 3986 | Bacteria | 0.22 |
| 980 | 1074044.1 | uncultured phage WW-nAnB | Viruses | 0.22 |
| 981 | 469617 | Fusobacterium ulcerans ATCC 49185 | Bacteria | 0.22 |
| 982 | 1511761.1 | Leuconostoc mesenteroides subsp. suionicum strain DSM | Bacteria | 0.22 |
| 983 | 1125779 | Corynebacterium pyruviciproducens ATCC BAA-1742 | Bacteria | 0.22 |
| 984 | 1778.5 | Mycobacterium gordonae strain HMC_M15 | Bacteria | 0.22 |
| 985 | 1739251.1 | Fusobacterium sp. HMSC073F01 | Bacteria | 0.22 |
| 986 | 479436 | Veillonella parvula DSM 2008 | Bacteria | 0.21 |
| 987 | 397288.1 | Lachnospiraceae bacterium 3-1 | Bacteria | 0.21 |
| 988 | 563191.1 | Acidaminococcus sp. D21 | Bacteria | 0.21 |
| 989 | 646413.1 | Streptococcus phage 5093 | Viruses | 0.21 |
| 990 | 645512 | Jonquetella anthropi E3_33 E1 | Bacteria | 0.21 |
| 991 | 565040 | Bifidobacterium longum subsp. infantis 157F | Bacteria | 0.21 |
| 992 | 12263.1 | Squash mosaic virus isolate CH | Viruses | 0.21 |
| 993 | 35350.4 | Apple stem pitting virus | Viruses | 0.20 |
| 994 | 12317.6 | Tobacco streak virus isolate 1973 | Viruses | 0.20 |
| 995 | 243563.2 | Strawberry necrotic shock virus isolate Florida-4 | Viruses | 0.20 |
| 996 | 1776381.1 | Olsenella sp. KHD7 sp. KHD7 | Bacteria | 0.20 |
| 997 | 760570 | Streptococcus parasanguinis ATCC 15912 | Bacteria | 0.20 |
| 998 | 1653434.1 | Sellimonas intestinalis strain BR72 | Bacteria | 0.20 |
| 999 | 44008.8 | Enterococcus cecorum strain G-29 | Bacteria | 0.20 |
| 1000 | 322159 | Streptococcus thermophilus LMD-9 | Bacteria | 0.20 |
| 1001 | 537288.1 | Megasphaera sp. DJF_B143 | Bacteria | 0.20 |
| 1002 | 1583.1 | Weissella confusa strain DSM | Bacteria | 0.20 |
| 1003 | 1852372.1 | Varibaculum sp. Marseille-P2802 sp. Marseille-P2802 | Bacteria | 0.20 |
| 1004 | 907931 | Leuconostoc fallax KCTC 3537 | Bacteria | 0.20 |
| 1005 | 1261.3 | Peptostreptococcus anaerobius strain MJR8628A | Bacteria | 0.20 |
| 1006 | 1681.1 | Bifidobacterium bifidum strain 791 | Bacteria | 0.19 |
| 1007 | 938288.1 | Fenollaria massiliensis | Bacteria | 0.19 |
| 1008 | 270498.1 | Catabacter hongkongensis strain ABBA15k | Bacteria | 0.19 |
| 1009 | 634994 | Leptotrichia hofstadii F0254 | Bacteria | 0.19 |
| 1010 | 1501391.1 | Alistipes inops strain 627 | Bacteria | 0.19 |
| 1011 | 1216932.1 | Clostridium bornimense strain M2/40T | Bacteria | 0.19 |
| 1012 | 546262 | Neisseria cinerea ATCC 14685 | Bacteria | 0.19 |
| 1013 | 12242.1 | Tobacco mosaic virus | Viruses | 0.19 |
| 1014 | 12263.2 | Squash mosaic virus | Viruses | 0.19 |

| | | | | |
|---|---|---|---|---|
| 1015 | 1236497 | Prevotella oulorum JCM 14966 | Bacteria | 0.19 |
| 1016 | 596329 | Peptostreptococcus anaerobius 653-L | Bacteria | 0.18 |
| 1017 | 1720317.1 | Porphyromonadaceae bacterium FC4 | Bacteria | 0.18 |
| 1018 | 1236689 | Candidatus Methanomethylophilus alvus Mx1201 | Archaea | 0.18 |
| 1019 | 1871014.1 | Arcanobacterium urinimassiliense sp. Marseille-P3248 | Bacteria | 0.18 |
| 1020 | 1391466 | Enterococcus faecium NEF1 | Bacteria | 0.18 |
| 1021 | 1658108.1 | Niameybacter massiliensis | Bacteria | 0.18 |
| 1022 | 209529.6 | Aphid lethal paralysis virus isolate AP | Viruses | 0.18 |
| 1023 | 712122.1 | Actinomyces sp. oral taxon 414 strain F0588 | Bacteria | 0.18 |
| 1024 | 1432052.5 | Eisenbergiella tayi strain NML110678 | Bacteria | 0.18 |
| 1025 | 143387.2 | Fusobacterium necrophorum subsp. funduliforme strain LS_1272 | Bacteria | 0.18 |
| 1026 | 1449336 | Carnobacterium divergens DSM 20623 | Bacteria | 0.18 |
| 1027 | 1871002.1 | Acidaminococcus timonensis strain Marseille-P2764 | Bacteria | 0.18 |
| 1028 | 35350.7 | Apple stem pitting virus isolate apple | Viruses | 0.17 |
| 1029 | 1720204.1 | Collinsella ihuae sp. GD7 | Bacteria | 0.17 |
| 1030 | 1347366.1 | Clostridium sp. ND2 | Bacteria | 0.17 |
| 1031 | 1838287.1 | Gammaproteobacteria bacterium 2W06 | Bacteria | 0.17 |
| 1032 | 12458 | Garlic latent virus | Viruses | 0.17 |
| 1033 | 78448.1 | Bifidobacterium pullorum strain LMG | Bacteria | 0.17 |
| 1034 | 1871033.1 | Olsenella sp. Marseille-P3197 sp. Marseille-P3197 | Bacteria | 0.17 |
| 1035 | 1280689 | Clostridium paraputrificum AGR2156 | Bacteria | 0.17 |
| 1036 | 367928 | Bifidobacterium adolescentis ATCC 15703 | Bacteria | 0.17 |
| 1037 | 1073386 | Bacteroides fragilis HMW 610 | Bacteria | 0.17 |
| 1038 | 1497955.1 | Clostridiales bacterium KA00274 | Bacteria | 0.17 |
| 1039 | 1136138 | Pseudomonas fragi B25 | Bacteria | 0.17 |
| 1040 | 1050201 | Allobaculum stercoricanis DSM 13633 | Bacteria | 0.17 |
| 1041 | 525919 | Anaerococcus prevotii DSM 20548 | Bacteria | 0.17 |
| 1042 | 1105031.1 | Clostridium sp. MSTE9 | Bacteria | 0.17 |
| 1043 | 1680.4 | Bifidobacterium adolescentis strain 2789STDY5608824 | Bacteria | 0.17 |
| 1044 | 817.9 | Bacteroides fragilis strain 20793-3 | Bacteria | 0.17 |
| 1045 | 1852362.1 | Bacteroides ihuae strain Marseille-P2824 | Bacteria | 0.17 |
| 1046 | 1778.3 | Mycobacterium gordonae strain 1245752.6 | Bacteria | 0.17 |
| 1047 | 1304.5 | Streptococcus salivarius strain 726_SSAL | Bacteria | 0.17 |
| 1048 | 1739406.1 | Actinomyces sp. HMSC035G02 | Bacteria | 0.17 |
| 1049 | 1401075 | Prevotella disiens DNF00882 | Bacteria | 0.17 |
| 1050 | 1280674.1 | Prevotella sp. AGR2160 | Bacteria | 0.17 |
| 1051 | 1321782 | Oribacterium sp. oral taxon 078 str. F0263 | Bacteria | 0.17 |
| 1052 | 1167629 | 27673 | Bacteria | 0.17 |
| 1053 | 1497953.1 | Bacteroidales bacterium KA00251 | Bacteria | 0.17 |
| 1054 | 1805470.1 | Clostridium sp. Marseille-P2414 sp. Marseille-P2414 | Bacteria | 0.17 |
| 1055 | 28026.5 | Bifidobacterium pseudocatenulatum strain CA-K29b | Bacteria | 0.17 |

| | | | | |
|---|---|---|---|---|
| 1056 | 103724.1 | Grapevine asteroid mosaic-associated virus isolate GV30 | Viruses | 0.17 |
| 1057 | 936588.1 | Veillonella sp. ACP1 | Bacteria | 0.17 |
| 1058 | 927691 | Leuconostoc gelidum subsp. gelidum KCTC 3527 | Bacteria | 0.17 |
| 1059 | 287.9 | Pseudomonas aeruginosa strain Pae_CF67.12q | Bacteria | 0.17 |
| 1060 | 1715051.1 | Streptococcus sp. HMSC068F04 | Bacteria | 0.17 |
| 1061 | 205913 | Bifidobacterium longum DJO10A | Bacteria | 0.17 |
| 1062 | 1352.145 | Enterococcus faecium isolate Hp_7-8 | Bacteria | 0.16 |
| 1063 | 1561.3 | Clostridium baratii strain 2789STDY5834956 | Bacteria | 0.16 |
| 1064 | 1581080.1 | Streptococcus sp. HMSC10E12 | Bacteria | 0.16 |
| 1065 | 1111454.1 | Megasphaera sp. BV3C16-1 | Bacteria | 0.16 |
| 1066 | 742766 | Dysgonomonas gadei ATCC BAA-286 | Bacteria | 0.16 |
| 1067 | 1339288 | Bacteroides fragilis str. 3988 T1 | Bacteria | 0.16 |
| 1068 | 1032506.1 | Prevotella sp. MSX73 | Bacteria | 0.16 |
| 1069 | 1655645.1 | Parabacteroides phage YZ-2015b | Viruses | 0.16 |
| 1070 | 457416.1 | Veillonella sp. 3_1_44 | Bacteria | 0.16 |
| 1071 | 1118062 | Peptoniphilus obesi ph1 | Bacteria | 0.16 |
| 1072 | 1623.1 | Lactobacillus ruminis strain WC1T17 | Bacteria | 0.16 |
| 1073 | 1588753.1 | Coriobacteriales bacterium DNF00809 | Bacteria | 0.16 |
| 1074 | 546269 | Filifactor alocis ATCC 35896 | Bacteria | 0.16 |
| 1075 | 139208.1 | Isoptericola variabilis strain 871_IVAR | Bacteria | 0.16 |
| 1076 | 1203593.1 | Veillonella sp. HPA0037 | Bacteria | 0.16 |
| 1077 | 879243 | Porphyromonas asaccharolytica DSM 20707 | Bacteria | 0.15 |
| 1078 | 817.23 | Bacteroides fragilis strain BOB25 | Bacteria | 0.15 |
| 1079 | 867080.1 | Paenibacillus sp. IHB B 3415 | Bacteria | 0.15 |
| 1080 | 1720315.1 | Eggerthellaceae bacterium AT8 | Bacteria | 0.15 |
| 1081 | 1410673 | Selenomonas bovis 8-14-1 | Bacteria | 0.15 |
| 1082 | 525146 | Desulfovibrio desulfuricans subsp. desulfuricans str. ATCC 27774 | Bacteria | 0.15 |
| 1083 | 1352.101 | Enterococcus faecium strain XH877 | Bacteria | 0.15 |
| 1084 | 1423747 | Lactobacillus fuchuensis DSM 14340 = JCM 11249 | Bacteria | 0.15 |
| 1085 | 1302.13 | Streptococcus gordonii strain M5 | Bacteria | 0.15 |
| 1086 | 873513 | Prevotella buccae ATCC 33574 | Bacteria | 0.15 |
| 1087 | 184922 | Giardia lamblia ATCC 50803 | Eukaryota | 0.15 |
| 1088 | 1339306 | Bacteroides fragilis str. 3719 T6 | Bacteria | 0.15 |
| 1089 | 1871023.1 | Rikenella sp. Marseille-P3215 sp. Marseille-P3215 | Bacteria | 0.15 |
| 1090 | 548908.1 | Fig fleck-associated virus | Viruses | 0.15 |
| 1091 | 28347.12 | Apple stem grooving virus clone ASGVp12 | Viruses | 0.15 |
| 1092 | 883069 | Actinomyces europaeus ACS-120-V-Col10b | Bacteria | 0.15 |
| 1093 | 1293039 | Methanobrevibacter arboriphilus JCM 9315 | Archaea | 0.15 |
| 1094 | 206672 | Bifidobacterium longum NCC2705 | Bacteria | 0.15 |
| 1095 | 1206566.2 | Blueberry virus A isolate Elliot | Viruses | 0.14 |
| 1096 | 1161409.1 | Bifidobacterium sp. MSTE12 | Bacteria | 0.14 |
| 1097 | 1410621.1 | Lachnospiraceae bacterium AD3010 | Bacteria | 0.14 |
| 1098 | 742740 | [Clostridium] symbiosum WAL-14163 | Bacteria | 0.14 |
| 1099 | 12172.3 | Shallot latent virus isolate WA-1 | Viruses | 0.14 |
| 1100 | 112227.2 | Cactus virus X strain SCM51431 | Viruses | 0.14 |

| | | | | |
|---|---|---|---|---|
| 1101 | 1308.1 | Streptococcus thermophilus strain KLDS | Bacteria | 0.14 |
| 1102 | 1351.93 | Enterococcus faecalis isolate Hp_74-d5 | Bacteria | 0.14 |
| 1103 | 1629.2 | Weissella viridescens strain NCDO | Bacteria | 0.14 |
| 1104 | 1623.2 | Lactobacillus ruminis strain DPC | Bacteria | 0.14 |
| 1105 | 469618 | Fusobacterium varium ATCC 27725 | Bacteria | 0.14 |
| 1106 | 879305 | Anaerococcus prevotii ACS-065-V-Col13 | Bacteria | 0.14 |
| 1107 | 1052902 | Tobacco mosaic virus strain Ohio V | Viruses | 0.14 |
| 1108 | 1739525.1 | Peptoniphilus sp. HMSC075B08 | Bacteria | 0.14 |
| 1109 | 1401072 | Prevotella bivia DNF00650 | Bacteria | 0.13 |
| 1110 | 1294025 | Cellulosilyticum ruminicola JCM 14822 | Bacteria | 0.13 |
| 1111 | 37733.7 | Prunus necrotic ringspot virus | Viruses | 0.13 |
| 1112 | 393921.1 | Porphyromonas crevioricanis strain COT-253 | Bacteria | 0.13 |
| 1113 | 1496.602 | Clostridioides difficile isolate VL_0285 | Bacteria | 0.13 |
| 1114 | 1943580.1 | Pyramidobacter sp. C12-8 | Bacteria | 0.13 |
| 1115 | 1410663 | Megasphaera elsdenii T81 | Bacteria | 0.13 |
| 1116 | 1236517 | Prevotella fusca JCM 17724 | Bacteria | 0.13 |
| 1117 | 168135.1 | Apium virus Y | Viruses | 0.13 |
| 1118 | 1739522.1 | Haemophilus sp. HMSC068C11 | Bacteria | 0.13 |
| 1119 | 544580.13 | Actinomyces oris strain P6N | Bacteria | 0.13 |
| 1120 | 1382366 | Lactobacillus plantarum 4_3 | Bacteria | 0.13 |
| 1121 | 1035185 | Streptococcus parasanguinis SK236 | Bacteria | 0.13 |
| 1122 | 404196.1 | Blackberry yellow vein-associated virus | Viruses | 0.13 |
| 1123 | 12172.1 | Shallot latent virus isolate SW3 | Viruses | 0.13 |
| 1124 | 28026.4 | Bifidobacterium pseudocatenulatum strain CA-05 | Bacteria | 0.13 |
| 1125 | 686660 | Veillonella parvula ATCC 17745 | Bacteria | 0.13 |
| 1126 | 1679.13 | Bifidobacterium longum subsp. longum strain VMKB44 | Bacteria | 0.13 |
| 1127 | 1122981 | Prevotella corporis DSM 18810 = JCM 8529 | Bacteria | 0.13 |
| 1128 | 469588.1 | Bacteroides sp. 2_1_22 | Bacteria | 0.13 |
| 1129 | 1235558 | Herbaspirillum huttiense subsp. putei IAM 15032 | Bacteria | 0.13 |
| 1130 | 1401244 | Methanobrevibacter arboriphilus ANOR1 | Archaea | 0.13 |
| 1131 | 1261.2 | Peptostreptococcus anaerobius strain KA00810 | Bacteria | 0.13 |
| 1132 | 1408472 | Prevotella brevis ATCC 19188 | Bacteria | 0.13 |
| 1133 | 1579342.1 | Streptococcus sp. 343_SSPC | Bacteria | 0.13 |
| 1134 | 910312 | Porphyromonas asaccharolytica PR426713P-I | Bacteria | 0.13 |
| 1135 | 180332.2 | Robinsoniella peoriensis isolate 6600698 | Bacteria | 0.13 |
| 1136 | 1630.2 | Kandleria vitulina strain S3b | Bacteria | 0.13 |
| 1137 | 1322347 | Bifidobacterium longum E18 | Bacteria | 0.13 |
| 1138 | 936375.1 | Mogibacterium sp. CM50 | Bacteria | 0.12 |
| 1139 | 999422 | Prevotella maculosa OT 289 | Bacteria | 0.12 |
| 1140 | 12056.1 | Tobacco necrosis virus D | Viruses | 0.12 |
| 1141 | 193121.1 | Pea enation mosaic virus-1 | Viruses | 0.12 |
| 1142 | 1423746 | Lactobacillus frumenti DSM 13145 | Bacteria | 0.12 |
| 1143 | 154288.1 | Turicibacter sanguinis strain 2789STDY5834851 | Bacteria | 0.12 |
| 1144 | 1449896.1 | Uncultured phage WW-nAnB strain 2 strain 2 | Viruses | 0.12 |
| 1145 | 1236521 | Porphyromonas macacae JCM 15984 | Bacteria | 0.12 |
| 1146 | 59201.8 | Salmonella enterica subsp. enterica strain ADRDL-LA-5-2014 | Bacteria | 0.12 |

| | | | | |
|---|---|---|---|---|
| 1147 | 1286820 | [Clostridium] methoxybenzovorans SR3 | Bacteria | 0.12 |
| 1148 | 1402207 | Lactobacillus ruminis S23 | Bacteria | 0.12 |
| 1149 | 1121950 | Hespellia stercorisuis DSM 15480 | Bacteria | 0.12 |
| 1150 | 445974 | Erysipelatoclostridium ramosum DSM 1402 | Bacteria | 0.12 |
| 1151 | 727.74 | Haemophilus influenzae strain 841_HINF | Bacteria | 0.12 |
| 1152 | 1495144.1 | methanogenic archaeon ISO4-H5 | Archaea | 0.12 |
| 1153 | 12319.3 | Apple mosaic virus isolate Apple | Viruses | 0.12 |
| 1154 | 31504.1 | Tobacco ringspot virus satellite RNA | Viruses | 0.12 |
| 1155 | 502393.1 | Gemella asaccharolytica strain KA00071 | Bacteria | 0.12 |
| 1156 | 1155766 | Enterococcus faecium Aus0004 | Bacteria | 0.12 |
| 1157 | 11987.1 | Melon necrotic spot virus strain MNSV/USA/2016 | Viruses | 0.11 |
| 1158 | 1105171.1 | Bacteroides phage B124-14 | Viruses | 0.11 |
| 1159 | 1341156 | Ruminococcus albus SY3 | Bacteria | 0.11 |
| 1160 | 1347790 | Prevotella intermedia ZT | Bacteria | 0.11 |
| 1161 | 1639.215 | Listeria monocytogenes strain FDA00009837 | Bacteria | 0.11 |
| 1162 | 1852379.1 | Veillonellaceae bacterium Marseille-P2911 | Bacteria | 0.11 |
| 1163 | 521393 | Actinomyces timonensis DSM 23838 | Bacteria | 0.11 |
| 1164 | 1401078 | Prevotella buccalis DNF00985 | Bacteria | 0.11 |
| 1165 | 216816.14 | Bifidobacterium longum isolate Bifido_04 | Bacteria | 0.11 |
| 1166 | 562981 | Gemella haemolysans M341 | Bacteria | 0.11 |
| 1167 | 1232449.1 | Clostridiales bacterium VE202-08 | Bacteria | 0.11 |
| 1168 | 563008 | Prevotella oris C735 | Bacteria | 0.11 |
| 1169 | 1768874.1 | Sinapis alba cryptic virus 1 isolate LTBJ | Viruses | 0.11 |
| 1170 | 47770.6 | Lactobacillus crispatus strain VMC7 | Bacteria | 0.11 |
| 1171 | 1128111 | Veillonella atypica KON | Bacteria | 0.11 |
| 1172 | 862965 | Haemophilus parainfluenzae T3T1 | Bacteria | 0.11 |
| 1173 | 1391465 | Enterococcus faecium 10/96A | Bacteria | 0.11 |
| 1174 | 1122984 | Prevotella intermedia ATCC 25611 = DSM 20706 | Bacteria | 0.11 |
| 1175 | 305.9 | Ralstonia solanacearum strain 58_RSOL | Bacteria | 0.11 |
| 1176 | 1319815 | Cetobacterium somerae ATCC BAA-474 | Bacteria | 0.11 |
| 1177 | 1712675.1 | Turicibacter sp. H121 sp. H121 | Bacteria | 0.11 |
| 1178 | 469605 | Fusobacterium gonidiaformans 3-1-5R | Bacteria | 0.11 |
| 1179 | 180332.1 | Robinsoniella peoriensis strain WT | Bacteria | 0.11 |
| 1180 | 1190620.1 | Atopobium sp. ICM42b | Bacteria | 0.11 |
| 1181 | 397290.1 | Lachnospiraceae bacterium A2 | Bacteria | 0.11 |
| 1182 | 1423782 | Lactobacillus panis DSM 6035 | Bacteria | 0.10 |
| 1183 | 887929 | Pseudoramibacter alactolyticus ATCC 23263 | Bacteria | 0.10 |
| 1184 | 1280686.1 | Butyrivibrio sp. MC2013 | Bacteria | 0.10 |
| 1185 | 585506 | Weissella paramesenteroides ATCC 33313 | Bacteria | 0.10 |
| 1186 | 1423769 | Lactobacillus manihotivorans DSM 13343 = JCM 12514 | Bacteria | 0.10 |
| 1187 | 839.3 | Prevotella ruminicola strain D31d | Bacteria | 0.10 |
| 1188 | 367121.1 | Grapevine leafroll-associated virus 10 | Viruses | 0.10 |
| 1189 | 1685.22 | Bifidobacterium breve strain LMC520 | Bacteria | 0.10 |
| 1190 | 167634.1 | Grapevine rootstock stem lesion associated virus | Viruses | 0.10 |
| 1191 | 1122978 | Prevotella albensis DSM 11370 = JCM 12258 | Bacteria | 0.10 |
| 1192 | 626369 | Granulicatella elegans ATCC 700633 | Bacteria | 0.10 |
| 1193 | 1595998.12 | Human smacovirus 1 isolate France/3/2009/4191 | Viruses | 0.10 |

| | | | | |
|---|---|---|---|---|
| 1194 | 1122991 | Prevotella shahii DSM 15611 = JCM 12083 | Bacteria | 0.10 |
| 1195 | 12175.4 | Apple chlorotic leaf spot virus strain AC-ind | Viruses | 0.10 |
| 1196 | 873127 | Enterococcus faecium E4453 | Bacteria | 0.10 |
| 1197 | 1054217.1 | Thermoplasmatales archaeon BRNA1 | Archaea | 0.10 |
| 1198 | 1127131 | Weissella confusa LBAE C39-2 | Bacteria | 0.10 |
| 1199 | 548480 | Bifidobacterium longum subsp. longum ATCC 55813 | Bacteria | 0.10 |
| 1200 | 318464.1 | Clostridium sulfidigenes strain 113A | Bacteria | 0.10 |
| 1201 | 649743 | Actinomyces sp. oral taxon 848 str. F0332 | Bacteria | 0.10 |
| 1202 | 1384065 | Ruminococcus albus AD2013 | Bacteria | 0.10 |
| 1203 | 33945.1 | Enterococcus avium strain 639_EFCM | Bacteria | 0.10 |
| 1204 | 39681.1 | Asparagus virus 2 | Viruses | 0.10 |
| 1205 | 936591.1 | Veillonella sp. ICM51a | Bacteria | 0.10 |
| 1206 | 1218148 | Salmonella enterica subsp. enterica serovar Typhimurium str. STm8 | Bacteria | 0.10 |
| 1207 | 119219.2 | Cupriavidus metallidurans strain NA4 | Bacteria | 0.10 |
| 1208 | 1423801 | Lactobacillus satsumensis DSM 16230 = JCM 12392 | Bacteria | 0.10 |
| 1209 | 1496.628 | Clostridioides difficile isolate VL_0086 | Bacteria | 0.09 |
| 1210 | 35350.1 | Apple stem pitting virus isolate PM8 | Viruses | 0.09 |
| 1211 | 91753.2 | Cucurbit aphid-borne yellows virus | Viruses | 0.09 |
| 1212 | 1581114.1 | Enterococcus sp. HMSC05C03 | Bacteria | 0.09 |
| 1213 | 99179.1 | Bacteroides phage B40-8 | Viruses | 0.09 |
| 1214 | 35350.2 | Apple stem pitting virus isolate YT | Viruses | 0.09 |
| 1215 | 1702221.2 | Faecalibaculum rodentium strain Alo17 | Bacteria | 0.09 |
| 1216 | 983966 | Cyberlindnera jadinii NRRL Y-1542 | Eukaryota | 0.09 |
| 1217 | 35350.1 | Apple stem pitting virus isolate Hannover | Viruses | 0.09 |
| 1218 | 288000.3 | Bradyrhizobium sp. BTAi1 sp. BTAi1 | Bacteria | 0.09 |
| 1219 | 1321772 | Aggregatibacter sp. oral taxon 458 str. W10330 | Bacteria | 0.09 |
| 1220 | 203168.1 | Grapevine leafroll-associated virus 6 | Viruses | 0.09 |
| 1221 | 90410.1 | Streptococcus phage DT1 | Viruses | 0.09 |
| 1222 | 1318.7 | Streptococcus parasanguinis strain 392_SPAR | Bacteria | 0.09 |
| 1223 | 82135.2 | Atopobium vaginae strain CMW7778A | Bacteria | 0.09 |
| 1224 | 1561964.1 | Methanosphaera sp. WGK6 | Archaea | 0.09 |
| 1225 | 28026.2 | C29 | Bacteria | 0.09 |
| 1226 | 479713 | Primula malacoides virus China/Mar2007 | Viruses | 0.09 |
| 1227 | 29272.3 | Turnip vein-clearing virus | Viruses | 0.09 |
| 1228 | 1588751.1 | Tissierellia bacterium KA00581 | Bacteria | 0.09 |
| 1229 | 1282664 | Streptococcus oralis subsp. tigurinus AZ_3a | Bacteria | 0.09 |
| 1230 | 927694 | Leuconostoc inhae KCTC 3774 | Bacteria | 0.09 |
| 1231 | 348151.1 | Lactobacillus siliginis strain DSM | Bacteria | 0.09 |
| 1232 | 1165892 | Leuconostoc gelidum subsp. gasicomitatum KG16-1 | Bacteria | 0.09 |
| 1233 | 585530 | Brevibacterium mcbrellneri ATCC 49030 | Bacteria | 0.09 |
| 1234 | 1121126 | Brochothrix thermosphacta DSM 20171 = FSL F6-1036 | Bacteria | 0.09 |
| 1235 | 1430896 | Haemophilus parahaemolyticus G321 | Bacteria | 0.09 |
| 1236 | 1121132 | Butyrivibrio hungatei DSM 14810 | Bacteria | 0.08 |

| | | | | |
|---|---|---|---|---|
| 1237 | 11612 | Impatiens necrotic spot virus | Viruses | 0.08 |
| 1238 | 1235835.1 | Anaerotruncus sp. G3(2012) | Bacteria | 0.08 |
| 1239 | 1661.1 | Trueperella pyogenes strain 331_TPYO | Bacteria | 0.08 |
| 1240 | 134358.2 | Westerdykella cylindrica strain ATCC | Eukaryota | 0.08 |
| 1241 | 1267.2 | Clostridium ventriculi strain 2789STDY5834858 | Bacteria | 0.08 |
| 1242 | 1605.2 | Lactobacillus animalis strain 381-IL-28 | Bacteria | 0.08 |
| 1243 | 196400.1 | Grapevine rupestris stem pitting-associated virus isolate SK704-B | Viruses | 0.08 |
| 1244 | 470.537 | Acinetobacter baumannii strain XH803 | Bacteria | 0.08 |
| 1245 | 907.1 | Megasphaera elsdenii strain 24-50 | Bacteria | 0.08 |
| 1246 | 28347.4 | Apple stem grooving virus isolate M219-3 | Viruses | 0.08 |
| 1247 | 180957.7 | Pectobacterium carotovorum subsp. brasiliense strain BD255 | Bacteria | 0.08 |
| 1248 | 190721.2 | Ralstonia insidiosa strain ATCC | Bacteria | 0.08 |
| 1249 | 1176736.1 | Pitaya virus X isolate P37 | Viruses | 0.08 |
| 1250 | 1755753.1 | Penicillium aurantiogriseum foetidus-like virus | Viruses | 0.08 |
| 1251 | 1673726.1 | Clostridiales bacterium SIT11 | Bacteria | 0.08 |
| 1252 | 1111133.1 | Peptoniphilus sp. BV3AC2 | Bacteria | 0.08 |
| 1253 | 1123303 | Streptococcus ferus DSM 20646 | Bacteria | 0.08 |
| 1254 | 28901.27 | Salmonella enterica strain NGUA09 | Bacteria | 0.08 |
| 1255 | 1352.151 | Enterococcus faecium isolate Hp_21-21 | Bacteria | 0.08 |
| 1256 | 1226531 | Saccharomyces cerevisiae virus L-A-lus | Viruses | 0.08 |
| 1257 | 1235802 | Eubacterium plexicaudatum ASF492 | Bacteria | 0.08 |
| 1258 | 1324352.1 | Chryseobacterium gallinarum strain DSM | Bacteria | 0.08 |
| 1259 | 485724.1 | Melon severe mosaic tospovirus isolate VE440-A | Viruses | 0.08 |
| 1260 | 1203556.1 | Actinomyces sp. HPA0247 | Bacteria | 0.08 |
| 1261 | 1357399 | Helicobacter canis NCTC 12740 | Bacteria | 0.08 |
| 1262 | 1840217.1 | Candidatus Arthromitus sp. SFB-turkey isolate UMNCA01 | Bacteria | 0.08 |
| 1263 | 796944 | Oribacterium asaccharolyticum ACB7 | Bacteria | 0.08 |
| 1264 | 12615.8 | Cherry leaf roll virus isolate Olm1 | Viruses | 0.08 |
| 1265 | 67761.3 | Cowpea mild mottle virus isolate CPMMV:BR:GO:01:1 | Viruses | 0.08 |
| 1266 | 1401068 | Prevotella bivia DNF00320 | Bacteria | 0.08 |
| 1267 | 1660.1 | Actinomyces odontolyticus strain XH001 | Bacteria | 0.07 |
| 1268 | 35350.8 | Apple stem pitting virus isolate HB-HN1 | Viruses | 0.07 |
| 1269 | 64003.5 | 93/955 | Viruses | 0.07 |
| 1270 | 1118059 | Kallipyga massiliensis ph2 | Bacteria | 0.07 |
| 1271 | 936574.1 | Shuttleworthia sp. MSX8B | Bacteria | 0.07 |
| 1272 | 37733.5 | Prunus necrotic ringspot virus isolate ChrYL | Viruses | 0.07 |
| 1273 | 1385512 | Pontibacillus litoralis JSM 072002 | Bacteria | 0.07 |
| 1274 | 12143.1 | Cucumber necrosis virus | Viruses | 0.07 |
| 1275 | 1407052 | Lactobacillus fermentum L930BB | Bacteria | 0.07 |
| 1276 | 202566.1 | Cherry rasp leaf virus | Viruses | 0.07 |
| 1277 | 1121111 | Bifidobacterium thermacidophilum subsp. thermacidophilum DSM 15837 | Bacteria | 0.07 |
| 1278 | 328061.2 | Radish mosaic virus | Viruses | 0.07 |
| 1279 | 1335.1 | Streptococcus equinus strain ICDDRB-NRC-S6 | Bacteria | 0.07 |

| | | | | |
|---|---|---|---|---|
| 1280 | 37734.4 | Enterococcus casseliflavus strain PAVET15 | Bacteria | 0.07 |
| 1281 | 28347.8 | Apple stem grooving virus isolate HPKu-2 | Viruses | 0.07 |
| 1282 | 1720298.1 | Peptoniphilus phoceensis strain SIT15 | Bacteria | 0.07 |
| 1283 | 1339293 | Bacteroides fragilis str. 1007-1-F #8 | Bacteria | 0.07 |
| 1284 | 1280670.1 | Butyrivibrio sp. AD3002 | Bacteria | 0.07 |
| 1285 | 1679.3 | Bifidobacterium longum subsp. longum strain MC-42 | Bacteria | 0.07 |
| 1286 | 888057 | Aggregatibacter segnis ATCC 33393 | Bacteria | 0.07 |
| 1287 | 1402208 | Lactobacillus ruminis DPC 6832 | Bacteria | 0.07 |
| 1288 | 888062 | Dialister micraerophilus DSM 19965 | Bacteria | 0.07 |
| 1289 | 1435051 | Bifidobacterium moukalabense DSM 27321 | Bacteria | 0.07 |
| 1290 | 11008.1 | Saccharomyces cerevisiae virus L-A | Viruses | 0.07 |
| 1291 | 997898 | Clostridium butyricum 60E.3 | Bacteria | 0.07 |
| 1292 | 467210.1 | Lachnoanaerobaculum saburreum strain DNF00896 | Bacteria | 0.07 |
| 1293 | 111105.1 | Porphyromonas gulae strain COT-052 | Bacteria | 0.07 |
| 1294 | 679192 | Bulleidia extructa W1219 | Bacteria | 0.07 |
| 1295 | 767453 | Lactobacillus fermentum F-6 | Bacteria | 0.07 |
| 1296 | 294747 | Candida tropicalis MYA-3404 | Eukaryota | 0.07 |
| 1297 | 326941.1 | Raspberry leaf mottle virus isolate HCRL | Viruses | 0.07 |
| 1298 | 697329 | Ruminococcus albus 7 = DSM 20455 | Bacteria | 0.07 |
| 1299 | 33029.1 | Anaerococcus hydrogenalis strain MJR7738A | Bacteria | 0.07 |
| 1300 | 469615 | Fusobacterium gonidiaformans ATCC 25563 | Bacteria | 0.07 |
| 1301 | 936549.1 | Actinomyces sp. ICM54 | Bacteria | 0.07 |
| 1302 | 879308 | Peptoniphilus sp. oral taxon 375 str. F0436 | Bacteria | 0.07 |
| 1303 | 54067.1 | Xylophilus ampelinus strain CCH5-B3 | Bacteria | 0.07 |
| 1304 | 1230730.1 | Tissierellia bacterium S5-A11 | Bacteria | 0.07 |
| 1305 | 12433.2 | Garlic virus A isolate WA7 | Viruses | 0.07 |
| 1306 | 1697797.1 | Actinobacteria bacterium UC5.1-1B11 | Bacteria | 0.07 |
| 1307 | 1695.2 | Bifidobacterium longum subsp. suis strain BSM11-5 | Bacteria | 0.07 |
| 1308 | 1348632 | Lactococcus plantarum NBRC 100936 | Bacteria | 0.07 |
| 1309 | 253700.1 | Schlumbergera virus X | Viruses | 0.07 |
| 1310 | 1347393.1 | Bacteroides neonati strain MS4 | Bacteria | 0.07 |
| 1311 | 663278 | Ethanoligenens harbinense YUAN-3 | Bacteria | 0.07 |
| 1312 | 1437611 | Bifidobacterium saeculare DSM 6531 = LMG 14934 | Bacteria | 0.07 |
| 1313 | 1211814 | Bacillus massilioanorexius AP8 | Bacteria | 0.07 |
| 1314 | 1496.534 | Clostridioides difficile isolate VL_0083 | Bacteria | 0.06 |
| 1315 | 1739371.1 | Streptococcus sp. HMSC064H09 | Bacteria | 0.06 |
| 1316 | 33763.1 | Peanut mottle virus | Viruses | 0.06 |
| 1317 | 1161744 | Bifidobacterium longum subsp. longum 1-6B | Bacteria | 0.06 |
| 1318 | 936589.1 | Veillonella sp. AS16 | Bacteria | 0.06 |
| 1319 | 904306 | Streptococcus vestibularis F0396 | Bacteria | 0.06 |
| 1320 | 537971 | Helicobacter cinaedi CCUG 18818 = ATCC BAA-847 | Bacteria | 0.06 |
| 1321 | 1080349.1 | Saccharomyces eubayanus strain FM1318 | Eukaryota | 0.06 |
| 1322 | 1265.3 | Ruminococcus flavefaciens strain YRD2003 | Bacteria | 0.06 |

| | | | | |
|---|---|---|---|---|
| 1323 | 985691 | Sapovirus Hu/GI.2/BR-DF01/BRA/2009 | Viruses | 0.06 |
| 1324 | 12430.3 | Garlic virus D isolate GarVD-SW10 | Viruses | 0.06 |
| 1325 | 1215061 | Clostridioides difficile E10 | Bacteria | 0.06 |
| 1326 | 1350473 | Bifidobacterium longum subsp. longum 17-1B | Bacteria | 0.06 |
| 1327 | 1299998.1 | Olsenella scatoligenes strain SK9K4 | Bacteria | 0.06 |
| 1328 | 1118058.1 | Actinomyces sp. ph3 | Bacteria | 0.06 |
| 1329 | 1046402.2 | Potato virus H isolate YN | Viruses | 0.06 |
| 1330 | 1127690 | Actinomyces sp. oral taxon 181 str. F0379 | Bacteria | 0.06 |
| 1331 | 696281 | Desulfotomaculum ruminis DSM 2154 | Bacteria | 0.06 |
| 1332 | 866774 | Atopobium vaginae PB189-T1-4 | Bacteria | 0.06 |
| 1333 | 866776 | Veillonella atypica ACS-049-V-Sch6 | Bacteria | 0.06 |
| 1334 | 1122151 | Lactobacillus paralimentarius DSM 13238 = JCM 10415 | Bacteria | 0.06 |
| 1335 | 1410616.1 | Pseudobutyrivibrio sp. MD2005 | Bacteria | 0.06 |
| 1336 | 37734.2 | Enterococcus casseliflavus strain NLAE-zl-G268 | Bacteria | 0.06 |
| 1337 | 1679.12 | W11 | Bacteria | 0.06 |
| 1338 | 36874.1 | Porphyromonas cangingivalis strain COT-109 | Bacteria | 0.06 |
| 1339 | 1852378.1 | Veillonellaceae bacterium Marseille-P2974 | Bacteria | 0.06 |
| 1340 | 679200 | Johnsonella ignava ATCC 51276 | Bacteria | 0.06 |
| 1341 | 767031 | Prevotella denticola F0289 | Bacteria | 0.06 |
| 1342 | 1914544 | Norovirus Hu/USA/2014/GI.P7_GI.7/GA5043 | Viruses | 0.06 |
| 1343 | 284592 | Debaryomyces hansenii CBS767 | Eukaryota | 0.06 |
| 1344 | 1930605 | Saccharomyces cerevisiae virus L-BC-2 | Viruses | 0.06 |
| 1345 | 1774276.1 | Hordeum vulgare endornavirus | Viruses | 0.06 |
| 1346 | 1235798.1 | Dorea sp. 5-2 | Bacteria | 0.06 |
| 1347 | 537937 | Bifidobacterium longum subsp. infantis CCUG 52486 | Bacteria | 0.06 |
| 1348 | 1235790.1 | Eubacterium sp. 14-2 | Bacteria | 0.06 |
| 1349 | 31713.1 | Lettuce infectious yellows virus | Viruses | 0.06 |
| 1350 | 208084.1 | Grapevine Algerian latent virus | Viruses | 0.06 |
| 1351 | 253700.3 | Schlumbergera virus X isolate Palma-PE | Viruses | 0.06 |
| 1352 | 253700.2 | Schlumbergera virus X isolate nopal | Viruses | 0.06 |
| 1353 | 1175296 | Methanomassiliicoccus luminyensis B10 | Archaea | 0.06 |
| 1354 | 1401079 | Mogibacterium timidum ATCC 33093 | Bacteria | 0.06 |
| 1355 | 1203602 | Atopobium sp. oral taxon 199 str. F0494 | Bacteria | 0.06 |
| 1356 | 33036.1 | Anaerococcus tetradius strain MJR8151 | Bacteria | 0.06 |
| 1357 | 1423799 | Lactobacillus salivarius DSM 20555 = ATCC 11741 | Bacteria | 0.06 |
| 1358 | 1358418.1 | Sinorhizobium sp. GL28 | Bacteria | 0.06 |
| 1359 | 1401066 | Prevotella amnii DNF00058 | Bacteria | 0.06 |
| 1360 | 1739537.1 | Anaerococcus sp. HMSC075B03 | Bacteria | 0.06 |
| 1361 | 1423764 | Lactobacillus kefiri DSM 20587 = JCM 5818 | Bacteria | 0.06 |
| 1362 | 196400.1 | Grapevine rupestris stem pitting-associated virus strain Syrah | Viruses | 0.06 |
| 1363 | 11987.4 | Melon necrotic spot virus isolate N | Viruses | 0.06 |
| 1364 | 35350.9 | Apple stem pitting virus isolate PB66 | Viruses | 0.06 |
| 1365 | 1423815 | Lactobacillus versmoldensis DSM 14857 = KCTC 3814 | Bacteria | 0.06 |

| | | | | |
|---|---|---|---|---|
| 1366 | 1423783 | Lactobacillus pantheris DSM 15945 = JCM 12539 = NBRC 106106 | Bacteria | 0.06 |
| 1367 | 1302.12 | Streptococcus gordonii strain MB666 | Bacteria | 0.06 |
| 1368 | 888826 | Campylobacter upsaliensis JV21 | Bacteria | 0.06 |
| 1369 | 1579341.1 | Streptococcus sp. 400_SSPC | Bacteria | 0.06 |
| 1370 | 348449.1 | Raphanus sativus cryptic virus 1 | Viruses | 0.06 |
| 1371 | 1161412.1 | Prevotella sp. ICM33 | Bacteria | 0.06 |
| 1372 | 1613.2 | Lactobacillus fermentum | Bacteria | 0.06 |
| 1373 | 1000568.1 | Megasphaera sp. UPII 199-6 | Bacteria | 0.06 |
| 1374 | 12430.2 | Garlic virus D isolate GarVD-SW9 | Viruses | 0.06 |
| 1375 | 1095733 | Streptococcus parasanguinis F0449 | Bacteria | 0.06 |
| 1376 | 379892.1 | Passiflora latent carlavirus | Viruses | 0.06 |
| 1377 | 1442158 | Saccharomyces cerevisiae virus L-A-2 | Viruses | 0.06 |
| 1378 | 1123250 | Selenomonas bovis DSM 23594 | Bacteria | 0.06 |
| 1379 | 1261640.1 | Eubacterium sp. 68-3-10 | Bacteria | 0.06 |
| 1380 | 42004.6 | Leek yellow stripe virus isolate LYSV-MG | Viruses | 0.06 |
| 1381 | 42882.2 | Cherry virus A isolate ChYT52 | Viruses | 0.06 |
| 1382 | 158787.2 | Bifidobacterium scardovii strain LMG | Bacteria | 0.06 |
| 1383 | 60456 | Iris yellow spot virus | Viruses | 0.06 |
| 1384 | 553175 | Porphyromonas endodontalis ATCC 35406 | Bacteria | 0.06 |
| 1385 | 1123754.1 | nopal | Viruses | 0.06 |
| 1386 | 1297865.1 | Bradyrhizobium sp. OHSU_III | Bacteria | 0.06 |
| 1387 | 2702.11 | Gardnerella vaginalis strain CMW7778B | Bacteria | 0.06 |
| 1388 | 562.547 | Escherichia coli strain LM17584/1 | Bacteria | 0.06 |
| 1389 | 767100 | Parvimonas sp. oral taxon 110 str. F0139 | Bacteria | 0.06 |
| 1390 | 1151384 | Clostridioides difficile Y266 | Bacteria | 0.06 |
| 1391 | 1686287.1 | Kallipyga gabonensis strain GM4 | Bacteria | 0.06 |
| 1392 | 1276.1 | Kytococcus sedentarius strain 262_KSED | Bacteria | 0.06 |
| 1393 | 1080072.1 | Streptococcus dentasini | Bacteria | 0.06 |
| 1394 | 105219.3 | Ralstonia mannitolilytica strain SN83A39 | Bacteria | 0.06 |
| 1395 | 31537.1 | Lactococcus phage c2 | Viruses | 0.06 |
| 1396 | 1852373.1 | Murdochiella sp. Marseille-P2341 strain Marseille-P2341T | Bacteria | 0.06 |
| 1397 | 1469950.1 | Robinsoniella sp. KNHs210 | Bacteria | 0.06 |
| 1398 | 72000.1 | Kocuria rhizophila strain 14ASP | Bacteria | 0.06 |
| 1399 | 1525173.1 | Human circovirus VS6600022 | Viruses | 0.06 |
| 1400 | 134605.1 | Fusobacterium equinum strain CMW8396 | Bacteria | 0.06 |
| 1401 | 234601 | Sapovirus Mc10 | Viruses | 0.06 |
| 1402 | 626523 | Shuttleworthia satelles DSM 14600 | Bacteria | 0.06 |
| 1403 | 936550.1 | Atopobium sp. BS2 | Bacteria | 0.06 |
| 1404 | 1348.8 | Streptococcus parauberis strain SP-llh | Bacteria | 0.06 |
| 1405 | 425279.2 | Rehmannia mosaic virus | Viruses | 0.06 |
| 1406 | 1777865.1 | Weissella sp. DD23 | Bacteria | 0.06 |
| 1407 | 196375.8 | Beet black scorch virus isolate CO | Viruses | 0.06 |
| 1408 | 39777.1 | Veillonella atypica strain CMW7756B | Bacteria | 0.06 |
| 1409 | 655811 | Anaerococcus vaginalis ATCC 51170 | Bacteria | 0.06 |
| 1410 | 1715211.1 | Haemophilus sp. HMSC061E01 | Bacteria | 0.06 |
| 1411 | 45071.2 | Legionella parisiensis strain DSM | Bacteria | 0.06 |

| | | | | |
|---|---|---|---|---|
| 1412 | 1235800.1 | Lachnospiraceae bacterium 10-1 | Bacteria | 0.06 |
| 1413 | 862517 | Peptoniphilus duerdenii ATCC BAA-1640 | Bacteria | 0.06 |
| 1414 | 1711684.1 | Hot pepper endornavirus isolate CS | Viruses | 0.06 |
| 1415 | 1497954.1 | Bacteroidales bacterium KA00344 | Bacteria | 0.06 |
| 1416 | 1280700 | Butyrivibrio fibrisolvens FE2007 | Bacteria | 0.06 |
| 1417 | 869209 | Treponema succinifaciens DSM 2489 | Bacteria | 0.05 |
| 1418 | 729.7 | Haemophilus parainfluenzae strain 901_HPAR | Bacteria | 0.05 |
| 1419 | 328396.1 | Enterococcus aquimarinus strain DSM | Bacteria | 0.05 |
| 1420 | 1232454.1 | Clostridiales bacterium VE202-26 | Bacteria | 0.05 |
| 1421 | 1425364.1 | Carrot torradovirus 1 isolate CTV-1_RNA2_H6 | Viruses | 0.05 |
| 1422 | 396268.1 | Lactobacillus secaliphilus strain DSM | Bacteria | 0.05 |
| 1423 | 1074482 | Bifidobacterium breve DPC 6330 | Bacteria | 0.05 |
| 1424 | 1190621 | Olsenella uli MSTE5 | Bacteria | 0.05 |
| 1425 | 1121100 | Bacteroides pyogenes DSM 20611 = JCM 6294 | Bacteria | 0.05 |
| 1426 | 1095727.1 | Streptococcus sp. SK643 | Bacteria | 0.05 |
| 1427 | 888832 | Prevotella salivae DSM 15606 | Bacteria | 0.05 |
| 1428 | 595895.1 | Drosophila A virus | Viruses | 0.05 |
| 1429 | 1265.5 | Ruminococcus flavefaciens strain XPD3002 | Bacteria | 0.05 |
| 1430 | 1265.1 | Ruminococcus flavefaciens strain YL228 | Bacteria | 0.05 |
| 1431 | 1405296 | Chlamydia suis MD56 | Bacteria | 0.05 |
| 1432 | 1654927.1 | Klebsiella phage PKP126 | Viruses | 0.05 |
| 1433 | 1074494 | Streptococcus salivarius M18 | Bacteria | 0.05 |
| 1434 | 12432.1 | Garlic virus B isolate Mesi | Viruses | 0.05 |
| 1435 | 1739422.1 | Streptococcus sp. HMSC065C01 | Bacteria | 0.05 |
| 1436 | 1520332.1 | Blueberry mosaic associated virus | Viruses | 0.05 |
| 1437 | 907.3 | Megasphaera elsdenii strain YE34 | Bacteria | 0.05 |
| 1438 | 29363.1 | Clostridium paraputrificum strain 373-A1 | Bacteria | 0.05 |
| 1439 | 1236509 | Prevotella dentasini JCM 15908 | Bacteria | 0.05 |
| 1440 | 866778 | Veillonella atypica ACS-134-V-Col7a | Bacteria | 0.05 |
| 1441 | 216816.16 | Bifidobacterium longum | Bacteria | 0.05 |
| 1442 | 1739286.1 | Anaerococcus sp. HMSC068A02 | Bacteria | 0.05 |
| 1443 | 1167628 | Actinomyces massiliensis 4401292 | Bacteria | 0.05 |
| 1444 | 311413.1 | Lettuce big-vein associated virus | Viruses | 0.05 |
| 1445 | 42882.4 | Cherry virus A isolate JK | Viruses | 0.05 |
| 1446 | 729.4 | Haemophilus parainfluenzae strain 209_HPAR | Bacteria | 0.05 |
| 1447 | 739.1 | Aggregatibacter segnis strain 933_AAPH | Bacteria | 0.05 |
| 1448 | 1122147 | Lactobacillus harbinensis DSM 16991 | Bacteria | 0.05 |
| 1449 | 1284686 | Anaerococcus lactolyticus S7-1-13 | Bacteria | 0.05 |
| 1450 | 1658779.2 | Porphyromonadaceae bacterium H1 | Bacteria | 0.05 |
| 1451 | 12280.8 | Tomato ringspot virus isolate Rasp1-2014 | Viruses | 0.05 |
| 1452 | 936381.1 | Selenomonas sp. CM52 | Bacteria | 0.05 |
| 1453 | 1331064 | Escherichia coli CE516 | Bacteria | 0.05 |
| 1454 | 1123754.2 | Rattail cactus necrosis-associated virus | Viruses | 0.05 |
| 1455 | 284593 | Candida glabrata CBS 138 | Eukaryota | 0.05 |
| 1456 | 284590 | Kluyveromyces lactis NRRL Y-1140 | Eukaryota | 0.05 |
| 1457 | 682370.1 | Streptococcus phage Alq132 | Viruses | 0.05 |
| 1458 | 729.16 | Haemophilus parainfluenzae strain 65114 | Bacteria | 0.05 |
| 1459 | 1267000 | Mycoplasma hominis ATCC 27545 | Bacteria | 0.05 |

| | | | | |
|---|---|---|---|---|
| 1460 | 768724 | Peptoniphilus sp. oral taxon 836 str. F0141 | Bacteria | 0.05 |
| 1461 | 84135.1 | Gemella sanguinis strain 1094_BTHU | Bacteria | 0.05 |
| 1462 | 553207 | Corynebacterium matruchotii ATCC 14266 | Bacteria | 0.05 |
| 1463 | 1323524.1 | Red clover cryptic virus 2 isolate IPP_Nemaro | Viruses | 0.05 |
| 1464 | 1719140.1 | Klebsiella phage vB_KpnM_KB57 | Viruses | 0.05 |
| 1465 | 173976.1 | Cycas necrotic stunt virus | Viruses | 0.05 |
| 1466 | 1122979 | Prevotella amnii DSM 23384 = JCM 14753 | Bacteria | 0.05 |
| 1467 | 1351755 | Clostridium chauvoei JF4335 | Bacteria | 0.05 |
| 1468 | 1715086.1 | Streptococcus sp. HMSC078H03 | Bacteria | 0.05 |
| 1469 | 290314 | Sapovirus C12 | Viruses | 0.05 |
| 1470 | 1121321 | Asaccharospora irregularis DSM 2635 | Bacteria | 0.05 |
| 1471 | 876478.1 | Tepidiphilus thermophilus strain JCM | Bacteria | 0.05 |
| 1472 | 471858.1 | Helicobacter magdeburgensis strain MIT | Bacteria | 0.05 |
| 1473 | 397287.1 | Lachnospiraceae bacterium 28-4 | Bacteria | 0.05 |
| 1474 | 35281.1 | Paprika mild mottle virus strain israeli | Viruses | 0.05 |
| 1475 | 767029 | Pseudopropionibacterium propionicum F0230a | Bacteria | 0.05 |
| 1476 | 1840518.1 | Gardnerella sp. 30-4 | Bacteria | 0.05 |
| 1477 | 87541.1 | Aerococcus christensenii strain KA00635 | Bacteria | 0.05 |
| 1478 | 1051074 | Streptococcus thermophilus JIM 8232 | Bacteria | 0.05 |
| 1479 | 1035817 | Bifidobacterium longum subsp. longum KACC 91563 | Bacteria | 0.05 |
| 1480 | 1675609.1 | Klebsiella phage Sushi | Viruses | 0.05 |
| 1481 | 1496.3 | Clostridioides difficile isolate VL_0125 | Bacteria | 0.05 |
| 1482 | 525256 | Atopobium vaginae DSM 15829 | Bacteria | 0.05 |
| 1483 | 457388.1 | Parabacteroides sp. 2_1_7 | Bacteria | 0.05 |
| 1484 | 664639.1 | Kocuria salsicia | Bacteria | 0.05 |
| 1485 | 1006000 | Kluyvera ascorbata ATCC 33433 | Bacteria | 0.05 |
| 1486 | 883113 | Facklamia languida CCUG 37842 | Bacteria | 0.05 |
| 1487 | 1505.39 | Paeniclostridium sordellii strain R32462 | Bacteria | 0.05 |
| 1488 | 1601.1 | Lactobacillus agilis | Bacteria | 0.05 |
| 1489 | 1287474 | Atopobium parvulum DNF00906 | Bacteria | 0.05 |
| 1490 | 883062 | Bifidobacterium bifidum S17 | Bacteria | 0.05 |
| 1491 | 1384078.1 | Prevotella sp. DNF00663 | Bacteria | 0.05 |
| 1492 | 42478.1 | Saccharomyces cerevisiae virus L-BC (La) | Viruses | 0.05 |
| 1493 | 905.2 | Acidaminococcus fermentans strain WCC6 | Bacteria | 0.05 |
| 1494 | 71030.1 | Chayote mosaic virus | Viruses | 0.05 |
| 1495 | 883112 | Facklamia ignava CCUG 37419 | Bacteria | 0.05 |
| 1496 | 1744.1 | Propionibacterium freudenreichii | Bacteria | 0.04 |
| 1497 | 1131702.1 | Persea americana endornavirus 1 | Viruses | 0.04 |
| 1498 | 33763.2 | Peanut mottle virus isolate Habin | Viruses | 0.04 |
| 1499 | 55951.9 | Grapevine leafroll-associated virus 3 clone 3 | Viruses | 0.04 |
| 1500 | 591001 | Acidaminococcus fermentans DSM 20731 | Bacteria | 0.04 |
| 1501 | 273677.2 | Microbacterium oleivorans strain Wellendorf | Bacteria | 0.04 |
| 1502 | 392504.1 | Turnip ringspot virus isolate B | Viruses | 0.04 |
| 1503 | 1592930.1 | Yam latent virus isolate SG1 | Viruses | 0.04 |
| 1504 | 1403344 | Xylella fastidiosa Mul-MD | Bacteria | 0.04 |
| 1505 | 1423770 | Lactobacillus mindensis DSM 14500 | Bacteria | 0.04 |
| 1506 | 1121102 | Campylobacter ureolyticus DSM 20703 | Bacteria | 0.04 |

| | | | | |
|---|---|---|---|---|
| 1507 | 1674942.1 | Caenibacillus caldisaponilyticus strain B157 | Bacteria | 0.04 |
| 1508 | 137591.1 | Weissella cibaria strain FBL5 | Bacteria | 0.04 |
| 1509 | 1457183 | Bifidobacterium longum subsp. infantis EK3 | Bacteria | 0.04 |
| 1510 | 4909.1 | Pichia kudriavzevii | Eukaryota | 0.04 |
| 1511 | 1120996 | Anaerosporobacter mobilis DSM 15930 | Bacteria | 0.04 |
| 1512 | 944565 | Parvimonas sp. oral taxon 393 str. F0440 | Bacteria | 0.04 |
| 1513 | 1318.15 | Streptococcus parasanguinis strain 512_SPAR | Bacteria | 0.04 |
| 1514 | 1321823 | Porphyromonas gingivalis W4087 | Bacteria | 0.04 |
| 1515 | 360104 | Campylobacter concisus 13826 | Bacteria | 0.04 |
| 1516 | 1323525.1 | White clover cryptic virus 2 isolate IPP_Lirepa | Viruses | 0.04 |
| 1517 | 999411 | Clostridium colicanis 209318 | Bacteria | 0.04 |
| 1518 | 1862960.1 | Lactococcus phage M5938 | Viruses | 0.04 |
| 1519 | 1597976.1 | Enterococcus phage EFDG1 | Viruses | 0.04 |
| 1520 | 61592.1 | Corynebacterium durum strain CCH6-D9 | Bacteria | 0.04 |
| 1521 | 907.2 | Megasphaera elsdenii strain J1 | Bacteria | 0.04 |
| 1522 | 1423750 | Lactobacillus ghanensis DSM 18630 | Bacteria | 0.04 |
| 1523 | 45634.1 | Streptococcus cristatus strain DD08 | Bacteria | 0.04 |
| 1524 | 1739356.1 | Anaerococcus sp. HMSC065G05 | Bacteria | 0.04 |
| 1525 | 638301 | Granulicatella adiacens ATCC 49175 | Bacteria | 0.04 |
| 1526 | 298.2 | Pseudomonas marginalis strain BS2952 | Bacteria | 0.04 |
| 1527 | 1165092.1 | Lachnospiraceae bacterium JC7 | Bacteria | 0.04 |
| 1528 | 97139.1 | Clostridium sp. ASF502 | Bacteria | 0.04 |
| 1529 | 28347.9 | Apple stem grooving virus isolate HH | Viruses | 0.04 |
| 1530 | 1410676 | Succinivibrio dextrinosolvens H5 | Bacteria | 0.04 |
| 1531 | 1579339.1 | Streptococcus sp. 449_SSPC | Bacteria | 0.04 |
| 1532 | 1739413.1 | Alloscardovia sp. HMSC034E08 | Bacteria | 0.04 |
| 1533 | 1433844.1 | Prevotella sp. HJM029 | Bacteria | 0.04 |
| 1534 | 1336241 | Eubacterium xylanophilum ATCC 35991 | Bacteria | 0.04 |
| 1535 | 42004.2 | Leek yellow stripe virus isolate SW8 | Viruses | 0.04 |
| 1536 | 1316412.1 | Streptococcus sp. HSISS3 | Bacteria | 0.04 |
| 1537 | 525254 | Anaerococcus lactolyticus ATCC 51172 | Bacteria | 0.04 |
| 1538 | 888052 | Actinomyces sp. oral taxon 180 str. F0310 | Bacteria | 0.04 |
| 1539 | 1235799.1 | Lachnospiraceae bacterium 3-2 | Bacteria | 0.04 |
| 1540 | 1214168 | Streptococcus suis 88-1861 | Bacteria | 0.04 |
| 1541 | 1805472.1 | Clostridium sp. Marseille-P2434 sp. Marseille-P2434 | Bacteria | 0.04 |
| 1542 | 1860161.1 | Streptococcus sp. CCUG 49591 | Bacteria | 0.04 |
| 1543 | 12275.4 | Tomato black ring virus isolate TBRV-P1 | Viruses | 0.04 |
| 1544 | 1815509.1 | Bacillus phage AR9 | Viruses | 0.04 |
| 1545 | 1445607 | Bacteroides reticulotermitis JCM 10512 | Bacteria | 0.04 |
| 1546 | 1122949 | Peptoniphilus lacrimalis DSM 7455 | Bacteria | 0.04 |
| 1547 | 1158603 | Enterococcus flavescens ATCC 49996 | Bacteria | 0.04 |
| 1548 | 1496.14 | Clostridioides difficile strain CD8-15 | Bacteria | 0.04 |
| 1549 | 1739452.1 | Haemophilus sp. HMSC073C03 | Bacteria | 0.04 |
| 1550 | 1682.4 | Bifidobacterium longum subsp. infantis strain TPY12-1 | Bacteria | 0.04 |
| 1551 | 186189.1 | Xylanimonas cellulosilytica strain 352_XCEL | Bacteria | 0.04 |
| 1552 | 306264 | Campylobacter upsaliensis RM3195 | Bacteria | 0.04 |

| | | | | |
|---|---|---|---|---|
| 1553 | 212365.1 | Bifidobacterium thermacidophilum subsp. porcinum strain LMG | Bacteria | 0.04 |
| 1554 | 197614.4 | Streptococcus pasteurianus strain GED7275A | Bacteria | 0.04 |
| 1555 | 1101373.1 | Tepidimonas fonticaldi strain PL17 | Bacteria | 0.04 |
| 1556 | 1496.439 | Clostridioides difficile strain VRECD0155 | Bacteria | 0.04 |
| 1557 | 1423718 | Lactobacillus agilis DSM 20509 | Bacteria | 0.04 |
| 1558 | 213633.2 | Providence virus | Viruses | 0.04 |
| 1559 | 28026.8 | Bifidobacterium pseudocatenulatum strain CECT | Bacteria | 0.04 |
| 1560 | 997893 | [Clostridium] bolteae 90A5 | Bacteria | 0.04 |
| 1561 | 1122993 | Prevotella veroralis DSM 19559 = JCM 6290 | Bacteria | 0.04 |
| 1562 | 1930298.1 | Chicken stool-associated gemycircularvirus strain RS/BR/2015 | Viruses | 0.04 |
| 1563 | 1679.16 | Bifidobacterium longum subsp. longum strain AH1206 | Bacteria | 0.04 |
| 1564 | 216816.6 | Bifidobacterium longum isolate Bifido_06 | Bacteria | 0.04 |
| 1565 | 12558.1 | Sesbania mosaic virus | Viruses | 0.04 |
| 1566 | 12041.1 | Bean leafroll virus strain Manfredi | Viruses | 0.04 |
| 1567 | 399795 | Comamonas testosteroni KF-1 | Bacteria | 0.04 |
| 1568 | 277.1 | Meiothermus ruber strain TC-1 | Bacteria | 0.04 |
| 1569 | 1122219 | Megasphaera cerevisiae DSM 20462 | Bacteria | 0.04 |
| 1570 | 1282.6 | Staphylococcus epidermidis strain MF1789 | Bacteria | 0.04 |
| 1571 | 1280688 | Pseudobutyrivibrio ruminis CF1b | Bacteria | 0.04 |
| 1572 | 890402 | Bifidobacterium longum subsp. longum BBMN68 | Bacteria | 0.04 |
| 1573 | 11983.2 | Norwalk virus | Viruses | 0.04 |
| 1574 | 1384082.1 | Veillonellaceae bacterium DNF00751 | Bacteria | 0.04 |
| 1575 | 1794912.1 | Anaerosporomusa subterranea strain RU4 | Bacteria | 0.04 |
| 1576 | 1454376 | Bifidobacterium pseudocatenulatum IPLA36007 | Bacteria | 0.04 |
| 1577 | 1392486.1 | Prevotella sp. HUN102 | Bacteria | 0.04 |
| 1578 | 546271 | Selenomonas sputigena ATCC 35185 | Bacteria | 0.04 |
| 1579 | 306902 | Clavispora lusitaniae ATCC 42720 | Eukaryota | 0.04 |
| 1580 | 879301 | Lactobacillus iners LEAF 2053A-b | Bacteria | 0.04 |
| 1581 | 1914543 | Norovirus Hu/USA/2011/GI.P7_GI.7/CS5567 | Viruses | 0.04 |
| 1582 | 29385.22 | Staphylococcus saprophyticus strain JB027 | Bacteria | 0.04 |
| 1583 | 1161417.1 | Streptococcus sp. SR4 | Bacteria | 0.04 |
| 1584 | 1352.198 | Enterococcus faecium strain E161 | Bacteria | 0.04 |
| 1585 | 1339350 | Bacteroides vulgatus str. 3775 SL(B) 10 (iv) | Bacteria | 0.04 |
| 1586 | 54005.1 | Peptoniphilus harei strain CMW7756A | Bacteria | 0.04 |
| 1587 | 39397.1 | Candida sake strain CBS | Eukaryota | 0.04 |
| 1588 | 378833.2 | Sowbane mosaic virus isolate SoMV-WA | Viruses | 0.04 |
| 1589 | 1408415 | Acholeplasma equifetale ATCC 29724 | Bacteria | 0.04 |
| 1590 | 1408440 | Gemella sanguinis ATCC 700632 | Bacteria | 0.04 |
| 1591 | 1739381.1 | Streptococcus sp. HMSC072D03 | Bacteria | 0.04 |
| 1592 | 90371.51 | Salmonella enterica subsp. enterica serovar Typhimurium strain CFSAN033866 | Bacteria | 0.04 |
| 1593 | 49266.1 | Fucus vesiculosus | Eukaryota | 0.04 |
| 1594 | 529.5 | Ochrobactrum anthropi strain OAB | Bacteria | 0.04 |
| 1595 | 28901.23 | Salmonella enterica strain NGUA11 | Bacteria | 0.04 |
| 1596 | 575612 | Prevotella melaninogenica D18 | Bacteria | 0.04 |

| | | | | |
|---|---|---|---|---|
| 1597 | 12317.18 | Tobacco streak virus isolate 2334 | Viruses | 0.04 |
| 1598 | 1870984.1 | Anaerococcus mediterraneensis strain Marseille-P2765T sp. Marseille-P2765 | Bacteria | 0.04 |
| 1599 | 47736.1 | Carrot mottle mimic virus | Viruses | 0.04 |
| 1600 | 225992.1 | Comamonas kerstersii strain J29 | Bacteria | 0.04 |
| 1601 | 12263.4 | Squash mosaic virus strain Kimble | Viruses | 0.04 |
| 1602 | 888743 | Prevotella multiformis DSM 16608 | Bacteria | 0.03 |
| 1603 | 95609.1 | Herbaspirillum sp. B39 | Bacteria | 0.03 |
| 1604 | 1458705 | Norovirus Hu/GII.6/HS245/2010/USA | Viruses | 0.03 |
| 1605 | 1756285.1 | Maize associated totivirus isolate EC_Portoviejo | Viruses | 0.03 |
| 1606 | 1121947 | Helcococcus sueciensis DSM 17243 | Bacteria | 0.03 |
| 1607 | 62059.1 | Shallot yellow stripe virus | Viruses | 0.03 |
| 1608 | 1304.19 | Streptococcus salivarius strain 37-09 | Bacteria | 0.03 |
| 1609 | 112436.1 | Celery mosaic virus | Viruses | 0.03 |
| 1610 | 1118057 | Peptoniphilus grossensis ph5 | Bacteria | 0.03 |
| 1611 | 1435146.1 | Morganella sp. EGD-HP17 | Bacteria | 0.03 |
| 1612 | 1235479.1 | Pelosinus sp. HCF1 | Bacteria | 0.03 |
| 1613 | 1294024 | Calditerricola satsumensis JCM 14719 | Bacteria | 0.03 |
| 1614 | 1685.9 | Bifidobacterium breve strain BR-10 | Bacteria | 0.03 |
| 1615 | 1161918 | Brachyspira pilosicoli WesB | Bacteria | 0.03 |
| 1616 | 1681.5 | Bifidobacterium bifidum strain LMG | Bacteria | 0.03 |
| 1617 | 12139.3 | Southern bean mosaic virus isolate Sao | Viruses | 0.03 |
| 1618 | 12450.1 | Saccharomyces cerevisiae killer virus M1 | Viruses | 0.03 |
| 1619 | 1267.1 | Clostridium ventriculi | Bacteria | 0.03 |
| 1620 | 1352.164 | Enterococcus faecium strain E3 | Bacteria | 0.03 |
| 1621 | 5353.1 | Lentinula edodes | Eukaryota | 0.03 |
| 1622 | 1859694.1 | Haemophilus sp. CCUG 66565 | Bacteria | 0.03 |
| 1623 | 883092 | Lactobacillus crispatus FB077-07 | Bacteria | 0.03 |
| 1624 | 111015.1 | Actinomyces radicidentis strain CCUG | Bacteria | 0.03 |
| 1625 | 85154.1 | Streptococcus phage O1205 | Viruses | 0.03 |
| 1626 | 679937 | Bacteroides coprosuis DSM 18011 | Bacteria | 0.03 |
| 1627 | 1496.503 | Clostridioides difficile strain VRECD0027 | Bacteria | 0.03 |
| 1628 | 253701.2 | Zygocactus virus X | Viruses | 0.03 |
| 1629 | 253701.1 | Zygocactus virus X isolate P39 | Viruses | 0.03 |
| 1630 | 243563.1 | Strawberry necrotic shock virus isolate 840 | Viruses | 0.03 |
| 1631 | 1120988 | Anaerobiospirillum succiniciproducens DSM 6400 | Bacteria | 0.03 |
| 1632 | 2748.1 | Carnobacterium divergens isolate CDIV41 | Bacteria | 0.03 |
| 1633 | 1339280 | Bacteroides fragilis str. 2-F-2 #4 | Bacteria | 0.03 |
| 1634 | 1141139.1 | Enterobacteria phage vB_EcoP_ACG-C91 | Viruses | 0.03 |
| 1635 | 1624.2 | Lactobacillus salivarius strain 778_LSAL | Bacteria | 0.03 |
| 1636 | 458233 | Macrococcus caseolyticus JCSC5402 | Bacteria | 0.03 |
| 1637 | 1111734 | 4900 | Bacteria | 0.03 |
| 1638 | 1581148.1 | Clostridium sp. HMSC19A10 | Bacteria | 0.03 |
| 1639 | 33037.1 | Anaerococcus vaginalis strain PH9 | Bacteria | 0.03 |
| 1640 | 257758.5 | Streptococcus pseudopneumoniae strain 315_SPSE | Bacteria | 0.03 |
| 1641 | 419005.1 | Prevotella amnii strain DNF00307 | Bacteria | 0.03 |
| 1642 | 1307833 | Tannerella forsythia KS16 | Bacteria | 0.03 |

| | | | | |
|---|---|---|---|---|
| 1643 | 883165 | Campylobacter ureolyticus ACS-301-V-Sch3b | Bacteria | 0.03 |
| 1644 | 729.6 | Haemophilus parainfluenzae strain CCUG | Bacteria | 0.03 |
| 1645 | 72539.1 | Physalis mottle virus | Viruses | 0.03 |
| 1646 | 908338 | Peptoniphilus harei ACS-146-V-Sch2b | Bacteria | 0.03 |
| 1647 | 12313.1 | Peanut stunt virus | Viruses | 0.03 |
| 1648 | 1287488.1 | Prevotella sp. S7 MS 2 | Bacteria | 0.03 |
| 1649 | 1113547 | Salmonella phage 9NA | Viruses | 0.03 |
| 1650 | 470.622 | Acinetobacter baumannii strain XH545 | Bacteria | 0.03 |
| 1651 | 578361.3 | Soybean yellow mottle mosaic virus isolate New | Viruses | 0.03 |
| 1652 | 1141663 | Providencia rettgeri Dmel1 | Bacteria | 0.03 |
| 1653 | 1932006.1 | Chicken associated smacovirus strain RS/BR/2015/3 | Viruses | 0.03 |
| 1654 | 1496.648 | Clostridioides difficile isolate VL_0231 | Bacteria | 0.03 |
| 1655 | 369963 | Sapovirus Hu/Ehime475/2004/JP | Viruses | 0.03 |
| 1656 | 1120932 | Actinotignum schaalii DSM 15541 | Bacteria | 0.03 |
| 1657 | 388038.1 | Cucumber mottle virus | Viruses | 0.03 |
| 1658 | 1723382.1 | Peptoniphilaceae bacterium FC2 | Bacteria | 0.03 |
| 1659 | 469553.1 | Delftia sp. JD2 | Bacteria | 0.03 |
| 1660 | 112023.1 | Streptococcus phage 7201 | Viruses | 0.03 |
| 1661 | 1608898.1 | Haemophilus sp. HMSC71H05 | Bacteria | 0.03 |
| 1662 | 1658008.1 | Dysgonomonas sp. BGC7 | Bacteria | 0.03 |
| 1663 | 1379858 | Mucispirillum schaedleri ASF457 | Bacteria | 0.03 |
| 1664 | 1073387 | Bacteroides fragilis HMW 615 | Bacteria | 0.03 |
| 1665 | 698486 | Escherichia phage K1-dep(4) | Viruses | 0.03 |
| 1666 | 12274.29 | Grapevine fanleaf virus isolate GFLV-SDHN | Viruses | 0.03 |
| 1667 | 693746 | Oscillibacter valericigenes Sjm18-20 | Bacteria | 0.03 |
| 1668 | 1681.12 | Bifidobacterium bifidum strain 156B | Bacteria | 0.03 |
| 1669 | 1315956.1 | Shigella phage pSf-1 | Viruses | 0.03 |
| 1670 | 470.475 | Acinetobacter baumannii strain IHIT17966 | Bacteria | 0.03 |
| 1671 | 1123488 | Varibaculum cambriense DSM 15806 | Bacteria | 0.03 |
| 1672 | 1328.3 | Streptococcus anginosus strain 557_SANG | Bacteria | 0.03 |
| 1673 | 1318.5 | Streptococcus parasanguinis strain 886_SPAR | Bacteria | 0.03 |
| 1674 | 1318.4 | Streptococcus parasanguinis strain 65_SPAR | Bacteria | 0.03 |
| 1675 | 1423778 | Lactobacillus oligofermentans DSM 15707 = LMG 22743 | Bacteria | 0.03 |
| 1676 | 298338.1 | Lactobacillus phage LP65 | Viruses | 0.03 |
| 1677 | 1218109 | Propionibacterium propionicum NBRC 14587 | Bacteria | 0.03 |
| 1678 | 1492.4 | Clostridium butyricum strain HM-68 | Bacteria | 0.03 |
| 1679 | 665952 | Bacillus smithii 7_3_47FAA | Bacteria | 0.03 |
| 1680 | 1689.1 | Bifidobacterium dentium strain DE-29 | Bacteria | 0.03 |
| 1681 | 1225194 | Streptococcus mutans B24Sm2 | Bacteria | 0.03 |
| 1682 | 1526.2 | [Clostridium] aminophilum strain KH1P1 | Bacteria | 0.03 |
| 1683 | 1685.8 | Bifidobacterium breve strain BR-21 | Bacteria | 0.03 |
| 1684 | 1122986 | Prevotella maculosa DSM 19339 = JCM 15638 | Bacteria | 0.03 |
| 1685 | 344022.1 | Escherichia virus K1E | Viruses | 0.03 |
| 1686 | 35288.1 | Grapevine virus A isolate 3138-03 | Viruses | 0.03 |
| 1687 | 1871031.1 | Olsenella sp. Marseille-P3256 sp. Marseille-P3256 | Bacteria | 0.03 |
| 1688 | 879310 | Selenomonas sp. oral taxon 137 str. F0430 | Bacteria | 0.03 |

| | | | | |
|---|---|---|---|---|
| 1689 | 1129191 | Klebsiella phage KP36 | Viruses | 0.03 |
| 1690 | 12470.2 | Lucerne transient streak virus | Viruses | 0.03 |
| 1691 | 1111121.1 | Atopobium sp. BV3Ac4 | Bacteria | 0.03 |
| 1692 | 1922434.2 | Beihai mollusks virus 1 strain WZSLuoI86140 | Viruses | 0.03 |
| 1693 | 51354.1 | Maize chlorotic dwarf virus strain Severe | Viruses | 0.03 |
| 1694 | 521000 | Providencia rettgeri DSM 1131 | Bacteria | 0.03 |
| 1695 | 521002 | Methanobrevibacter smithii DSM 2374 | Archaea | 0.03 |
| 1696 | 1027232.1 | Groundnut ringspot and Tomato chlorotic spot virus reassortant | Viruses | 0.03 |
| 1697 | 1922513.1 | HOU234589 | Viruses | 0.03 |
| 1698 | 1114922 | Citrobacter farmeri GTC 1319 | Bacteria | 0.03 |
| 1699 | 866771 | Prevotella disiens FB035-09AN | Bacteria | 0.03 |
| 1700 | 988645.1 | Raspberry leaf blotch virus | Viruses | 0.03 |
| 1701 | 439334.28 | Mycobacterium avium subsp. hominissuis strain MAH-E-104-1 | Bacteria | 0.03 |
| 1702 | 237561 | Candida albicans SC5314 | Eukaryota | 0.03 |
| 1703 | 1121933 | Granulicoccus phenolivorans DSM 17626 | Bacteria | 0.03 |
| 1704 | 1739309.1 | Streptococcus sp. HMSC034E03 | Bacteria | 0.03 |
| 1705 | 1590596.1 | Sphingobacterium sp. T2 | Bacteria | 0.03 |
| 1706 | 1659298.2 | Nectarine stem pitting-associated virus isolate NSPaV/12P42 | Viruses | 0.03 |
| 1707 | 42783 | Coxsackievirus A22 | Viruses | 0.03 |
| 1708 | 1156433.1 | Streptococcus sp. I-P16 sp. I-P16 | Bacteria | 0.03 |
| 1709 | 1496.68 | Clostridioides difficile isolate VL_0167 | Bacteria | 0.03 |
| 1710 | 1304.26 | Streptococcus salivarius strain GED7778A | Bacteria | 0.03 |
| 1711 | 1718158.1 | Enterococcus phage IME-EFm5 | Viruses | 0.03 |
| 1712 | 1287476 | Prevotella bivia DNF00188 | Bacteria | 0.03 |
| 1713 | 1683.2 | Bifidobacterium angulatum strain GT102 | Bacteria | 0.03 |
| 1714 | 106008.1 | Curvibasidium cygneicollum | Eukaryota | 0.03 |
| 1715 | 1567484.1 | Lactobacillus phage LfeInf | Viruses | 0.03 |
| 1716 | 12165.1 | Chrysanthemum virus B | Viruses | 0.03 |
| 1717 | 290399.2 | Arthrobacter sp. FB24 | Bacteria | 0.03 |
| 1718 | 47715.3 | Lactobacillus rhamnosus strain Lrh22 | Bacteria | 0.03 |
| 1719 | 1891289.1 | Sporanaerobacter sp. PP17-6a isolate PP176A | Bacteria | 0.03 |
| 1720 | 1496.74 | Clostridioides difficile strain CD105KSE1 | Bacteria | 0.03 |
| 1721 | 879212 | Desulfobacter postgatei 2ac9 | Bacteria | 0.03 |
| 1722 | 1930509.1 | Husavirus sp. isolate 16370_59, sp. isolate | Viruses | 0.03 |
| 1723 | 529.4 | Ochrobactrum anthropi strain ML7 | Bacteria | 0.03 |
| 1724 | 340.1 | Xanthomonas campestris pv. campestris | Bacteria | 0.03 |
| 1725 | 1702287.1 | Negativicoccus massiliensis strain AT7 | Bacteria | 0.03 |
| 1726 | 1130798 | Lactobacillus mucosae LM1 | Bacteria | 0.03 |
| 1727 | 12208.1 | Pea seed-borne mosaic virus | Viruses | 0.02 |
| 1728 | 12431.3 | Garlic virus C isolate SW3.3B | Viruses | 0.02 |
| 1729 | 469594.1 | Bifidobacterium sp. 12_1_47BFAA | Bacteria | 0.02 |
| 1730 | 1398.2 | Bacillus coagulans strain B4098 | Bacteria | 0.02 |
| 1731 | 1423780 | Lactobacillus otakiensis DSM 19908 = JCM 15040 | Bacteria | 0.02 |
| 1732 | 1608882.1 | Haemophilus sp. HMSC61B11 | Bacteria | 0.02 |

| | | | | |
|---|---|---|---|---|
| 1733 | 596330 | Peptoniphilus lacrimalis 315-B | Bacteria | 0.02 |
| 1734 | 424716.1 | Salmonella phage Vi II-E1 | Viruses | 0.02 |
| 1735 | 483913.1 | Bacillus subtilis subsp. inaquosorum strain DE111 | Bacteria | 0.02 |
| 1736 | 91753.8 | Cucurbit aphid-borne yellows virus strain Sq/2005/9.2 | Viruses | 0.02 |
| 1737 | 693582.1 | Pseudomonas phage phi-2 | Viruses | 0.02 |
| 1738 | 1517899.1 | Kytococcus sp. CUA-901 | Bacteria | 0.02 |
| 1739 | 243164 | Dehalococcoides mccartyi 195 | Bacteria | 0.02 |
| 1740 | 1922435.1 | Beihai mollusks virus 2 strain WZSLuoI86113 | Viruses | 0.02 |
| 1741 | 1759399.1 | Streptococcus sp. A12 sp. A12 | Bacteria | 0.02 |
| 1742 | 1118056 | Anaerococcus obesiensis ph10 | Bacteria | 0.02 |
| 1743 | 726.4 | Haemophilus haemolyticus strain 27P25 | Bacteria | 0.02 |
| 1744 | 392504.3 | Turnip ringspot virus | Viruses | 0.02 |
| 1745 | 392504.4 | Turnip ringspot virus isolate M12 | Viruses | 0.02 |
| 1746 | 36015.1 | Pichia kluyveri strain CBS | Eukaryota | 0.02 |
| 1747 | 1215915 | Lactococcus raffinolactis 4877 | Bacteria | 0.02 |
| 1748 | 1301100.1 | [Clostridium] dakarense strain FF1 | Bacteria | 0.02 |
| 1749 | 67962.3 | Carrot red leaf luteovirus associated RNA clone sigma | Viruses | 0.02 |
| 1750 | 936578.1 | Streptococcus sp. AS20 | Bacteria | 0.02 |
| 1751 | 1379.2 | Gemella haemolysans strain DNF01167 | Bacteria | 0.02 |
| 1752 | 79604.1 | Denitrobacterium detoxificans strain DSM | Bacteria | 0.02 |
| 1753 | 1496722.1 | Butyrivibrio sp. AE2005 | Bacteria | 0.02 |
| 1754 | 1123404 | Tissierella praeacuta DSM 18095 | Bacteria | 0.02 |
| 1755 | 137591.4 | Weissella cibaria strain MG1 | Bacteria | 0.02 |
| 1756 | 137591.5 | Weissella cibaria strain AB3b | Bacteria | 0.02 |
| 1757 | 1457186 | Bifidobacterium longum subsp. longum EK13 | Bacteria | 0.02 |
| 1758 | 562.2176 | Escherichia coli strain MN067 | Bacteria | 0.02 |
| 1759 | 1613.5 | Lactobacillus fermentum strain LfQi6 | Bacteria | 0.02 |
| 1760 | 1613.6 | Lactobacillus fermentum strain RI-508 | Bacteria | 0.02 |
| 1761 | 1078773 | Herbaspirillum rubrisubalbicans M1 | Bacteria | 0.02 |
| 1762 | 134632.1 | American plum line pattern virus | Viruses | 0.02 |
| 1763 | 485.83 | Neisseria gonorrhoeae strain m07.05 | Bacteria | 0.02 |
| 1764 | 1502.2 | Clostridium perfringens strain 1207_CPER | Bacteria | 0.02 |
| 1765 | 1496.601 | Clostridioides difficile isolate VL_0330 | Bacteria | 0.02 |
| 1766 | 1236526 | Porphyromonas gingivicanis JCM 15907 | Bacteria | 0.02 |
| 1767 | 196400.5 | Grapevine rupestris stem pitting-associated virus isolate GRSPaV-JF | Viruses | 0.02 |
| 1768 | 12162.44 | Citrus tristeza virus isolate T68-1 | Viruses | 0.02 |
| 1769 | 12024.1 | Pseudomonas phage PRR1 | Viruses | 0.02 |
| 1770 | 1423760 | Lactobacillus ingluviei DSM 15946 | Bacteria | 0.02 |
| 1771 | 1235793.1 | Lachnospiraceae bacterium COE1 | Bacteria | 0.02 |
| 1772 | 12430.1 | Garlic virus D isolate Mesi | Viruses | 0.02 |
| 1773 | 184870.1 | Varibaculum cambriense strain DNF00696 | Bacteria | 0.02 |
| 1774 | 1410662 | Lachnospira multipara MC2003 | Bacteria | 0.02 |
| 1775 | 402626 | Ralstonia pickettii 12J | Bacteria | 0.02 |
| 1776 | 1095730 | Streptococcus constellatus subsp. constellatus SK53 | Bacteria | 0.02 |

| | | | | |
|---|---|---|---|---|
| 1777 | 1445858.1 | Enterococcus phage IME-EFm1 | Viruses | 0.02 |
| 1778 | 1596.8 | Lactobacillus gasseri strain AL5 | Bacteria | 0.02 |
| 1779 | 339420.3 | Blackberry chlorotic ringspot virus | Viruses | 0.02 |
| 1780 | 1908263.1 | Rodentibacter trehalosifermentans strain H1987082031 | Bacteria | 0.02 |
| 1781 | 762051 | Leuconostoc kimchii IMSNU 11154 | Bacteria | 0.02 |
| 1782 | 260742.1 | Streptomyces sp. SS | Bacteria | 0.02 |
| 1783 | 1236514 | Bacteroides stercorirosoris JCM 17103 | Bacteria | 0.02 |
| 1784 | 1496.181 | Clostridioides difficile isolate VL_0239 | Bacteria | 0.02 |
| 1785 | 140626.1 | Lachnobacterium bovis strain S1b | Bacteria | 0.02 |
| 1786 | 1923170.1 | Hubei polero-like virus 2 strain QTM26674 | Viruses | 0.02 |
| 1787 | 1448141 | Trueperella pyogenes MS249 | Bacteria | 0.02 |
| 1788 | 1272.1 | Kocuria varians strain G6 | Bacteria | 0.02 |
| 1789 | 632112.1 | Lactobacillus phage Lb338-1 | Viruses | 0.02 |
| 1790 | 1385385.1 | Streptococcus phage TP-778L | Viruses | 0.02 |
| 1791 | 4950.2 | Torulaspora delbrueckii | Eukaryota | 0.02 |
| 1792 | 573061 | Clostridium cellulovorans 743B | Bacteria | 0.02 |
| 1793 | 1598.19 | Lactobacillus reuteri strain I49 | Bacteria | 0.02 |
| 1794 | 1255.2 | Pediococcus pentosaceus strain NKYL15 | Bacteria | 0.02 |
| 1795 | 562.1794 | Escherichia coli strain GN02091 | Bacteria | 0.02 |
| 1796 | 1739491.1 | Streptococcus sp. HMSC067H01 | Bacteria | 0.02 |
| 1797 | 154339.1 | Little cherry virus 2 | Viruses | 0.02 |
| 1798 | 562.979 | Escherichia coli strain upec-127 | Bacteria | 0.02 |
| 1799 | 637909 | Streptococcus gallolyticus UCN34 | Bacteria | 0.02 |
| 1800 | 65467.5 | Cherry green ring mottle virus isolate S10 | Viruses | 0.02 |
| 1801 | 1321816 | Alloscardovia omnicolens F0580 | Bacteria | 0.02 |
| 1802 | 1229757 | Escherichia phage EC6 | Viruses | 0.02 |
| 1803 | 1120942 | Actinomyces georgiae DSM 6843 | Bacteria | 0.02 |
| 1804 | 1658779.1 | Porphyromonadaceae bacterium H1 strain H2 | Bacteria | 0.02 |
| 1805 | 218923.3 | Turnip rosette virus isolate TRoV-2 | Viruses | 0.02 |
| 1806 | 1304.18 | Streptococcus salivarius strain 40-02 | Bacteria | 0.02 |
| 1807 | 1529923 | Norovirus GII/Hu/JP/2010/GII.P7_GII.7/Musashimurayama/TAKAsanKimchi | Viruses | 0.02 |
| 1808 | 1280693 | Clostridium butyricum AGR2140 | Bacteria | 0.02 |
| 1809 | 12242.49 | Tobacco mosaic virus strain pet-TW | Viruses | 0.02 |
| 1810 | 999405 | [Clostridium] clostridioforme 90A4 | Bacteria | 0.02 |
| 1811 | 1889813.1 | Anaerolineaceae bacterium oral taxon 439 strain W11661 | Bacteria | 0.02 |
| 1812 | 1122154 | Lactococcus chungangensis CAU 28 = DSM 22330 | Bacteria | 0.02 |
| 1813 | 55951.12 | Grapevine leafroll-associated virus 3 isolate GH24 | Viruses | 0.02 |
| 1814 | 1229753.1 | Escherichia phage phAPEC8 | Viruses | 0.02 |
| 1815 | 1496.402 | Clostridioides difficile strain VRECD0053 | Bacteria | 0.02 |
| 1816 | 1033733 | Anaerococcus senegalensis JC48 | Bacteria | 0.02 |
| 1817 | 42882.3 | Cherry virus A | Viruses | 0.02 |
| 1818 | 729.8 | Haemophilus parainfluenzae strain 432_HPAR | Bacteria | 0.02 |
| 1819 | 729.5 | Haemophilus parainfluenzae strain 488_HPAR | Bacteria | 0.02 |

| | | | | |
|---|---|---|---|---|
| 1820 | 95340 | Norwalk-like virus | Viruses | 0.02 |
| 1821 | 1739465.1 | Enterococcus sp. HMSC076E04 | Bacteria | 0.02 |
| 1822 | 353496 | Lactobacillus delbrueckii subsp. bulgaricus 2038 | Bacteria | 0.02 |
| 1823 | 1330524.3 | Salivirus A isolate BN-5 | Viruses | 0.02 |
| 1824 | 28131.1 | Prevotella intermedia strain ATCC | Bacteria | 0.02 |
| 1825 | 980518 | Enterobacter mori LMG 25706 | Bacteria | 0.02 |
| 1826 | 1477000.1 | Peptoniphilus sp. DNF00840 | Bacteria | 0.02 |
| 1827 | 1564903.1 | Strawberry polerovirus 1 isolate AB5301 | Viruses | 0.02 |
| 1828 | 443746.1 | Asparagus virus 1 isolate DSMZ | Viruses | 0.02 |
| 1829 | 1795648.1 | Picornavirales Tottori-HG1 | Viruses | 0.02 |
| 1830 | 682382.1 | HMO Astrovirus A | Viruses | 0.02 |
| 1831 | 1423730 | Lactobacillus camelliae DSM 22697 = JCM 13995 | Bacteria | 0.02 |
| 1832 | 1423739 | Lactobacillus diolivorans DSM 14421 | Bacteria | 0.02 |
| 1833 | 1647408.1 | Klebsiella phage KLPN1 | Viruses | 0.02 |
| 1834 | 1316931 | Clostridium butyricum DSM 10702 | Bacteria | 0.02 |
| 1835 | 749551 | Selenomonas artemidis F0399 | Bacteria | 0.02 |
| 1836 | 944557 | Prevotella denticola CRIS 18C-A | Bacteria | 0.02 |
| 1837 | 1035189 | Streptococcus infantis SK970 | Bacteria | 0.02 |
| 1838 | 12317.5 | Tobacco streak virus isolate OK | Viruses | 0.02 |
| 1839 | 1318.21 | Streptococcus parasanguinis strain DD19 | Bacteria | 0.02 |
| 1840 | 1200793 | Streptococcus salivarius K12 | Bacteria | 0.02 |
| 1841 | 553218 | Campylobacter rectus RM3267 | Bacteria | 0.02 |
| 1842 | 1932006.2 | Chicken associated smacovirus strain RS/BR/2015/4 | Viruses | 0.02 |
| 1843 | 1206545.1 | Klebsiella phage 0507-KN2-1 | Viruses | 0.02 |
| 1844 | 685899.1 | Papaya lethal yellowing virus | Viruses | 0.02 |
| 1845 | 1095768 | Enterobacter massiliensis JC163 | Bacteria | 0.02 |
| 1846 | 300717 | Sapovirus NongKhai-24/Thailand | Viruses | 0.02 |
| 1847 | 5478.1 | [Candida] glabrata | Eukaryota | 0.02 |
| 1848 | 1597.12 | Lactobacillus paracasei strain DSM | Bacteria | 0.02 |
| 1849 | 1410625.1 | Lachnospiraceae bacterium MD2004 | Bacteria | 0.02 |
| 1850 | 936577.1 | Streptococcus sp. AS14 | Bacteria | 0.02 |
| 1851 | 12275.1 | Tomato black ring virus strain ED) | Viruses | 0.02 |
| 1852 | 1073353 | Campylobacter showae CC57C | Bacteria | 0.02 |
| 1853 | 1444211 | Escherichia coli 5-172-05_S4_C1 | Bacteria | 0.02 |
| 1854 | 150055.1 | Streptococcus lutetiensis strain DD06 | Bacteria | 0.02 |
| 1855 | 1172562 | Helicobacter cinaedi PAGU611 | Bacteria | 0.02 |
| 1856 | 948870.1 | Enterobacteria phage phi92 | Viruses | 0.02 |
| 1857 | 33964.1 | Leuconostoc citreum strain 1301_LGAS | Bacteria | 0.02 |
| 1858 | 1353.2 | Enterococcus gallinarum strain SKF1 | Bacteria | 0.02 |
| 1859 | 1111137.1 | Slackia sp. CM382 | Bacteria | 0.02 |
| 1860 | 1123309 | Streptococcus minor DSM 17118 | Bacteria | 0.02 |
| 1861 | 1682.1 | CECT | Bacteria | 0.02 |
| 1862 | 331278.1 | Yersinia phage phiR1-37 | Viruses | 0.02 |
| 1863 | 471285.1 | Lettuce yellow mottle virus | Viruses | 0.02 |
| 1864 | 1095770 | Peptoniphilus timonensis JC401 | Bacteria | 0.02 |
| 1865 | 1475062.1 | Porcine stool-associated circular virus 4 isolate CP2 | Viruses | 0.02 |

| | | | | |
|---|---|---|---|---|
| 1866 | 689781.1 | Oribacterium sp. NK2B42 | Bacteria | 0.02 |
| 1867 | 1354300.1 | Peptoniphilus sp. ChDC B134 | Bacteria | 0.02 |
| 1868 | 1276.2 | Kytococcus sedentarius strain 1083_KSED | Bacteria | 0.02 |
| 1869 | 1197906 | Afipia birgiae 34632 | Bacteria | 0.02 |
| 1870 | 12041.2 | Bean leafroll virus | Viruses | 0.02 |
| 1871 | 613026 | Helicobacter bilis ATCC 43879 | Bacteria | 0.02 |
| 1872 | 56879.1 | Oat blue dwarf virus isolate OBDV-2r | Viruses | 0.02 |
| 1873 | 1339351 | Bacteroides vulgatus str. 3775 SR(B) 19 | Bacteria | 0.02 |
| 1874 | 1073366 | Prevotella nigrescens CC14M | Bacteria | 0.02 |
| 1875 | 112227.3 | Cactus virus X isolate NTU | Viruses | 0.02 |
| 1876 | 1354264 | Kluyvera georgiana ATCC 51603 | Bacteria | 0.02 |
| 1877 | 1381124 | Lactobacillus fermentum 3872 | Bacteria | 0.02 |
| 1878 | 575594 | Lactobacillus coleohominis 101-4-CHN | Bacteria | 0.02 |
| 1879 | 1679.11 | Bifidobacterium longum subsp. longum strain LO-10 | Bacteria | 0.02 |
| 1880 | 525363 | Lactobacillus sakei subsp. carnosus DSM 15831 | Bacteria | 0.02 |
| 1881 | 1599.3 | Lactobacillus sakei strain RI-412 | Bacteria | 0.02 |
| 1882 | 29388.7 | Staphylococcus capitis strain 129_SAUR | Bacteria | 0.02 |
| 1883 | 1283279 | Saccharopolyspora rectivirgula DSM 43747 | Bacteria | 0.02 |
| 1884 | 28141.1 | Cronobacter sakazakii strain HPB5174 | Bacteria | 0.02 |
| 1885 | 1857568.1 | Macellibacteroides sp. HH-ZS | Bacteria | 0.02 |
| 1886 | 35703.8 | Citrobacter amalonaticus strain FDAARGOS_122 | Bacteria | 0.02 |
| 1887 | 51288.1 | Kluyvera ascorbata strain WCH1410 | Bacteria | 0.02 |
| 1888 | 1318.6 | Streptococcus parasanguinis strain 766_SPAR | Bacteria | 0.02 |
| 1889 | 218667.1 | Oyster mushroom spherical virus | Viruses | 0.02 |
| 1890 | 1522179.1 | Asterionellopsis glacialis RNA virus | Viruses | 0.02 |
| 1891 | 1339301 | Bacteroides fragilis str. 3397 N3 | Bacteria | 0.02 |
| 1892 | 1581071.1 | Granulicatella sp. HMSC30F09 | Bacteria | 0.02 |
| 1893 | 1685.1 | Bifidobacterium breve strain BR-L29 | Bacteria | 0.02 |
| 1894 | 596322 | Streptococcus salivarius SK126 | Bacteria | 0.02 |
| 1895 | 683173 | Astrovirus VA2 | Viruses | 0.02 |
| 1896 | 52253.1 | Candida sojae | Eukaryota | 0.02 |
| 1897 | 758863 | Sapovirus Hu/GI/Sapporo/MT-2010/1982 | Viruses | 0.02 |
| 1898 | 39443.2 | Carnation Italian ringspot virus isolate CZ | Viruses | 0.02 |
| 1899 | 1195163.1 | Amazon lily mild mottle virus | Viruses | 0.02 |
| 1900 | 12165.3 | Chrysanthemum virus B isolate Tamil | Viruses | 0.02 |
| 1901 | 315405.1 | Streptococcus gallolyticus strain ICDDRB-NRC-S3 | Bacteria | 0.02 |
| 1902 | 34073.1 | Variovorax paradoxus strain H112 | Bacteria | 0.02 |
| 1903 | 1785087.1 | Candidatus Protochlamydia sp. W-9 sp. W-9 | Bacteria | 0.02 |
| 1904 | 1122988 | Prevotella nanceiensis DSM 19126 = JCM 15639 | Bacteria | 0.02 |
| 1905 | 1303.18 | Streptococcus oralis strain DD24 | Bacteria | 0.02 |
| 1906 | 641149 | Neisseria sp. oral taxon 014 str. F0314 | Bacteria | 0.02 |
| 1907 | 29363.3 | Clostridium paraputrificum strain 2789STDY5834857 | Bacteria | 0.02 |
| 1908 | 1715012.1 | Enterococcus sp. HMSC072H05 | Bacteria | 0.02 |
| 1909 | 1852625.1 | Klebsiella phage vB_KpnM_KpV477 | Viruses | 0.02 |
| 1910 | 1081640 | Sphingomonas elodea ATCC 31461 | Bacteria | 0.02 |

| | | | | |
|---|---|---|---|---|
| 1911 | 1127128 | Leuconostoc citreum LBAE C11 | Bacteria | 0.02 |
| 1912 | 28348.2 | Sweet clover necrotic mosaic virus | Viruses | 0.02 |
| 1913 | 1665651 | Norovirus GII/Hu/NL/2014/GII.2/Groningen | Viruses | 0.02 |
| 1914 | 1211388.1 | Apple green crinkle associated virus isolate Aurora-1 | Viruses | 0.02 |
| 1915 | 50948.1 | Enterobacteria phage RB49 | Viruses | 0.02 |
| 1916 | 1261065 | Gardnerella vaginalis JCP8070 | Bacteria | 0.02 |
| 1917 | 727.77 | Haemophilus influenzae strain 159_HINF | Bacteria | 0.02 |
| 1918 | 1354263 | Hafnia paralvei ATCC 29927 | Bacteria | 0.02 |
| 1919 | 1913024.1 | White clover mottle virus | Viruses | 0.02 |
| 1920 | 227507.1 | Strawberry pallidosis-associated virus | Viruses | 0.02 |
| 1921 | 1193128 | Parascardovia denticolens IPLA 20019 | Bacteria | 0.02 |
| 1922 | 1280675.1 | Bifidobacterium sp. AGR2158 | Bacteria | 0.02 |
| 1923 | 1739450.1 | Actinomyces sp. HMSC062G12 | Bacteria | 0.02 |
| 1924 | 879306 | Anaerococcus hydrogenalis ACS-025-V-Sch4 | Bacteria | 0.02 |
| 1925 | 997352 | Prevotella nigrescens ATCC 33563 | Bacteria | 0.02 |
| 1926 | 1630.1 | Kandleria vitulina strain WCC7 | Bacteria | 0.02 |
| 1927 | 1906334.1 | Corynebacterium sp. NML140438 | Bacteria | 0.02 |
| 1928 | 596153 | Alicycliphilus denitrificans BC | Bacteria | 0.02 |
| 1929 | 46170.38 | Staphylococcus aureus subsp. aureus strain SA-120 | Bacteria | 0.02 |
| 1930 | 1610829 | Klebsiella phage 1513 | Viruses | 0.02 |
| 1931 | 888064 | Enterococcus italicus DSM 15952 | Bacteria | 0.02 |
| 1932 | 1923554.1 | Wenzhou bivalvia virus 2 strain beimix75763 | Viruses | 0.02 |
| 1933 | 585.5 | Proteus vulgaris strain ATCC | Bacteria | 0.02 |
| 1934 | 199.3 | Campylobacter concisus strain ATCC | Bacteria | 0.02 |
| 1935 | 469607 | Fusobacterium nucleatum subsp. animalis 4_8 | Bacteria | 0.02 |
| 1936 | 1416758 | Morganella morganii H1r | Bacteria | 0.02 |
| 1937 | 1547495.1 | Salivirus FHB | Viruses | 0.02 |
| 1938 | 1204476 | Fusobacterium nucleatum CTI-3 | Bacteria | 0.02 |
| 1939 | 702439 | Prevotella nigrescens F0103 | Bacteria | 0.02 |
| 1940 | 1868658.1 | Human astrovirus | Viruses | 0.02 |
| 1941 | 1327956 | Escherichia phage JES2013 | Viruses | 0.02 |
| 1942 | 12055.1 | Tobacco necrosis virus A | Viruses | 0.02 |
| 1943 | 868164 | Escherichia coli DEC6E | Bacteria | 0.02 |
| 1944 | 570949.1 | Carrot mottle mimic virus satellite RNA | Viruses | 0.02 |
| 1945 | 1408324.1 | Lachnospiraceae bacterium MC2017 | Bacteria | 0.02 |
| 1946 | 582.3 | Morganella morganii strain L3 | Bacteria | 0.02 |
| 1947 | 28038.4 | Lactobacillus curvatus strain RI-406 | Bacteria | 0.02 |
| 1948 | 1339291 | Bacteroides fragilis str. S23 R14 | Bacteria | 0.02 |
| 1949 | 10590 | Human papillomavirus type 42 | Viruses | 0.02 |
| 1950 | 351495.1 | Raphanus sativus cryptic virus 2 | Viruses | 0.02 |
| 1951 | 12433.3 | Garlic virus A isolate SW3.1A | Viruses | 0.02 |
| 1952 | 12433.4 | Garlic virus A isolate GarVA-SP | Viruses | 0.02 |
| 1953 | 12049.1 | Soybean dwarf virus | Viruses | 0.02 |
| 1954 | 1590.151 | Lactobacillus plantarum strain MF1298 | Bacteria | 0.02 |
| 1955 | 12431.2 | Garlic virus C isolate SW3.3A | Viruses | 0.02 |
| 1956 | 399183 | Escherichia phage V5 | Viruses | 0.02 |

| | | | | |
|---|---|---|---|---|
| 1957 | 1698360.1 | Klebsiella phage JD18 | Viruses | 0.02 |
| 1958 | 1304.6 | Streptococcus salivarius strain 1003_SOLI | Bacteria | 0.02 |
| 1959 | 1852368.1 | Prevotellaceae bacterium Marseille-P2826 | Bacteria | 0.02 |
| 1960 | 315405.11 | Streptococcus gallolyticus strain DD02 | Bacteria | 0.02 |
| 1961 | 722911 | Bifidobacterium longum subsp. longum F8 | Bacteria | 0.02 |
| 1962 | 1563222.1 | Citrobacter pasteurii strain CIP | Bacteria | 0.02 |
| 1963 | 217686.1 | Little cherry virus 1 | Viruses | 0.02 |
| 1964 | 1339347 | Bacteroides ovatus str. 3725 D9 iii | Bacteria | 0.02 |
| 1965 | 28347.11 | Apple stem grooving virus isolate ASGV-CHN | Viruses | 0.02 |
| 1966 | 499174 | Clostridioides difficile QCD-23m63 | Bacteria | 0.02 |
| 1967 | 1537165.1 | Porcine stool-associated circular virus 6 isolate XP1 | Viruses | 0.02 |
| 1968 | 1163671.1 | Clostridium sp. 12(A) | Bacteria | 0.02 |
| 1969 | 156976.1 | Corynebacterium riegelii strain PUDD_83A45 | Bacteria | 0.02 |
| 1970 | 1737425.1 | Corynebacterium provencense strain SN15 | Bacteria | 0.02 |
| 1971 | 1032457.1 | Passion fruit mosaic virus | Viruses | 0.02 |
| 1972 | 53655.1 | Pichia fermentans | Eukaryota | 0.02 |
| 1973 | 1654357.1 | La Jolla virus | Viruses | 0.02 |
| 1974 | 1002805 | Helicobacter bizzozeronii CCUG 35545 | Bacteria | 0.02 |
| 1975 | 1127129 | Leuconostoc citreum LBAE E16 | Bacteria | 0.02 |
| 1976 | 1654356.1 | Thika virus | Viruses | 0.02 |
| 1977 | 562.496 | Escherichia coli strain 592_ECOL | Bacteria | 0.02 |
| 1978 | 1581074.1 | Granulicatella sp. HMSC31F03 | Bacteria | 0.02 |
| 1979 | 1131317 | Escherichia phage FV3 | Viruses | 0.02 |
| 1980 | 1665644 | Norovirus GI/Hu/NL/2011/GI.4/Groningen | Viruses | 0.02 |
| 1981 | 1183241.1 | Persimmon cryptic virus isolate SSPI | Viruses | 0.02 |
| 1982 | 359987.1 | Rhizosolenia setigera RNA virus 01 | Viruses | 0.02 |
| 1983 | 59241.1 | Streptococcus phage Dp-1 | Viruses | 0.02 |
| 1984 | 1211817 | Clostridium ihumii AP5 | Bacteria | 0.02 |
| 1985 | 1923116.1 | Hubei picorna-like virus 36 strain SCM50248 | Viruses | 0.02 |
| 1986 | 1121269 | Campylobacter upsaliensis DSM 5365 | Bacteria | 0.02 |
| 1987 | 1123489 | Veillonella magna DSM 19857 | Bacteria | 0.02 |
| 1988 | 562.1822 | Escherichia coli strain SF-491 | Bacteria | 0.02 |
| 1989 | 1351.11 | Enterococcus faecalis | Bacteria | 0.02 |
| 1990 | 92395.1 | Black queen cell virus strain PP | Viruses | 0.01 |
| 1991 | 196400.16 | Grapevine rupestris stem pitting-associated virus isolate GRSPaV-MG | Viruses | 0.01 |
| 1992 | 1218493.1 | Lactobacillus kullabergensis strain Biut2 | Bacteria | 0.01 |
| 1993 | 1429888 | Leuconostoc mesenteroides P45 | Bacteria | 0.01 |
| 1994 | 1284702 | Mageeibacillus indolicus 0009-5 | Bacteria | 0.01 |
| 1995 | 64003.1 | Grapevine leafroll-associated virus 2 isolate GLRaV-2-SG | Viruses | 0.01 |
| 1996 | 1229204.1 | alpha proteobacterium L41A | Bacteria | 0.01 |
| 1997 | 1093141 | Nannochloropsis gaditana CCMP526 | Eukaryota | 0.01 |
| 1998 | 336306.3 | Enterobacter cloacae subsp. cloacae strain GN02616 | Bacteria | 0.01 |
| 1999 | 11987.3 | Melon necrotic spot virus isolate Malfa5 | Viruses | 0.01 |
| 2000 | 216816.19 | Bifidobacterium longum strain BG7 | Bacteria | 0.01 |

| | | | | |
|---|---|---|---|---|
| 2001 | 1505227.1 | Aeromonas phage pAh6-C | Viruses | 0.01 |
| 2002 | 329.1 | Ralstonia pickettii strain H2Cu5 | Bacteria | 0.01 |
| 2003 | 349519 | Leuconostoc citreum KM20 | Bacteria | 0.01 |
| 2004 | 1551.1 | Clostridium aurantibutyricum strain DSM | Bacteria | 0.01 |
| 2005 | 169292.4 | Corynebacterium aurimucosum strain 620_CAUR | Bacteria | 0.01 |
| 2006 | 1566990.1 | Streptococcus phage SpSL1 | Viruses | 0.01 |
| 2007 | 1502.38 | Clostridium perfringens strain FORC_025 | Bacteria | 0.01 |
| 2008 | 641487.1 | Lactococcus phage P087 | Viruses | 0.01 |
| 2009 | 1235801 | Lactobacillus murinus ASF361 | Bacteria | 0.01 |
| 2010 | 649760 | Prevotella oris F0302 | Bacteria | 0.01 |
| 2011 | 577.2 | Raoultella terrigena strain NZ133 | Bacteria | 0.01 |
| 2012 | 1297581 | Anoxybacillus flavithermus AK1 | Bacteria | 0.01 |
| 2013 | 562.1581 | Escherichia coli strain GN02461 | Bacteria | 0.01 |
| 2014 | 1304.15 | Streptococcus salivarius strain 84-12 | Bacteria | 0.01 |
| 2015 | 546.1 | Citrobacter freundii strain CF7_ST91 | Bacteria | 0.01 |
| 2016 | 620833 | Fusobacterium periodonticum D10 | Bacteria | 0.01 |
| 2017 | 1302.16 | Streptococcus gordonii strain 1116_SGOR | Bacteria | 0.01 |
| 2018 | 762550 | Leuconostoc gelidum subsp. gasicomitatum LMG 18811 | Bacteria | 0.01 |
| 2019 | 633135.1 | Streptococcus phage Abc2 | Viruses | 0.01 |
| 2020 | 448384.1 | Enterobacteria phage Phi1 | Viruses | 0.01 |
| 2021 | 1868652.2 | High Plains wheat mosaic virus | Viruses | 0.01 |
| 2022 | 1868652.1 | High Plains wheat mosaic virus isolate Nebraska | Viruses | 0.01 |
| 2023 | 1284663 | Lactobacillus plantarum ZJ316 | Bacteria | 0.01 |
| 2024 | 1365969 | Bifidobacterium breve MCC 1605 | Bacteria | 0.01 |
| 2025 | 1787.2 | Mycobacterium szulgai strain ACS1160 | Bacteria | 0.01 |
| 2026 | 1898961.1 | Kluyvera intestini strain GT-16 | Bacteria | 0.01 |
| 2027 | 12317.4 | Tobacco streak virus isolate dp | Viruses | 0.01 |
| 2028 | 12188.1 | White clover mosaic virus | Viruses | 0.01 |
| 2029 | 1425364.2 | Carrot torradovirus 1 isolate CTV-1_RNA1_H6 | Viruses | 0.01 |
| 2030 | 392504.2 | Turnip ringspot virus isolate CV-3 | Viruses | 0.01 |
| 2031 | 1496.635 | Clostridioides difficile isolate VL_0468 | Bacteria | 0.01 |
| 2032 | 1231072 | Clostridium tetani 12124569 | Bacteria | 0.01 |
| 2033 | 501571.1 | Butyricicoccus | Bacteria | 0.01 |
| 2034 | 877422 | Butyrivibrio hungatei NK4A153 | Bacteria | 0.01 |
| 2035 | 1089447 | Aggregatibacter actinomycetemcomitans RhAA1 | Bacteria | 0.01 |
| 2036 | 113574.1 | Hyphomicrobium sp. GJ21 sp. GJ21 | Bacteria | 0.01 |
| 2037 | 537970 | Helicobacter canadensis MIT 98-5491 | Bacteria | 0.01 |
| 2038 | 936554.1 | Campylobacter sp. FOBRC14 | Bacteria | 0.01 |
| 2039 | 1496.457 | Clostridioides difficile strain VRECD0002 | Bacteria | 0.01 |
| 2040 | 1437607 | Bifidobacterium saguini DSM 23967 | Bacteria | 0.01 |
| 2041 | 932006 | Bacillus subtilis subsp. spizizenii RFWG1A3 | Bacteria | 0.01 |
| 2042 | 1913031 | Papaya ringspot virus-W | Viruses | 0.01 |
| 2043 | 1739479.1 | Actinomyces sp. HMSC065F12 | Bacteria | 0.01 |
| 2044 | 1437600 | Bifidobacterium pullorum DSM 20433 | Bacteria | 0.01 |
| 2045 | 185902 | Human rhinovirus A31 | Viruses | 0.01 |
| 2046 | 185905 | Human rhinovirus A34 | Viruses | 0.01 |
| 2047 | 817.25 | Bacteroides fragilis strain O:21 | Bacteria | 0.01 |

| | | | | |
|---|---|---|---|---|
| 2048 | 1685.4 | Bifidobacterium breve strain BR-I29 | Bacteria | 0.01 |
| 2049 | 1354.1 | Enterococcus hirae strain 19m | Bacteria | 0.01 |
| 2050 | 1681.3 | 2789STDY5608877 | Bacteria | 0.01 |
| 2051 | 303541.1 | Lactobacillus apis strain R-53131 | Bacteria | 0.01 |
| 2052 | 562.78 | Escherichia coli strain VL127 | Bacteria | 0.01 |
| 2053 | 1051661 | Lactobacillus casei UW4 | Bacteria | 0.01 |
| 2054 | 562.77 | Escherichia coli strain AF7658-2 | Bacteria | 0.01 |
| 2055 | 195.1 | Campylobacter coli | Bacteria | 0.01 |
| 2056 | 1923308.1 | Hubei toti-like virus 2 strain arthropodmix13450 | Viruses | 0.01 |
| 2057 | 32629.1 | Indian peanut clump virus | Viruses | 0.01 |
| 2058 | 1123243 | Schwartzia succinivorans DSM 10502 | Bacteria | 0.01 |
| 2059 | 135750 | Tomato bushy stunt virus satellite RNA B10 | Viruses | 0.01 |
| 2060 | 1568973.1 | Botrytis cinerea RNA virus 1 strain BerBc-1 | Viruses | 0.01 |
| 2061 | 1302.23 | Streptococcus gordonii strain G9B | Bacteria | 0.01 |
| 2062 | 1439319.1 | Citrobacter sp. MGH 55 | Bacteria | 0.01 |
| 2063 | 137591.6 | Weissella cibaria strain CH2 | Bacteria | 0.01 |
| 2064 | 1222338.1 | Enterobacteria phage GEC-3S | Viruses | 0.01 |
| 2065 | 1293592 | Lactobacillus curvatus JCM 1096 = DSM 20019 | Bacteria | 0.01 |
| 2066 | 1496.325 | Clostridioides difficile strain VRECD0035 | Bacteria | 0.01 |
| 2067 | 1244083 | Campylobacter showae CSUNSWCD | Bacteria | 0.01 |
| 2068 | 1613.3 | Lactobacillus fermentum strain DSM | Bacteria | 0.01 |
| 2069 | 1423813 | Lactobacillus vaccinostercus DSM 20634 | Bacteria | 0.01 |
| 2070 | 910996 | Hafnia alvei ATCC 13337 | Bacteria | 0.01 |
| 2071 | 1588750.1 | Clostridiales bacterium KA00134 | Bacteria | 0.01 |
| 2072 | 1219360 | Erwinia persicina NBRC 102418 | Bacteria | 0.01 |
| 2073 | 1605.1 | Lactobacillus animalis strain P38 | Bacteria | 0.01 |
| 2074 | 1081904 | Prevotella pleuritidis F0068 | Bacteria | 0.01 |
| 2075 | 291175 | Sapovirus Hu/Dresden/pJG-Sap01/DE | Viruses | 0.01 |
| 2076 | 1283.11 | Staphylococcus haemolyticus strain 1292_SHAE | Bacteria | 0.01 |
| 2077 | 1232440 | Hungatella hathewayi VE202-04 | Bacteria | 0.01 |
| 2078 | 562.134 | Escherichia coli isolate YS | Bacteria | 0.01 |
| 2079 | 1504.1 | Clostridium septicum strain P1044 | Bacteria | 0.01 |
| 2080 | 267135.9 | Porphyrobacter donghaensis strain CCH1-A1 | Bacteria | 0.01 |
| 2081 | 1450518 | Sphingobium lucknowense F2 | Bacteria | 0.01 |
| 2082 | 12196.45 | Bean common mosaic virus isolate 1755a | Viruses | 0.01 |
| 2083 | 1702221.1 | Faecalibaculum rodentium strain NYU-BL-K8 | Bacteria | 0.01 |
| 2084 | 1193182 | Tetrasphaera australiensis Ben110 | Bacteria | 0.01 |
| 2085 | 328061.1 | Radish mosaic virus isolate DH-1 | Viruses | 0.01 |
| 2086 | 439016.1 | Marine RNA virus JP-B | Viruses | 0.01 |
| 2087 | 151043.1 | Tulare apple mosaic virus | Viruses | 0.01 |
| 2088 | 28037.3 | Streptococcus mitis strain SK629 | Bacteria | 0.01 |
| 2089 | 1414721.1 | Clostridium jeddahense strain JCD | Bacteria | 0.01 |
| 2090 | 1715019.1 | Enterococcus sp. HMSC064A12 | Bacteria | 0.01 |
| 2091 | 1242969 | Campylobacter concisus ATCC 51562 | Bacteria | 0.01 |
| 2092 | 92444.3 | Acute bee paralysis virus isolate Hungary | Viruses | 0.01 |
| 2093 | 1739351.1 | Corynebacterium sp. HMSC074C03 | Bacteria | 0.01 |
| 2094 | 267135.1 | Porphyrobacter donghaensis strain CCH7-A10 | Bacteria | 0.01 |
| 2095 | 1120992 | Anaeromusa acidaminophila DSM 3853 | Bacteria | 0.01 |

| | | | | |
|---|---|---|---|---|
| 2096 | 1223528 | Microbacterium oleivorans NBRC 103075 | Bacteria | 0.01 |
| 2097 | 747.4 | Pasteurella multocida strain 306_PMUL | Bacteria | 0.01 |
| 2098 | 1715007.1 | Rothia sp. HMSC071B01 | Bacteria | 0.01 |
| 2099 | 1281072 | Escherichia coli HVH 139 (4-3192644) | Bacteria | 0.01 |
| 2100 | 1739310.1 | Turicella sp. HMSC076G08 | Bacteria | 0.01 |
| 2101 | 1077464.1 | Streptococcus oralis subsp. tigurinus strain DGIIBVI | Bacteria | 0.01 |
| 2102 | 12056.2 | Tobacco necrosis virus D isolate TNV-DP | Viruses | 0.01 |
| 2103 | 1220025.4 | Pokeweed mosaic virus isolate PkMV-NJ | Viruses | 0.01 |
| 2104 | 1235640.1 | Enterobacteria phage M | Viruses | 0.01 |
| 2105 | 857154 | Streptococcus mutans 1ID3 | Bacteria | 0.01 |
| 2106 | 944560 | Actinomyces sp. oral taxon 175 str. F0384 | Bacteria | 0.01 |
| 2107 | 1714265.1 | Klebsiella sp. KGM-IMP216 | Bacteria | 0.01 |
| 2108 | 12235.13 | Cucumber green mottle mosaic virus isolate Ec | Viruses | 0.01 |
| 2109 | 586220 | Leuconostoc mesenteroides subsp. cremoris ATCC 19254 | Bacteria | 0.01 |
| 2110 | 44135 | Human rhinovirus A65 | Viruses | 0.01 |
| 2111 | 1519399.1 | Sewage-associated gemycircularvirus 4 isolate BS3913 | Viruses | 0.01 |
| 2112 | 187978.1 | Peru tomato mosaic virus | Viruses | 0.01 |
| 2113 | 1923604.1 | Wenzhou picorna-like virus 2 strain beimix73672 | Viruses | 0.01 |
| 2114 | 586419.2 | Human cosavirus A strain CMH-N199-11 | Viruses | 0.01 |
| 2115 | 45972.1 | Staphylococcus pasteuri strain 915_SPAS | Bacteria | 0.01 |
| 2116 | 134533.1 | Acinetobacter parvus strain CM11 | Bacteria | 0.01 |
| 2117 | 1318.1 | Streptococcus parasanguinis strain 349_SPAR | Bacteria | 0.01 |
| 2118 | 1318.18 | Streptococcus parasanguinis strain C1A | Bacteria | 0.01 |
| 2119 | 76860.1 | Streptococcus constellatus strain 317_SINT | Bacteria | 0.01 |
| 2120 | 5755.1 | Acanthamoeba castellanii | Eukaryota | 0.01 |
| 2121 | 1410666 | Prevotella brevis P6B11 | Bacteria | 0.01 |
| 2122 | 1610835 | Klebsiella phage vB_KpnP_SU552A | Viruses | 0.01 |
| 2123 | 1679.15 | Bifidobacterium longum subsp. longum strain NCIMB8809 | Bacteria | 0.01 |
| 2124 | 569.3 | Hafnia alvei strain GB001 | Bacteria | 0.01 |
| 2125 | 1321820 | Gemella bergeriae ATCC 700627 | Bacteria | 0.01 |
| 2126 | 630199.1 | Grapevine Syrah virus 1 isolate VF-BR | Viruses | 0.01 |
| 2127 | 1433126.1 | Mucinivorans hirudinis | Bacteria | 0.01 |
| 2128 | 11986.1 | Carnation mottle virus | Viruses | 0.01 |
| 2129 | 1095736 | Streptococcus mitis SK575 | Bacteria | 0.01 |
| 2130 | 1095737 | Streptococcus mitis SK579 | Bacteria | 0.01 |
| 2131 | 1321950 | Clostridium butyricum CWBI1009 | Bacteria | 0.01 |
| 2132 | 12275.3 | Tomato black ring virus | Viruses | 0.01 |
| 2133 | 1598.13 | Lactobacillus reuteri strain MD | Bacteria | 0.01 |
| 2134 | 35350.11 | Apple stem pitting virus strain N | Viruses | 0.01 |
| 2135 | 1768792.1 | Erythrobacter sp. CCH5-A1 | Bacteria | 0.01 |
| 2136 | 1208313 | Piscicoccus intestinalis NBRC 104926 | Bacteria | 0.01 |
| 2137 | 1270.3 | Micrococcus luteus strain O'kane | Bacteria | 0.01 |
| 2138 | 837.8 | Porphyromonas gingivalis strain 84_3 isolate 84_3 | Bacteria | 0.01 |
| 2139 | 908337 | Eremococcus coleocola ACS-139-V-Col8 | Bacteria | 0.01 |

| | | | | |
|---|---|---|---|---|
| 2140 | 1122129 | Jeotgalicoccus psychrophilus DSM 19085 | Bacteria | 0.01 |
| 2141 | 999415 | Eggerthia catenaformis OT 569 = DSM 20559 | Bacteria | 0.01 |
| 2142 | 4896.1 | Schizosaccharomyces pombe | Eukaryota | 0.01 |
| 2143 | 140626.2 | Lachnobacterium bovis strain AE2004 | Bacteria | 0.01 |
| 2144 | 1739400.1 | Corynebacterium sp. HMSC069E04 | Bacteria | 0.01 |
| 2145 | 1258574 | Streptococcus gallolyticus subsp. gallolyticus DSM 16831 | Bacteria | 0.01 |
| 2146 | 1715164.1 | Streptococcus sp. HMSC074F05 | Bacteria | 0.01 |
| 2147 | 255248.1 | Leuconostoc garlicum strain KFRI01 | Bacteria | 0.01 |
| 2148 | 1444120 | Escherichia coli 2-156-04_S3_C1 | Bacteria | 0.01 |
| 2149 | 1129192.1 | Bacillus phage BCP8-2 | Viruses | 0.01 |
| 2150 | 1496.477 | Clostridioides difficile strain VRECD0180 | Bacteria | 0.01 |
| 2151 | 1496.476 | Clostridioides difficile strain VRECD0025 | Bacteria | 0.01 |
| 2152 | 1423759 | Lactobacillus hordei DSM 19519 | Bacteria | 0.01 |
| 2153 | 1423753 | Lactobacillus hammesii DSM 16381 | Bacteria | 0.01 |
| 2154 | 548.55 | Klebsiella aerogenes strain 35003 | Bacteria | 0.01 |
| 2155 | 1353980 | Shigella dysenteriae SD1D | Bacteria | 0.01 |
| 2156 | 1219585.1 | Arcanobacterium sp. S3PF19 | Bacteria | 0.01 |
| 2157 | 736.1 | Haemophilus paraphrohaemolyticus strain CCUG | Bacteria | 0.01 |
| 2158 | 857143 | Streptococcus mutans 11VS1 | Bacteria | 0.01 |
| 2159 | 398513 | Bifidobacterium bifidum NCIMB 41171 | Bacteria | 0.01 |
| 2160 | 1542743.1 | Caribou feces-associated gemycircularvirus | Viruses | 0.01 |
| 2161 | 971.3 | Selenomonas ruminantium strain S137 | Bacteria | 0.01 |
| 2162 | 1255.5 | Pediococcus pentosaceus strain LP28 | Bacteria | 0.01 |
| 2163 | 1255.4 | Pediococcus pentosaceus | Bacteria | 0.01 |
| 2164 | 195.518 | Campylobacter coli isolate M2D2 | Bacteria | 0.01 |
| 2165 | 28141.47 | Cronobacter sakazakii strain MOD1-Md33g | Bacteria | 0.01 |
| 2166 | 12232.13 | Zucchini yellow mosaic virus | Viruses | 0.01 |
| 2167 | 1529.1 | Clostridium cadaveris strain NLAE-zl-G419 | Bacteria | 0.01 |
| 2168 | 1167632 | Staphylococcus vitulinus F1028 | Bacteria | 0.01 |
| 2169 | 563037.1 | Streptococcus sp. M143 | Bacteria | 0.01 |
| 2170 | 438780.1 | Lactobacillus phage phiPYB5 | Viruses | 0.01 |
| 2171 | 1210046 | Janibacter hoylei PVAS-1 | Bacteria | 0.01 |
| 2172 | 195103 | Clostridium perfringens ATCC 13124 | Bacteria | 0.01 |
| 2173 | 1064592 | Naumovozyma castellii CBS 4309 | Eukaryota | 0.01 |
| 2174 | 1652048 | Sapovirus Hu/GI.1/Seoul/ROK62/2013/KOR | Viruses | 0.01 |
| 2175 | 1410674 | Sharpea azabuensis DSM 18934 | Bacteria | 0.01 |
| 2176 | 1410672 | Ruminococcus flavefaciens ND2009 | Bacteria | 0.01 |
| 2177 | 1541211.1 | Cripavirus NB-1/2011/HUN | Viruses | 0.01 |
| 2178 | 29363.2 | Clostridium paraputrificum strain 2789STDY5834955 | Bacteria | 0.01 |
| 2179 | 562.1739 | Escherichia coli strain M8 | Bacteria | 0.01 |
| 2180 | 1269760 | Lactobacillus delbrueckii subsp. lactis CRL581 | Bacteria | 0.01 |
| 2181 | 1051676.1 | Erwinia phage vB_EamM-Y2 | Viruses | 0.01 |
| 2182 | 550.13 | Enterobacter cloacae strain MNCRE12 | Bacteria | 0.01 |
| 2183 | 1795832.1 | Eikenella sp. NML130454 | Bacteria | 0.01 |
| 2184 | 656024.1 | Frankia symbiont of Datisca glomerata | Bacteria | 0.01 |
| 2185 | 1293.1 | Staphylococcus gallinarum strain DSM | Bacteria | 0.01 |

| | | | | |
|---|---|---|---|---|
| 2186 | 1647391.1 | Streptococcus phage APCM01 | Viruses | 0.01 |
| 2187 | 1280697 | Butyrivibrio fibrisolvens AB2020 | Bacteria | 0.01 |
| 2188 | 1280692 | Clostridium cadaveris AGR2141 | Bacteria | 0.01 |
| 2189 | 48296.22 | Acinetobacter pittii strain ABBL074 | Bacteria | 0.01 |
| 2190 | 1236504 | Prevotella histicola JCM 15637 = DNF00424 | Bacteria | 0.01 |
| 2191 | 866773 | Finegoldia magna BVS033A4 | Bacteria | 0.01 |
| 2192 | 1122157 | Laribacter hongkongensis DSM 14985 | Bacteria | 0.01 |
| 2193 | 1335616.1 | Lactobacillus wasatchensis strain WDC04 | Bacteria | 0.01 |
| 2194 | 35288.5 | Grapevine virus A isolate GTR1-1 | Viruses | 0.01 |
| 2195 | 1408894.1 | Red clover cryptic virus 1 isolate IPP_Nemaro | Viruses | 0.01 |
| 2196 | 158787.1 | Bifidobacterium scardovii strain 981_BLON | Bacteria | 0.01 |
| 2197 | 1055192.1 | Comamonas sp. B-9 | Bacteria | 0.01 |
| 2198 | 571.8 | Klebsiella oxytoca strain 2880STDY5682571 | Bacteria | 0.01 |
| 2199 | 419015.3 | Alloscardovia omnicolens strain 476_GVAG | Bacteria | 0.01 |
| 2200 | 419015.4 | Alloscardovia omnicolens strain 350_GVAG | Bacteria | 0.01 |
| 2201 | 1622070.1 | Paenibacillus sp. GM2 sp. GM2 | Bacteria | 0.01 |
| 2202 | 1496.311 | Clostridioides difficile strain CD10 | Bacteria | 0.01 |
| 2203 | 28125.2 | Prevotella bivia strain GED7760C | Bacteria | 0.01 |
| 2204 | 633147 | Olsenella uli DSM 7084 | Bacteria | 0.01 |
| 2205 | 563038.1 | Streptococcus sp. M334 | Bacteria | 0.01 |
| 2206 | 1415765 | Streptococcus mitis 21/39 | Bacteria | 0.01 |
| 2207 | 905067 | Streptococcus parasanguinis F0405 | Bacteria | 0.01 |
| 2208 | 1581143.1 | Arthrobacter sp. HMSC08H08 | Bacteria | 0.01 |
| 2209 | 1692238.1 | Enterobacter sp. FY-07 sp. FY-07 | Bacteria | 0.01 |
| 2210 | 1692238.2 | Enterobacter sp. FY-07 | Bacteria | 0.01 |
| 2211 | 185932 | Human rhinovirus A77 | Viruses | 0.01 |
| 2212 | 1293441.1 | Lysinibacillus contaminans strain DSM | Bacteria | 0.01 |
| 2213 | 947969 | Cellulomonas carbonis T26 | Bacteria | 0.01 |
| 2214 | 1294274 | Streptococcus equinus JB1 | Bacteria | 0.01 |
| 2215 | 1080365 | Pediococcus acidilactici MA18/5M | Bacteria | 0.01 |
| 2216 | 1817674.1 | Geobacillus sp. 8 | Bacteria | 0.01 |
| 2217 | 328430.1 | Chickpea chlorotic stunt virus | Viruses | 0.01 |
| 2218 | 1384081.1 | Veillonella sp. DNF00869 | Bacteria | 0.01 |
| 2219 | 1599.7 | Lactobacillus sakei strain RI-409 | Bacteria | 0.01 |
| 2220 | 307486.1 | Tepidimonas taiwanensis strain MB2 | Bacteria | 0.01 |
| 2221 | 42004.3 | Leek yellow stripe virus isolate AG1 | Viruses | 0.01 |
| 2222 | 1446490 | Escherichia phage FFH2 | Viruses | 0.01 |
| 2223 | 1912598.1 | Cherry associated luteovirus | Viruses | 0.01 |
| 2224 | 1871025.1 | Ndongobacter massiliensis strain Marseille-P3170T sp. Marseille-P3170 | Bacteria | 0.01 |
| 2225 | 222805.6 | Mycobacterium chimaera strain AH16 | Bacteria | 0.01 |
| 2226 | 571.33 | Klebsiella oxytoca strain k142b | Bacteria | 0.01 |
| 2227 | 1748.2 | F3E8 | Bacteria | 0.01 |
| 2228 | 675631 | Desulfurococcus mobilis DSM 2161 | Archaea | 0.01 |
| 2229 | 871203.2 | Caballeronia zhejiangensis strain CEIB | Bacteria | 0.01 |
| 2230 | 941824 | Thermobrachium celere DSM 8682 | Bacteria | 0.01 |
| 2231 | 1280676.1 | Butyrivibrio sp. WCD3002 | Bacteria | 0.01 |
| 2232 | 61647.4 | Pluralibacter gergoviae strain DL84A27 | Bacteria | 0.01 |

| | | | | |
|---|---|---|---|---|
| 2233 | 209529.2 | Aphid lethal paralysis virus isolate ALPV-An | Viruses | 0.01 |
| 2234 | 1873985.1 | Salmonella phage IME207 | Viruses | 0.01 |
| 2235 | 1739536.1 | Corynebacterium sp. HMSC073D01 | Bacteria | 0.01 |
| 2236 | 1402966 | Mycobacterium avium subsp. hominissuis 100 | Bacteria | 0.01 |
| 2237 | 936562.1 | Fusobacterium sp. CM21 | Bacteria | 0.01 |
| 2238 | 1686381.1 | Citrobacter sp. MGH106 | Bacteria | 0.01 |
| 2239 | 28115.1 | Porphyromonas macacae strain COT-192 | Bacteria | 0.01 |
| 2240 | 381764 | Fervidobacterium nodosum Rt17-B1 | Bacteria | 0.01 |
| 2241 | 936572.1 | Selenomonas sp. FOBRC6 | Bacteria | 0.01 |
| 2242 | 1462608.1 | Pseudomonas phage KPP25 | Viruses | 0.01 |
| 2243 | 47770.11 | Lactobacillus crispatus strain C037 | Bacteria | 0.01 |
| 2244 | 47770.12 | Lactobacillus crispatus strain PSS7772C | Bacteria | 0.01 |
| 2245 | 1089548 | Thermicanus aegyptius DSM 12793 | Bacteria | 0.01 |
| 2246 | 544580.12 | Actinomyces oris strain G53E | Bacteria | 0.01 |
| 2247 | 544580.16 | Actinomyces oris strain F4D1 | Bacteria | 0.01 |
| 2248 | 1768771.1 | Streptococcus sp. CCH8-G7 | Bacteria | 0.01 |
| 2249 | 498216 | Lactobacillus casei str. Zhang | Bacteria | 0.01 |
| 2250 | 1581089.1 | Corynebacterium sp. HMSC11E11 | Bacteria | 0.01 |
| 2251 | 1169321.1 | Escherichia sp. KTE114 | Bacteria | 0.01 |
| 2252 | 1739496.1 | Prevotella sp. HMSC069G02 | Bacteria | 0.01 |
| 2253 | 12268.1 | Carnation ringspot virus | Viruses | 0.01 |
| 2254 | 2162.2 | Methanobacterium formicicum | Archaea | 0.01 |
| 2255 | 1545701.1 | Lactobacillus sp. wkB10 | Bacteria | 0.01 |
| 2256 | 1423738 | Lactobacillus dextrinicus DSM 20335 | Bacteria | 0.01 |
| 2257 | 1408226 | Vagococcus lutrae LBD1 | Bacteria | 0.01 |
| 2258 | 12280.2 | Tomato ringspot virus | Viruses | 0.01 |
| 2259 | 78541.1 | Streptococcus phage Sfi11 | Viruses | 0.01 |
| 2260 | 1675603.1 | Citrobacter phage Michonne | Viruses | 0.01 |
| 2261 | 29484.2 | Yersinia frederiksenii strain FCF208 | Bacteria | 0.01 |
| 2262 | 1340495 | Lactobacillus reuteri I5007 | Bacteria | 0.01 |
| 2263 | 1204529 | Salmonella phage SSE121 | Viruses | 0.01 |
| 2264 | 1216979 | Aerococcus urinae NBRC 15544 = CCUG 36881 | Bacteria | 0.01 |
| 2265 | 888055 | Leptotrichia wadei F0279 | Bacteria | 0.01 |
| 2266 | 1029718 | Candidatus Arthromitus sp. SFB-mouse-Japan | Bacteria | 0.01 |
| 2267 | 1481465.1 | Tomato necrotic dwarf virus isolate R | Viruses | 0.01 |
| 2268 | 1318.22 | Streptococcus parasanguinis strain MGH413 | Bacteria | 0.01 |
| 2269 | 553174 | Prevotella melaninogenica ATCC 25845 | Bacteria | 0.01 |
| 2270 | 273525 | Pear black necrotic leaf spot virus | Viruses | 0.01 |
| 2271 | 562.703 | Escherichia coli strain CVM | Bacteria | 0.01 |
| 2272 | 1932006.4 | Chicken associated smacovirus strain RS/BR/2015/2 | Viruses | 0.01 |
| 2273 | 1134687.28 | Klebsiella michiganensis strain 97_38 | Bacteria | 0.01 |
| 2274 | 1512.2 | [Clostridium] symbiosum | Bacteria | 0.01 |
| 2275 | 1280.912 | Staphylococcus aureus strain 01-19 | Bacteria | 0.01 |
| 2276 | 1514105.1 | Erysipelothrix larvae sp. LV19 | Bacteria | 0.01 |
| 2277 | 1138898 | Enterococcus faecium EnGen0001 | Bacteria | 0.01 |
| 2278 | 169292.2 | 1237_CAUR | Bacteria | 0.01 |

| | | | | |
|---|---|---|---|---|
| 2279 | 1529909 | Norovirus GI/Hu/JP/2007/GI.P3_GI.3/Shimizu/KK2866 | Viruses | 0.01 |
| 2280 | 1302863 | Streptococcus cristatus AS 1.3089 | Bacteria | 0.01 |
| 2281 | 1922438.1 | Beihai narna-like virus 11 strain BWBFG39775 | Viruses | 0.01 |
| 2282 | 1590.44 | Lactobacillus plantarum strain FBR4 | Bacteria | 0.01 |
| 2283 | 216816.2 | Bifidobacterium longum strain 379 | Bacteria | 0.01 |
| 2284 | 300715 | Sapovirus Chanthaburi-74/Thailand | Viruses | 0.01 |
| 2285 | 228582.1 | Cereal yellow dwarf virus-RPS | Viruses | 0.01 |
| 2286 | 1122172 | Leptotrichia shahii DSM 19757 | Bacteria | 0.01 |
| 2287 | 33936.1 | Aeribacillus pallidus strain 8m3 | Bacteria | 0.01 |
| 2288 | 646010.2 | Suakwa aphid-borne yellows virus isolate DL76 | Viruses | 0.01 |
| 2289 | 1814960.1 | Streptococcus virus 9874 | Viruses | 0.01 |
| 2290 | 713030.1 | Selenomonas sp. oral taxon 136 strain F0591 | Bacteria | 0.01 |
| 2291 | 2702.1 | Gardnerella vaginalis strain GV37 | Bacteria | 0.01 |
| 2292 | 1423720 | Lactobacillus alimentarius DSM 20249 | Bacteria | 0.01 |
| 2293 | 936594.1 | Lachnoanaerobaculum sp. ICM7 | Bacteria | 0.01 |
| 2294 | 12275.2 | Tomato black ring virus isolate TBRV-Mirs | Viruses | 0.01 |
| 2295 | 550.373 | Enterobacter cloacae strain WCHECl-C4 | Bacteria | 0.01 |
| 2296 | 562973 | Actinomyces viscosus C505 | Bacteria | 0.01 |
| 2297 | 257464.2 | Potato black ringspot virus isolate PRI-Ec | Viruses | 0.01 |
| 2298 | 1673719.1 | Anaerococcus sp. SB3 | Bacteria | 0.01 |
| 2299 | 1825924.1 | Barley virus G isolate Gimje | Viruses | 0.01 |
| 2300 | 33966.1 | Leuconostoc mesenteroides subsp. dextranicum strain LbE15 | Bacteria | 0.01 |
| 2301 | 1158602 | Enterococcus raffinosus ATCC 49464 | Bacteria | 0.01 |
| 2302 | 1358009 | Xanthomonas campestris pv. campestris str. CFBP 5817 | Bacteria | 0.01 |
| 2303 | 1246.1 | Leuconostoc lactis strain WIKIM21 | Bacteria | 0.01 |
| 2304 | 232846.1 | IITR89 | Bacteria | 0.01 |
| 2305 | 1121307 | Clostridium cylindrosporum DSM 605 | Bacteria | 0.01 |
| 2306 | 46256.1 | Weissella hellenica strain R-53116 | Bacteria | 0.01 |
| 2307 | 1770210.1 | Micrococcus sp. CH7 | Bacteria | 0.01 |
| 2308 | 1496.11 | Clostridioides difficile strain VRECD0055 | Bacteria | 0.01 |
| 2309 | 712538.1 | Selenomonas sp. oral taxon 478 | Bacteria | 0.01 |
| 2310 | 1353.4 | Enterococcus gallinarum | Bacteria | 0.01 |
| 2311 | 1410654 | Corynebacterium vitaeruminis Ga6A13 | Bacteria | 0.01 |
| 2312 | 150285.1 | Garlic virus E | Viruses | 0.01 |
| 2313 | 1195085.1 | Cronobacter phage CR5 | Viruses | 0.01 |
| 2314 | 156978.1 | Corynebacterium imitans strain DSM | Bacteria | 0.01 |
| 2315 | 44008.7 | Enterococcus cecorum strain BB-66 | Bacteria | 0.01 |
| 2316 | 44008.6 | Enterococcus cecorum strain CB-32 | Bacteria | 0.01 |
| 2317 | 44008.3 | Enterococcus cecorum strain CL-1 | Bacteria | 0.01 |
| 2318 | 546.23 | Citrobacter freundii strain 804_CKOS | Bacteria | 0.01 |
| 2319 | 1682.3 | Bifidobacterium longum subsp. infantis strain IN-F29 | Bacteria | 0.01 |
| 2320 | 1169330 | Escherichia coli KTE10 | Bacteria | 0.01 |
| 2321 | 928328 | Lactobacillus iners UPII 60-B | Bacteria | 0.01 |
| 2322 | 44562.1 | Pothos latent virus | Viruses | 0.01 |

| | | | | |
|---|---|---|---|---|
| 2323 | 1379702.1 | Methanobacterium sp. MB1 | Archaea | 0.01 |
| 2324 | 1540094.1 | Citrobacter phage Moogle | Viruses | 0.01 |
| 2325 | 129395.2 | Botrytis virus F | Viruses | 0.01 |
| 2326 | 1423726 | Lactobacillus bifermentans DSM 20003 | Bacteria | 0.01 |
| 2327 | 1304.28 | Streptococcus salivarius strain UC3162 | Bacteria | 0.01 |
| 2328 | 1256988 | Providencia alcalifaciens 205/92 | Bacteria | 0.01 |
| 2329 | 550.31 | Enterobacter cloacae strain CB2 | Bacteria | 0.01 |
| 2330 | 31631 | Human coronavirus OC43 | Viruses | 0.01 |
| 2331 | 1567453.1 | Lactobacillus phage LfeSau | Viruses | 0.01 |
| 2332 | 888808 | Streptococcus sanguinis SK49 | Bacteria | 0.01 |
| 2333 | 582.15 | Morganella morganii strain FDAARGOS_172 | Bacteria | 0.01 |
| 2334 | 463676.6 | Rhinovirus C isolate 1570-MY-10 | Viruses | 0.01 |
| 2335 | 388452.1 | Lactococcus phage KSY1 | Viruses | 0.01 |
| 2336 | 2094.1 | Mycoplasma arginini | Bacteria | 0.01 |
| 2337 | 729.2 | Haemophilus parainfluenzae strain ATCC | Bacteria | 0.01 |
| 2338 | 56879.2 | Oat blue dwarf virus | Viruses | 0.01 |
| 2339 | 1264.2 | Ruminococcus albus strain KH2T6 | Bacteria | 0.01 |
| 2340 | 131082.2 | Beet chlorosis virus isolate BChV-CR | Viruses | 0.01 |
| 2341 | 1639.45 | Listeria monocytogenes strain CFSAN026586 | Bacteria | 0.01 |
| 2342 | 1681197.1 | Arthrobacter sp. RIT-PI-e | Bacteria | 0.01 |
| 2343 | 1736702.1 | Enterobacter sp. K66-74 | Bacteria | 0.01 |
| 2344 | 28901.16 | Salmonella enterica strain NGUA31 | Bacteria | 0.01 |
| 2345 | 796937.3 | Peptoanaerobacter stomatis strain ACC19a | Bacteria | 0.01 |
| 2346 | 1612.1 | Lactobacillus farciminis | Bacteria | 0.01 |
| 2347 | 796937.4 | Peptoanaerobacter stomatis strain CM2 | Bacteria | 0.01 |
| 2348 | 1496.438 | Clostridioides difficile strain VRECD0005 | Bacteria | 0.01 |
| 2349 | 1496.434 | Clostridioides difficile strain VRECD0080 | Bacteria | 0.01 |
| 2350 | 29379.1 | Staphylococcus auricularis strain DSM | Bacteria | 0.01 |
| 2351 | 698957 | Gardnerella vaginalis 1500E | Bacteria | 0.01 |
| 2352 | 1302272 | Lactobacillus kimchicus JCM 15530 | Bacteria | 0.01 |
| 2353 | 904296 | Oribacterium sp. oral taxon 108 str. F0425 | Bacteria | 0.01 |
| 2354 | 1423715 | Lactobacillus acidifarinae DSM 19394 | Bacteria | 0.01 |
| 2355 | 373058.1 | Tomato bushy stunt virus satellite RNA | Viruses | 0.01 |
| 2356 | 1218113 | Kluyvera intermedia NBRC 102594 = ATCC 33110 | Bacteria | 0.01 |
| 2357 | 1218112 | Kluyvera cryocrescens NBRC 102467 | Bacteria | 0.01 |
| 2358 | 1280.1861 | Staphylococcus aureus strain C2485 | Bacteria | 0.01 |
| 2359 | 1303256.1 | Sphingobium sp. DC-2 | Bacteria | 0.01 |
| 2360 | 99565 | Chiba virus | Viruses | 0.01 |
| 2361 | 187764.1 | Escherichia virus K1-5 | Viruses | 0.01 |
| 2362 | 698487 | Escherichia phage K1-dep(1) | Viruses | 0.01 |
| 2363 | 1121315 | Terrisporobacter glycolicus ATCC 14880 = DSM 1288 | Bacteria | 0.01 |
| 2364 | 91753.24 | Cucurbit aphid-borne yellows virus isolate CABYV-R-TW82 | Viruses | 0.01 |
| 2365 | 1745712.1 | Anaerococcus sp. Marseille-P2143 strain Marseille-P2143, sp. FC4 | Bacteria | 0.01 |
| 2366 | 329852.1 | Escherichia virus MS2 isolate J20 | Viruses | 0.01 |

| | | | | |
|---|---|---|---|---|
| 2367 | 31770.2 | Shallot virus X strain Russian | Viruses | 0.01 |
| 2368 | 1131442 | Mycobacterium marinum E11 | Bacteria | 0.01 |
| 2369 | 1400823 | Enterococcus faecium UC7256 | Bacteria | 0.01 |
| 2370 | 1028804 | Haemophilus haemolyticus M21127 | Bacteria | 0.01 |
| 2371 | 1028806 | Haemophilus haemolyticus M21639 | Bacteria | 0.01 |
| 2372 | 1115515 | Escherichia vulneris NBRC 102420 | Bacteria | 0.01 |
| 2373 | 1122997 | Acidipropionibacterium jensenii DSM 20535 | Bacteria | 0.01 |
| 2374 | 1239384 | Escherichia phage IME11 | Viruses | 0.01 |
| 2375 | 857292 | Streptococcus intermedius F0395 | Bacteria | 0.01 |
| 2376 | 347253 | Streptococcus salivarius JIM8777 | Bacteria | 0.01 |
| 2377 | 575599 | Lactobacillus fermentum 28-3-CHN | Bacteria | 0.01 |
| 2378 | 1156431.1 | Streptococcus sp. I-G2 sp. I-G2 | Bacteria | 0.01 |
| 2379 | 1661745.1 | Haemophilus sp. C1 | Bacteria | 0.01 |
| 2380 | 12295.8 | Tobacco rattle virus isolate Rostock | Viruses | 0.01 |
| 2381 | 525369 | Proteus mirabilis ATCC 29906 | Bacteria | 0.01 |
| 2382 | 473784.1 | Opium poppy mosaic virus isolate PHEL5235 | Viruses | 0.01 |
| 2383 | 12295.6 | Tobacco rattle virus isolate MI-1 | Viruses | 0.01 |
| 2384 | 11226.4 | Human parainfluenza virus 4b strain QLD-01 | Viruses | 0.01 |
| 2385 | 1008452 | Streptococcus mitis SK1073 | Bacteria | 0.01 |
| 2386 | 1879023.1 | Mycobacterium sp. djl-10 sp. djl-10 | Bacteria | 0.01 |
| 2387 | 76859.1 | Fusobacterium nucleatum subsp. animalis strain KCOM | Bacteria | 0.01 |
| 2388 | 35703.1 | Citrobacter amalonaticus strain L8A | Bacteria | 0.01 |
| 2389 | 1813769.1 | Salmonella phage 64795_sal3 | Viruses | 0.01 |
| 2390 | 244366.24 | Klebsiella variicola isolate T29A | Bacteria | 0.01 |
| 2391 | 1328.2 | Streptococcus anginosus strain 1080_SANG | Bacteria | 0.01 |
| 2392 | 1391428 | Escherichia phage 4MG | Viruses | 0.01 |
| 2393 | 1112212 | Sphingomonas echinoides ATCC 14820 | Bacteria | 0.01 |
| 2394 | 1581067.1 | Kytococcus sp. HMSC28H12 | Bacteria | 0.01 |
| 2395 | 228578.6 | Youcai mosaic virus isolate Br | Viruses | 0.01 |
| 2396 | 889204 | Streptococcus infantis ATCC 700779 | Bacteria | 0.01 |
| 2397 | 338473.1 | Actinomyces virus Av1 | Viruses | 0.01 |
| 2398 | 5082.1 | Penicillium roqueforti strain UASWS | Eukaryota | 0.01 |
| 2399 | 1685.13 | Bifidobacterium breve strain BR-15 | Bacteria | 0.01 |
| 2400 | 1631871.1 | Weissella jogaejeotgali strain FOL01 | Bacteria | 0.01 |
| 2401 | 1423771 | Lactobacillus mucosae DSM 13345 | Bacteria | 0.01 |
| 2402 | 1608993.1 | Pseudomonas sp. DSM 28140 | Bacteria | 0.01 |
| 2403 | 12274.5 | Grapevine fanleaf virus isolate GHu | Viruses | 0.01 |
| 2404 | 646010.1 | Suakwa aphid-borne yellows virus isolate SABYV-TW19 | Viruses | 0.01 |
| 2405 | 1316587 | Fusobacterium nucleatum CTI-6 | Bacteria | 0.01 |
| 2406 | 1073373 | Streptococcus sanguinis CC94A | Bacteria | 0.01 |
| 2407 | 105219.2 | Ralstonia mannitolilytica strain SN82F48 | Bacteria | 0.01 |
| 2408 | 1581069.1 | Corynebacterium sp. HMSC29G08 | Bacteria | 0.01 |
| 2409 | 889206 | Streptococcus vestibularis ATCC 49124 | Bacteria | 0.01 |
| 2410 | 546.15 | Citrobacter freundii strain FDAARGOS_61 | Bacteria | 0.01 |
| 2411 | 28347.3 | Apple stem grooving virus isolate M220 | Viruses | 0.01 |
| 2412 | 1492.2 | Clostridium butyricum strain SU1 | Bacteria | 0.01 |

| | | | | |
|---|---|---|---|---|
| 2413 | 1492.5 | Clostridium butyricum strain JKY6D1 | Bacteria | 0.01 |
| 2414 | 1492.8 | Clostridium butyricum strain CDC_51208 | Bacteria | 0.01 |
| 2415 | 1585.4 | Lactobacillus delbrueckii subsp. bulgaricus strain LBB.B5 | Bacteria | 0.01 |
| 2416 | 864563 | Selenomonas sp. oral taxon 149 str. 67H29BP | Bacteria | 0.01 |
| 2417 | 315405.6 | Streptococcus gallolyticus strain VTM2R47 | Bacteria | 0.01 |
| 2418 | 12169.1 | Potato virus S isolate 89.249 | Viruses | 0.01 |
| 2419 | 1496.31 | Clostridioides difficile strain 106 | Bacteria | 0.01 |
| 2420 | 1122980 | Prevotella baroniae DSM 16972 = JCM 13447 | Bacteria | 0.01 |
| 2421 | 1581075.1 | Neisseria sp. HMSC31F04 | Bacteria | 0.01 |
| 2422 | 546.4 | Citrobacter freundii strain GED7749C | Bacteria | 0.01 |
| 2423 | 10752.1 | Escherichia phage N4 | Viruses | 0.01 |
| 2424 | 104263.2 | Hop latent virus | Viruses | 0.01 |
| 2425 | 1756832.2 | Phasey bean mild yellows virus isolate NSWCP15 | Viruses | 0.01 |
| 2426 | 1303.1 | Streptococcus oralis strain 918_SORA | Bacteria | 0.01 |
| 2427 | 1042417 | Brachyspira pilosicoli P43/6/78 | Bacteria | 0.01 |
| 2428 | 727.66 | Haemophilus influenzae strain 839_HINF | Bacteria | 0.01 |
| 2429 | 1496.406 | Clostridioides difficile strain CD33 | Bacteria | 0.01 |
| 2430 | 1051503 | Bacillus subtilis subsp. spizizenii DV1-B-1 | Bacteria | 0.01 |
| 2431 | 4959.1 | Debaryomyces hansenii | Eukaryota | 0.01 |
| 2432 | 1581072.1 | Corynebacterium sp. HMSC30G07 | Bacteria | 0.01 |
| 2433 | 644284 | Arcanobacterium haemolyticum DSM 20595 | Bacteria | 0.01 |
| 2434 | 1655.5 | Actinomyces naeslundii strain R8152 | Bacteria | 0.01 |
| 2435 | 12197.1 | Bean yellow mosaic virus | Viruses | 0.01 |
| 2436 | 162.1 | Treponema phagedenis | Bacteria | 0.01 |
| 2437 | 1116231 | Streptococcus macedonicus ACA-DC 198 | Bacteria | 0.01 |
| 2438 | 1401659 | Cronobacter sakazakii CMCC 45402 | Bacteria | 0.01 |
| 2439 | 1923725.1 | Wuhan insect virus 21 strain WHCCII13077 | Viruses | 0.01 |
| 2440 | 1206110.1 | Lactobacillus phage phiAQ113 | Viruses | 0.01 |
| 2441 | 576789.1 | Enterobacteria phage JSE | Viruses | 0.01 |
| 2442 | 469599 | Fusobacterium periodonticum 2_1_31 | Bacteria | 0.01 |
| 2443 | 1203573.1 | Propionibacterium sp. KPL1844 | Bacteria | 0.01 |
| 2444 | 28348.1 | Sweet clover necrotic mosaic virus strain 38 | Viruses | 0.01 |
| 2445 | 936140 | Lactobacillus farciminis KCTC 3681 = DSM 20184 | Bacteria | 0.01 |
| 2446 | 1527519.1 | Escherichia phage Av-05 | Viruses | 0.01 |
| 2447 | 235443 | Cryptococcus neoformans var. grubii H99 | Eukaryota | 0.01 |
| 2448 | 1035839 | Haemophilus sputorum CCUG 13788 | Bacteria | 0.01 |
| 2449 | 488537 | Clostridium perfringens D str. JGS1721 | Bacteria | 0.01 |
| 2450 | 562.535 | Escherichia coli strain 444_ECOL | Bacteria | 0.01 |
| 2451 | 673375 | Sodalis phage SO1 | Viruses | 0.01 |
| 2452 | 294746 | Meyerozyma guilliermondii ATCC 6260 | Eukaryota | 0.01 |
| 2453 | 11988.2 | Turnip crinkle virus | Viruses | 0.01 |
| 2454 | 1414738 | Shigella phage pSb-1 | Viruses | 0.01 |
| 2455 | 556267 | Helicobacter winghamensis ATCC BAA-430 | Bacteria | 0.01 |
| 2456 | 1095749 | Pasteurella bettyae CCUG 2042 | Bacteria | 0.01 |
| 2457 | 1381464.1 | Blackberry vein banding associated virus isolate Mississippi1 | Viruses | 0.01 |

| | | | | |
|---|---|---|---|---|
| 2458 | 1496.46 | Clostridioides difficile isolate VL_0218 | Bacteria | 0.01 |
| 2459 | 185891 | Human rhinovirus A9 | Viruses | 0.01 |
| 2460 | 1224748 | Solibacillus isronensis B3W22 | Bacteria | 0.01 |
| 2461 | 185897 | Human rhinovirus A24 | Viruses | 0.01 |
| 2462 | 185896 | Human rhinovirus A22 | Viruses | 0.01 |
| 2463 | 709323.1 | Fructobacillus tropaeoli | Bacteria | 0.01 |
| 2464 | 102684.1 | Streptococcus infantarius strain ICDDRB-NRC-S5 | Bacteria | 0.01 |
| 2465 | 142843.1 | Hop mosaic virus | Viruses | 0.01 |
| 2466 | 45634.5 | Streptococcus cristatus strain JPIIBBV4 | Bacteria | 0.01 |
| 2467 | 1121421 | Desulfotomaculum aeronauticum DSM 10349 | Bacteria | 0.01 |
| 2468 | 12470.1 | Lucerne transient streak virus isolate LTSV-Can | Viruses | 0.01 |
| 2469 | 1580.8 | Lactobacillus brevis strain DmCS_003 | Bacteria | 0.01 |
| 2470 | 1580.9 | Lactobacillus brevis | Bacteria | 0.01 |
| 2471 | 33759 | Citrus tatter leaf virus | Viruses | 0.01 |
| 2472 | 33750 | Hawaii calicivirus | Viruses | 0.01 |
| 2473 | 1768743.1 | Blastomonas sp. CCH8-A3 | Bacteria | 0.01 |
| 2474 | 111970.2 | Kyuri green mottle mosaic virus strain Yodo | Viruses | 0.01 |
| 2475 | 1232427.1 | Corynebacterium ihumii strain GD7 | Bacteria | 0.01 |
| 2476 | 1552735.1 | Lactobacillus phage Ldl1 | Viruses | 0.01 |
| 2477 | 1124962 | Salmonella enterica subsp. enterica serovar Poona str. ATCC BAA-1673 | Bacteria | 0.01 |
| 2478 | 456999.1 | Rhizoctonia solani strain AG3 | Eukaryota | 0.01 |
| 2479 | 1100043.1 | Apis mellifera filamentous virus isolate CH-CO5 | Viruses | 0.01 |
| 2480 | 565655 | Enterococcus casseliflavus EC20 | Bacteria | 0.01 |
| 2481 | 1871034.1 | Propionimicrobium sp. Marseille-P3275 strain Marseille-P3275T | Bacteria | 0.01 |
| 2482 | 1768764.1 | Streptococcus sp. CCH5-D3 | Bacteria | 0.01 |
| 2483 | 1581113.1 | Corynebacterium sp. HMSC05C01 | Bacteria | 0.01 |
| 2484 | 1064535 | Megasphaera elsdenii DSM 20460 | Bacteria | 0.01 |
| 2485 | 1790.3 | Mycobacterium asiaticum strain 1276495.2 | Bacteria | 0.01 |
| 2486 | 644.11 | Aeromonas hydrophila strain AH-1 | Bacteria | 0.01 |
| 2487 | 732.2 | Aggregatibacter aphrophilus strain W10433 | Bacteria | 0.01 |
| 2488 | 1923094.2 | Hubei picorna-like virus 15 strain QTM27139 | Viruses | 0.01 |
| 2489 | 57706.2 | Citrobacter braakii strain GTA-CB04 | Bacteria | 0.01 |
| 2490 | 36343.1 | Lactococcus phage bIL67 | Viruses | 0.01 |
| 2491 | 39804.3 | Escherichia virus FI strain BR8 | Viruses | 0.01 |
| 2492 | 537874.1 | Streptococcus phage PH15 | Viruses | 0.01 |
| 2493 | 39804.5 | Escherichia virus FI strain BR1 | Viruses | 0.01 |
| 2494 | 1123249 | Selenomonas artemidis DSM 19719 | Bacteria | 0.01 |
| 2495 | 327277.1 | Bifidobacterium crudilactis strain LMG | Bacteria | 0.01 |
| 2496 | 1408431 | Butyrivibrio proteoclasticus FD2007 | Bacteria | 0.01 |
| 2497 | 73422.1 | Streptococcus phage TP-J34 | Viruses | 0.01 |
| 2498 | 1444235 | Escherichia coli 2-316-03_S4_C2 | Bacteria | 0.01 |
| 2499 | 71032.3 | Grapevine leafroll-associated virus 5 isolate TRAJ1-BR | Viruses | 0.01 |
| 2500 | 1226633 | Fusobacterium necrophorum subsp. funduliforme B35 | Bacteria | 0.01 |
| 2501 | 936046 | Agaricus bisporus var. bisporus H97 | Eukaryota | 0.01 |

| | | | | |
|---|---|---|---|---|
| 2502 | 1182762.1 | Enterococcus sp. C1 | Bacteria | 0.01 |
| 2503 | 1392849 | Escherichia coli M3 | Bacteria | 0.01 |
| 2504 | 1873990.1 | Escherichia phage vB_EcoM_Alf5 | Viruses | 0.01 |
| 2505 | 1236508 | Prevotella aurantiaca JCM 15754 | Bacteria | 0.01 |
| 2506 | 576791 | Escherichia phage wV8 | Viruses | 0.01 |
| 2507 | 317010.1 | Enterococcus canintestini strain DSM | Bacteria | 0.01 |
| 2508 | 796937.2 | Peptoanaerobacter stomatis strain CM5 | Bacteria | 0.01 |
| 2509 | 546270 | Gemella haemolysans ATCC 10379 | Bacteria | 0.01 |
| 2510 | 12267.1 | Red clover necrotic mosaic virus | Viruses | 0.01 |
| 2511 | 546275 | Fusobacterium periodonticum ATCC 33693 | Bacteria | 0.01 |
| 2512 | 1295140 | Haemophilus influenzae CGSHiCZ412602 | Bacteria | 0.01 |
| 2513 | 1285582 | Lactobacillus sakei subsp. sakei LS25 | Bacteria | 0.01 |
| 2514 | 37961.1 | Atkinsonella hypoxylon virus | Viruses | 0.01 |
| 2515 | 515622 | Butyrivibrio proteoclasticus B316 | Bacteria | 0.01 |
| 2516 | 331679.1 | Pediococcus stilesii strain DSM | Bacteria | 0.01 |
| 2517 | 255238.1 | Fragaria chiloensis latent virus | Viruses | 0.01 |
| 2518 | 550.203 | Enterobacter cloacae strain e1639 | Bacteria | 0.01 |
| 2519 | 12319.5 | Apple mosaic virus | Viruses | 0.01 |
| 2520 | 1823756.1 | Actinomycetaceae bacterium BA112 | Bacteria | 0.01 |
| 2521 | 1768770.1 | Caulobacter sp. CCH5-E12 | Bacteria | 0.01 |
| 2522 | 33964.3 | Leuconostoc citreum | Bacteria | 0.01 |
| 2523 | 1233383.1 | Human cosavirus isolate Cosavirus_Amsterdam_1994 | Viruses | 0.01 |
| 2524 | 1033736.1 | Brevibacterium senegalense sp. JC43 | Bacteria | 0.01 |
| 2525 | 1914546 | Norovirus Hu/USA/2016/GI.P9_GI.9/SC6350 | Viruses | 0.01 |
| 2526 | 563194 | Pediococcus acidilactici 7_4 | Bacteria | 0.01 |
| 2527 | 484020 | Bifidobacterium bifidum BGN4 | Bacteria | 0.01 |
| 2528 | 1505.13 | Paeniclostridium sordellii strain W2922 | Bacteria | 0.01 |
| 2529 | 997353 | Prevotella pallens ATCC 700821 | Bacteria | 0.01 |
| 2530 | 561177 | Anaerococcus hydrogenalis DSM 7454 | Bacteria | 0.01 |
| 2531 | 457403 | Fusobacterium nucleatum subsp. animalis 11_3_2 | Bacteria | 0.01 |
| 2532 | 28375.1 | Soil-borne wheat mosaic virus | Viruses | 0.01 |
| 2533 | 56407.1 | Hanseniaspora occidentalis | Eukaryota | 0.01 |
| 2534 | 167161.6 | Strawberry mottle virus | Viruses | 0.01 |
| 2535 | 167161.2 | Strawberry mottle virus isolate NSper17 | Viruses | 0.01 |
| 2536 | 167161.3 | Strawberry mottle virus isolate NSper51 | Viruses | 0.01 |
| 2537 | 1305618 | Porphyromonas crevioricanis JCM 13913 | Bacteria | 0.01 |
| 2538 | 264483.1 | Phaffia rhodozyma | Eukaryota | 0.01 |
| 2539 | 2371.8 | Xylella fastidiosa strain CFBP8073 | Bacteria | 0.01 |
| 2540 | 1739264.1 | Corynebacterium sp. HMSC065D07 | Bacteria | 0.01 |
| 2541 | 1439318 | Citrobacter freundii MGH 56 | Bacteria | 0.01 |
| 2542 | 451754 | Clostridium perfringens B str. ATCC 3626 | Bacteria | 0.01 |
| 2543 | 451755 | Clostridium perfringens E str. JGS1987 | Bacteria | 0.01 |
| 2544 | 665550.1 | Dietzia alimentaria strain BP | Bacteria | 0.01 |
| 2545 | 68033.1 | Carrot mottle virus | Viruses | 0.01 |
| 2546 | 1161906.1 | Weissella phage phiYS61 | Viruses | 0.01 |
| 2547 | 1329838.1 | Enterobacter sp. BIDMC 26 | Bacteria | 0.01 |
| 2548 | 1451189 | Corynebacterium falsenii DSM 44353 | Bacteria | 0.01 |

| | | | | |
|---|---|---|---|---|
| 2549 | 979982.1 | Leuconostoc sp. C2 sp. C2 | Bacteria | 0.01 |
| 2550 | 590403.1 | Red clover vein mosaic virus isolate NZ | Viruses | 0.01 |
| 2551 | 585.1 | Proteus vulgaris strain CSUR | Bacteria | 0.01 |
| 2552 | 36745.1 | Clostridium saccharoperbutylacetonicum strain N1-504 | Bacteria | 0.01 |
| 2553 | 571.67 | Klebsiella oxytoca strain CHS143 | Bacteria | 0.01 |
| 2554 | 861454 | Lachnospiraceae bacterium oral taxon 082 str. F0431 | Bacteria | 0.01 |
| 2555 | 1444222 | Escherichia coli 3-105-05_S4_C1 | Bacteria | 0.01 |
| 2556 | 74381.1 | Undaria pinnatifida | Eukaryota | 0.01 |
| 2557 | 1505.8 | Paeniclostridium sordellii strain SSCC32135 | Bacteria | 0.01 |
| 2558 | 37206.1 | Helicoverpa armigera stunt virus | Viruses | 0.01 |
| 2559 | 1923266.1 | Hubei tombus-like virus 2 strain WHSFII19265 | Viruses | 0.01 |
| 2560 | 1416754 | Klebsiella michiganensis H1g | Bacteria | 0.01 |
| 2561 | 1171373 | Acidipropionibacterium acidipropionici ATCC 4875 | Bacteria | 0.01 |
| 2562 | 35350.3 | Apple stem pitting virus isolate PR1 | Viruses | 0.01 |
| 2563 | 10829.2 | Squash leaf curl virus | Viruses | 0.01 |
| 2564 | 817.1 | Bacteroides fragilis strain DCMSKEJBY0001B | Bacteria | 0.01 |
| 2565 | 1457184 | Bifidobacterium longum subsp. longum 72B | Bacteria | 0.01 |
| 2566 | 1408287 | Fusobacterium nucleatum W1481 | Bacteria | 0.01 |
| 2567 | 702438 | Prevotella oulorum F0390 | Bacteria | 0.01 |
| 2568 | 264076.1 | Horseradish latent virus | Viruses | 0.01 |
| 2569 | 851.3 | Fusobacterium nucleatum strain MJR7757B | Bacteria | 0.01 |
| 2570 | 1229751.1 | Lactococcus phage BM13 | Viruses | 0.01 |
| 2571 | 1965306.1 | Lagenaria siceraria endornavirus-Hubei isolate JZ | Viruses | 0.01 |
| 2572 | 12055.2 | Tobacco necrosis virus A isolate Velence | Viruses | 0.01 |
| 2573 | 1805471.1 | Clostridium sp. Marseille-P2415 sp. Marseille-P2415 | Bacteria | 0.01 |
| 2574 | 1423822 | Lactobacillus coryniformis subsp. torquens DSM 20004 = KCTC 3535 | Bacteria | 0.01 |
| 2575 | 1050107.1 | Lactobacillus delbrueckii subsp. sunkii strain JCM | Bacteria | 0.01 |
| 2576 | 1441736 | Fusobacterium necrophorum BFTR-2 | Bacteria | 0.01 |
| 2577 | 938293.1 | Anaerococcus provenciensis sp. 9402080 | Bacteria | 0.01 |
| 2578 | 1175299 | Dickeya zeae ZJU1202 | Bacteria | 0.01 |
| 2579 | 1522060.1 | Pantoea sp. 3.5.1 | Bacteria | 0.01 |
| 2580 | 1005705 | Streptococcus infantis SK1076 | Bacteria | 0.01 |
| 2581 | 368736.1 | Maracuja mosaic virus | Viruses | 0.01 |
| 2582 | 1496.7 | Clostridioides difficile isolate VL_0350 | Bacteria | 0.01 |
| 2583 | 439334.34 | Mycobacterium avium subsp. hominissuis strain MAH-P-0913 | Bacteria | 0.01 |
| 2584 | 35841.1 | Bacillus thermoamylovorans strain 1A1 | Bacteria | 0.01 |
| 2585 | 326202.1 | Vanilla distortion mosaic virus isolate VDMV-Cor | Viruses | 0.01 |
| 2586 | 1382301 | Lactobacillus plantarum EGD-AQ4 | Bacteria | 0.01 |
| 2587 | 553190 | Gardnerella vaginalis 409-05 | Bacteria | 0.01 |
| 2588 | 553198 | Propionibacterium acidifaciens F0233 | Bacteria | 0.01 |
| 2589 | 1502.4 | Clostridium perfringens strain CP15 | Bacteria | 0.01 |

| | | | | |
|---|---|---|---|---|
| 2590 | 471872 | Streptococcus infantarius subsp. infantarius ATCC BAA-102 | Bacteria | 0.01 |
| 2591 | 909827.2 | Pepper vein yellows virus isolate 12KNX1 | Viruses | 0.01 |
| 2592 | 699248 | Streptococcus ratti FA-1 = DSM 20564 | Bacteria | 0.01 |
| 2593 | 12230.2 | Turnip mosaic virus | Viruses | 0.01 |
| 2594 | 1134687.39 | Klebsiella michiganensis strain MGH | Bacteria | 0.01 |
| 2595 | 12431.1 | Garlic virus C | Viruses | 0.01 |
| 2596 | 1071395 | Bacillus coagulans XZL4 | Bacteria | 0.01 |
| 2597 | 1778.4 | Mycobacterium gordonae strain 1275229.4 | Bacteria | 0.01 |
| 2598 | 1381091 | Streptococcus equi subsp. zooepidemicus SzAM60 | Bacteria | 0.01 |
| 2599 | 1302.3 | Streptococcus gordonii strain Channon | Bacteria | 0.01 |
| 2600 | 143387.17 | Fusobacterium necrophorum subsp. funduliforme strain F1250 | Bacteria | 0.01 |
| 2601 | 1196034.1 | Klebsiella sp. 10982 | Bacteria | 0.01 |
| 2602 | 1739315.1 | Globicatella sp. HMSC072A10 | Bacteria | 0.01 |
| 2603 | 185893 | Human rhinovirus A13 | Viruses | 0.01 |
| 2604 | 37662.1 | Brettanomyces anomalus | Eukaryota | 0.01 |
| 2605 | 933356 | Enterococcus faecium E4452 | Bacteria | 0.01 |
| 2606 | 883066 | Actinobaculum massiliense ACS-171-V-Col2 | Bacteria | 0.01 |
| 2607 | 1400137 | Citrobacter freundii UCI 32 | Bacteria | 0.01 |
| 2608 | 1122171 | Leptotrichia hofstadii DSM 21651 | Bacteria | 0.01 |
| 2609 | 1675607.1 | Klebsiella phage Matisse | Viruses | 0.01 |
| 2610 | 315405.1 | Streptococcus gallolyticus strain ICDDRB-NRC-S1 | Bacteria | 0.01 |
| 2611 | 562.1112 | Escherichia coli strain upec-274 | Bacteria | 0.01 |
| 2612 | 1051660 | Lactobacillus casei UW1 | Bacteria | 0.01 |
| 2613 | 425279.3 | Rehmannia mosaic virus isolate Shanxi | Viruses | 0.01 |
| 2614 | 28037.19 | Streptococcus mitis strain SVGS_061 | Bacteria | 0.01 |
| 2615 | 28037.15 | Streptococcus mitis strain SK642 | Bacteria | 0.01 |
| 2616 | 28037.16 | Streptococcus mitis strain SK1126 | Bacteria | 0.01 |
| 2617 | 1358413 | Lactobacillus plantarum AY01 | Bacteria | 0.01 |
| 2618 | 131083.2 | Turnip yellows virus isolate WA-1 | Viruses | 0.01 |
| 2619 | 523844 | Methanosarcina thermophila TM-1 | Archaea | 0.01 |
| 2620 | 334413 | Finegoldia magna ATCC 29328 | Bacteria | 0.01 |
| 2621 | 638302 | Selenomonas flueggei ATCC 43531 | Bacteria | 0.01 |
| 2622 | 1509.18 | Clostridium sporogenes strain CDC23284 | Bacteria | 0.01 |
| 2623 | 1870933.1 | Enterobacter cloacae complex sp. 20432 | Bacteria | 0.01 |
| 2624 | 752790 | Escherichia coli CUMT8 | Bacteria | 0.01 |
| 2625 | 1914861.1 | Enterobacter sp. Sa187 | Bacteria | 0.01 |
| 2626 | 1849383.1 | Psychrobacter sp. SHUES1 | Bacteria | 0.01 |
| 2627 | 1932006.3 | Chicken associated smacovirus strain RS/BR/2015/1 | Viruses | 0.01 |
| 2628 | 1123306 | Streptococcus marimammalium DSM 18627 | Bacteria | 0.01 |
| 2629 | 1423828 | Lactobacillus kefiranofaciens subsp. kefirgranum DSM 10550 = JCM 8572 | Bacteria | 0.01 |
| 2630 | 12165.2 | Chrysanthemum virus B isolate Uttarakhand | Viruses | 0.01 |
| 2631 | 253702.1 | Opuntia virus X isolate nopal | Viruses | 0.01 |
| 2632 | 1096935.1 | Veillonella sp. OK1 | Bacteria | 0.01 |

| | | | | |
|---|---|---|---|---|
| 2633 | 1888167.1 | Enterobacter sp. ku-bf2 | Bacteria | 0.01 |
| 2634 | 391774 | Desulfovibrio vulgaris DP4 | Bacteria | 0.01 |
| 2635 | 1325933.1 | Clostridium polynesiense sp. MS1 | Bacteria | 0.01 |
| 2636 | 682148 | Gardnerella vaginalis 5-1 | Bacteria | 0.01 |
| 2637 | 1930302.1 | Chicken stool-associated circular virus strain RS/BR/2015 | Viruses | 0.01 |
| 2638 | 90371.59 | Salmonella enterica subsp. enterica serovar Typhimurium strain CFSAN033859 | Bacteria | 0.01 |
| 2639 | 443906 | Clavibacter michiganensis subsp. michiganensis NCPPB 382 | Bacteria | 0.01 |
| 2640 | 1496.62 | Clostridioides difficile strain CD105KSE9 | Bacteria | 0.01 |
| 2641 | 1613.1 | Lactobacillus fermentum strain 90 | Bacteria | 0.01 |
| 2642 | 445334 | Clostridium perfringens C str. JGS1495 | Bacteria | 0.01 |
| 2643 | 91353.1 | Campylobacter hyointestinalis subsp. lawsonii strain LMG | Bacteria | 0.01 |
| 2644 | 1739611.1 | Lactobacillus phage iLp1308 | Viruses | 0.01 |
| 2645 | 879297 | Lactobacillus iners LactinV 01V1-a | Bacteria | 0.01 |
| 2646 | 1776109.1 | Goose dicistrovirus isolate UW1 | Viruses | 0.01 |
| 2647 | 562.705 | Escherichia coli strain upec-282 | Bacteria | 0.01 |
| 2648 | 1907766.1 | Pseudomonas sp. BS-2016 strain AU14541 | Bacteria | 0.01 |
| 2649 | 796943 | Oribacterium parvum ACB1 | Bacteria | 0.01 |
| 2650 | 1211480.1 | Persimmon virus A | Viruses | 0.01 |
| 2651 | 425010.1 | Botryotinia fuckeliana partitivirus 1 | Viruses | 0.01 |
| 2652 | 1385940 | Bifidobacterium breve JCM 7019 | Bacteria | 0.01 |
| 2653 | 1123721 | Weissella koreensis KCTC 3621 | Bacteria | 0.01 |
| 2654 | 1246.4 | Leuconostoc lactis strain WiKim40 | Bacteria | 0.01 |
| 2655 | 888810 | Streptococcus sanguinis SK115 | Bacteria | 0.01 |
| 2656 | 28037.1 | Streptococcus mitis strain SK608 | Bacteria | 0.01 |
| 2657 | 1118963.2 | Arthrobacter sp. Rue61a | Bacteria | 0.01 |
| 2658 | 33010.7 | Cutibacterium avidum strain DPC | Bacteria | 0.01 |
| 2659 | 1840644.1 | Psophocarpus tetragonolobus endornavirus | Viruses | 0.01 |
| 2660 | 35841.3 | Bacillus thermoamylovorans strain B4064 | Bacteria | 0.01 |
| 2661 | 5412.1 | Cystofilobasidium capitatum | Eukaryota | 0.01 |
| 2662 | 469621 | Fusobacterium periodonticum 1_1_41FAA | Bacteria | 0.01 |
| 2663 | 1141136.1 | Cronobacter phage vB_CsaM_GAP32 | Viruses | 0.01 |
| 2664 | 1640536.1 | Arthrospira sp. TJSD091 | Bacteria | 0.01 |
| 2665 | 1169350.1 | Citrobacter sp. KTE32 | Bacteria | 0.01 |
| 2666 | 571.41 | Klebsiella oxytoca strain 2880STDY5682436 | Bacteria | 0.01 |
| 2667 | 35281.2 | Paprika mild mottle virus | Viruses | 0.01 |
| 2668 | 46076.3 | Artichoke latent virus isolate FR37 | Viruses | 0.01 |
| 2669 | 1280.449 | Staphylococcus aureus strain C2304 | Bacteria | 0.01 |
| 2670 | 28038.7 | Lactobacillus curvatus strain WiKim52 | Bacteria | 0.01 |
| 2671 | 1050903.3 | Pepper cryptic virus 2 isolate HW-01 | Viruses | 0.01 |
| 2672 | 1581133.1 | Actinomyces sp. HMSC08A09 | Bacteria | 0.01 |
| 2673 | 147712.1 | Rhinovirus B strain HRV-B06_p011_sT0384_2007 | Viruses | 0.01 |
| 2674 | 1305.7 | Streptococcus sanguinis strain 2908 | Bacteria | 0.01 |
| 2675 | 1305.6 | Streptococcus sanguinis strain 216_SSAN | Bacteria | 0.01 |

| | | | | |
|---|---|---|---|---|
| 2676 | 861360 | Glutamicibacter arilaitensis Re117 | Bacteria | 0.01 |
| 2677 | 1768759.1 | Bradyrhizobium sp. CCH4-A6 | Bacteria | 0.01 |
| 2678 | 1423792 | Lactobacillus perolens DSM 12744 | Bacteria | 0.01 |
| 2679 | 936563.1 | Fusobacterium sp. CM22 | Bacteria | 0.01 |
| 2680 | 185928 | Human rhinovirus A73 | Viruses | 0.01 |
| 2681 | 1411141 | Serratia ficaria NBRC 102596 | Bacteria | 0.01 |
| 2682 | 1923133.1 | Hubei picorna-like virus 51 strain QTM27291 | Viruses | 0.01 |
| 2683 | 12348.1 | Lactobacillus phage LL-H | Viruses | 0.01 |
| 2684 | 1200547.1 | Prevotella sp. RM4 | Bacteria | 0.01 |
| 2685 | 1121268 | Campylobacter curvus DSM 6644 | Bacteria | 0.01 |
| 2686 | 272633 | Mycoplasma penetrans HF-2 | Bacteria | 0.01 |
| 2687 | 1441735 | Fusobacterium necrophorum DAB | Bacteria | 0.01 |
| 2688 | 910314 | Dialister microaerophilus UPII 345-E | Bacteria | 0.01 |
| 2689 | 1386970 | Clostridium perfringens JJC | Bacteria | 0.01 |
| 2690 | 1599.4 | Lactobacillus sakei strain RI-394 | Bacteria | 0.01 |
| 2691 | 12305.35 | Cucumber mosaic virus isolate Rom | Viruses | 0.01 |
| 2692 | 883067 | Actinotignum schaalii FB123-CNA-2 | Bacteria | 0.01 |
| 2693 | 562.803 | Escherichia coli strain G5 | Bacteria | 0.01 |
| 2694 | 1401064 | Corynebacterium tuscaniense DNF00037 | Bacteria | 0.01 |
| 2695 | 1151428 | Clostridioides difficile P59 | Bacteria | 0.01 |
| 2696 | 1686394.1 | Enterobacter sp. MGH128 | Bacteria | 0.01 |
| 2697 | 525271 | Enterococcus faecalis ATCC 29200 | Bacteria | 0.01 |
| 2698 | 562.937 | Escherichia coli strain upec-34 | Bacteria | 0.01 |
| 2699 | 883119 | Klebsiella oxytoca 10-5244 | Bacteria | 0.01 |
| 2700 | 883114 | Helcococcus kunzii ATCC 51366 | Bacteria | 0.01 |
| 2701 | 132477.1 | Kalanchoe latent virus | Viruses | 0.01 |
| 2702 | 633.1 | Yersinia pseudotuberculosis strain CEB14_0017 | Bacteria | 0.01 |
| 2703 | 888811 | Streptococcus sanguinis SK150 | Bacteria | 0.01 |
| 2704 | 888813 | Streptococcus sanguinis SK330 | Bacteria | 0.01 |
| 2705 | 1406147 | Norovirus Hu/GI.2/Jingzhou/2013401/CHN | Viruses | 0.01 |
| 2706 | 1549858.2 | Sphingomonas taxi strain 30a | Bacteria | 0.01 |
| 2707 | 582.2 | Morganella morganii strain MRSN22709 | Bacteria | 0.01 |
| 2708 | 1923594.2 | Wenzhou picorna-like virus 10 strain WZRBX43164 | Viruses | 0.01 |
| 2709 | 1161421 | Streptococcus oralis SK304 | Bacteria | 0.01 |
| 2710 | 246432.4 | Staphylococcus equorum strain 738_7 | Bacteria | 0.01 |
| 2711 | 656083.1 | Barley yellow striate mosaic virus strain Hebei | Viruses | 0.01 |
| 2712 | 1134805 | Enterococcus faecium 504 | Bacteria | 0.01 |
| 2713 | 12402.1 | Streptococcus phage EJ-1 | Viruses | 0.01 |
| 2714 | 1280703 | Butyrivibrio fibrisolvens MD2001 | Bacteria | 0.01 |
| 2715 | 1280701 | Bifidobacterium pseudolongum AGR2145 | Bacteria | 0.01 |
| 2716 | 1365959 | Bifidobacterium animalis subsp. animalis ATCC 27672 | Bacteria | 0.01 |
| 2717 | 575604 | Lactobacillus gasseri SV-16A-US | Bacteria | 0.01 |
| 2718 | 544580.4 | Actinomyces oris strain R23275 | Bacteria | 0.01 |
| 2719 | 544580.7 | Actinomyces oris strain A19A-1 | Bacteria | 0.01 |
| 2720 | 1770265.1 | Alfalfa enamovirus-1 isolate Manfredi | Viruses | 0.01 |
| 2721 | 1307832 | Tannerella forsythia 3313 | Bacteria | 0.01 |

| | | | | |
|---|---|---|---|---|
| 2722 | 681573 | Streptococcus equi subsp. zooepidemicus BHS5 | Bacteria | 0.01 |
| 2723 | 294.4 | Pseudomonas fluorescens strain ATCC | Bacteria | 0.01 |

Table S3: All species identified in 10,000 human stool samples

| | Taxonomy ID | Species name | SuperKingdom | Prevalence in 10,000 samples, % |
|---|---|---|---|---|
| 1 | 821 | Bacteroides vulgatus | Bacteria | 97.13 |
| 2 | 470 | Acinetobacter baumannii | Bacteria | 96.82 |
| 3 | 853 | Faecalibacterium prausnitzii | Bacteria | 96.37 |
| 4 | 820 | Bacteroides uniformis | Bacteria | 95.38 |
| 5 | 84112 | Eggerthella lenta | Bacteria | 91.76 |
| 6 | 39488 | [Eubacterium] hallii | Bacteria | 91.51 |
| 7 | 169435 | Anaerotruncus colihominis | Bacteria | 91.42 |
| 8 | 1650661 | Clostridium phoceensis | Bacteria | 90.84 |
| 9 | 39778 | Veillonella dispar | Bacteria | 88.99 |
| 10 | 1150298 | Fusicatenibacter saccharivorans | Bacteria | 87.26 |
| 11 | 823 | Parabacteroides distasonis | Bacteria | 87.07 |
| 12 | 817 | Bacteroides fragilis | Bacteria | 85.93 |
| 13 | 214856 | Alistipes finegoldii | Bacteria | 83.35 |
| 14 | 301301 | Roseburia hominis | Bacteria | 83.35 |
| 15 | 40520 | Blautia obeum | Bacteria | 80.81 |
| 16 | 28116 | Bacteroides ovatus | Bacteria | 80.13 |
| 17 | 28117 | Alistipes putredinis | Bacteria | 79.90 |
| 18 | 33039 | [Ruminococcus] torques | Bacteria | 79.50 |
| 19 | 818 | Bacteroides thetaiotaomicron | Bacteria | 79.32 |
| 20 | 1737424 | Blautia massiliensis | Bacteria | 78.86 |
| 21 | 35833 | Bilophila wadsworthia | Bacteria | 78.74 |
| 22 | 28118 | Odoribacter splanchnicus | Bacteria | 78.07 |
| 23 | 1715004 | Clostridiales bacterium KLE1615 | Bacteria | 76.47 |
| 24 | 328814 | Alistipes shahii | Bacteria | 76.45 |
| 25 | 39485 | [Eubacterium] eligens | Bacteria | 76.01 |
| 26 | 88431 | Dorea longicatena | Bacteria | 75.73 |
| 27 | 46503 | Parabacteroides merdae | Bacteria | 75.09 |
| 28 | 39491 | [Eubacterium] rectale | Bacteria | 74.80 |
| 29 | 357276 | Bacteroides dorei | Bacteria | 74.59 |
| 30 | 410072 | Coprococcus comes | Bacteria | 74.49 |
| 31 | 166486 | Roseburia intestinalis | Bacteria | 74.28 |
| 32 | 1519439 | Oscillibacter sp. ER4 | Bacteria | 73.94 |
| 33 | 1522 | [Clostridium] innocuum | Bacteria | 73.77 |
| 34 | 360807 | Roseburia inulinivorans | Bacteria | 73.50 |
| 35 | 214851 | Subdoligranulum variabile | Bacteria | 72.43 |
| 36 | 446660 | Adlercreutzia equolifaciens | Bacteria | 70.41 |
| 37 | 1535 | [Clostridium] leptum | Bacteria | 70.30 |
| 38 | 47678 | Bacteroides caccae | Bacteria | 68.21 |
| 39 | 301302 | Roseburia faecis | Bacteria | 67.03 |
| 40 | 1297617 | Intestinimonas butyriciproducens | Bacteria | 66.95 |
| 41 | 471189 | Gordonibacter pamelaeae | Bacteria | 65.09 |
| 42 | 418240 | Blautia wexlerae | Bacteria | 64.05 |
| 43 | 1697794 | Clostridia bacterium UC5.1-1D1 | Bacteria | 63.33 |
| 44 | 74426 | Collinsella aerofaciens | Bacteria | 63.25 |
| 45 | 39490 | Eubacterium ramulus | Bacteria | 63.11 |

| | | | | |
|---|---|---|---|---|
| 46 | 12239 | Pepper mild mottle virus | Viruses | 62.57 |
| 47 | 1118061 | Alistipes obesi | Bacteria | 62.41 |
| 48 | 1871035 | Ruminococcus sp. Marseille-P3213 | Bacteria | 61.59 |
| 49 | 1352 | Enterococcus faecium | Bacteria | 59.67 |
| 50 | 46506 | Bacteroides stercoris | Bacteria | 59.62 |
| 51 | 1288121 | Alistipes senegalensis | Bacteria | 59.38 |
| 52 | 328813 | Alistipes onderdonkii | Bacteria | 57.51 |
| 53 | 544645 | Butyricimonas virosa | Bacteria | 57.45 |
| 54 | 39486 | Dorea formicigenerans | Bacteria | 56.78 |
| 55 | 204516 | Bacteroides massiliensis | Bacteria | 56.41 |
| 56 | 1917876 | Blautia sp. Marseille-P3087 | Bacteria | 54.65 |
| 57 | 1673721 | Intestinimonas massiliensis | Bacteria | 54.48 |
| 58 | 246787 | Bacteroides cellulosilyticus | Bacteria | 54.15 |
| 59 | 649756 | Anaerostipes hadrus | Bacteria | 54.09 |
| 60 | 39492 | [Eubacterium] siraeum | Bacteria | 53.84 |
| 61 | 552398 | Ruminococcaceae bacterium D16 | Bacteria | 53.81 |
| 62 | 1841867 | Phocea massiliensis | Bacteria | 53.60 |
| 63 | 46228 | Ruminococcus lactaris | Bacteria | 52.95 |
| 64 | 626932 | Alistipes indistinctus | Bacteria | 52.71 |
| 65 | 1232459 | Oscillospiraceae bacterium VE202-24 | Bacteria | 52.69 |
| 66 | 1160721 | Ruminococcus bicirculans | Bacteria | 52.47 |
| 67 | 239935 | Akkermansia muciniphila | Bacteria | 51.28 |
| 68 | 487174 | Barnesiella intestinihominis | Bacteria | 50.92 |
| 69 | 1432052 | Eisenbergiella tayi | Bacteria | 50.61 |
| 70 | 338188 | Bacteroides finegoldii | Bacteria | 50.31 |
| 71 | 329854 | Bacteroides intestinalis | Bacteria | 50.25 |
| 72 | 469610 | Burkholderiales bacterium 1_1_47 | Bacteria | 49.56 |
| 73 | 29348 | [Clostridium] spiroforme | Bacteria | 49.14 |
| 74 | 28052 | Lachnospira pectinoschiza | Bacteria | 48.84 |
| 75 | 84135 | Gemella sanguinis | Bacteria | 48.02 |
| 76 | 39496 | Eubacterium ventriosum | Bacteria | 48.00 |
| 77 | 1095771 | Ruminococcus sp. JC304 | Bacteria | 47.35 |
| 78 | 53443 | Blautia hydrogenotrophica | Bacteria | 43.97 |
| 79 | 665956 | Subdoligranulum sp. 4_3_54A2FAA | Bacteria | 43.55 |
| 80 | 487175 | Parasutterella excrementihominis | Bacteria | 43.55 |
| 81 | 1470347 | Alistipes ihumii | Bacteria | 43.30 |
| 82 | 1703332 | Lachnospiraceae bacterium TF01-11 | Bacteria | 43.21 |
| 83 | 1504823 | bacterium LF-3 | Bacteria | 42.51 |
| 84 | 908612 | Alistipes sp. HGB5 | Bacteria | 42.51 |
| 85 | 292800 | Flavonifractor plautii | Bacteria | 42.43 |
| 86 | 28111 | Bacteroides eggerthii | Bacteria | 42.17 |
| 87 | 1496 | Clostridioides difficile | Bacteria | 41.84 |
| 88 | 371601 | Bacteroides xylanisolvens | Bacteria | 41.60 |
| 89 | 665949 | Tannerella sp. 6_1_58FAA_CT1 | Bacteria | 41.22 |
| 90 | 106588 | Pseudoflavonifractor capillosus | Bacteria | 41.19 |
| 91 | 450746 | Coprobacillus sp. 8_1_38FAA | Bacteria | 40.56 |
| 92 | 658087 | Lachnospiraceae bacterium 7_1_58FAA | Bacteria | 39.20 |
| 93 | 387661 | Parabacteroides johnsonii | Bacteria | 38.96 |

| | | | | |
|---|---|---|---|---|
| 94 | 40545 | Sutterella wadsworthensis | Bacteria | 38.55 |
| 95 | 61171 | Holdemania filiformis | Bacteria | 38.48 |
| 96 | 154046 | Hungatella hathewayi | Bacteria | 38.27 |
| 97 | 1720200 | Anaerotruncus rubiinfantis | Bacteria | 38.19 |
| 98 | 411486 | Clostridium sp. M62/1 | Bacteria | 37.78 |
| 99 | 33043 | Coprococcus eutactus | Bacteria | 37.47 |
| 100 | 1499682 | Alistipes sp. AL-1 | Bacteria | 37.15 |
| 101 | 585543 | Bacteroides sp. D20 | Bacteria | 37.00 |
| 102 | 1232439 | Clostridiales bacterium VE202-03 | Bacteria | 36.73 |
| 103 | 1720194 | Clostridium sp. AT4 | Bacteria | 35.95 |
| 104 | 33038 | [Ruminococcus] gnavus | Bacteria | 35.90 |
| 105 | 1739298 | Bacteroides sp. HMSC067B03 | Bacteria | 33.55 |
| 106 | 1750560 | Parabacteroides sp. SN4 | Bacteria | 33.50 |
| 107 | 1232453 | Clostridiales bacterium VE202-21 | Bacteria | 33.30 |
| 108 | 1232438 | Clostridiales bacterium VE202-01 | Bacteria | 33.14 |
| 109 | 649724 | Clostridium sp. ATCC BAA-442 | Bacteria | 32.80 |
| 110 | 154288 | Turicibacter sanguinis | Bacteria | 32.57 |
| 111 | 1550024 | Ruthenibacterium lactatiformans | Bacteria | 32.08 |
| 112 | 626929 | Bacteroides clarus | Bacteria | 31.85 |
| 113 | 1697793 | Clostridia bacterium UC5.1-1E11 | Bacteria | 31.74 |
| 114 | 742722 | Collinsella sp. 4_8_47FAA | Bacteria | 31.43 |
| 115 | 291645 | Bacteroides nordii | Bacteria | 30.62 |
| 116 | 457412 | Ruminococcus sp. 5_1_39BFAA | Bacteria | 30.13 |
| 117 | 454154 | Paraprevotella clara | Bacteria | 29.87 |
| 118 | 1871020 | Clostridium sp. Marseille-P3244 | Bacteria | 28.76 |
| 119 | 1871018 | Angelakisella massiliensis | Bacteria | 28.62 |
| 120 | 1852384 | Ruminococcaceae bacterium Marseille-P2963 | Bacteria | 28.40 |
| 121 | 1531 | [Clostridium] clostridioforme | Bacteria | 28.28 |
| 122 | 1308 | Streptococcus thermophilus | Bacteria | 28.16 |
| 123 | 55565 | Actinomyces graevenitzii | Bacteria | 27.79 |
| 124 | 454155 | Paraprevotella xylaniphila | Bacteria | 27.73 |
| 125 | 208479 | [Clostridium] bolteae | Bacteria | 27.73 |
| 126 | 1776382 | Neglecta timonensis | Bacteria | 27.46 |
| 127 | 562 | Escherichia coli | Bacteria | 27.40 |
| 128 | 328812 | Parabacteroides goldsteinii | Bacteria | 27.15 |
| 129 | 33035 | Blautia producta | Bacteria | 27.11 |
| 130 | 40519 | Ruminococcus callidus | Bacteria | 27.08 |
| 131 | 1211417 | uncultured phage crAssphage | Viruses | 26.67 |
| 132 | 1870991 | Massilioclostridium coli | Bacteria | 26.41 |
| 133 | 1776384 | Emergencia timonensis | Bacteria | 26.10 |
| 134 | 1470345 | Bacteroides timonensis | Bacteria | 25.96 |
| 135 | 214853 | Anaerofustis stercorihominis | Bacteria | 25.78 |
| 136 | 457389 | Bacteroides sp. 3_1_13 | Bacteria | 25.52 |
| 137 | 1816676 | Alistipes sp. Marseille-P2431 | Bacteria | 25.21 |
| 138 | 1627893 | Ruminococcaceae bacterium cv2 | Bacteria | 25.08 |
| 139 | 1163670 | Bacteroides sp. 14(A) | Bacteria | 25.00 |
| 140 | 310298 | Bacteroides coprocola | Bacteria | 24.97 |
| 141 | 469592 | Bacteroides sp. 3_1_19 | Bacteria | 24.83 |

| | | | | |
|---|---|---|---|---|
| 142 | 291644 | Bacteroides salyersiae | Bacteria | 24.49 |
| 143 | 165179 | Prevotella copri | Bacteria | 24.39 |
| 144 | 39484 | Butyricicoccus desmolans | Bacteria | 24.38 |
| 145 | 658662 | Parabacteroides sp. D26 | Bacteria | 24.36 |
| 146 | 563193 | Parabacteroides sp. D13 | Bacteria | 23.93 |
| 147 | 29347 | [Clostridium] scindens | Bacteria | 23.91 |
| 148 | 45851 | Butyrivibrio crossotus | Bacteria | 23.91 |
| 149 | 1852366 | Holdemania sp. Marseille-P2844 | Bacteria | 23.81 |
| 150 | 1697787 | Clostridia bacterium UC5.1-1D10 | Bacteria | 23.63 |
| 151 | 626940 | Phascolarctobacterium succinatutens | Bacteria | 23.61 |
| 152 | 12241 | Tobacco mild green mosaic virus | Viruses | 23.52 |
| 153 | 160404 | [Clostridium] lactatifermentans | Bacteria | 23.31 |
| 154 | 544644 | Butyricimonas synergistica | Bacteria | 22.96 |
| 155 | 341220 | Lactonifactor longoviformis | Bacteria | 22.74 |
| 156 | 100886 | Catenibacterium mitsuokai | Bacteria | 22.05 |
| 157 | 1735 | Holdemanella biformis | Bacteria | 21.82 |
| 158 | 12253 | Tomato mosaic virus | Viruses | 21.63 |
| 159 | 556259 | Bacteroides sp. D2 | Bacteria | 21.38 |
| 160 | 2173 | Methanobrevibacter smithii | Archaea | 21.15 |
| 161 | 936548 | Actinomyces sp. ICM47 | Bacteria | 20.98 |
| 162 | 1226324 | Blautia sp. KLE 1732 | Bacteria | 20.52 |
| 163 | 457393 | Bacteroides sp. 4_1_36 | Bacteria | 20.25 |
| 164 | 218538 | Dialister invisus | Bacteria | 20.17 |
| 165 | 1473216 | Senegalimassilia anaerobia | Bacteria | 19.57 |
| 166 | 457395 | Bacteroides sp. 9_1_42FAA | Bacteria | 19.06 |
| 167 | 1468449 | Holdemania massiliensis | Bacteria | 18.92 |
| 168 | 411489 | Clostridium sp. L2-50 | Bacteria | 18.88 |
| 169 | 1465754 | Alistipes timonensis | Bacteria | 18.85 |
| 170 | 1720300 | Ruminococcus sp. AT10 | Bacteria | 18.39 |
| 171 | 457394 | Bacteroides sp. 4_3_47FAA | Bacteria | 18.17 |
| 172 | 501571 | Butyricicoccus pullicaecorum | Bacteria | 17.83 |
| 173 | 847 | Oxalobacter formigenes | Bacteria | 17.78 |
| 174 | 693988 | Bilophila sp. 4_1_30 | Bacteria | 17.76 |
| 175 | 1226325 | Clostridium sp. KLE 1755 | Bacteria | 17.72 |
| 176 | 84026 | [Clostridium] methylpentosum | Bacteria | 17.29 |
| 177 | 1574262 | Sutterella sp. KLE1602 | Bacteria | 17.25 |
| 178 | 376806 | Bacteroides gallinarum | Bacteria | 17.21 |
| 179 | 1870993 | Tyzzerella sp. Marseille-P3062 | Bacteria | 17.05 |
| 180 | 1739319 | Bacteroides sp. HMSC068A09 | Bacteria | 16.66 |
| 181 | 438033 | Ruminococcus gauvreauii | Bacteria | 16.59 |
| 182 | 1280669 | Dorea sp. AGR2135 | Bacteria | 16.46 |
| 183 | 1078089 | Bacteroides sp. HPS0048 | Bacteria | 16.45 |
| 184 | 216816 | Bifidobacterium longum | Bacteria | 16.01 |
| 185 | 457390 | Bacteroides sp. 3_1_23 | Bacteria | 15.74 |
| 186 | 270498 | Catabacter hongkongensis | Bacteria | 15.70 |
| 187 | 1232443 | Clostridiales bacterium VE202-13 | Bacteria | 15.56 |
| 188 | 1232457 | Clostridiales bacterium VE202-27 | Bacteria | 15.48 |
| 189 | 1653435 | Clostridium sp. BR31 | Bacteria | 15.43 |

| | | | | |
|---|---|---|---|---|
| 190 | 387090 | Bacteroides coprophilus | Bacteria | 15.29 |
| 191 | 457391 | Bacteroides sp. 3_1_33FAA | Bacteria | 15.06 |
| 192 | 29361 | Tyzzerella nexilis | Bacteria | 14.64 |
| 193 | 112229 | Pepino mosaic virus | Viruses | 14.57 |
| 194 | 658655 | Lachnospiraceae bacterium 1_4_56FAA | Bacteria | 14.54 |
| 195 | 469590 | Bacteroides sp. 2_2_4 | Bacteria | 14.31 |
| 196 | 310297 | Bacteroides plebeius | Bacteria | 14.21 |
| 197 | 105841 | Anaerostipes caccae | Bacteria | 14.20 |
| 198 | 674529 | Bacteroides faecis | Bacteria | 14.13 |
| 199 | 384638 | [Bacteroides] pectinophilus | Bacteria | 14.02 |
| 200 | 457415 | Synergistes sp. 3_1_syn1 | Bacteria | 13.94 |
| 201 | 910311 | Eggerthella sp. HGA1 | Bacteria | 13.94 |
| 202 | 1650663 | Fournierella massiliensis | Bacteria | 13.45 |
| 203 | 363265 | Prevotella stercorea | Bacteria | 13.19 |
| 204 | 871324 | Bacteroides stercorirosoris | Bacteria | 13.17 |
| 205 | 34073 | Variovorax paradoxus | Bacteria | 13.08 |
| 206 | 592978 | Ruminococcus faecis | Bacteria | 12.99 |
| 207 | 1841855 | Bacteroides sp. Marseille-P2653 | Bacteria | 12.97 |
| 208 | 1745713 | Bariatricus massiliensis | Bacteria | 12.65 |
| 209 | 1574263 | Candidatus Stoquefichus sp. KLE1796 | Bacteria | 12.63 |
| 210 | 1099853 | Coprobacter fastidiosus | Bacteria | 12.40 |
| 211 | 469589 | Bacteroides sp. 2_1_33B | Bacteria | 12.40 |
| 212 | 1512 | [Clostridium] symbiosum | Bacteria | 12.26 |
| 213 | 1680 | Bifidobacterium adolescentis | Bacteria | 12.25 |
| 214 | 1907659 | Blautia sp. Marseille-P3201T | Bacteria | 11.90 |
| 215 | 1805476 | Blautia sp. Marseille-P2398 | Bacteria | 11.75 |
| 216 | 1034346 | Dielma fastidiosa | Bacteria | 11.73 |
| 217 | 1852361 | Actinomyces sp. Marseille-P2825 | Bacteria | 11.71 |
| 218 | 28901 | Salmonella enterica | Bacteria | 11.71 |
| 219 | 1796613 | Bacteroides caecimuris | Bacteria | 11.59 |
| 220 | 901 | Desulfovibrio piger | Bacteria | 11.48 |
| 221 | 333367 | [Clostridium] asparagiforme | Bacteria | 11.29 |
| 222 | 1681 | Bifidobacterium bifidum | Bacteria | 11.28 |
| 223 | 35281 | Paprika mild mottle virus | Viruses | 11.23 |
| 224 | 469586 | Bacteroides sp. 1_1_6 | Bacteria | 10.86 |
| 225 | 712888 | Actinobaculum sp. oral taxon 183 | Bacteria | 10.86 |
| 226 | 658659 | Erysipelotrichaceae bacterium 3_1_53 | Bacteria | 10.66 |
| 227 | 12172 | Shallot latent virus | Viruses | 10.54 |
| 228 | 341225 | [Clostridium] saccharogumia | Bacteria | 10.49 |
| 229 | 1161942 | Ruminococcus champanellensis | Bacteria | 10.48 |
| 230 | 147207 | Collinsella intestinalis | Bacteria | 10.36 |
| 231 | 626937 | Christensenella minuta | Bacteria | 10.21 |
| 232 | 469614 | Erysipelotrichaceae bacterium 6_1_45 | Bacteria | 10.21 |
| 233 | 469593 | Bacteroides sp. 3_1_40A | Bacteria | 10.17 |
| 234 | 1309 | Streptococcus mutans | Bacteria | 9.92 |
| 235 | 261299 | Intestinibacter bartlettii | Bacteria | 9.88 |
| 236 | 412467 | Entamoeba nuttalli | Eukaryota | 9.87 |
| 237 | 457387 | Bacteroides sp. 1_1_30 | Bacteria | 9.64 |

| | | | | |
|---|---|---|---|---|
| 238 | 469591 | Parabacteroides sp. 20_3 | Bacteria | 9.45 |
| 239 | 28025 | Bifidobacterium animalis | Bacteria | 9.36 |
| 240 | 626930 | Bacteroides fluxus | Bacteria | 9.23 |
| 241 | 1188792 | Phaseolus vulgaris endornavirus 1 | Viruses | 8.90 |
| 242 | 1852370 | Prevotellamassilia timonensis | Bacteria | 8.84 |
| 243 | 1776379 | Prevotella sp. KHD1 | Bacteria | 8.72 |
| 244 | 1871006 | Bacteroides sp. Marseille-P3132 | Bacteria | 8.71 |
| 245 | 168384 | Marvinbryantia formatexigens | Bacteria | 8.54 |
| 246 | 89153 | [Clostridium] hylemonae | Bacteria | 8.43 |
| 247 | 658089 | Lachnospiraceae bacterium 5_1_63FAA | Bacteria | 8.33 |
| 248 | 1871021 | Lachnoclostridium phocaeense | Bacteria | 8.30 |
| 249 | 1260 | Finegoldia magna | Bacteria | 8.29 |
| 250 | 37733 | Prunus necrotic ringspot virus | Viruses | 8.22 |
| 251 | 876 | Desulfovibrio desulfuricans | Bacteria | 7.97 |
| 252 | 1506471 | Sutterellaceae bacterium ND3 | Bacteria | 7.92 |
| 253 | 143393 | [Eubacterium] sulci | Bacteria | 7.84 |
| 254 | 658086 | Lachnospiraceae bacterium 3_1_57FAA_CT1 | Bacteria | 7.81 |
| 255 | 1841856 | Bacteroides mediterraneensis | Bacteria | 7.77 |
| 256 | 12235 | Cucumber green mottle mosaic virus | Viruses | 7.73 |
| 257 | 626931 | Bacteroides oleiciplenus | Bacteria | 7.73 |
| 258 | 1232444 | Clostridiales bacterium VE202-15 | Bacteria | 7.67 |
| 259 | 574930 | Parabacteroides gordonii | Bacteria | 7.56 |
| 260 | 31971 | [Eubacterium] dolichum | Bacteria | 7.55 |
| 261 | 1318 | Streptococcus parasanguinis | Bacteria | 7.54 |
| 262 | 665950 | Lachnospiraceae bacterium 3_1_46FAA | Bacteria | 7.50 |
| 263 | 36834 | Clostridium celatum | Bacteria | 7.49 |
| 264 | 1852381 | Sutterellaceae bacterium Marseille-P2968 | Bacteria | 7.48 |
| 265 | 457421 | Clostridiales bacterium 1_7_47FAA | Bacteria | 7.42 |
| 266 | 1245 | Leuconostoc mesenteroides | Bacteria | 7.30 |
| 267 | 1903263 | Traorella massiliensis | Bacteria | 7.29 |
| 268 | 1917883 | Bacteroides sp. Marseille-P3166 | Bacteria | 7.16 |
| 269 | 85831 | Bacteroides acidifaciens | Bacteria | 7.04 |
| 270 | 626934 | Slackia piriformis | Bacteria | 6.83 |
| 271 | 28026 | Bifidobacterium pseudocatenulatum | Bacteria | 6.83 |
| 272 | 1834205 | Burkholderiales bacterium YL45 | Bacteria | 6.81 |
| 273 | 84024 | Clostridium disporicum | Bacteria | 6.68 |
| 274 | 1547597 | Sanguibacteroides justesenii | Bacteria | 6.61 |
| 275 | 938289 | Levyella massiliensis | Bacteria | 6.59 |
| 276 | 457397 | Clostridium sp. 1_1_41A1FAA | Bacteria | 6.54 |
| 277 | 1871013 | Parabacteroides sp. Marseille-P3236 | Bacteria | 6.51 |
| 278 | 457402 | Eubacterium sp. 3_1_31 | Bacteria | 6.49 |
| 279 | 1115692 | Cannabis cryptic virus | Viruses | 6.43 |
| 280 | 626933 | Odoribacter laneus | Bacteria | 6.39 |
| 281 | 12968 | Blastocystis hominis | Eukaryota | 6.38 |
| 282 | 1676614 | Prevotella sp. 109 | Bacteria | 6.35 |
| 283 | 102148 | Solobacterium moorei | Bacteria | 6.32 |
| 284 | 342942 | [Clostridium] glycyrrhizinilyticum | Bacteria | 6.25 |
| 285 | 871325 | Bacteroides faecichinchillae | Bacteria | 6.24 |

| | | | | |
|---|---|---|---|---|
| 286 | 47900 | Garlic common latent virus | Viruses | 6.21 |
| 287 | 585544 | Bacteroides sp. D22 | Bacteria | 6.19 |
| 288 | 1602172 | Prevotella sp. P5-125 | Bacteria | 6.14 |
| 289 | 1697795 | Clostridia bacterium UC5.1-2H11 | Bacteria | 6.14 |
| 290 | 186772 | Saccharomyces 20S RNA narnavirus | Viruses | 6.10 |
| 291 | 1499681 | Collinsella sp. MS5 | Bacteria | 6.10 |
| 292 | 76517 | Campylobacter hominis | Bacteria | 5.90 |
| 293 | 1284708 | Tissierellia bacterium S7-1-4 | Bacteria | 5.88 |
| 294 | 1579343 | Streptococcus sp. 263_SSPC | Bacteria | 5.67 |
| 295 | 362693 | Oryza sativa endornavirus | Viruses | 5.65 |
| 296 | 1736 | Eubacterium limosum | Bacteria | 5.62 |
| 297 | 1078087 | Parabacteroides sp. HGS0025 | Bacteria | 5.58 |
| 298 | 39483 | Faecalitalea cylindroides | Bacteria | 5.54 |
| 299 | 358743 | [Clostridium] citroniae | Bacteria | 5.44 |
| 300 | 1816678 | Christensenella timonensis | Bacteria | 5.40 |
| 301 | 658085 | Lachnospiraceae bacterium 5_1_57FAA | Bacteria | 5.36 |
| 302 | 665940 | Clostridium sp. 7_3_54FAA | Bacteria | 5.24 |
| 303 | 1406512 | Candidatus Methanomassiliicoccus intestinalis | Archaea | 5.16 |
| 304 | 944170 | Blastocystis sp. subtype 4 | Eukaryota | 5.08 |
| 305 | 28125 | Prevotella bivia | Bacteria | 4.93 |
| 306 | 1778 | Mycobacterium gordonae | Bacteria | 4.90 |
| 307 | 1658112 | Eubacterium sp. SB2 | Bacteria | 4.85 |
| 308 | 487173 | Dialister succinatiphilus | Bacteria | 4.70 |
| 309 | 1625 | Lactobacillus sanfranciscensis | Bacteria | 4.62 |
| 310 | 290052 | Acetivibrio ethanolgignens | Bacteria | 4.54 |
| 311 | 1304 | Streptococcus salivarius | Bacteria | 4.49 |
| 312 | 1261 | Peptostreptococcus anaerobius | Bacteria | 4.44 |
| 313 | 1391702 | Tomato mottle mosaic virus | Viruses | 4.43 |
| 314 | 33964 | Leuconostoc citreum | Bacteria | 4.38 |
| 315 | 43997 | Catonella morbi | Bacteria | 4.32 |
| 316 | 1519438 | Eubacterium sp. ER2 | Bacteria | 4.30 |
| 317 | 272548 | Actinomyces dentalis | Bacteria | 4.23 |
| 318 | 28128 | Prevotella corporis | Bacteria | 4.18 |
| 319 | 327387 | Tropical soda apple mosaic virus | Viruses | 4.16 |
| 320 | 1761477 | Tomato brown rugose fruit virus | Viruses | 4.09 |
| 321 | 658656 | Lachnospiraceae bacterium 6_1_37FAA | Bacteria | 4.08 |
| 322 | 12175 | Apple chlorotic leaf spot virus | Viruses | 4.07 |
| 323 | 1841865 | Mediterranea massiliensis | Bacteria | 4.05 |
| 324 | 28123 | Porphyromonas asaccharolytica | Bacteria | 4.04 |
| 325 | 1871003 | Tidjanibacter massiliensis | Bacteria | 4.02 |
| 326 | 1671366 | Ruminococcus sp. DSM 100440 | Bacteria | 4.00 |
| 327 | 80879 | Curvibacter delicatus | Bacteria | 3.97 |
| 328 | 936595 | Lachnoanaerobaculum sp. OBRC5-5 | Bacteria | 3.97 |
| 329 | 12305 | Cucumber mosaic virus | Viruses | 3.95 |
| 330 | 658657 | Erysipelotrichaceae bacterium 21_3 | Bacteria | 3.92 |
| 331 | 12242 | Tobacco mosaic virus | Viruses | 3.92 |
| 332 | 376805 | Bacteroides salanitronis | Bacteria | 3.91 |
| 333 | 1723384 | Fenollaria timonensis | Bacteria | 3.88 |

| | | | | |
|---|---|---|---|---|
| 334 | 1871030 | Merdibacter massiliensis | Bacteria | 3.87 |
| 335 | 4932 | Saccharomyces cerevisiae | Eukaryota | 3.87 |
| 336 | 626935 | Collinsella tanakaei | Bacteria | 3.78 |
| 337 | 1501392 | Coprobacter secundus | Bacteria | 3.67 |
| 338 | 1255 | Pediococcus pentosaceus | Bacteria | 3.67 |
| 339 | 747056 | Blueberry shock virus | Viruses | 3.66 |
| 340 | 729 | Haemophilus parainfluenzae | Bacteria | 3.58 |
| 341 | 51330 | Cucurbit yellow stunting disorder virus | Viruses | 3.55 |
| 342 | 712411 | Olsenella sp. oral taxon 807 | Bacteria | 3.54 |
| 343 | 1816694 | Clostridium sp. Marseille-P2538 | Bacteria | 3.47 |
| 344 | 33763 | Peanut mottle virus | Viruses | 3.46 |
| 345 | 397865 | Barnesiella viscericola | Bacteria | 3.45 |
| 346 | 1197717 | Cloacibacillus porcorum | Bacteria | 3.44 |
| 347 | 1602168 | Prevotella sp. P4-65 | Bacteria | 3.43 |
| 348 | 1261635 | Roseburia sp. 831b | Bacteria | 3.28 |
| 349 | 368735 | Bell pepper mottle virus | Viruses | 3.27 |
| 350 | 376804 | Bacteroides barnesiae | Bacteria | 3.27 |
| 351 | 1660 | Actinomyces odontolyticus | Bacteria | 3.27 |
| 352 | 1609975 | Clostridium sp. FS41 | Bacteria | 3.26 |
| 353 | 580026 | Enterorhabdus mucosicola | Bacteria | 3.25 |
| 354 | 1686296 | Gabonia massiliensis | Bacteria | 3.25 |
| 355 | 1903262 | Bacteroides sp. Marseille-P3108 | Bacteria | 3.25 |
| 356 | 1261634 | Roseburia sp. 499 | Bacteria | 3.24 |
| 357 | 47246 | [Clostridium] viride | Bacteria | 3.20 |
| 358 | 28130 | Prevotella disiens | Bacteria | 3.18 |
| 359 | 386414 | Prevotella timonensis | Bacteria | 3.18 |
| 360 | 44742 | Desulfovibrio fairfieldensis | Bacteria | 3.16 |
| 361 | 1265 | Ruminococcus flavefaciens | Bacteria | 3.10 |
| 362 | 1744 | Propionibacterium freudenreichii | Bacteria | 3.07 |
| 363 | 1584 | Lactobacillus delbrueckii | Bacteria | 3.07 |
| 364 | 157777 | Pea streak virus | Viruses | 3.07 |
| 365 | 1491 | Clostridium botulinum | Bacteria | 3.06 |
| 366 | 1322 | Blautia hansenii | Bacteria | 3.03 |
| 367 | 1907658 | Bacteroides sp. Marseille-P3208T | Bacteria | 2.97 |
| 368 | 480391 | Pediococcus argentinicus | Bacteria | 2.97 |
| 369 | 1232460 | Clostridiales bacterium VE202-28 | Bacteria | 2.95 |
| 370 | 311413 | Lettuce big-vein associated virus | Viruses | 2.95 |
| 371 | 712124 | Actinomyces sp. oral taxon 448 | Bacteria | 2.91 |
| 372 | 39482 | [Eubacterium] contortum | Bacteria | 2.89 |
| 373 | 665938 | Bacteroides sp. 2_1_56FAA | Bacteria | 2.89 |
| 374 | 665937 | Anaerostipes sp. 3_2_56FAA | Bacteria | 2.87 |
| 375 | 1583 | Weissella confusa | Bacteria | 2.87 |
| 376 | 1232447 | Clostridiales bacterium VE202-09 | Bacteria | 2.86 |
| 377 | 2317 | Methanosphaera stadtmanae | Archaea | 2.79 |
| 378 | 28139 | Rikenella microfusus | Bacteria | 2.78 |
| 379 | 508460 | Cloacibacillus evryensis | Bacteria | 2.76 |
| 380 | 1870994 | Urmitella timonensis | Bacteria | 2.75 |
| 381 | 457422 | Erysipelotrichaceae bacterium 2_2_44A | Bacteria | 2.73 |

| | | | | |
|---|---|---|---|---|
| 382 | 147206 | Collinsella stercoris | Bacteria | 2.71 |
| 383 | 1232442 | Clostridiales bacterium VE202-06 | Bacteria | 2.71 |
| 384 | 658083 | Lachnospiraceae bacterium 6_1_63FAA | Bacteria | 2.71 |
| 385 | 281920 | Porphyromonas uenonis | Bacteria | 2.68 |
| 386 | 469597 | Coprobacillus sp. 8_2_54BFAA | Bacteria | 2.67 |
| 387 | 1816677 | Butyricimonas sp. Marseille-P2440 | Bacteria | 2.60 |
| 388 | 658088 | Lachnospiraceae bacterium 9_1_43BFAA | Bacteria | 2.53 |
| 389 | 35787 | Lactobacillus pontis | Bacteria | 2.52 |
| 390 | 28347 | Apple stem grooving virus | Viruses | 2.52 |
| 391 | 665941 | Coprobacillus sp. 3_3_56FAA | Bacteria | 2.50 |
| 392 | 944168 | Blastocystis sp. subtype 3 | Eukaryota | 2.49 |
| 393 | 12319 | Apple mosaic virus | Viruses | 2.46 |
| 394 | 491921 | Megamonas rupellensis | Bacteria | 2.43 |
| 395 | 1046403 | Brassica yellows virus | Viruses | 2.41 |
| 396 | 1472761 | Enorma massiliensis | Bacteria | 2.39 |
| 397 | 1574264 | Akkermansia sp. KLE1797 | Bacteria | 2.37 |
| 398 | 1602171 | Prevotella sp. P5-119 | Bacteria | 2.37 |
| 399 | 1235797 | Oscillibacter sp. 1-3 | Bacteria | 2.29 |
| 400 | 1623 | Lactobacillus ruminis | Bacteria | 2.28 |
| 401 | 1852383 | Ruminococcaceae bacterium Marseille-P2935 | Bacteria | 2.28 |
| 402 | 11987 | Melon necrotic spot virus | Viruses | 2.28 |
| 403 | 52227 | Prevotella dentalis | Bacteria | 2.26 |
| 404 | 1599 | Lactobacillus sakei | Bacteria | 2.26 |
| 405 | 501496 | Porphyromonas bennonis | Bacteria | 2.22 |
| 406 | 290053 | Bacteroides helcogenes | Bacteria | 2.15 |
| 407 | 2702 | Gardnerella vaginalis | Bacteria | 2.13 |
| 408 | 1739394 | Porphyromonas sp. HMSC065F10 | Bacteria | 2.13 |
| 409 | 944036 | Blastocystis sp. subtype 1 | Eukaryota | 2.12 |
| 410 | 28127 | Prevotella buccalis | Bacteria | 2.10 |
| 411 | 1796616 | Blautia sp. YL58 | Bacteria | 2.07 |
| 412 | 187327 | Acidaminococcus intestini | Bacteria | 2.04 |
| 413 | 469587 | Bacteroides sp. 2_1_16 | Bacteria | 2.04 |
| 414 | 461393 | Actinomyces massiliensis | Bacteria | 2.03 |
| 415 | 12169 | Potato virus S | Viruses | 2.02 |
| 416 | 91753 | Cucurbit aphid-borne yellows virus | Viruses | 2.02 |
| 417 | 502558 | Eggerthella sp. YY7918 | Bacteria | 2.02 |
| 418 | 671266 | Enterorhabdus caecimuris | Bacteria | 2.01 |
| 419 | 1230734 | Clostridiales bacterium S5-A14a | Bacteria | 2.01 |
| 420 | 1658109 | Candidatus Stoquefichus sp. SB1 | Bacteria | 2.01 |
| 421 | 5722 | Trichomonas vaginalis | Eukaryota | 2.00 |
| 422 | 419015 | Alloscardovia omnicolens | Bacteria | 1.99 |
| 423 | 665942 | Desulfovibrio sp. 6_1_46AFAA | Bacteria | 1.98 |
| 424 | 442302 | Porcine picobirnavirus | Viruses | 1.96 |
| 425 | 1585974 | Beduini massiliensis | Bacteria | 1.96 |
| 426 | 35350 | Apple stem pitting virus | Viruses | 1.95 |
| 427 | 310300 | Bacteroides pyogenes | Bacteria | 1.94 |
| 428 | 1590 | Lactobacillus plantarum | Bacteria | 1.92 |
| 429 | 131111 | Actinomyces turicensis | Bacteria | 1.91 |

| | | | | |
|---|---|---|---|---|
| 430 | 1502 | Clostridium perfringens | Bacteria | 1.90 |
| 431 | 187326 | Megasphaera micronuciformis | Bacteria | 1.89 |
| 432 | 665943 | Eggerthella sp. 1_3_56FAA | Bacteria | 1.88 |
| 433 | 243563 | Strawberry necrotic shock virus | Viruses | 1.84 |
| 434 | 827 | Campylobacter ureolyticus | Bacteria | 1.83 |
| 435 | 1603888 | Megasphaera sp. MJR8396C | Bacteria | 1.83 |
| 436 | 1564113 | Sphingomonas sp. Ant H11 | Bacteria | 1.82 |
| 437 | 1323 | Faecalicoccus pleomorphus | Bacteria | 1.82 |
| 438 | 1903261 | Desulfovibrio sp. Marseille-P3199 | Bacteria | 1.79 |
| 439 | 1611875 | RNA | Viruses | 1.79 |
| 440 | 509923 | Beet cryptic virus 1 | Viruses | 1.79 |
| 441 | 1602169 | Prevotella sp. P4-76 | Bacteria | 1.78 |
| 442 | 1632013 | Drancourtella massiliensis | Bacteria | 1.78 |
| 443 | 1871016 | Collinsella sp. Marseille-P3245 | Bacteria | 1.77 |
| 444 | 187979 | Mitsuokella jalaludinii | Bacteria | 1.77 |
| 445 | 252598 | Saccharomyces sp. 'boulardii' | Eukaryota | 1.76 |
| 446 | 706435 | Capnocytophaga sp. oral taxon 329 | Bacteria | 1.75 |
| 447 | 1078090 | Coprococcus sp. HPP0074 | Bacteria | 1.74 |
| 448 | 1907654 | Collinsella sp. Marseille-P3296T | Bacteria | 1.67 |
| 449 | 1871022 | Libanicoccus massiliensis | Bacteria | 1.67 |
| 450 | 1720313 | Bittarella massiliensis | Bacteria | 1.66 |
| 451 | 1685 | Bifidobacterium breve | Bacteria | 1.66 |
| 452 | 220618 | Cucumber Bulgarian virus | Viruses | 1.63 |
| 453 | 39950 | Dialister pneumosintes | Bacteria | 1.62 |
| 454 | 378833 | Sowbane mosaic virus | Viruses | 1.58 |
| 455 | 1232452 | Clostridiales bacterium VE202-14 | Bacteria | 1.57 |
| 456 | 742723 | Lachnospiraceae bacterium 2_1_46FAA | Bacteria | 1.57 |
| 457 | 1244 | Leuconostoc gelidum | Bacteria | 1.57 |
| 458 | 518643 | Bifidobacterium mongoliense | Bacteria | 1.56 |
| 459 | 28135 | Prevotella oris | Bacteria | 1.56 |
| 460 | 1834196 | Lachnoclostridium sp. YL32 | Bacteria | 1.54 |
| 461 | 1323529 | Dill cryptic virus 2 | Viruses | 1.54 |
| 462 | 1499684 | Clostridium sp. CL-2 | Bacteria | 1.54 |
| 463 | 28038 | Lactobacillus curvatus | Bacteria | 1.53 |
| 464 | 1778580 | Nectarine virus M | Viruses | 1.52 |
| 465 | 755172 | Peptoniphilus coxii | Bacteria | 1.52 |
| 466 | 33760 | Prune dwarf virus | Viruses | 1.52 |
| 467 | 658665 | Dorea sp. D27 | Bacteria | 1.52 |
| 468 | 552396 | Erysipelotrichaceae bacterium 5_2_54FAA | Bacteria | 1.51 |
| 469 | 354328 | Bell pepper endornavirus | Viruses | 1.50 |
| 470 | 1602170 | Prevotella sp. P5-60 | Bacteria | 1.49 |
| 471 | 1714570 | Blueberry shoestring virus | Viruses | 1.48 |
| 472 | 29466 | Veillonella parvula | Bacteria | 1.47 |
| 473 | 1111120 | Acidaminococcus sp. BV3L6 | Bacteria | 1.47 |
| 474 | 165432 | Cucumber leaf spot virus | Viruses | 1.46 |
| 475 | 1598 | Lactobacillus reuteri | Bacteria | 1.42 |
| 476 | 1841857 | Culturomica massiliensis | Bacteria | 1.40 |
| 477 | 47770 | Lactobacillus crispatus | Bacteria | 1.39 |

| | | | | |
|---|---|---|---|---|
| 478 | 76122 | Alloprevotella tannerae | Bacteria | 1.38 |
| 479 | 12321 | Alfalfa mosaic virus | Viruses | 1.37 |
| 480 | 437897 | Megamonas funiformis | Bacteria | 1.35 |
| 481 | 666 | Vibrio cholerae | Bacteria | 1.31 |
| 482 | 1613 | Lactobacillus fermentum | Bacteria | 1.29 |
| 483 | 1739517 | Bacteroides sp. HMSC073E02 | Bacteria | 1.29 |
| 484 | 1105029 | Actinomyces sp. ICM39 | Bacteria | 1.29 |
| 485 | 1917878 | Prevotella ihumii | Bacteria | 1.28 |
| 486 | 12302 | Brome mosaic virus | Viruses | 1.28 |
| 487 | 67761 | Cowpea mild mottle virus | Viruses | 1.27 |
| 488 | 1284775 | Prevotella sp. S7-1-8 | Bacteria | 1.27 |
| 489 | 1470354 | Candidatus Soleaferrea massiliensis | Bacteria | 1.26 |
| 490 | 31741 | Wheat streak mosaic virus | Viruses | 1.26 |
| 491 | 1581179 | Clostridium sp. HMSC19A11 | Bacteria | 1.26 |
| 492 | 1078091 | Coprococcus sp. HPP0048 | Bacteria | 1.25 |
| 493 | 1739408 | Streptococcus sp. HMSC072G04 | Bacteria | 1.24 |
| 494 | 105612 | Lactobacillus algidus | Bacteria | 1.24 |
| 495 | 45634 | Streptococcus cristatus | Bacteria | 1.24 |
| 496 | 1338 | Streptococcus intermedius | Bacteria | 1.24 |
| 497 | 544581 | Actinomyces johnsonii | Bacteria | 1.23 |
| 498 | 341694 | Peptostreptococcus stomatis | Bacteria | 1.21 |
| 499 | 1042156 | Clostridium sp. SY8519 | Bacteria | 1.21 |
| 500 | 850 | Fusobacterium mortiferum | Bacteria | 1.20 |
| 501 | 861 | Fusobacterium ulcerans | Bacteria | 1.20 |
| 502 | 658082 | Lachnospiraceae bacterium 2_1_58FAA | Bacteria | 1.20 |
| 503 | 1739304 | Anaerosphaera sp. HMSC064C01 | Bacteria | 1.19 |
| 504 | 1232428 | Megasphaera massiliensis | Bacteria | 1.19 |
| 505 | 1870997 | Mogibacterium sp. Marseille-P3115 | Bacteria | 1.19 |
| 506 | 1264 | Ruminococcus albus | Bacteria | 1.18 |
| 507 | 52769 | Actinomyces gerencseriae | Bacteria | 1.17 |
| 508 | 1697788 | Clostridia bacterium UC5.1-2H6 | Bacteria | 1.17 |
| 509 | 1261636 | Anaerostipes sp. 494a | Bacteria | 1.16 |
| 510 | 28132 | Prevotella melaninogenica | Bacteria | 1.15 |
| 511 | 28126 | Prevotella buccae | Bacteria | 1.15 |
| 512 | 147802 | Lactobacillus iners | Bacteria | 1.15 |
| 513 | 322095 | Porphyromonas somerae | Bacteria | 1.14 |
| 514 | 1351 | Enterococcus faecalis | Bacteria | 1.13 |
| 515 | 1398 | Bacillus coagulans | Bacteria | 1.10 |
| 516 | 52226 | Mitsuokella multacida | Bacteria | 1.10 |
| 517 | 230143 | Scardovia wiggsiae | Bacteria | 1.10 |
| 518 | 1851429 | Christensenella sp. AF73-05CM02 | Bacteria | 1.09 |
| 519 | 1305 | Streptococcus sanguinis | Bacteria | 1.07 |
| 520 | 31722 | Blueberry scorch virus | Viruses | 1.07 |
| 521 | 242750 | Prevotella bergensis | Bacteria | 1.07 |
| 522 | 137838 | Clostridium neonatale | Bacteria | 1.06 |
| 523 | 12470 | Lucerne transient streak virus | Viruses | 1.06 |
| 524 | 1933298 | Tomato spotted wilt tospovirus | Viruses | 1.05 |
| 525 | 180164 | Blautia schinkii | Bacteria | 1.04 |

| | | | | |
|---|---|---|---|---|
| 526 | 1408895 | Dill cryptic virus 1 | Viruses | 1.04 |
| 527 | 1549 | [Clostridium] sporosphaeroides | Bacteria | 1.03 |
| 528 | 77095 | Prevotella bryantii | Bacteria | 1.02 |
| 529 | 55951 | Grapevine leafroll-associated virus 3 | Viruses | 1.00 |
| 530 | 198589 | Beet western yellows ST9 associated virus | Viruses | 1.00 |
| 531 | 1624 | Lactobacillus salivarius | Bacteria | 0.99 |
| 532 | 1280685 | Butyrivibrio sp. NC3005 | Bacteria | 0.99 |
| 533 | 1364 | Lactococcus piscium | Bacteria | 0.98 |
| 534 | 1834207 | Erysipelotrichaceae bacterium I46 | Bacteria | 0.97 |
| 535 | 457398 | Desulfovibrio sp. 3_1_syn3 | Bacteria | 0.97 |
| 536 | 29391 | Gemella morbillorum | Bacteria | 0.96 |
| 537 | 1307 | Streptococcus suis | Bacteria | 0.95 |
| 538 | 1414720 | Clostridium saudiense | Bacteria | 0.95 |
| 539 | 29363 | Clostridium paraputrificum | Bacteria | 0.94 |
| 540 | 884684 | Mageeibacillus indolicus | Bacteria | 0.94 |
| 541 | 556261 | Clostridium sp. D5 | Bacteria | 0.93 |
| 542 | 907 | Megasphaera elsdenii | Bacteria | 0.91 |
| 543 | 1750 | Pseudopropionibacterium propionicum | Bacteria | 0.89 |
| 544 | 1329795 | Clostridiaceae bacterium MS3 | Bacteria | 0.89 |
| 545 | 228604 | Prevotella salivae | Bacteria | 0.89 |
| 546 | 287 | Pseudomonas aeruginosa | Bacteria | 0.89 |
| 547 | 329 | Ralstonia pickettii | Bacteria | 0.88 |
| 548 | 309120 | Dialister micraerophilus | Bacteria | 0.86 |
| 549 | 1528099 | Lawsonella clevelandensis | Bacteria | 0.86 |
| 550 | 146500 | Watermelon mosaic virus | Viruses | 0.86 |
| 551 | 1111134 | Peptoniphilus sp. BV3C26 | Bacteria | 0.85 |
| 552 | 82135 | Atopobium vaginae | Bacteria | 0.84 |
| 553 | 1267 | Clostridium ventriculi | Bacteria | 0.83 |
| 554 | 1235792 | Lachnospiraceae bacterium M18-1 | Bacteria | 0.82 |
| 555 | 78345 | Bifidobacterium merycicum | Bacteria | 0.82 |
| 556 | 196400 | Grapevine rupestris stem pitting-associated virus | Viruses | 0.82 |
| 557 | 11983 | Norwalk virus | Viruses | 0.80 |
| 558 | 837 | Porphyromonas gingivalis | Bacteria | 0.79 |
| 559 | 905 | Acidaminococcus fermentans | Bacteria | 0.78 |
| 560 | 133926 | Olsenella uli | Bacteria | 0.78 |
| 561 | 999425 | Streptococcus sp. F0442 | Bacteria | 0.77 |
| 562 | 1579 | Lactobacillus acidophilus | Bacteria | 0.76 |
| 563 | 1504822 | bacterium OL-1 | Bacteria | 0.76 |
| 564 | 37734 | Enterococcus casseliflavus | Bacteria | 0.76 |
| 565 | 1797112 | Olsenella sp. kh2p3 | Bacteria | 0.76 |
| 566 | 12162 | Citrus tristeza virus | Viruses | 0.76 |
| 567 | 12042 | Beet western yellows virus | Viruses | 0.75 |
| 568 | 1254 | Pediococcus acidilactici | Bacteria | 0.75 |
| 569 | 1378168 | Firmicutes bacterium ASF500 | Bacteria | 0.75 |
| 570 | 46681 | Entamoeba dispar | Eukaryota | 0.74 |
| 571 | 877411 | Ruminococcus sp. NK3A76 | Bacteria | 0.74 |
| 572 | 1232448 | Clostridiales bacterium VE202-07 | Bacteria | 0.73 |
| 573 | 70177 | Grapevine leafroll-associated virus 4 | Viruses | 0.72 |

| | | | | |
|---|---|---|---|---|
| 574 | 1564903 | Strawberry polerovirus 1 | Viruses | 0.72 |
| 575 | 12282 | Tobacco ringspot virus | Viruses | 0.72 |
| 576 | 425279 | Rehmannia mosaic virus | Viruses | 0.71 |
| 577 | 199 | Campylobacter concisus | Bacteria | 0.70 |
| 578 | 156456 | Anaeroglobus geminatus | Bacteria | 0.70 |
| 579 | 1078480 | Haemophilus sputorum | Bacteria | 0.70 |
| 580 | 732242 | Bacteroides paurosaccharolyticus | Bacteria | 0.70 |
| 581 | 437898 | Sutterella parvirubra | Bacteria | 0.69 |
| 582 | 46125 | Abiotrophia defectiva | Bacteria | 0.69 |
| 583 | 42680 | Spinach latent virus | Viruses | 0.69 |
| 584 | 53442 | Eubacterium callanderi | Bacteria | 0.69 |
| 585 | 1110546 | Veillonella tobetsuensis | Bacteria | 0.68 |
| 586 | 12451 | Raspberry bushy dwarf virus | Viruses | 0.66 |
| 587 | 1776081 | Megasphaera sp. DISK 18 | Bacteria | 0.65 |
| 588 | 1310 | Streptococcus sobrinus | Bacteria | 0.65 |
| 589 | 2748 | Carnobacterium divergens | Bacteria | 0.65 |
| 590 | 1673717 | Anaeromassilibacillus senegalensis | Bacteria | 0.65 |
| 591 | 556499 | Propionibacterium acidifaciens | Bacteria | 0.65 |
| 592 | 1203555 | Acidaminococcus sp. HPA0509 | Bacteria | 0.64 |
| 593 | 12844 | Sweet potato feathery mottle virus | Viruses | 0.64 |
| 594 | 515414 | Prevotella falsenii | Bacteria | 0.64 |
| 595 | 67754 | Tomato chlorosis virus | Viruses | 0.63 |
| 596 | 1588755 | Parvimonas sp. KA00067 | Bacteria | 0.63 |
| 597 | 60920 | Sanguibacter keddieii | Bacteria | 0.62 |
| 598 | 51123 | [Eubacterium] saphenum | Bacteria | 0.62 |
| 599 | 228578 | Youcai mosaic virus | Viruses | 0.62 |
| 600 | 1655 | Actinomyces naeslundii | Bacteria | 0.62 |
| 601 | 419005 | Prevotella amnii | Bacteria | 0.61 |
| 602 | 12280 | Tomato ringspot virus | Viruses | 0.60 |
| 603 | 187101 | Sneathia amnii | Bacteria | 0.60 |
| 604 | 184870 | Varibaculum cambriense | Bacteria | 0.59 |
| 605 | 39029 | Megasphaera cerevisiae | Bacteria | 0.59 |
| 606 | 1577792 | Terrisporobacter othiniensis | Bacteria | 0.59 |
| 607 | 1173061 | Geotrichum candidum | Eukaryota | 0.59 |
| 608 | 155892 | Caulobacter vibrioides | Bacteria | 0.58 |
| 609 | 1261637 | Anaerostipes sp. 992a | Bacteria | 0.58 |
| 610 | 12317 | Tobacco streak virus | Viruses | 0.57 |
| 611 | 61592 | Corynebacterium durum | Bacteria | 0.57 |
| 612 | 712117 | Actinomyces sp. oral taxon 170 | Bacteria | 0.57 |
| 613 | 89152 | [Clostridium] hiranonis | Bacteria | 0.56 |
| 614 | 1871336 | Criibacterium bergeronii | Bacteria | 0.56 |
| 615 | 72149 | Aichivirus A | Viruses | 0.55 |
| 616 | 12433 | Garlic virus A | Viruses | 0.55 |
| 617 | 12187 | Strawberry mild yellow edge virus | Viruses | 0.55 |
| 618 | 1263547 | Oscillibacter ruminantium | Bacteria | 0.55 |
| 619 | 477666 | Bacteroides graminisolvens | Bacteria | 0.55 |
| 620 | 712118 | Actinomyces sp. oral taxon 172 | Bacteria | 0.55 |
| 621 | 1720195 | Gabonibacter massiliensis | Bacteria | 0.54 |

| | | | | |
|---|---|---|---|---|
| 622 | 1605 | Lactobacillus animalis | Bacteria | 0.53 |
| 623 | 33032 | Anaerococcus lactolyticus | Bacteria | 0.53 |
| 624 | 12216 | Potato virus Y | Viruses | 0.52 |
| 625 | 1105030 | Actinomyces sp. ICM58 | Bacteria | 0.52 |
| 626 | 1393034 | Atopobium deltae | Bacteria | 0.51 |
| 627 | 1532180 | Penicillium roqueforti ssRNA mycovirus 1 | Viruses | 0.50 |
| 628 | 1697790 | Clostridia bacterium UC5.1-1C12 | Bacteria | 0.50 |
| 629 | 39441 | Methanobrevibacter arboriphilus | Archaea | 0.50 |
| 630 | 1051631 | Streptococcus phage YMC-2011 | Viruses | 0.50 |
| 631 | 1805478 | Olsenella sp. Marseille-P2300 | Bacteria | 0.49 |
| 632 | 1852386 | Olsenella sp. Marseille-P2912 | Bacteria | 0.49 |
| 633 | 1111135 | Coriobacteriaceae bacterium BV3Ac1 | Bacteria | 0.49 |
| 634 | 112227 | Cactus virus X | Viruses | 0.48 |
| 635 | 556270 | Coprobacillus sp. D7 | Bacteria | 0.48 |
| 636 | 877414 | Clostridiales bacterium NK3B98 | Bacteria | 0.48 |
| 637 | 1697792 | Clostridia bacterium UC5.1-2F7 | Bacteria | 0.48 |
| 638 | 1937817 | Spinach cryptic virus 1 | Viruses | 0.48 |
| 639 | 35518 | [Eubacterium] nodatum | Bacteria | 0.48 |
| 640 | 1739529 | Porphyromonas sp. HMSC077F02 | Bacteria | 0.48 |
| 641 | 1561 | Clostridium baratii | Bacteria | 0.48 |
| 642 | 1232446 | Clostridiales bacterium VE202-18 | Bacteria | 0.47 |
| 643 | 1531429 | Coriobacteriaceae bacterium 68-1-3 | Bacteria | 0.47 |
| 644 | 237576 | Oribacterium sinus | Bacteria | 0.47 |
| 645 | 642478 | Lettuce chlorosis virus | Viruses | 0.47 |
| 646 | 1852374 | Ezakiella massiliensis | Bacteria | 0.46 |
| 647 | 181487 | Actinomyces cardiffensis | Bacteria | 0.46 |
| 648 | 824 | Campylobacter gracilis | Bacteria | 0.46 |
| 649 | 66200 | Carrot red leaf virus | Viruses | 0.46 |
| 650 | 39777 | Veillonella atypica | Bacteria | 0.45 |
| 651 | 12263 | Squash mosaic virus | Viruses | 0.45 |
| 652 | 1673723 | Murdochiella massiliensis | Bacteria | 0.45 |
| 653 | 580254 | Raphanus sativus cryptic virus 3 | Viruses | 0.45 |
| 654 | 1630 | Kandleria vitulina | Bacteria | 0.45 |
| 655 | 12205 | Papaya ringspot virus | Viruses | 0.44 |
| 656 | 290055 | [Eubacterium] fissicatena | Bacteria | 0.44 |
| 657 | 1392494 | Lachnospiraceae bacterium AC2012 | Bacteria | 0.43 |
| 658 | 1470350 | Candidatus Stoquefichus massiliensis | Bacteria | 0.42 |
| 659 | 54005 | Peptoniphilus harei | Bacteria | 0.42 |
| 660 | 431317 | Triticum mosaic virus | Viruses | 0.42 |
| 661 | 1226323 | Oscillibacter sp. KLE 1745 | Bacteria | 0.42 |
| 662 | 544580 | Actinomyces oris | Bacteria | 0.42 |
| 663 | 1392836 | Lachnospiraceae bacterium TWA4 | Bacteria | 0.42 |
| 664 | 727 | Haemophilus influenzae | Bacteria | 0.42 |
| 665 | 1232445 | Clostridiales bacterium VE202-16 | Bacteria | 0.41 |
| 666 | 1601 | Lactobacillus agilis | Bacteria | 0.41 |
| 667 | 1595998 | Human smacovirus 1 | Viruses | 0.41 |
| 668 | 28037 | Streptococcus mitis | Bacteria | 0.41 |
| 669 | 1489836 | Atlantic salmon calicivirus | Viruses | 0.40 |

| | | | | |
|---|---|---|---|---|
| 670 | 52229 | [Hallella] seregens | Bacteria | 0.40 |
| 671 | 712991 | Lachnospiraceae bacterium oral taxon 500 | Bacteria | 0.40 |
| 672 | 10641 | Cauliflower mosaic virus | Viruses | 0.40 |
| 673 | 1852375 | Acidaminococcus massiliensis | Bacteria | 0.39 |
| 674 | 1465757 | Peptoniphilus senegalensis | Bacteria | 0.39 |
| 675 | 5007 | Brettanomyces bruxellensis | Eukaryota | 0.39 |
| 676 | 1479 | Bacillus smithii | Bacteria | 0.39 |
| 677 | 76875 | Broad bean wilt virus 2 | Viruses | 0.39 |
| 678 | 1689 | Bifidobacterium dentium | Bacteria | 0.39 |
| 679 | 29322 | [Eubacterium] cellulosolvens | Bacteria | 0.39 |
| 680 | 1776177 | Cucumis melo endornavirus | Viruses | 0.39 |
| 681 | 322505 | Sharpea azabuensis | Bacteria | 0.38 |
| 682 | 736 | Haemophilus paraphrohaemolyticus | Bacteria | 0.38 |
| 683 | 457396 | Clostridium sp. 7_2_43FAA | Bacteria | 0.37 |
| 684 | 1683 | Bifidobacterium angulatum | Bacteria | 0.37 |
| 685 | 103722 | Grapevine fleck virus | Viruses | 0.37 |
| 686 | 1382 | Atopobium parvulum | Bacteria | 0.36 |
| 687 | 1581061 | Abiotrophia sp. HMSC24B09 | Bacteria | 0.36 |
| 688 | 739 | Aggregatibacter segnis | Bacteria | 0.36 |
| 689 | 467210 | Lachnoanaerobaculum saburreum | Bacteria | 0.35 |
| 690 | 859 | Fusobacterium necrophorum | Bacteria | 0.35 |
| 691 | 95342 | Sapporo virus | Viruses | 0.35 |
| 692 | 210 | Helicobacter pylori | Bacteria | 0.35 |
| 693 | 397544 | Squash vein yellowing virus | Viruses | 0.35 |
| 694 | 56774 | [Eubacterium] infirmum | Bacteria | 0.35 |
| 695 | 84163 | Cryptobacterium curtum | Bacteria | 0.35 |
| 696 | 1499683 | Clostridium sp. CL-6 | Bacteria | 0.35 |
| 697 | 1676617 | Ralstonia sp. MD27 | Bacteria | 0.35 |
| 698 | 310514 | Prevotella multisaccharivorax | Bacteria | 0.34 |
| 699 | 12274 | Grapevine fanleaf virus | Viruses | 0.34 |
| 700 | 908340 | Clostridium sp. HGF2 | Bacteria | 0.34 |
| 701 | 35517 | [Eubacterium] brachy | Bacteria | 0.34 |
| 702 | 638849 | Pyramidobacter piscolens | Bacteria | 0.33 |
| 703 | 28051 | Lachnospira multipara | Bacteria | 0.33 |
| 704 | 145856 | Human picobirnavirus | Viruses | 0.33 |
| 705 | 113107 | Streptococcus australis | Bacteria | 0.33 |
| 706 | 29360 | Cellulosilyticum lentocellum | Bacteria | 0.32 |
| 707 | 47671 | Lautropia mirabilis | Bacteria | 0.32 |
| 708 | 45254 | Dysgonomonas capnocytophagoides | Bacteria | 0.31 |
| 709 | 12056 | Tobacco necrosis virus D | Viruses | 0.31 |
| 710 | 1219626 | Peptostreptococcus sp. MV1 | Bacteria | 0.31 |
| 711 | 1284680 | Actinomyces sp. S6-Spd3 | Bacteria | 0.31 |
| 712 | 39046 | Cassava common mosaic virus | Viruses | 0.31 |
| 713 | 209529 | Aphid lethal paralysis virus | Viruses | 0.31 |
| 714 | 138595 | Olsenella profusa | Bacteria | 0.31 |
| 715 | 137591 | Weissella cibaria | Bacteria | 0.31 |
| 716 | 33034 | Anaerococcus prevotii | Bacteria | 0.31 |
| 717 | 12615 | Cherry leaf roll virus | Viruses | 0.31 |

| | | | | |
|---|---|---|---|---|
| 718 | 796942 | Stomatobaculum longum | Bacteria | 0.31 |
| 719 | 1366 | Lactococcus raffinolactis | Bacteria | 0.31 |
| 720 | 1776390 | Peptoniphilus sp. KHD5 | Bacteria | 0.30 |
| 721 | 163665 | Dysgonomonas mossii | Bacteria | 0.30 |
| 722 | 97478 | Lactobacillus mucosae | Bacteria | 0.30 |
| 723 | 12196 | Bean common mosaic virus | Viruses | 0.30 |
| 724 | 231049 | Lactobacillus rossiae | Bacteria | 0.30 |
| 725 | 78258 | Parascardovia denticolens | Bacteria | 0.29 |
| 726 | 626938 | Succinatimonas hippei | Bacteria | 0.29 |
| 727 | 1871012 | Mobilibacterium timonense | Bacteria | 0.29 |
| 728 | 32625 | Mushroom bacilliform virus | Viruses | 0.29 |
| 729 | 1686 | Bifidobacterium catenulatum | Bacteria | 0.29 |
| 730 | 193118 | Lucerne transient streak virus satellite RNA | Viruses | 0.29 |
| 731 | 1739305 | Peptoniphilus sp. HMSC062D09 | Bacteria | 0.28 |
| 732 | 43768 | Corynebacterium matruchotii | Bacteria | 0.28 |
| 733 | 558690 | Cucurbit chlorotic yellows virus | Viruses | 0.28 |
| 734 | 1252 | Leuconostoc carnosum | Bacteria | 0.28 |
| 735 | 1689303 | Lagierella massiliensis | Bacteria | 0.28 |
| 736 | 1379 | Gemella haemolysans | Bacteria | 0.27 |
| 737 | 563031 | Prevotella sp. C561 | Bacteria | 0.27 |
| 738 | 1588754 | Veillonellaceae bacterium DNF00626 | Bacteria | 0.27 |
| 739 | 12436 | Cucumber mosaic virus satellite RNA | Viruses | 0.27 |
| 740 | 82688 | Lactobacillus nagelii | Bacteria | 0.27 |
| 741 | 930881 | Sweet potato virus C | Viruses | 0.27 |
| 742 | 36841 | Terrisporobacter glycolicus | Bacteria | 0.26 |
| 743 | 67962 | Carrot red leaf luteovirus associated RNA | Viruses | 0.26 |
| 744 | 42004 | Leek yellow stripe virus | Viruses | 0.26 |
| 745 | 131083 | Turnip yellows virus | Viruses | 0.26 |
| 746 | 1871017 | Peptoniphilus urinimassiliensis | Bacteria | 0.26 |
| 747 | 1596 | Lactobacillus gasseri | Bacteria | 0.26 |
| 748 | 54062 | Pediococcus parvulus | Bacteria | 0.26 |
| 749 | 198599 | Saccharomyces 23S RNA narnavirus | Viruses | 0.26 |
| 750 | 42478 | Saccharomyces cerevisiae virus L-BC (La) | Viruses | 0.26 |
| 751 | 40543 | Sneathia sanguinegens | Bacteria | 0.25 |
| 752 | 1449897 | Uncultured phage WW-nAnB strain 3 | Viruses | 0.25 |
| 753 | 1852387 | Streptococcus timonensis | Bacteria | 0.25 |
| 754 | 78343 | Bifidobacterium boum | Bacteria | 0.25 |
| 755 | 1169032 | Wasabi mottle virus | Viruses | 0.25 |
| 756 | 29833 | Hanseniaspora uvarum | Eukaryota | 0.25 |
| 757 | 1302 | Streptococcus gordonii | Bacteria | 0.25 |
| 758 | 158847 | Megamonas hypermegale | Bacteria | 0.25 |
| 759 | 1659 | Actinomyces israelii | Bacteria | 0.25 |
| 760 | 1301220 | Citrus vein enation virus | Viruses | 0.25 |
| 761 | 150284 | Garlic virus X | Viruses | 0.25 |
| 762 | 851 | Fusobacterium nucleatum | Bacteria | 0.25 |
| 763 | 1502943 | Veillonella seminalis | Bacteria | 0.25 |
| 764 | 42882 | Cherry virus A | Viruses | 0.25 |
| 765 | 450749 | Veillonella sp. 6_1_27 | Bacteria | 0.25 |

| | | | | |
|---|---|---|---|---|
| 766 | 12295 | Tobacco rattle virus | Viruses | 0.25 |
| 767 | 180332 | Robinsoniella peoriensis | Bacteria | 0.25 |
| 768 | 244362 | Ruminococcus sp. YE71 | Bacteria | 0.25 |
| 769 | 1805477 | Clostridium sp. Marseille-P299 | Bacteria | 0.24 |
| 770 | 12232 | Zucchini yellow mosaic virus | Viruses | 0.24 |
| 771 | 109790 | Lactobacillus jensenii | Bacteria | 0.24 |
| 772 | 12041 | Bean leafroll virus | Viruses | 0.24 |
| 773 | 180903 | Pineapple mealybug wilt-associated virus 1 | Viruses | 0.24 |
| 774 | 12145 | Tomato bushy stunt virus | Viruses | 0.24 |
| 775 | 1633 | Lactobacillus vaginalis | Bacteria | 0.24 |
| 776 | 45661 | Banana bract mosaic virus | Viruses | 0.23 |
| 777 | 83771 | Succinivibrio dextrinosolvens | Bacteria | 0.23 |
| 778 | 167161 | Strawberry mottle virus | Viruses | 0.23 |
| 779 | 114652 | Streptococcus orisratti | Bacteria | 0.23 |
| 780 | 598660 | Corynebacterium pyruviciproducens | Bacteria | 0.23 |
| 781 | 1415630 | Pseudomonas sp. TKP | Bacteria | 0.23 |
| 782 | 253700 | Schlumbergera virus X | Viruses | 0.22 |
| 783 | 1463935 | Streptomyces sp. NRRL WC-3744 | Bacteria | 0.22 |
| 784 | 165096 | Weissella koreensis | Bacteria | 0.22 |
| 785 | 537288 | Megasphaera sp. DJF_B143 | Bacteria | 0.22 |
| 786 | 735 | Haemophilus parahaemolyticus | Bacteria | 0.22 |
| 787 | 397288 | Lachnospiraceae bacterium 3-1 | Bacteria | 0.22 |
| 788 | 1074044 | uncultured phage WW-nAnB | Viruses | 0.22 |
| 789 | 646413 | Streptococcus phage 5093 | Viruses | 0.21 |
| 790 | 1501391 | Alistipes inops | Bacteria | 0.21 |
| 791 | 1492 | Clostridium butyricum | Bacteria | 0.21 |
| 792 | 11008 | Saccharomyces cerevisiae virus L-A | Viruses | 0.21 |
| 793 | 849 | Fusobacterium gonidiaformans | Bacteria | 0.21 |
| 794 | 1739251 | Fusobacterium sp. HMSC073F01 | Bacteria | 0.21 |
| 795 | 1251 | Leuconostoc fallax | Bacteria | 0.21 |
| 796 | 563191 | Acidaminococcus sp. D21 | Bacteria | 0.21 |
| 797 | 1852372 | Varibaculum sp. Marseille-P2802 | Bacteria | 0.20 |
| 798 | 1776381 | Olsenella sp. KHD7 | Bacteria | 0.20 |
| 799 | 2051 | Mobiluncus curtisii | Bacteria | 0.20 |
| 800 | 1653434 | Sellimonas intestinalis | Bacteria | 0.20 |
| 801 | 483 | Neisseria cinerea | Bacteria | 0.20 |
| 802 | 428712 | Jonquetella anthropi | Bacteria | 0.20 |
| 803 | 675077 | Plum bark necrosis stem pitting-associated virus | Viruses | 0.20 |
| 804 | 2130 | Ureaplasma urealyticum | Bacteria | 0.20 |
| 805 | 938288 | Fenollaria massiliensis | Bacteria | 0.20 |
| 806 | 416586 | Selenomonas bovis | Bacteria | 0.20 |
| 807 | 304207 | Lactobacillus harbinensis | Bacteria | 0.20 |
| 808 | 1216932 | Clostridium bornimense | Bacteria | 0.20 |
| 809 | 196375 | Beet black scorch virus | Viruses | 0.19 |
| 810 | 1206566 | Blueberry virus A | Viruses | 0.19 |
| 811 | 28136 | Prevotella oulorum | Bacteria | 0.19 |
| 812 | 1658108 | Niameybacter massiliensis | Bacteria | 0.19 |
| 813 | 1720204 | Collinsella ihuae | Bacteria | 0.19 |

| | | | | |
|---|---|---|---|---|
| 814 | 712122 | Actinomyces sp. oral taxon 414 | Bacteria | 0.18 |
| 815 | 1505 | Paeniclostridium sordellii | Bacteria | 0.18 |
| 816 | 1497955 | Clostridiales bacterium KA00274 | Bacteria | 0.18 |
| 817 | 78448 | Bifidobacterium pullorum | Bacteria | 0.18 |
| 818 | 1291540 | Candidatus Methanomethylophilus alvus | Archaea | 0.18 |
| 819 | 1871014 | Arcanobacterium urinimassiliense | Bacteria | 0.18 |
| 820 | 1280674 | Prevotella sp. AGR2160 | Bacteria | 0.18 |
| 821 | 1580 | Lactobacillus brevis | Bacteria | 0.18 |
| 822 | 1871002 | Acidaminococcus timonensis | Bacteria | 0.18 |
| 823 | 174709 | Allobaculum stercoricanis | Bacteria | 0.18 |
| 824 | 1347366 | Clostridium sp. ND2 | Bacteria | 0.18 |
| 825 | 1303 | Streptococcus oralis | Bacteria | 0.18 |
| 826 | 1720317 | Porphyromonadaceae bacterium FC4 | Bacteria | 0.18 |
| 827 | 936588 | Veillonella sp. ACP1 | Bacteria | 0.18 |
| 828 | 1739406 | Actinomyces sp. HMSC035G02 | Bacteria | 0.17 |
| 829 | 296 | Pseudomonas fragi | Bacteria | 0.17 |
| 830 | 1852362 | Bacteroides ihuae | Bacteria | 0.17 |
| 831 | 64003 | Grapevine leafroll-associated virus 2 | Viruses | 0.17 |
| 832 | 1871033 | Olsenella sp. Marseille-P3197 | Bacteria | 0.17 |
| 833 | 157688 | Leptotrichia hofstadii | Bacteria | 0.17 |
| 834 | 1659298 | Nectarine stem pitting-associated virus | Viruses | 0.17 |
| 835 | 1715051 | Streptococcus sp. HMSC068F04 | Bacteria | 0.17 |
| 836 | 29272 | Turnip vein-clearing virus | Viruses | 0.17 |
| 837 | 1805470 | Clostridium sp. Marseille-P2414 | Bacteria | 0.16 |
| 838 | 1655645 | Parabacteroides phage YZ-2015b | Viruses | 0.16 |
| 839 | 305 | Ralstonia solanacearum | Bacteria | 0.16 |
| 840 | 66228 | Actinomyces europaeus | Bacteria | 0.16 |
| 841 | 139208 | Isoptericola variabilis | Bacteria | 0.16 |
| 842 | 1497953 | Bacteroidales bacterium KA00251 | Bacteria | 0.16 |
| 843 | 156974 | Dysgonomonas gadei | Bacteria | 0.16 |
| 844 | 47903 | Strawberry vein banding virus | Viruses | 0.16 |
| 845 | 852 | Fusobacterium perfoetens | Bacteria | 0.16 |
| 846 | 5741 | Giardia intestinalis | Eukaryota | 0.16 |
| 847 | 28133 | Prevotella nigrescens | Bacteria | 0.16 |
| 848 | 46619 | Sweet potato virus G | Viruses | 0.16 |
| 849 | 103724 | Grapevine asteroid mosaic-associated virus | Viruses | 0.16 |
| 850 | 1111454 | Megasphaera sp. BV3C16-1 | Bacteria | 0.15 |
| 851 | 1472765 | Peptoniphilus obesi | Bacteria | 0.15 |
| 852 | 104263 | Hop latent virus | Viruses | 0.15 |
| 853 | 204 | Campylobacter showae | Bacteria | 0.15 |
| 854 | 1032506 | Prevotella sp. MSX73 | Bacteria | 0.15 |
| 855 | 1334 | Streptococcus dysgalactiae | Bacteria | 0.15 |
| 856 | 1838287 | Gammaproteobacteria bacterium 2W06 | Bacteria | 0.15 |
| 857 | 1720315 | Eggerthellaceae bacterium AT8 | Bacteria | 0.15 |
| 858 | 1588753 | Coriobacteriales bacterium DNF00809 | Bacteria | 0.15 |
| 859 | 1046402 | Potato virus H | Viruses | 0.15 |
| 860 | 457416 | Veillonella sp. 3_1_44 | Bacteria | 0.15 |
| 861 | 1739522 | Haemophilus sp. HMSC068C11 | Bacteria | 0.15 |

| | | | | |
|---|---|---|---|---|
| 862 | 87541 | Aerococcus christensenii | Bacteria | 0.15 |
| 863 | 1343 | Streptococcus vestibularis | Bacteria | 0.15 |
| 864 | 1581080 | Streptococcus sp. HMSC10E12 | Bacteria | 0.15 |
| 865 | 652706 | Oribacterium sp. oral taxon 078 | Bacteria | 0.15 |
| 866 | 1203593 | Veillonella sp. HPA0037 | Bacteria | 0.15 |
| 867 | 143361 | Filifactor alocis | Bacteria | 0.15 |
| 868 | 1679444 | Akkermansia glycaniphila | Bacteria | 0.15 |
| 869 | 1871023 | Rikenella sp. Marseille-P3215 | Bacteria | 0.15 |
| 870 | 2098 | Mycoplasma hominis | Bacteria | 0.15 |
| 871 | 44008 | Enterococcus cecorum | Bacteria | 0.15 |
| 872 | 197614 | Streptococcus pasteurianus | Bacteria | 0.15 |
| 873 | 1410621 | Lachnospiraceae bacterium AD3010 | Bacteria | 0.15 |
| 874 | 867080 | Paenibacillus sp. IHB B 3415 | Bacteria | 0.15 |
| 875 | 1311 | Streptococcus agalactiae | Bacteria | 0.15 |
| 876 | 12430 | Garlic virus D | Viruses | 0.15 |
| 877 | 548908 | Fig fleck-associated virus | Viruses | 0.15 |
| 878 | 732 | Aggregatibacter aphrophilus | Bacteria | 0.15 |
| 879 | 12154 | Turnip yellow mosaic virus | Viruses | 0.15 |
| 880 | 164393 | Lactobacillus fuchuensis | Bacteria | 0.15 |
| 881 | 856 | Fusobacterium varium | Bacteria | 0.14 |
| 882 | 2096 | Mycoplasma gallisepticum | Bacteria | 0.14 |
| 883 | 267818 | Lactobacillus kefiranofaciens | Bacteria | 0.14 |
| 884 | 43769 | Corynebacterium propinquum | Bacteria | 0.14 |
| 885 | 5341 | Agaricus bisporus | Eukaryota | 0.14 |
| 886 | 66219 | Lachnoclostridium phytofermentans | Bacteria | 0.14 |
| 887 | 315405 | Streptococcus gallolyticus | Bacteria | 0.14 |
| 888 | 1739525 | Peptoniphilus sp. HMSC075B08 | Bacteria | 0.14 |
| 889 | 1161409 | Bifidobacterium sp. MSTE12 | Bacteria | 0.14 |
| 890 | 114922 | Carrot thin leaf virus | Viruses | 0.14 |
| 891 | 404196 | Blackberry yellow vein-associated virus | Viruses | 0.13 |
| 892 | 1943580 | Pyramidobacter sp. C12-8 | Bacteria | 0.13 |
| 893 | 1449896 | Uncultured phage WW-nAnB strain 2 | Viruses | 0.13 |
| 894 | 589436 | Prevotella fusca | Bacteria | 0.13 |
| 895 | 28080 | Campylobacter upsaliensis | Bacteria | 0.13 |
| 896 | 1501329 | Oribacterium parvum | Bacteria | 0.13 |
| 897 | 168135 | Apium virus Y | Viruses | 0.13 |
| 898 | 213 | Helicobacter cinaedi | Bacteria | 0.13 |
| 899 | 469588 | Bacteroides sp. 2_1_22 | Bacteria | 0.13 |
| 900 | 180311 | Hespellia stercorisuis | Bacteria | 0.13 |
| 901 | 193121 | Pea enation mosaic virus-1 | Viruses | 0.13 |
| 902 | 33036 | Anaerococcus tetradius | Bacteria | 0.13 |
| 903 | 863372 | Herbaspirillum huttiense | Bacteria | 0.13 |
| 904 | 1579342 | Streptococcus sp. 343_SSPC | Bacteria | 0.13 |
| 905 | 267135 | Porphyrobacter donghaensis | Bacteria | 0.13 |
| 906 | 1123754 | Rattail cactus necrosis-associated virus | Viruses | 0.12 |
| 907 | 1547 | Erysipelatoclostridium ramosum | Bacteria | 0.12 |
| 908 | 81424 | [Clostridium] methoxybenzovorans | Bacteria | 0.12 |
| 909 | 1495144 | methanogenic archaeon ISO4-H5 | Archaea | 0.12 |

| | | | | |
|---|---|---|---|---|
| 910 | 502393 | Gemella asaccharolytica | Bacteria | 0.12 |
| 911 | 147711 | Rhinovirus A | Viruses | 0.12 |
| 912 | 936375 | Mogibacterium sp. CM50 | Bacteria | 0.12 |
| 913 | 367121 | Grapevine leafroll-associated virus 10 | Viruses | 0.12 |
| 914 | 31504 | Tobacco ringspot virus satellite RNA | Viruses | 0.12 |
| 915 | 104955 | Lactobacillus frumenti | Bacteria | 0.12 |
| 916 | 83526 | Lactobacillus paralimentarius | Bacteria | 0.12 |
| 917 | 1335 | Streptococcus equinus | Bacteria | 0.12 |
| 918 | 425254 | Cellulosilyticum ruminicola | Bacteria | 0.12 |
| 919 | 397290 | Lachnospiraceae bacterium A2 | Bacteria | 0.12 |
| 920 | 1632 | Lactobacillus oris | Bacteria | 0.12 |
| 921 | 1288391 | Actinomyces timonensis | Bacteria | 0.11 |
| 922 | 1768874 | Sinapis alba cryptic virus 1 | Viruses | 0.11 |
| 923 | 77768 | Prevotella albensis | Bacteria | 0.11 |
| 924 | 1852379 | Veillonellaceae bacterium Marseille-P2911 | Bacteria | 0.11 |
| 925 | 1190620 | Atopobium sp. ICM42b | Bacteria | 0.11 |
| 926 | 546 | Citrobacter freundii | Bacteria | 0.11 |
| 927 | 1211388 | Apple green crinkle associated virus | Viruses | 0.11 |
| 928 | 1280686 | Butyrivibrio sp. MC2013 | Bacteria | 0.11 |
| 929 | 1105031 | Clostridium sp. MSTE9 | Bacteria | 0.11 |
| 930 | 59505 | Actinotignum schaalii | Bacteria | 0.11 |
| 931 | 188913 | Cetobacterium somerae | Bacteria | 0.11 |
| 932 | 65467 | Cherry green ring mottle virus | Viruses | 0.11 |
| 933 | 1105171 | Bacteroides phage B124-14 | Viruses | 0.11 |
| 934 | 28129 | Prevotella denticola | Bacteria | 0.11 |
| 935 | 218923 | Turnip rosette virus | Viruses | 0.11 |
| 936 | 37128 | Potato mop-top virus | Viruses | 0.11 |
| 937 | 1348 | Streptococcus parauberis | Bacteria | 0.11 |
| 938 | 1232449 | Clostridiales bacterium VE202-08 | Bacteria | 0.11 |
| 939 | 1712675 | Turicibacter sp. H121 | Bacteria | 0.11 |
| 940 | 439703 | Prevotella maculosa | Bacteria | 0.10 |
| 941 | 1629 | Weissella viridescens | Bacteria | 0.10 |
| 942 | 1922434 | Beihai mollusks virus 1 | Viruses | 0.10 |
| 943 | 42817 | Corynebacterium argentoratense | Bacteria | 0.10 |
| 944 | 52584 | Brachyspira pilosicoli | Bacteria | 0.10 |
| 945 | 47493 | Lactobacillus panis | Bacteria | 0.10 |
| 946 | 33945 | Enterococcus avium | Bacteria | 0.10 |
| 947 | 649739 | Actinomyces sp. oral taxon 848 | Bacteria | 0.10 |
| 948 | 113287 | Pseudoramibacter alactolyticus | Bacteria | 0.10 |
| 949 | 53363 | Brevibacterium mcbrellneri | Bacteria | 0.10 |
| 950 | 68033 | Carrot mottle virus | Viruses | 0.10 |
| 951 | 28131 | Prevotella intermedia | Bacteria | 0.10 |
| 952 | 12431 | Garlic virus C | Viruses | 0.10 |
| 953 | 228603 | Prevotella shahii | Bacteria | 0.10 |
| 954 | 318464 | Clostridium sulfidigenes | Bacteria | 0.10 |
| 955 | 88233 | Lactobacillus manihotivorans | Bacteria | 0.10 |
| 956 | 881 | Desulfovibrio vulgaris | Bacteria | 0.10 |
| 957 | 47715 | Lactobacillus rhamnosus | Bacteria | 0.10 |

| | | | | |
|---|---|---|---|---|
| 958 | 137732 | Granulicatella elegans | Bacteria | 0.10 |
| 959 | 1249 | Weissella paramesenteroides | Bacteria | 0.10 |
| 960 | 936591 | Veillonella sp. ICM51a | Bacteria | 0.10 |
| 961 | 167634 | Grapevine rootstock stem lesion associated virus | Viruses | 0.10 |
| 962 | 39681 | Asparagus virus 2 | Viruses | 0.10 |
| 963 | 71032 | Grapevine leafroll-associated virus 5 | Viruses | 0.10 |
| 964 | 259059 | Lactobacillus satsumensis | Bacteria | 0.09 |
| 965 | 178001 | Leuconostoc inhae | Bacteria | 0.09 |
| 966 | 712148 | Aggregatibacter sp. oral taxon 458 | Bacteria | 0.09 |
| 967 | 12045 | Potato leafroll virus | Viruses | 0.09 |
| 968 | 1328 | Streptococcus anginosus | Bacteria | 0.09 |
| 969 | 56879 | Oat blue dwarf virus | Viruses | 0.09 |
| 970 | 99179 | Bacteroides phage B40-8 | Viruses | 0.09 |
| 971 | 1501332 | Oribacterium asaccharolyticum | Bacteria | 0.09 |
| 972 | 12040 | Barley yellow dwarf virus-PAV | Viruses | 0.09 |
| 973 | 1054217 | Thermoplasmatales archaeon BRNA1 | Archaea | 0.09 |
| 974 | 1511840 | Primula malacoides virus 1 | Viruses | 0.09 |
| 975 | 2751 | Carnobacterium maltaromaticum | Bacteria | 0.09 |
| 976 | 90410 | Streptococcus phage DT1 | Viruses | 0.09 |
| 977 | 1247 | Oenococcus oeni | Bacteria | 0.09 |
| 978 | 246618 | Bifidobacterium thermacidophilum | Bacteria | 0.09 |
| 979 | 33029 | Anaerococcus hydrogenalis | Bacteria | 0.09 |
| 980 | 348151 | Lactobacillus siliginis | Bacteria | 0.09 |
| 981 | 1588751 | Tissierellia bacterium KA00581 | Bacteria | 0.09 |
| 982 | 33037 | Anaerococcus vaginalis | Bacteria | 0.09 |
| 983 | 714 | Aggregatibacter actinomycetemcomitans | Bacteria | 0.09 |
| 984 | 288000 | Bradyrhizobium sp. BTAi1 | Bacteria | 0.09 |
| 985 | 582 | Morganella morganii | Bacteria | 0.09 |
| 986 | 68892 | Streptococcus infantis | Bacteria | 0.09 |
| 987 | 52280 | Chilli veinal mottle virus | Viruses | 0.09 |
| 988 | 4903 | Cyberlindnera jadinii | Eukaryota | 0.09 |
| 989 | 1933294 | Impatiens necrotic spot tospovirus | Viruses | 0.09 |
| 990 | 203168 | Grapevine leafroll-associated virus 6 | Viruses | 0.09 |
| 991 | 1581114 | Enterococcus sp. HMSC05C03 | Bacteria | 0.09 |
| 992 | 208086 | Moroccan pepper virus | Viruses | 0.08 |
| 993 | 12138 | Maize chlorotic mottle virus | Viruses | 0.08 |
| 994 | 1755753 | Penicillium aurantiogriseum foetidus-like virus | Viruses | 0.08 |
| 995 | 5476 | Candida albicans | Eukaryota | 0.08 |
| 996 | 1840217 | Candidatus Arthromitus sp. SFB-turkey | Bacteria | 0.08 |
| 997 | 190721 | Ralstonia insidiosa | Bacteria | 0.08 |
| 998 | 529 | Ochrobactrum anthropi | Bacteria | 0.08 |
| 999 | 97253 | Eubacterium plexicaudatum | Bacteria | 0.08 |
| 1000 | 1673726 | Clostridiales bacterium SIT11 | Bacteria | 0.08 |
| 1001 | 273677 | Microbacterium oleivorans | Bacteria | 0.08 |
| 1002 | 29419 | Helicobacter canis | Bacteria | 0.08 |
| 1003 | 89463 | Sacbrood virus | Viruses | 0.08 |
| 1004 | 1347393 | Bacteroides neonati | Bacteria | 0.08 |
| 1005 | 860 | Fusobacterium periodonticum | Bacteria | 0.08 |

| | | | | |
|---|---|---|---|---|
| 1006 | 1203556 | Actinomyces sp. HPA0247 | Bacteria | 0.08 |
| 1007 | 348449 | Raphanus sativus cryptic virus 1 | Viruses | 0.08 |
| 1008 | 1235835 | Anaerotruncus sp. G3(2012) | Bacteria | 0.08 |
| 1009 | 1661 | Trueperella pyogenes | Bacteria | 0.08 |
| 1010 | 1246 | Leuconostoc lactis | Bacteria | 0.08 |
| 1011 | 516703 | Pontibacillus litoralis | Bacteria | 0.08 |
| 1012 | 1111133 | Peptoniphilus sp. BV3AC2 | Bacteria | 0.08 |
| 1013 | 1345 | Streptococcus ferus | Bacteria | 0.08 |
| 1014 | 485724 | Melon severe mosaic tospovirus | Viruses | 0.08 |
| 1015 | 1176736 | Pitaya virus X | Viruses | 0.08 |
| 1016 | 1324352 | Chryseobacterium gallinarum | Bacteria | 0.08 |
| 1017 | 37923 | Kocuria kristinae | Bacteria | 0.08 |
| 1018 | 1472764 | Kallipyga massiliensis | Bacteria | 0.07 |
| 1019 | 202566 | Cherry rasp leaf virus | Viruses | 0.07 |
| 1020 | 1852378 | Veillonellaceae bacterium Marseille-P2974 | Bacteria | 0.07 |
| 1021 | 43995 | Johnsonella ignava | Bacteria | 0.07 |
| 1022 | 430710 | Blueberry latent virus | Viruses | 0.07 |
| 1023 | 119219 | Cupriavidus metallidurans | Bacteria | 0.07 |
| 1024 | 1922435 | Beihai mollusks virus 2 | Viruses | 0.07 |
| 1025 | 12211 | Plum pox virus | Viruses | 0.07 |
| 1026 | 1232454 | Clostridiales bacterium VE202-26 | Bacteria | 0.07 |
| 1027 | 253701 | Zygocactus virus X | Viruses | 0.07 |
| 1028 | 12202 | Lettuce mosaic virus | Viruses | 0.07 |
| 1029 | 1561964 | Methanosphaera sp. WGK6 | Archaea | 0.07 |
| 1030 | 78257 | Bifidobacterium saeculare | Bacteria | 0.07 |
| 1031 | 839 | Prevotella ruminicola | Bacteria | 0.07 |
| 1032 | 5482 | Candida tropicalis | Eukaryota | 0.07 |
| 1033 | 1280670 | Butyrivibrio sp. AD3002 | Bacteria | 0.07 |
| 1034 | 118748 | Bulleidia extructa | Bacteria | 0.07 |
| 1035 | 33962 | Lactobacillus kefiri | Bacteria | 0.07 |
| 1036 | 1697797 | Actinobacteria bacterium UC5.1-1B11 | Bacteria | 0.07 |
| 1037 | 936549 | Actinomyces sp. ICM54 | Bacteria | 0.07 |
| 1038 | 1333651 | Bifidobacterium moukalabense | Bacteria | 0.07 |
| 1039 | 4959 | Debaryomyces hansenii | Eukaryota | 0.07 |
| 1040 | 591166 | Southern tomato virus | Viruses | 0.07 |
| 1041 | 1694 | Bifidobacterium pseudolongum | Bacteria | 0.07 |
| 1042 | 1468413 | Bacillus massilioanorexius | Bacteria | 0.07 |
| 1043 | 35288 | Grapevine virus A | Viruses | 0.07 |
| 1044 | 33031 | Peptoniphilus lacrimalis | Bacteria | 0.07 |
| 1045 | 712430 | Peptoniphilus sp. oral taxon 375 | Bacteria | 0.07 |
| 1046 | 326941 | Raspberry leaf mottle virus | Viruses | 0.07 |
| 1047 | 328061 | Radish mosaic virus | Viruses | 0.07 |
| 1048 | 12143 | Cucumber necrosis virus | Viruses | 0.07 |
| 1049 | 936574 | Shuttleworthia sp. MSX8B | Bacteria | 0.07 |
| 1050 | 253239 | Ethanoligenens harbinense | Bacteria | 0.07 |
| 1051 | 134358 | Westerdykella cylindrica | Eukaryota | 0.07 |
| 1052 | 1365 | Lactococcus plantarum | Bacteria | 0.07 |
| 1053 | 51288 | Kluyvera ascorbata | Bacteria | 0.06 |

| | | | | |
|---|---|---|---|---|
| 1054 | 1080349 | Saccharomyces eubayanus | Eukaryota | 0.06 |
| 1055 | 712121 | Actinomyces sp. oral taxon 181 | Bacteria | 0.06 |
| 1056 | 208084 | Grapevine Algerian latent virus | Viruses | 0.06 |
| 1057 | 1118058 | Actinomyces sp. ph3 | Bacteria | 0.06 |
| 1058 | 1280 | Staphylococcus aureus | Bacteria | 0.06 |
| 1059 | 712156 | Atopobium sp. oral taxon 199 | Bacteria | 0.06 |
| 1060 | 1235790 | Eubacterium sp. 14-2 | Bacteria | 0.06 |
| 1061 | 1235798 | Dorea sp. 5-2 | Bacteria | 0.06 |
| 1062 | 12222 | Soybean mosaic virus | Viruses | 0.06 |
| 1063 | 1299998 | Olsenella scatoligenes | Bacteria | 0.06 |
| 1064 | 1410616 | Pseudobutyrivibrio sp. MD2005 | Bacteria | 0.06 |
| 1065 | 1739371 | Streptococcus sp. HMSC064H09 | Bacteria | 0.06 |
| 1066 | 590403 | Red clover vein mosaic virus | Viruses | 0.06 |
| 1067 | 147712 | Rhinovirus B | Viruses | 0.06 |
| 1068 | 1276 | Kytococcus sedentarius | Bacteria | 0.06 |
| 1069 | 31713 | Lettuce infectious yellows virus | Viruses | 0.06 |
| 1070 | 118562 | Arthrospira platensis | Bacteria | 0.06 |
| 1071 | 1080712 | Methanomassiliicoccus luminyensis | Archaea | 0.06 |
| 1072 | 1720298 | Peptoniphilus phoceensis | Bacteria | 0.06 |
| 1073 | 172220 | Blueberry red ringspot virus | Viruses | 0.06 |
| 1074 | 614 | Serratia liquefaciens | Bacteria | 0.06 |
| 1075 | 1230730 | Tissierellia bacterium S5-A11 | Bacteria | 0.06 |
| 1076 | 1050903 | Pepper cryptic virus 2 | Viruses | 0.06 |
| 1077 | 54067 | Xylophilus ampelinus | Bacteria | 0.06 |
| 1078 | 35519 | Mogibacterium timidum | Bacteria | 0.06 |
| 1079 | 1529 | Clostridium cadaveris | Bacteria | 0.06 |
| 1080 | 453050 | Sweet potato virus 2 | Viruses | 0.06 |
| 1081 | 1739537 | Anaerococcus sp. HMSC075B03 | Bacteria | 0.06 |
| 1082 | 1161412 | Prevotella sp. ICM33 | Bacteria | 0.06 |
| 1083 | 392416 | Lactobacillus crustorum | Bacteria | 0.06 |
| 1084 | 1358418 | Sinorhizobium sp. GL28 | Bacteria | 0.06 |
| 1085 | 1774276 | Hordeum vulgare endornavirus | Viruses | 0.06 |
| 1086 | 936589 | Veillonella sp. AS16 | Bacteria | 0.06 |
| 1087 | 1564 | Desulfotomaculum ruminis | Bacteria | 0.06 |
| 1088 | 1852373 | Murdochiella sp. Marseille-P2341 | Bacteria | 0.05 |
| 1089 | 194326 | Lactobacillus versmoldensis | Bacteria | 0.05 |
| 1090 | 171523 | Lactobacillus pantheris | Bacteria | 0.05 |
| 1091 | 1281 | Staphylococcus carnosus | Bacteria | 0.05 |
| 1092 | 12139 | Southern bean mosaic virus | Viruses | 0.05 |
| 1093 | 1777865 | Weissella sp. DD23 | Bacteria | 0.05 |
| 1094 | 1610 | Lactobacillus coryniformis | Bacteria | 0.05 |
| 1095 | 37372 | Helicobacter bilis | Bacteria | 0.05 |
| 1096 | 1686287 | Kallipyga gabonensis | Bacteria | 0.05 |
| 1097 | 1497954 | Bacteroidales bacterium KA00344 | Bacteria | 0.05 |
| 1098 | 131082 | Beet chlorosis virus | Viruses | 0.05 |
| 1099 | 1297865 | Bradyrhizobium sp. OHSU_III | Bacteria | 0.05 |
| 1100 | 1261640 | Eubacterium sp. 68-3-10 | Bacteria | 0.05 |
| 1101 | 1587 | Lactobacillus helveticus | Bacteria | 0.05 |

| | | | | |
|---|---|---|---|---|
| 1102 | 1597 | Lactobacillus paracasei | Bacteria | 0.05 |
| 1103 | 1715211 | Haemophilus sp. HMSC061E01 | Bacteria | 0.05 |
| 1104 | 507750 | Peptoniphilus duerdenii | Bacteria | 0.05 |
| 1105 | 31537 | Lactococcus phage c2 | Viruses | 0.05 |
| 1106 | 92395 | Black queen cell virus | Viruses | 0.05 |
| 1107 | 195 | Campylobacter coli | Bacteria | 0.05 |
| 1108 | 28137 | Prevotella veroralis | Bacteria | 0.05 |
| 1109 | 119910 | Ryegrass mottle virus | Viruses | 0.05 |
| 1110 | 1343493 | Grapevine satellite virus | Viruses | 0.05 |
| 1111 | 379892 | Passiflora latent carlavirus | Viruses | 0.05 |
| 1112 | 28124 | Porphyromonas endodontalis | Bacteria | 0.05 |
| 1113 | 43130 | Onion yellow dwarf virus | Viruses | 0.05 |
| 1114 | 1235800 | Lachnospiraceae bacterium 10-1 | Bacteria | 0.05 |
| 1115 | 339420 | Blackberry chlorotic ringspot virus | Viruses | 0.05 |
| 1116 | 177972 | Shuttleworthia satelles | Bacteria | 0.05 |
| 1117 | 1525173 | Human circovirus VS6600022 | Viruses | 0.05 |
| 1118 | 12183 | Potato virus X | Viruses | 0.05 |
| 1119 | 11986 | Carnation mottle virus | Viruses | 0.05 |
| 1120 | 138950 | Enterovirus C | Viruses | 0.05 |
| 1121 | 137730 | Facklamia ignava | Bacteria | 0.05 |
| 1122 | 198112 | Deformed wing virus | Viruses | 0.05 |
| 1123 | 1579341 | Streptococcus sp. 400_SSPC | Bacteria | 0.05 |
| 1124 | 671230 | Parvimonas sp. oral taxon 110 | Bacteria | 0.05 |
| 1125 | 1933295 | Iris yellow spot tospovirus | Viruses | 0.05 |
| 1126 | 1353 | Enterococcus gallinarum | Bacteria | 0.05 |
| 1127 | 1354 | Enterococcus hirae | Bacteria | 0.05 |
| 1128 | 1932006 | Chicken associated smacovirus | Viruses | 0.05 |
| 1129 | 392504 | Turnip ringspot virus | Viruses | 0.05 |
| 1130 | 134605 | Fusobacterium equinum | Bacteria | 0.05 |
| 1131 | 1000568 | Megasphaera sp. UPII 199-6 | Bacteria | 0.05 |
| 1132 | 587 | Providencia rettgeri | Bacteria | 0.05 |
| 1133 | 46867 | Clostridium chauvoei | Bacteria | 0.05 |
| 1134 | 471858 | Helicobacter magdeburgensis | Bacteria | 0.05 |
| 1135 | 12238 | Odontoglossum ringspot virus | Viruses | 0.05 |
| 1136 | 876478 | Tepidiphilus thermophilus | Bacteria | 0.05 |
| 1137 | 28985 | Kluyveromyces lactis | Eukaryota | 0.05 |
| 1138 | 82347 | Facklamia languida | Bacteria | 0.05 |
| 1139 | 1080072 | Streptococcus dentasini | Bacteria | 0.05 |
| 1140 | 1764 | Mycobacterium avium | Bacteria | 0.05 |
| 1141 | 1840518 | Gardnerella sp. 30-4 | Bacteria | 0.05 |
| 1142 | 167 | Treponema succinifaciens | Bacteria | 0.05 |
| 1143 | 936550 | Atopobium sp. BS2 | Bacteria | 0.05 |
| 1144 | 1423347 | Sclerotinia sclerotiorum hypovirus 2 | Viruses | 0.05 |
| 1145 | 516956 | Grapevine virus E | Viruses | 0.05 |
| 1146 | 1739422 | Streptococcus sp. HMSC065C01 | Bacteria | 0.05 |
| 1147 | 5478 | [Candida] glabrata | Eukaryota | 0.05 |
| 1148 | 28348 | Sweet clover necrotic mosaic virus | Viruses | 0.05 |
| 1149 | 1384078 | Prevotella sp. DNF00663 | Bacteria | 0.05 |

| | | | | |
|---|---|---|---|---|
| 1150 | 1330491 | Cosavirus A | Viruses | 0.05 |
| 1151 | 140626 | Lachnobacterium bovis | Bacteria | 0.05 |
| 1152 | 173976 | Cycas necrotic stunt virus | Viruses | 0.05 |
| 1153 | 29359 | Asaccharospora irregularis | Bacteria | 0.05 |
| 1154 | 257464 | Potato black ringspot virus | Viruses | 0.05 |
| 1155 | 12271 | Arabis mosaic virus | Viruses | 0.05 |
| 1156 | 1425364 | Carrot torradovirus 1 | Viruses | 0.05 |
| 1157 | 195054 | Human parechovirus | Viruses | 0.05 |
| 1158 | 83559 | Chlamydia suis | Bacteria | 0.05 |
| 1159 | 1323524 | Red clover cryptic virus 2 | Viruses | 0.05 |
| 1160 | 225992 | Comamonas kerstersii | Bacteria | 0.05 |
| 1161 | 64958 | Parietaria mottle virus | Viruses | 0.05 |
| 1162 | 1719140 | Klebsiella phage vB_KpnM_KB57 | Viruses | 0.05 |
| 1163 | 682370 | Streptococcus phage Alq132 | Viruses | 0.05 |
| 1164 | 328396 | Enterococcus aquimarinus | Bacteria | 0.05 |
| 1165 | 53444 | Lactobacillus lindneri | Bacteria | 0.05 |
| 1166 | 1711684 | Hot pepper endornavirus | Viruses | 0.05 |
| 1167 | 595895 | Drosophila A virus | Viruses | 0.05 |
| 1168 | 41997 | Enterococcus saccharolyticus | Bacteria | 0.05 |
| 1169 | 909827 | Pepper vein yellows virus | Viruses | 0.05 |
| 1170 | 457388 | Parabacteroides sp. 2_1_7 | Bacteria | 0.05 |
| 1171 | 1675609 | Klebsiella phage Sushi | Viruses | 0.05 |
| 1172 | 12432 | Garlic virus B | Viruses | 0.05 |
| 1173 | 936381 | Selenomonas sp. CM52 | Bacteria | 0.05 |
| 1174 | 1715086 | Streptococcus sp. HMSC078H03 | Bacteria | 0.05 |
| 1175 | 193120 | Pea enation mosaic virus-2 | Viruses | 0.05 |
| 1176 | 664639 | Kocuria salsicia | Bacteria | 0.05 |
| 1177 | 374840 | Enterobacteria phage phiX174 sensu lato | Viruses | 0.05 |
| 1178 | 671216 | Peptoniphilus sp. oral taxon 836 | Bacteria | 0.05 |
| 1179 | 12165 | Chrysanthemum virus B | Viruses | 0.05 |
| 1180 | 12167 | Potato virus M | Viruses | 0.05 |
| 1181 | 12161 | Beet yellows virus | Viruses | 0.05 |
| 1182 | 1520332 | Blueberry mosaic associated virus | Viruses | 0.05 |
| 1183 | 589537 | Prevotella dentasini | Bacteria | 0.05 |
| 1184 | 1739286 | Anaerococcus sp. HMSC068A02 | Bacteria | 0.05 |
| 1185 | 12197 | Bean yellow mosaic virus | Viruses | 0.05 |
| 1186 | 1095727 | Streptococcus sp. SK643 | Bacteria | 0.05 |
| 1187 | 257758 | Streptococcus pseudopneumoniae | Bacteria | 0.05 |
| 1188 | 71030 | Chayote mosaic virus | Viruses | 0.05 |
| 1189 | 66834 | Rhopalosiphum padi virus | Viruses | 0.05 |
| 1190 | 97139 | Clostridium sp. ASF502 | Bacteria | 0.05 |
| 1191 | 57732 | Enterococcus asini | Bacteria | 0.05 |
| 1192 | 396268 | Lactobacillus secaliphilus | Bacteria | 0.05 |
| 1193 | 1654927 | Klebsiella phage PKP126 | Viruses | 0.05 |
| 1194 | 28197 | Arcobacter butzleri | Bacteria | 0.05 |
| 1195 | 298 | Pseudomonas marginalis | Bacteria | 0.04 |
| 1196 | 1805472 | Clostridium sp. Marseille-P2434 | Bacteria | 0.04 |
| 1197 | 1480694 | Spirochaeta lutea | Bacteria | 0.04 |

| | | | | |
|---|---|---|---|---|
| 1198 | 399370 | Lactobacillus ghanensis | Bacteria | 0.04 |
| 1199 | 1282 | Staphylococcus epidermidis | Bacteria | 0.04 |
| 1200 | 35841 | Bacillus thermoamylovorans | Bacteria | 0.04 |
| 1201 | 46124 | Granulicatella adiacens | Bacteria | 0.04 |
| 1202 | 405212 | Alicyclobacillus acidocaldarius | Bacteria | 0.04 |
| 1203 | 578361 | Soybean yellow mottle mosaic virus | Viruses | 0.04 |
| 1204 | 1235799 | Lachnospiraceae bacterium 3-2 | Bacteria | 0.04 |
| 1205 | 1101373 | Tepidimonas fonticaldi | Bacteria | 0.04 |
| 1206 | 1739413 | Alloscardovia sp. HMSC034E08 | Bacteria | 0.04 |
| 1207 | 545 | Citrobacter koseri | Bacteria | 0.04 |
| 1208 | 46206 | Pseudobutyrivibrio ruminis | Bacteria | 0.04 |
| 1209 | 1658779 | Porphyromonadaceae bacterium H1 | Bacteria | 0.04 |
| 1210 | 31749 | Obuda pepper virus | Viruses | 0.04 |
| 1211 | 79604 | Denitrobacterium detoxificans | Bacteria | 0.04 |
| 1212 | 39397 | Candida sake | Eukaryota | 0.04 |
| 1213 | 1923594 | Wenzhou picorna-like virus 10 | Viruses | 0.04 |
| 1214 | 1674942 | Caenibacillus caldisaponilyticus | Bacteria | 0.04 |
| 1215 | 39497 | Eubacterium xylanophilum | Bacteria | 0.04 |
| 1216 | 1469950 | Robinsoniella sp. KNHs210 | Bacteria | 0.04 |
| 1217 | 12275 | Tomato black ring virus | Viruses | 0.04 |
| 1218 | 129141 | Citrus leaf blotch virus | Viruses | 0.04 |
| 1219 | 1270 | Micrococcus luteus | Bacteria | 0.04 |
| 1220 | 29385 | Staphylococcus saprophyticus | Bacteria | 0.04 |
| 1221 | 713008 | Parvimonas sp. oral taxon 393 | Bacteria | 0.04 |
| 1222 | 1433844 | Prevotella sp. HJM029 | Bacteria | 0.04 |
| 1223 | 204933 | Grapevine rupestris vein feathering virus | Viruses | 0.04 |
| 1224 | 47736 | Carrot mottle mimic virus | Viruses | 0.04 |
| 1225 | 1579339 | Streptococcus sp. 449_SSPC | Bacteria | 0.04 |
| 1226 | 831 | Butyrivibrio fibrisolvens | Bacteria | 0.04 |
| 1227 | 1739452 | Haemophilus sp. HMSC073C03 | Bacteria | 0.04 |
| 1228 | 1727 | Corynebacterium variabile | Bacteria | 0.04 |
| 1229 | 1323525 | White clover cryptic virus 2 | Viruses | 0.04 |
| 1230 | 69823 | Selenomonas sputigena | Bacteria | 0.04 |
| 1231 | 102684 | Streptococcus infantarius | Bacteria | 0.04 |
| 1232 | 237258 | Cloacibacterium normanense | Bacteria | 0.04 |
| 1233 | 36911 | Clavispora lusitaniae | Eukaryota | 0.04 |
| 1234 | 106008 | Curvibasidium cygneicollum | Eukaryota | 0.04 |
| 1235 | 1921123 | Salmonella virus 9NA | Viruses | 0.04 |
| 1236 | 1815509 | Bacillus phage AR9 | Viruses | 0.04 |
| 1237 | 49266 | Fucus vesiculosus | Eukaryota | 0.04 |
| 1238 | 1794912 | Anaerosporomusa subterranea | Bacteria | 0.04 |
| 1239 | 1581148 | Clostridium sp. HMSC19A10 | Bacteria | 0.04 |
| 1240 | 152331 | Lactobacillus parabuchneri | Bacteria | 0.04 |
| 1241 | 683172 | Astrovirus MLB2 | Viruses | 0.04 |
| 1242 | 264463 | Anaerosporobacter mobilis | Bacteria | 0.04 |
| 1243 | 76860 | Streptococcus constellatus | Bacteria | 0.04 |
| 1244 | 1860161 | Streptococcus sp. CCUG 49591 | Bacteria | 0.04 |
| 1245 | 568715 | Astrovirus MLB1 | Viruses | 0.04 |

| | | | | |
|---|---|---|---|---|
| 1246 | 270256 | Eggplant mottled crinkle virus | Viruses | 0.04 |
| 1247 | 1739381 | Streptococcus sp. HMSC072D03 | Bacteria | 0.04 |
| 1248 | 12230 | Turnip mosaic virus | Viruses | 0.04 |
| 1249 | 1930298 | Chicken stool-associated gemycircularvirus | Viruses | 0.04 |
| 1250 | 1133319 | Bacteroides reticulotermitis | Bacteria | 0.04 |
| 1251 | 1384082 | Veillonellaceae bacterium DNF00751 | Bacteria | 0.04 |
| 1252 | 31721 | Beet necrotic yellow vein virus | Viruses | 0.04 |
| 1253 | 167481 | Lactobacillus mindensis | Bacteria | 0.04 |
| 1254 | 1316412 | Streptococcus sp. HSISS3 | Bacteria | 0.04 |
| 1255 | 1279099 | Sclerotinia sclerotiorum mitovirus 3 | Viruses | 0.04 |
| 1256 | 1161417 | Streptococcus sp. SR4 | Bacteria | 0.04 |
| 1257 | 1597976 | Enterococcus phage EFDG1 | Viruses | 0.04 |
| 1258 | 339 | Xanthomonas campestris | Bacteria | 0.04 |
| 1259 | 1330524 | Salivirus A | Viruses | 0.04 |
| 1260 | 28141 | Cronobacter sakazakii | Bacteria | 0.04 |
| 1261 | 10840 | Beet curly top virus | Viruses | 0.04 |
| 1262 | 138949 | Enterovirus B | Viruses | 0.04 |
| 1263 | 138948 | Enterovirus A | Viruses | 0.04 |
| 1264 | 584 | Proteus mirabilis | Bacteria | 0.04 |
| 1265 | 72750 | Beet pseudoyellows virus | Viruses | 0.04 |
| 1266 | 569 | Hafnia alvei | Bacteria | 0.04 |
| 1267 | 1165092 | Lachnospiraceae bacterium JC7 | Bacteria | 0.04 |
| 1268 | 1392486 | Prevotella sp. HUN102 | Bacteria | 0.04 |
| 1269 | 4909 | Pichia kudriavzevii | Eukaryota | 0.04 |
| 1270 | 1739356 | Anaerococcus sp. HMSC065G05 | Bacteria | 0.04 |
| 1271 | 186189 | Xylanimonas cellulosilytica | Bacteria | 0.04 |
| 1272 | 1862960 | Lactococcus phage M5938 | Viruses | 0.04 |
| 1273 | 571 | Klebsiella oxytoca | Bacteria | 0.04 |
| 1274 | 1423 | Bacillus subtilis | Bacteria | 0.04 |
| 1275 | 47985 | Grapevine leafroll-associated virus 1 | Viruses | 0.04 |
| 1276 | 213633 | Providence virus | Viruses | 0.04 |
| 1277 | 1519 | Clostridium tyrobutyricum | Bacteria | 0.04 |
| 1278 | 1131702 | Persea americana endornavirus 1 | Viruses | 0.04 |
| 1279 | 397287 | Lachnospiraceae bacterium 28-4 | Bacteria | 0.04 |
| 1280 | 1608898 | Haemophilus sp. HMSC71H05 | Bacteria | 0.04 |
| 1281 | 264634 | Acholeplasma equifetale | Bacteria | 0.04 |
| 1282 | 726 | Haemophilus haemolyticus | Bacteria | 0.04 |
| 1283 | 1748 | Acidipropionibacterium acidipropionici | Bacteria | 0.04 |
| 1284 | 12558 | Sesbania mosaic virus | Viruses | 0.04 |
| 1285 | 1592930 | Yam latent virus | Viruses | 0.04 |
| 1286 | 1930509 | Husavirus sp. | Viruses | 0.03 |
| 1287 | 85454 | Alternanthera mosaic virus | Viruses | 0.03 |
| 1288 | 294 | Pseudomonas fluorescens | Bacteria | 0.03 |
| 1289 | 218140 | Bifidobacterium psychraerophilum | Bacteria | 0.03 |
| 1290 | 290399 | Arthrobacter sp. FB24 | Bacteria | 0.03 |
| 1291 | 550 | Enterobacter cloacae | Bacteria | 0.03 |
| 1292 | 333754 | Alphapapillomavirus 10 | Viruses | 0.03 |
| 1293 | 1315956 | Shigella phage pSf-1 | Viruses | 0.03 |

| | | | | |
|---|---|---|---|---|
| 1294 | 69966 | Macrococcus caseolyticus | Bacteria | 0.03 |
| 1295 | 1027232 | Groundnut ringspot and Tomato chlorotic spot virus reassortant | Viruses | 0.03 |
| 1296 | 1922513 | Beihai permutotetra-like virus 2 | Viruses | 0.03 |
| 1297 | 179628 | Clostridium colicanis | Bacteria | 0.03 |
| 1298 | 1702221 | Faecalibaculum rodentium | Bacteria | 0.03 |
| 1299 | 1141139 | Enterobacteria phage vB_EcoP_ACG-C91 | Viruses | 0.03 |
| 1300 | 112436 | Celery mosaic virus | Viruses | 0.03 |
| 1301 | 548 | Klebsiella aerogenes | Bacteria | 0.03 |
| 1302 | 126385 | Providencia alcalifaciens | Bacteria | 0.03 |
| 1303 | 988645 | Raspberry leaf blotch virus | Viruses | 0.03 |
| 1304 | 1920861 | Klebsiella virus KP36 | Viruses | 0.03 |
| 1305 | 354259 | Lactococcus phage 936 sensu lato | Viruses | 0.03 |
| 1306 | 546367 | Hafnia paralvei | Bacteria | 0.03 |
| 1307 | 13335 | Anaerobiospirillum succiniciproducens | Bacteria | 0.03 |
| 1308 | 248039 | Mucispirillum schaedleri | Bacteria | 0.03 |
| 1309 | 651609 | Actinomyces sp. oral taxon 180 | Bacteria | 0.03 |
| 1310 | 12450 | Saccharomyces cerevisiae killer virus M1 | Viruses | 0.03 |
| 1311 | 1723382 | Peptoniphilaceae bacterium FC2 | Bacteria | 0.03 |
| 1312 | 373054 | Calditerricola satsumensis | Bacteria | 0.03 |
| 1313 | 241555 | Helcococcus sueciensis | Bacteria | 0.03 |
| 1314 | 1922835 | Hubei arthropod virus 1 | Viruses | 0.03 |
| 1315 | 682382 | HMO Astrovirus A | Viruses | 0.03 |
| 1316 | 12208 | Pea seed-borne mosaic virus | Viruses | 0.03 |
| 1317 | 78259 | Scardovia inopinata | Bacteria | 0.03 |
| 1318 | 31770 | Shallot virus X | Viruses | 0.03 |
| 1319 | 12055 | Tobacco necrosis virus A | Viruses | 0.03 |
| 1320 | 111015 | Actinomyces radicidentis | Bacteria | 0.03 |
| 1321 | 1465756 | Peptoniphilus grossensis | Bacteria | 0.03 |
| 1322 | 469553 | Delftia sp. JD2 | Bacteria | 0.03 |
| 1323 | 1870984 | Anaerococcus mediterraneensis | Bacteria | 0.03 |
| 1324 | 12313 | Peanut stunt virus | Viruses | 0.03 |
| 1325 | 1868658 | Human astrovirus | Viruses | 0.03 |
| 1326 | 157687 | Leptotrichia wadei | Bacteria | 0.03 |
| 1327 | 1582 | Lactobacillus casei | Bacteria | 0.03 |
| 1328 | 351091 | Oscillibacter valericigenes | Bacteria | 0.03 |
| 1329 | 35818 | Helicobacter pullorum | Bacteria | 0.03 |
| 1330 | 72539 | Physalis mottle virus | Viruses | 0.03 |
| 1331 | 1590596 | Sphingobacterium sp. T2 | Bacteria | 0.03 |
| 1332 | 28112 | Tannerella forsythia | Bacteria | 0.03 |
| 1333 | 1756285 | Maize associated totivirus | Viruses | 0.03 |
| 1334 | 260742 | Streptomyces sp. SS | Bacteria | 0.03 |
| 1335 | 1111121 | Atopobium sp. BV3Ac4 | Bacteria | 0.03 |
| 1336 | 1343920 | Apricot vein clearing associated virus | Viruses | 0.03 |
| 1337 | 47669 | Olive latent virus 1 | Viruses | 0.03 |
| 1338 | 1567484 | Lactobacillus phage LfeInf | Viruses | 0.03 |
| 1339 | 329853 | Escherichia virus BZ13 | Viruses | 0.03 |
| 1340 | 329852 | Escherichia virus MS2 | Viruses | 0.03 |

| | | | | |
|---|---|---|---|---|
| 1341 | 197 | Campylobacter jejuni | Bacteria | 0.03 |
| 1342 | 1156433 | Streptococcus sp. I-P16 | Bacteria | 0.03 |
| 1343 | 1718158 | Enterococcus phage IME-EFm5 | Viruses | 0.03 |
| 1344 | 1702287 | Negativicoccus massiliensis | Bacteria | 0.03 |
| 1345 | 1739309 | Streptococcus sp. HMSC034E03 | Bacteria | 0.03 |
| 1346 | 158787 | Bifidobacterium scardovii | Bacteria | 0.03 |
| 1347 | 5353 | Lentinula edodes | Eukaryota | 0.03 |
| 1348 | 1859694 | Haemophilus sp. CCUG 66565 | Bacteria | 0.03 |
| 1349 | 28134 | Prevotella oralis | Bacteria | 0.03 |
| 1350 | 227942 | Lactobacillus gastricus | Bacteria | 0.03 |
| 1351 | 293371 | Lactobacillus oligofermentans | Bacteria | 0.03 |
| 1352 | 298338 | Lactobacillus phage LP65 | Viruses | 0.03 |
| 1353 | 1435146 | Morganella sp. EGD-HP17 | Bacteria | 0.03 |
| 1354 | 471285 | Lettuce yellow mottle virus | Viruses | 0.03 |
| 1355 | 1891289 | Sporanaerobacter sp. PP17-6a | Bacteria | 0.03 |
| 1356 | 1911008 | Escherichia virus K1G | Viruses | 0.03 |
| 1357 | 1314 | Streptococcus pyogenes | Bacteria | 0.03 |
| 1358 | 630199 | Grapevine Syrah virus 1 | Viruses | 0.03 |
| 1359 | 712531 | Selenomonas sp. oral taxon 137 | Bacteria | 0.03 |
| 1360 | 111418 | Zucchini green mottle mosaic virus | Viruses | 0.03 |
| 1361 | 151276 | Bacteroides coprosuis | Bacteria | 0.03 |
| 1362 | 1871031 | Olsenella sp. Marseille-P3256 | Bacteria | 0.03 |
| 1363 | 469594 | Bifidobacterium sp. 12_1_47BFAA | Bacteria | 0.03 |
| 1364 | 95609 | Herbaspirillum sp. B39 | Bacteria | 0.03 |
| 1365 | 1134687 | Klebsiella michiganensis | Bacteria | 0.03 |
| 1366 | 51354 | Maize chlorotic dwarf virus | Viruses | 0.03 |
| 1367 | 35703 | Citrobacter amalonaticus | Bacteria | 0.03 |
| 1368 | 83231 | Prevotella brevis | Bacteria | 0.03 |
| 1369 | 388038 | Cucumber mottle virus | Viruses | 0.03 |
| 1370 | 463676 | Rhinovirus C | Viruses | 0.03 |
| 1371 | 282402 | Prevotella multiformis | Bacteria | 0.03 |
| 1372 | 57706 | Citrobacter braakii | Bacteria | 0.03 |
| 1373 | 908834 | Grapevine associated narnavirus-1 | Viruses | 0.03 |
| 1374 | 43675 | Rothia mucilaginosa | Bacteria | 0.03 |
| 1375 | 134632 | American plum line pattern virus | Viruses | 0.03 |
| 1376 | 1658008 | Dysgonomonas sp. BGC7 | Bacteria | 0.03 |
| 1377 | 112023 | Streptococcus phage 7201 | Viruses | 0.03 |
| 1378 | 85154 | Streptococcus phage O1205 | Viruses | 0.03 |
| 1379 | 2293 | Desulfobacter postgatei | Bacteria | 0.03 |
| 1380 | 67824 | Citrobacter farmeri | Bacteria | 0.03 |
| 1381 | 169292 | Corynebacterium aurimucosum | Bacteria | 0.03 |
| 1382 | 539813 | Enterobacter mori | Bacteria | 0.02 |
| 1383 | 1608882 | Haemophilus sp. HMSC61B11 | Bacteria | 0.02 |
| 1384 | 277 | Meiothermus ruber | Bacteria | 0.02 |
| 1385 | 1163671 | Clostridium sp. 12(A) | Bacteria | 0.02 |
| 1386 | 1914853 | Escherichia virus V5 | Viruses | 0.02 |
| 1387 | 1914855 | Escherichia virus FV3 | Viruses | 0.02 |
| 1388 | 1914854 | Escherichia virus JES2013 | Viruses | 0.02 |

| | | | | |
|---|---|---|---|---|
| 1389 | 136609 | Leuconostoc kimchii | Bacteria | 0.02 |
| 1390 | 151414 | Afipia birgiae | Bacteria | 0.02 |
| 1391 | 217686 | Little cherry virus 1 | Viruses | 0.02 |
| 1392 | 73098 | Kluyvera georgiana | Bacteria | 0.02 |
| 1393 | 52253 | Candida sojae | Eukaryota | 0.02 |
| 1394 | 1698360 | Klebsiella phage JD18 | Viruses | 0.02 |
| 1395 | 191217 | Cereal yellow dwarf virus-RPV | Viruses | 0.02 |
| 1396 | 1301100 | [Clostridium] dakarense | Bacteria | 0.02 |
| 1397 | 1795648 | Picornavirales Tottori-HG1 | Viruses | 0.02 |
| 1398 | 149016 | Streptococcus urinalis | Bacteria | 0.02 |
| 1399 | 1235793 | Lachnospiraceae bacterium COE1 | Bacteria | 0.02 |
| 1400 | 1852368 | Prevotellaceae bacterium Marseille-P2826 | Bacteria | 0.02 |
| 1401 | 12227 | Tobacco etch virus | Viruses | 0.02 |
| 1402 | 331278 | Yersinia phage phiR1-37 | Viruses | 0.02 |
| 1403 | 1612 | Lactobacillus farciminis | Bacteria | 0.02 |
| 1404 | 948870 | Enterobacteria phage phi92 | Viruses | 0.02 |
| 1405 | 114871 | Zygosaccharomyces bailii virus Z | Viruses | 0.02 |
| 1406 | 1920862 | Klebsiella virus 1513 | Viruses | 0.02 |
| 1407 | 570949 | Carrot mottle mimic virus satellite RNA | Viruses | 0.02 |
| 1408 | 337048 | Alphapapillomavirus 1 | Viruses | 0.02 |
| 1409 | 1913024 | White clover mottle virus | Viruses | 0.02 |
| 1410 | 71452 | Enterococcus raffinosus | Bacteria | 0.02 |
| 1411 | 1522179 | Asterionellopsis glacialis RNA virus | Viruses | 0.02 |
| 1412 | 1496722 | Butyrivibrio sp. AE2005 | Bacteria | 0.02 |
| 1413 | 1408324 | Lachnospiraceae bacterium MC2017 | Bacteria | 0.02 |
| 1414 | 1737425 | Corynebacterium provencense | Bacteria | 0.02 |
| 1415 | 4950 | Torulaspora delbrueckii | Eukaryota | 0.02 |
| 1416 | 1600 | Lactobacillus acetotolerans | Bacteria | 0.02 |
| 1417 | 179878 | Sphingomonas elodea | Bacteria | 0.02 |
| 1418 | 216463 | Lactobacillus spicheri | Bacteria | 0.02 |
| 1419 | 1715012 | Enterococcus sp. HMSC072H05 | Bacteria | 0.02 |
| 1420 | 1288120 | Anaerococcus senegalensis | Bacteria | 0.02 |
| 1421 | 539 | Eikenella corrodens | Bacteria | 0.02 |
| 1422 | 12024 | Pseudomonas phage PRR1 | Viruses | 0.02 |
| 1423 | 1195163 | Amazon lily mild mottle virus | Viruses | 0.02 |
| 1424 | 1923170 | Hubei polero-like virus 2 | Viruses | 0.02 |
| 1425 | 154339 | Little cherry virus 2 | Viruses | 0.02 |
| 1426 | 179636 | Alicycliphilus denitrificans | Bacteria | 0.02 |
| 1427 | 39804 | Escherichia virus FI | Viruses | 0.02 |
| 1428 | 246144 | Enterococcus italicus | Bacteria | 0.02 |
| 1429 | 685899 | Papaya lethal yellowing virus | Viruses | 0.02 |
| 1430 | 312295 | Cotton leafroll dwarf virus | Viruses | 0.02 |
| 1431 | 1739439 | Corynebacterium sp. HMSC076D02 | Bacteria | 0.02 |
| 1432 | 1493 | Clostridium cellulovorans | Bacteria | 0.02 |
| 1433 | 796937 | Peptoanaerobacter stomatis | Bacteria | 0.02 |
| 1434 | 470565 | Prevotella histicola | Bacteria | 0.02 |
| 1435 | 451457 | Lactococcus chungangensis | Bacteria | 0.02 |
| 1436 | 1229753 | Escherichia phage phAPEC8 | Viruses | 0.02 |

| | | | | |
|---|---|---|---|---|
| 1437 | 1581074 | Granulicatella sp. HMSC31F03 | Bacteria | 0.02 |
| 1438 | 1581071 | Granulicatella sp. HMSC30F09 | Bacteria | 0.02 |
| 1439 | 425941 | Prevotella nanceiensis | Bacteria | 0.02 |
| 1440 | 464322 | Veillonella magna | Bacteria | 0.02 |
| 1441 | 425010 | Botryotinia fuckeliana partitivirus 1 | Viruses | 0.02 |
| 1442 | 1470356 | Clostridium ihumii | Bacteria | 0.02 |
| 1443 | 1287488 | Prevotella sp. S7 MS 2 | Bacteria | 0.02 |
| 1444 | 443746 | Asparagus virus 1 | Viruses | 0.02 |
| 1445 | 157691 | Leptotrichia shahii | Bacteria | 0.02 |
| 1446 | 444193 | Botrytis cinerea mitovirus 1 | Viruses | 0.02 |
| 1447 | 29388 | Staphylococcus capitis | Bacteria | 0.02 |
| 1448 | 1385385 | Streptococcus phage TP-778L | Viruses | 0.02 |
| 1449 | 1354300 | Peptoniphilus sp. ChDC B134 | Bacteria | 0.02 |
| 1450 | 200 | Campylobacter curvus | Bacteria | 0.02 |
| 1451 | 203 | Campylobacter rectus | Bacteria | 0.02 |
| 1452 | 12049 | Soybean dwarf virus | Viruses | 0.02 |
| 1453 | 1923116 | Hubei picorna-like virus 36 | Viruses | 0.02 |
| 1454 | 10829 | Squash leaf curl virus | Viruses | 0.02 |
| 1455 | 1857568 | Macellibacteroides sp. HH-ZS | Bacteria | 0.02 |
| 1456 | 36343 | Lactococcus phage bIL67 | Viruses | 0.02 |
| 1457 | 359987 | Rhizosolenia setigera RNA virus 01 | Viruses | 0.02 |
| 1458 | 1206545 | Klebsiella phage 0507-KN2-1 | Viruses | 0.02 |
| 1459 | 1868652 | High Plains wheat mosaic virus | Viruses | 0.02 |
| 1460 | 1739450 | Actinomyces sp. HMSC062G12 | Bacteria | 0.02 |
| 1461 | 1547495 | Salivirus FHB | Viruses | 0.02 |
| 1462 | 59241 | Streptococcus phage Dp-1 | Viruses | 0.02 |
| 1463 | 156976 | Corynebacterium riegelii | Bacteria | 0.02 |
| 1464 | 1589 | Lactobacillus pentosus | Bacteria | 0.02 |
| 1465 | 1517899 | Kytococcus sp. CUA-901 | Bacteria | 0.02 |
| 1466 | 1280675 | Bifidobacterium sp. AGR2158 | Bacteria | 0.02 |
| 1467 | 1759399 | Streptococcus sp. A12 | Bacteria | 0.02 |
| 1468 | 1563222 | Citrobacter pasteurii | Bacteria | 0.02 |
| 1469 | 181675 | Lactobacillus coleohominis | Bacteria | 0.02 |
| 1470 | 357278 | Lactobacillus parabrevis | Bacteria | 0.02 |
| 1471 | 1692238 | Enterobacter sp. FY-07 | Bacteria | 0.02 |
| 1472 | 81931 | Sweet potato chlorotic stunt virus | Viruses | 0.02 |
| 1473 | 1739465 | Enterococcus sp. HMSC076E04 | Bacteria | 0.02 |
| 1474 | 179838 | Lactobacillus diolivorans | Bacteria | 0.02 |
| 1475 | 1410625 | Lachnospiraceae bacterium MD2004 | Bacteria | 0.02 |
| 1476 | 218667 | Oyster mushroom spherical virus | Viruses | 0.02 |
| 1477 | 39443 | Carnation Italian ringspot virus | Viruses | 0.02 |
| 1478 | 111105 | Porphyromonas gulae | Bacteria | 0.02 |
| 1479 | 12209 | Pepper mottle virus | Viruses | 0.02 |
| 1480 | 50948 | Enterobacteria phage RB49 | Viruses | 0.02 |
| 1481 | 381742 | Lactobacillus camelliae | Bacteria | 0.02 |
| 1482 | 485 | Neisseria gonorrhoeae | Bacteria | 0.02 |
| 1483 | 1647408 | Klebsiella phage KLPN1 | Viruses | 0.02 |
| 1484 | 693582 | Pseudomonas phage phi-2 | Viruses | 0.02 |

| | | | | |
|---|---|---|---|---|
| 1485 | 1693 | Bifidobacterium minimum | Bacteria | 0.02 |
| 1486 | 1272 | Kocuria varians | Bacteria | 0.02 |
| 1487 | 1739491 | Streptococcus sp. HMSC067H01 | Bacteria | 0.02 |
| 1488 | 1268254 | Peptoniphilus timonensis | Bacteria | 0.02 |
| 1489 | 1445858 | Enterococcus phage IME-EFm1 | Viruses | 0.02 |
| 1490 | 641148 | Neisseria sp. oral taxon 014 | Bacteria | 0.02 |
| 1491 | 36427 | Rotavirus C | Viruses | 0.02 |
| 1492 | 1183241 | Persimmon cryptic virus | Viruses | 0.02 |
| 1493 | 1852625 | Klebsiella phage vB_KpnM_KpV477 | Viruses | 0.02 |
| 1494 | 202789 | Actinobaculum massiliense | Bacteria | 0.02 |
| 1495 | 1485952 | Enterobacter massiliensis | Bacteria | 0.02 |
| 1496 | 1965376 | Escherichia virus EC6 | Viruses | 0.02 |
| 1497 | 36015 | Pichia kluyveri | Eukaryota | 0.02 |
| 1498 | 134821 | Ureaplasma parvum | Bacteria | 0.02 |
| 1499 | 61435 | Dehalococcoides mccartyi | Bacteria | 0.02 |
| 1500 | 375175 | Lactobacillus backii | Bacteria | 0.02 |
| 1501 | 632112 | Lactobacillus phage Lb338-1 | Viruses | 0.02 |
| 1502 | 43131 | Tissierella praeacuta | Bacteria | 0.02 |
| 1503 | 424716 | Salmonella phage Vi II-E1 | Viruses | 0.02 |
| 1504 | 150055 | Streptococcus lutetiensis | Bacteria | 0.02 |
| 1505 | 1908263 | Rodentibacter trehalosifermentans | Bacteria | 0.02 |
| 1506 | 105219 | Ralstonia mannitolilytica | Bacteria | 0.02 |
| 1507 | 33934 | Anoxybacillus flavithermus | Bacteria | 0.02 |
| 1508 | 435910 | Franconibacter pulveris | Bacteria | 0.02 |
| 1509 | 12179 | Foxtail mosaic virus | Viruses | 0.02 |
| 1510 | 689781 | Oribacterium sp. NK2B42 | Bacteria | 0.02 |
| 1511 | 1922725 | Beihai tombus-like virus 4 | Viruses | 0.02 |
| 1512 | 344022 | Escherichia virus K1E | Viruses | 0.02 |
| 1513 | 33905 | Bifidobacterium thermophilum | Bacteria | 0.02 |
| 1514 | 1654357 | La Jolla virus | Viruses | 0.02 |
| 1515 | 1654356 | Thika virus | Viruses | 0.02 |
| 1516 | 481720 | Lactobacillus otakiensis | Bacteria | 0.02 |
| 1517 | 1720495 | Escherichia phage slur16 | Viruses | 0.02 |
| 1518 | 5412 | Cystofilobasidium capitatum | Eukaryota | 0.02 |
| 1519 | 129875 | Human mastadenovirus A | Viruses | 0.02 |
| 1520 | 1756832 | Phasey bean mild yellows virus | Viruses | 0.02 |
| 1521 | 1906334 | Corynebacterium sp. NML140438 | Bacteria | 0.02 |
| 1522 | 753670 | Pea necrotic yellow dwarf virus | Viruses | 0.02 |
| 1523 | 53655 | Pichia fermentans | Eukaryota | 0.02 |
| 1524 | 1032457 | Passion fruit mosaic virus | Viruses | 0.02 |
| 1525 | 585 | Proteus vulgaris | Bacteria | 0.02 |
| 1526 | 580 | Kluyvera cryocrescens | Bacteria | 0.02 |
| 1527 | 1177630 | American hop latent virus | Viruses | 0.02 |
| 1528 | 1111137 | Slackia sp. CM382 | Bacteria | 0.02 |
| 1529 | 1349 | Streptococcus uberis | Bacteria | 0.02 |
| 1530 | 1509 | Clostridium sporogenes | Bacteria | 0.02 |
| 1531 | 351495 | Raphanus sativus cryptic virus 2 | Viruses | 0.02 |
| 1532 | 52768 | Actinomyces georgiae | Bacteria | 0.02 |

| | | | | |
|---|---|---|---|---|
| 1533 | 373058 | Tomato bushy stunt virus satellite RNA | Viruses | 0.02 |
| 1534 | 227507 | Strawberry pallidosis-associated virus | Viruses | 0.02 |
| 1535 | 1475062 | Porcine stool-associated circular virus 4 | Viruses | 0.02 |
| 1536 | 54291 | Raoultella ornithinolytica | Bacteria | 0.02 |
| 1537 | 1889813 | Anaerolineaceae bacterium oral taxon 439 | Bacteria | 0.02 |
| 1538 | 1287640 | Anaerococcus obesiensis | Bacteria | 0.02 |
| 1539 | 671224 | Selenomonas artemidis | Bacteria | 0.02 |
| 1540 | 1785087 | Candidatus Protochlamydia sp. W-9 | Bacteria | 0.02 |
| 1541 | 1517 | Thermoanaerobacterium thermosaccharolyticum | Bacteria | 0.02 |
| 1542 | 936577 | Streptococcus sp. AS14 | Bacteria | 0.02 |
| 1543 | 1537165 | Porcine stool-associated circular virus 6 | Viruses | 0.02 |
| 1544 | 1923554 | Wenzhou bivalvia virus 2 | Viruses | 0.02 |
| 1545 | 1631 | Fructobacillus fructosus | Bacteria | 0.02 |
| 1546 | 229549 | Streptococcus minor | Bacteria | 0.02 |
| 1547 | 27288 | Naumovozyma castellii | Eukaryota | 0.01 |
| 1548 | 1211480 | Persimmon virus A | Viruses | 0.01 |
| 1549 | 439016 | Marine RNA virus JP-B | Viruses | 0.01 |
| 1550 | 407975 | Prevotella pleuritidis | Bacteria | 0.01 |
| 1551 | 563038 | Streptococcus sp. M334 | Bacteria | 0.01 |
| 1552 | 1408894 | Red clover cryptic virus 1 | Viruses | 0.01 |
| 1553 | 60133 | Prevotella pallens | Bacteria | 0.01 |
| 1554 | 448384 | Enterobacteria phage Phi1 | Viruses | 0.01 |
| 1555 | 1781 | Mycobacterium marinum | Bacteria | 0.01 |
| 1556 | 113574 | Hyphomicrobium sp. GJ21 | Bacteria | 0.01 |
| 1557 | 1656 | Actinomyces viscosus | Bacteria | 0.01 |
| 1558 | 656024 | Frankia symbiont of Datisca glomerata | Bacteria | 0.01 |
| 1559 | 1805471 | Clostridium sp. Marseille-P2415 | Bacteria | 0.01 |
| 1560 | 313439 | Streptococcus massiliensis | Bacteria | 0.01 |
| 1561 | 938293 | Anaerococcus provenciensis | Bacteria | 0.01 |
| 1562 | 554 | Pectobacterium carotovorum | Bacteria | 0.01 |
| 1563 | 1739400 | Corynebacterium sp. HMSC069E04 | Bacteria | 0.01 |
| 1564 | 1631871 | Weissella jogaejeotgali | Bacteria | 0.01 |
| 1565 | 1581133 | Actinomyces sp. HMSC08A09 | Bacteria | 0.01 |
| 1566 | 178214 | Facklamia hominis | Bacteria | 0.01 |
| 1567 | 336988 | Oenococcus kitaharae | Bacteria | 0.01 |
| 1568 | 1914856 | Escherichia virus FFH2 | Viruses | 0.01 |
| 1569 | 1283 | Staphylococcus haemolyticus | Bacteria | 0.01 |
| 1570 | 1814960 | Streptococcus virus 9874 | Viruses | 0.01 |
| 1571 | 43263 | Pseudomonas alcaligenes | Bacteria | 0.01 |
| 1572 | 1745712 | Anaerococcus sp. Marseille-P2143 | Bacteria | 0.01 |
| 1573 | 185008 | Butyrivibrio hungatei | Bacteria | 0.01 |
| 1574 | 656083 | Barley yellow striate mosaic virus | Viruses | 0.01 |
| 1575 | 1923133 | Hubei picorna-like virus 51 | Viruses | 0.01 |
| 1576 | 2274 | Desulfurococcus mobilis | Archaea | 0.01 |
| 1577 | 53346 | Enterococcus mundtii | Bacteria | 0.01 |
| 1578 | 1504 | Clostridium septicum | Bacteria | 0.01 |
| 1579 | 12224 | Sugarcane mosaic virus | Viruses | 0.01 |
| 1580 | 1433126 | Mucinivorans hirudinis | Bacteria | 0.01 |

| | | | | |
|---|---|---|---|---|
| 1581 | 29379 | Staphylococcus auricularis | Bacteria | 0.01 |
| 1582 | 1568973 | Botrytis cinerea RNA virus 1 | Viruses | 0.01 |
| 1583 | 435842 | Streptococcus sp. C150 | Bacteria | 0.01 |
| 1584 | 33936 | Aeribacillus pallidus | Bacteria | 0.01 |
| 1585 | 1965385 | Escherichia virus wV8 | Viruses | 0.01 |
| 1586 | 1965384 | Erwinia virus Ea214 | Viruses | 0.01 |
| 1587 | 936563 | Fusobacterium sp. CM22 | Bacteria | 0.01 |
| 1588 | 129951 | Human mastadenovirus C | Viruses | 0.01 |
| 1589 | 713030 | Selenomonas sp. oral taxon 136 | Bacteria | 0.01 |
| 1590 | 59310 | Streptococcus macedonicus | Bacteria | 0.01 |
| 1591 | 1823756 | Actinomycetaceae bacterium BA112 | Bacteria | 0.01 |
| 1592 | 28227 | Mycoplasma penetrans | Bacteria | 0.01 |
| 1593 | 1383 | Atopobium rimae | Bacteria | 0.01 |
| 1594 | 228582 | Cereal yellow dwarf virus-RPS | Viruses | 0.01 |
| 1595 | 40091 | Helcococcus kunzii | Bacteria | 0.01 |
| 1596 | 1715007 | Rothia sp. HMSC071B01 | Bacteria | 0.01 |
| 1597 | 1169350 | Citrobacter sp. KTE32 | Bacteria | 0.01 |
| 1598 | 150285 | Garlic virus E | Viruses | 0.01 |
| 1599 | 1219585 | Arcanobacterium sp. S3PF19 | Bacteria | 0.01 |
| 1600 | 31973 | Eggerthia catenaformis | Bacteria | 0.01 |
| 1601 | 77635 | Bifidobacterium subtile | Bacteria | 0.01 |
| 1602 | 1608993 | Pseudomonas sp. DSM 28140 | Bacteria | 0.01 |
| 1603 | 2424 | Fervidobacterium nodosum | Bacteria | 0.01 |
| 1604 | 1675607 | Klebsiella phage Matisse | Viruses | 0.01 |
| 1605 | 94009 | Thermicanus aegyptius | Bacteria | 0.01 |
| 1606 | 204042 | Dickeya zeae | Bacteria | 0.01 |
| 1607 | 1922438 | Beihai narna-like virus 11 | Viruses | 0.01 |
| 1608 | 12215 | Potato virus A | Viruses | 0.01 |
| 1609 | 936554 | Campylobacter sp. FOBRC14 | Bacteria | 0.01 |
| 1610 | 1602 | Lactobacillus alimentarius | Bacteria | 0.01 |
| 1611 | 1607 | Lactobacillus bifermentans | Bacteria | 0.01 |
| 1612 | 1739310 | Turicella sp. HMSC076G08 | Bacteria | 0.01 |
| 1613 | 1739315 | Globicatella sp. HMSC072A10 | Bacteria | 0.01 |
| 1614 | 1566990 | Streptococcus phage SpSL1 | Viruses | 0.01 |
| 1615 | 388452 | Lactococcus phage KSY1 | Viruses | 0.01 |
| 1616 | 1888167 | Enterobacter sp. ku-bf2 | Bacteria | 0.01 |
| 1617 | 37206 | Helicoverpa armigera stunt virus | Viruses | 0.01 |
| 1618 | 1551 | Clostridium aurantibutyricum | Bacteria | 0.01 |
| 1619 | 1914894 | Escherichia virus 4MG | Viruses | 0.01 |
| 1620 | 1870933 | Enterobacter cloacae complex sp. 20432 | Bacteria | 0.01 |
| 1621 | 368736 | Maracuja mosaic virus | Viruses | 0.01 |
| 1622 | 132477 | Kalanchoe latent virus | Viruses | 0.01 |
| 1623 | 1715019 | Enterococcus sp. HMSC064A12 | Bacteria | 0.01 |
| 1624 | 1542743 | Caribou feces-associated gemycircularvirus | Viruses | 0.01 |
| 1625 | 130309 | Human mastadenovirus F | Viruses | 0.01 |
| 1626 | 253702 | Opuntia virus X | Viruses | 0.01 |
| 1627 | 1519399 | Sewage-associated gemycircularvirus 4 | Viruses | 0.01 |
| 1628 | 46076 | Artichoke latent virus | Viruses | 0.01 |

| | | | | |
|---|---|---|---|---|
| 1629 | 307486 | Tepidimonas taiwanensis | Bacteria | 0.01 |
| 1630 | 328430 | Chickpea chlorotic stunt virus | Viruses | 0.01 |
| 1631 | 246432 | Staphylococcus equorum | Bacteria | 0.01 |
| 1632 | 1567453 | Lactobacillus phage LfeSau | Viruses | 0.01 |
| 1633 | 81947 | Vagococcus lutrae | Bacteria | 0.01 |
| 1634 | 12203 | Maize dwarf mosaic virus | Viruses | 0.01 |
| 1635 | 1329838 | Enterobacter sp. BIDMC 26 | Bacteria | 0.01 |
| 1636 | 148604 | Lactobacillus ingluviei | Bacteria | 0.01 |
| 1637 | 470166 | Tomato marchitez virus | Viruses | 0.01 |
| 1638 | 1739430 | Streptococcus sp. HMSC078D09 | Bacteria | 0.01 |
| 1639 | 1495 | Clostridium cylindrosporum | Bacteria | 0.01 |
| 1640 | 129395 | Botrytis virus F | Viruses | 0.01 |
| 1641 | 2094 | Mycoplasma arginini | Bacteria | 0.01 |
| 1642 | 208962 | Escherichia albertii | Bacteria | 0.01 |
| 1643 | 157228 | Jeotgalicoccus psychrophilus | Bacteria | 0.01 |
| 1644 | 123841 | Helicobacter canadensis | Bacteria | 0.01 |
| 1645 | 37961 | Atkinsonella hypoxylon virus | Viruses | 0.01 |
| 1646 | 1335616 | Lactobacillus wasatchensis | Bacteria | 0.01 |
| 1647 | 74381 | Undaria pinnatifida | Eukaryota | 0.01 |
| 1648 | 1581075 | Neisseria sp. HMSC31F04 | Bacteria | 0.01 |
| 1649 | 1581072 | Corynebacterium sp. HMSC30G07 | Bacteria | 0.01 |
| 1650 | 1930302 | Chicken stool-associated circular virus | Viruses | 0.01 |
| 1651 | 1871034 | Propionimicrobium sp. Marseille-P3275 | Bacteria | 0.01 |
| 1652 | 160454 | Enterococcus pallens | Bacteria | 0.01 |
| 1653 | 688701 | Chiltepin yellow mosaic virus | Viruses | 0.01 |
| 1654 | 256701 | Glutamicibacter arilaitensis | Bacteria | 0.01 |
| 1655 | 111970 | Kyuri green mottle mosaic virus | Viruses | 0.01 |
| 1656 | 641487 | Lactococcus phage P087 | Viruses | 0.01 |
| 1657 | 1232427 | Corynebacterium ihumii | Bacteria | 0.01 |
| 1658 | 747 | Pasteurella multocida | Bacteria | 0.01 |
| 1659 | 1540094 | Citrobacter phage Moogle | Viruses | 0.01 |
| 1660 | 43305 | Butyrivibrio proteoclasticus | Bacteria | 0.01 |
| 1661 | 440518 | Sphingobium lucknowense | Bacteria | 0.01 |
| 1662 | 10726 | Escherichia virus T5 | Viruses | 0.01 |
| 1663 | 1923725 | Wuhan insect virus 21 | Viruses | 0.01 |
| 1664 | 2718 | Cardiobacterium hominis | Bacteria | 0.01 |
| 1665 | 712976 | Lachnospiraceae bacterium oral taxon 082 | Bacteria | 0.01 |
| 1666 | 1033736 | Brevibacterium senegalense | Bacteria | 0.01 |
| 1667 | 936578 | Streptococcus sp. AS20 | Bacteria | 0.01 |
| 1668 | 36809 | Mycobacterium abscessus | Bacteria | 0.01 |
| 1669 | 936572 | Selenomonas sp. FOBRC6 | Bacteria | 0.01 |
| 1670 | 1386092 | Cellulomonas carbonis | Bacteria | 0.01 |
| 1671 | 108486 | Corynebacterium falsenii | Bacteria | 0.01 |
| 1672 | 1581089 | Corynebacterium sp. HMSC11E11 | Bacteria | 0.01 |
| 1673 | 327277 | Bifidobacterium crudilactis | Bacteria | 0.01 |
| 1674 | 53422 | Thermobrachium celere | Bacteria | 0.01 |
| 1675 | 456999 | Rhizoctonia solani | Eukaryota | 0.01 |
| 1676 | 1770265 | Alfalfa enamovirus-1 | Viruses | 0.01 |

| | | | | |
|---|---|---|---|---|
| 1677 | 1930269 | Bermuda grass latent virus | Viruses | 0.01 |
| 1678 | 129143 | Cherry necrotic rusty mottle virus | Viruses | 0.01 |
| 1679 | 709323 | Fructobacillus tropaeoli | Bacteria | 0.01 |
| 1680 | 56689 | Mycobacterium mucogenicum | Bacteria | 0.01 |
| 1681 | 1640536 | Arthrospira sp. TJSD091 | Bacteria | 0.01 |
| 1682 | 295090 | Olive mild mosaic virus | Viruses | 0.01 |
| 1683 | 1581067 | Kytococcus sp. HMSC28H12 | Bacteria | 0.01 |
| 1684 | 1581069 | Corynebacterium sp. HMSC29G08 | Bacteria | 0.01 |
| 1685 | 622 | Shigella dysenteriae | Bacteria | 0.01 |
| 1686 | 1871025 | Ndongobacter massiliensis | Bacteria | 0.01 |
| 1687 | 1381007 | Grapevine red-blotch associated virus | Viruses | 0.01 |
| 1688 | 147709 | Carnobacterium inhibens | Bacteria | 0.01 |
| 1689 | 53413 | Xanthomonas axonopodis | Bacteria | 0.01 |
| 1690 | 61651 | Serratia ficaria | Bacteria | 0.01 |
| 1691 | 72520 | Nannochloropsis gaditana | Eukaryota | 0.01 |
| 1692 | 1545701 | Lactobacillus sp. wkB10 | Bacteria | 0.01 |
| 1693 | 1923266 | Hubei tombus-like virus 2 | Viruses | 0.01 |
| 1694 | 1119528 | Methyloversatilis discipulorum | Bacteria | 0.01 |
| 1695 | 154621 | Enterococcus phoeniculicola | Bacteria | 0.01 |
| 1696 | 1303256 | Sphingobium sp. DC-2 | Bacteria | 0.01 |
| 1697 | 201 | Campylobacter lari | Bacteria | 0.01 |
| 1698 | 1879023 | Mycobacterium sp. djl-10 | Bacteria | 0.01 |
| 1699 | 1923604 | Wenzhou picorna-like virus 2 | Viruses | 0.01 |
| 1700 | 135080 | Selenomonas flueggei | Bacteria | 0.01 |
| 1701 | 12188 | White clover mosaic virus | Viruses | 0.01 |
| 1702 | 364298 | Janibacter hoylei | Bacteria | 0.01 |
| 1703 | 1527519 | Escherichia phage Av-05 | Viruses | 0.01 |
| 1704 | 2210 | Methanosarcina thermophila | Archaea | 0.01 |
| 1705 | 1686381 | Citrobacter sp. MGH106 | Bacteria | 0.01 |
| 1706 | 936562 | Fusobacterium sp. CM21 | Bacteria | 0.01 |
| 1707 | 12268 | Carnation ringspot virus | Viruses | 0.01 |
| 1708 | 10752 | Escherichia phage N4 | Viruses | 0.01 |
| 1709 | 12267 | Red clover necrotic mosaic virus | Viruses | 0.01 |
| 1710 | 44562 | Pothos latent virus | Viruses | 0.01 |
| 1711 | 4929 | Meyerozyma guilliermondii | Eukaryota | 0.01 |
| 1712 | 303541 | Lactobacillus apis | Bacteria | 0.01 |
| 1713 | 269666 | Streptococcus marimammalium | Bacteria | 0.01 |
| 1714 | 78344 | Bifidobacterium gallinarum | Bacteria | 0.01 |
| 1715 | 83554 | Chlamydia psittaci | Bacteria | 0.01 |
| 1716 | 156978 | Corynebacterium imitans | Bacteria | 0.01 |
| 1717 | 1776109 | Goose dicistrovirus | Viruses | 0.01 |
| 1718 | 29394 | Dolosigranulum pigrum | Bacteria | 0.01 |
| 1719 | 633 | Yersinia pseudotuberculosis | Bacteria | 0.01 |
| 1720 | 1090134 | Salmonella phage SPN3US | Viruses | 0.01 |
| 1721 | 1549858 | Sphingomonas taxi | Bacteria | 0.01 |
| 1722 | 1505227 | Aeromonas phage pAh6-C | Viruses | 0.01 |
| 1723 | 1229751 | Lactococcus phage BM13 | Viruses | 0.01 |
| 1724 | 1100043 | Apis mellifera filamentous virus | Viruses | 0.01 |

| | | | | |
|---|---|---|---|---|
| 1725 | 12465 | Barley yellow mosaic virus | Viruses | 0.01 |
| 1726 | 61648 | Kluyvera intermedia | Bacteria | 0.01 |
| 1727 | 1193095 | Lactobacillus hokkaidonensis | Bacteria | 0.01 |
| 1728 | 1920759 | Escherichia virus IME11 | Viruses | 0.01 |
| 1729 | 1795832 | Eikenella sp. NML130454 | Bacteria | 0.01 |
| 1730 | 55211 | Erwinia persicina | Bacteria | 0.01 |
| 1731 | 55507 | Schwartzia succinivorans | Bacteria | 0.01 |
| 1732 | 712119 | Actinomyces sp. oral taxon 175 | Bacteria | 0.01 |
| 1733 | 4896 | Schizosaccharomyces pombe | Eukaryota | 0.01 |
| 1734 | 1686394 | Enterobacter sp. MGH128 | Bacteria | 0.01 |
| 1735 | 94625 | Ochrobactrum intermedium | Bacteria | 0.01 |
| 1736 | 1235479 | Pelosinus sp. HCF1 | Bacteria | 0.01 |
| 1737 | 1912598 | Cherry associated luteovirus | Viruses | 0.01 |
| 1738 | 400946 | Wohlfahrtiimonas chitiniclastica | Bacteria | 0.01 |
| 1739 | 1768771 | Streptococcus sp. CCH8-G7 | Bacteria | 0.01 |
| 1740 | 1739264 | Corynebacterium sp. HMSC065D07 | Bacteria | 0.01 |
| 1741 | 752 | Pasteurella bettyae | Bacteria | 0.01 |
| 1742 | 750 | Gallibacterium anatis | Bacteria | 0.01 |
| 1743 | 45972 | Staphylococcus pasteuri | Bacteria | 0.01 |
| 1744 | 1770210 | Micrococcus sp. CH7 | Bacteria | 0.01 |
| 1745 | 267364 | Lactobacillus acidifarinae | Bacteria | 0.01 |
| 1746 | 56407 | Hanseniaspora occidentalis | Eukaryota | 0.01 |
| 1747 | 646010 | Suakwa aphid-borne yellows virus | Viruses | 0.01 |
| 1748 | 1462681 | Pepo aphid-borne yellows virus | Viruses | 0.01 |
| 1749 | 1462682 | Luffa aphid-borne yellows virus | Viruses | 0.01 |
| 1750 | 32629 | Indian peanut clump virus | Viruses | 0.01 |
| 1751 | 43771 | Corynebacterium urealyticum | Bacteria | 0.01 |
| 1752 | 28264 | Arcanobacterium haemolyticum | Bacteria | 0.01 |
| 1753 | 198 | Campylobacter hyointestinalis | Bacteria | 0.01 |
| 1754 | 1907766 | Pseudomonas sp. BS-2016 | Bacteria | 0.01 |
| 1755 | 1156431 | Streptococcus sp. I-G2 | Bacteria | 0.01 |
| 1756 | 46436 | Beet soil-borne virus | Viruses | 0.01 |
| 1757 | 1220025 | Pokeweed mosaic virus | Viruses | 0.01 |
| 1758 | 1325933 | Clostridium polynesiense | Bacteria | 0.01 |
| 1759 | 438780 | Lactobacillus phage phiPYB5 | Viruses | 0.01 |
| 1760 | 2162 | Methanobacterium formicicum | Archaea | 0.01 |
| 1761 | 1933261 | Groundnut bud necrosis tospovirus | Viruses | 0.01 |
| 1762 | 1898961 | Kluyvera intestini | Bacteria | 0.01 |
| 1763 | 53343 | Desulfotomaculum aeronauticum | Bacteria | 0.01 |
| 1764 | 1739479 | Actinomyces sp. HMSC065F12 | Bacteria | 0.01 |
| 1765 | 1768764 | Streptococcus sp. CCH5-D3 | Bacteria | 0.01 |
| 1766 | 59803 | Sphingomonas echinoides | Bacteria | 0.01 |
| 1767 | 71237 | Staphylococcus vitulinus | Bacteria | 0.01 |
| 1768 | 78541 | Streptococcus phage Sfi11 | Viruses | 0.01 |
| 1769 | 1714265 | Klebsiella sp. KGM-IMP216 | Bacteria | 0.01 |
| 1770 | 264076 | Horseradish latent virus | Viruses | 0.01 |
| 1771 | 1661745 | Haemophilus sp. C1 | Bacteria | 0.01 |
| 1772 | 28447 | Clavibacter michiganensis | Bacteria | 0.01 |

| | | | | |
|---|---|---|---|---|
| 1773 | 1222338 | Enterobacteria phage GEC-3S | Viruses | 0.01 |
| 1774 | 1739496 | Prevotella sp. HMSC069G02 | Bacteria | 0.01 |
| 1775 | 1581143 | Arthrobacter sp. HMSC08H08 | Bacteria | 0.01 |
| 1776 | 1280676 | Butyrivibrio sp. WCD3002 | Bacteria | 0.01 |
| 1777 | 28042 | Saccharopolyspora rectivirgula | Bacteria | 0.01 |
| 1778 | 1622070 | Paenibacillus sp. GM2 | Bacteria | 0.01 |
| 1779 | 12402 | Streptococcus phage EJ-1 | Viruses | 0.01 |
| 1780 | 186538 | Zaire ebolavirus | Viruses | 0.01 |
| 1781 | 51680 | Ribgrass mosaic virus | Viruses | 0.01 |
| 1782 | 1541211 | Cripavirus NB-1/2011/HUN | Viruses | 0.01 |
| 1783 | 61647 | Pluralibacter gergoviae | Bacteria | 0.01 |
| 1784 | 1055192 | Comamonas sp. B-9 | Bacteria | 0.01 |
| 1785 | 1873985 | Salmonella phage IME207 | Viruses | 0.01 |
| 1786 | 1923094 | Hubei picorna-like virus 15 | Viruses | 0.01 |
| 1787 | 537874 | Streptococcus phage PH15 | Viruses | 0.01 |
| 1788 | 1920779 | Klebsiella virus SU552A | Viruses | 0.01 |
| 1789 | 1514105 | Erysipelothrix larvae | Bacteria | 0.01 |
| 1790 | 1914861 | Enterobacter sp. Sa187 | Bacteria | 0.01 |
| 1791 | 88132 | Eremococcus coleocola | Bacteria | 0.01 |
| 1792 | 1522060 | Pantoea sp. 3.5.1 | Bacteria | 0.01 |
| 1793 | 1825924 | Barley virus G | Viruses | 0.01 |
| 1794 | 2371 | Xylella fastidiosa | Bacteria | 0.01 |
| 1795 | 1195085 | Cronobacter phage CR5 | Viruses | 0.01 |
| 1796 | 762210 | Bifidobacterium saguini | Bacteria | 0.01 |
| 1797 | 665550 | Dietzia alimentaria | Bacteria | 0.01 |
| 1798 | 12348 | Lactobacillus phage LL-H | Viruses | 0.01 |
| 1799 | 694003 | Betacoronavirus 1 | Viruses | 0.01 |
| 1800 | 317010 | Enterococcus canintestini | Bacteria | 0.01 |
| 1801 | 59749 | Maize rayado fino virus | Viruses | 0.01 |
| 1802 | 53345 | Enterococcus durans | Bacteria | 0.01 |
| 1803 | 1206110 | Lactobacillus phage phiAQ113 | Viruses | 0.01 |
| 1804 | 1161906 | Weissella phage phiYS61 | Viruses | 0.01 |
| 1805 | 1229204 | alpha proteobacterium L41A | Bacteria | 0.01 |
| 1806 | 114090 | Pediococcus inopinatus | Bacteria | 0.01 |
| 1807 | 1675603 | Citrobacter phage Michonne | Viruses | 0.01 |
| 1808 | 1233383 | Human cosavirus | Viruses | 0.01 |
| 1809 | 255238 | Fragaria chiloensis latent virus | Viruses | 0.01 |
| 1810 | 1849383 | Psychrobacter sp. SHUES1 | Bacteria | 0.01 |
| 1811 | 264483 | Phaffia rhodozyma | Eukaryota | 0.01 |
| 1812 | 158836 | Enterobacter hormaechei | Bacteria | 0.01 |
| 1813 | 100468 | Lactobacillus perolens | Bacteria | 0.01 |
| 1814 | 712535 | Selenomonas sp. oral taxon 149 | Bacteria | 0.01 |
| 1815 | 11203 | Human parainfluenza virus 4 | Viruses | 0.01 |
| 1816 | 35289 | Grapevine virus B | Viruses | 0.01 |
| 1817 | 1873990 | Escherichia phage vB_EcoM_Alf5 | Viruses | 0.01 |
| 1818 | 93466 | Fervidobacterium pennivorans | Bacteria | 0.01 |
| 1819 | 563037 | Streptococcus sp. M143 | Bacteria | 0.01 |
| 1820 | 1499685 | Bacillus andreraoultii | Bacteria | 0.01 |

| | | | | |
|---|---|---|---|---|
| 1821 | 393921 | Porphyromonas crevioricanis | Bacteria | 0.01 |
| 1822 | 81857 | Lactobacillus selangorensis | Bacteria | 0.01 |
| 1823 | 176291 | Lactobacillus vaccinostercus | Bacteria | 0.01 |
| 1824 | 1129192 | Bacillus phage BCP8-2 | Viruses | 0.01 |
| 1825 | 481719 | Lactobacillus sunkii | Bacteria | 0.01 |
| 1826 | 1768743 | Blastomonas sp. CCH8-A3 | Bacteria | 0.01 |
| 1827 | 5207 | Cryptococcus neoformans | Eukaryota | 0.01 |
| 1828 | 1647391 | Streptococcus phage APCM01 | Viruses | 0.01 |
| 1829 | 1856642 | Maize yellow mosaic virus | Viruses | 0.01 |
| 1830 | 1141136 | Cronobacter phage vB_CsaM_GAP32 | Viruses | 0.01 |
| 1831 | 1552735 | Lactobacillus phage Ldl1 | Viruses | 0.01 |
| 1832 | 71451 | Enterococcus malodoratus | Bacteria | 0.01 |
| 1833 | 1481465 | Tomato necrotic dwarf virus | Viruses | 0.01 |
| 1834 | 1954380 | Sodalis virus SO1 | Viruses | 0.01 |
| 1835 | 1911010 | Escherichia virus K1H | Viruses | 0.01 |
| 1836 | 1965306 | Lagenaria siceraria endornavirus-Hubei | Viruses | 0.01 |
| 1837 | 193122 | Peanut stunt virus satellite RNA | Viruses | 0.01 |
| 1838 | 1736702 | Enterobacter sp. K66-74 | Bacteria | 0.01 |
| 1839 | 1384081 | Veillonella sp. DNF00869 | Bacteria | 0.01 |
| 1840 | 38305 | Corynebacterium vitaeruminis | Bacteria | 0.01 |
| 1841 | 712414 | Oribacterium sp. oral taxon 108 | Bacteria | 0.01 |
| 1842 | 73422 | Streptococcus phage TP-J34 | Viruses | 0.01 |
| 1843 | 338473 | Actinomyces virus Av1 | Viruses | 0.01 |
| 1844 | 1588750 | Clostridiales bacterium KA00134 | Bacteria | 0.01 |
| 1845 | 12294 | Pea early-browning virus | Viruses | 0.01 |
| 1846 | 1379702 | Methanobacterium sp. MB1 | Archaea | 0.01 |
| 1847 | 1414721 | Clostridium jeddahense | Bacteria | 0.01 |
| 1848 | 187978 | Peru tomato mosaic virus | Viruses | 0.01 |
| 1849 | 99480 | Tetrasphaera australiensis | Bacteria | 0.01 |
| 1850 | 11988 | Turnip crinkle virus | Viruses | 0.01 |
| 1851 | 596085 | Prevotella aurantiaca | Bacteria | 0.01 |
| 1852 | 48296 | Acinetobacter pittii | Bacteria | 0.01 |
| 1853 | 746033 | Piscicoccus intestinalis | Bacteria | 0.01 |
| 1854 | 1118963 | Arthrobacter sp. Rue61a | Bacteria | 0.01 |
| 1855 | 1681197 | Arthrobacter sp. RIT-PI-e | Bacteria | 0.01 |
| 1856 | 1293441 | Lysinibacillus contaminans | Bacteria | 0.01 |
| 1857 | 1196034 | Klebsiella sp. 10982 | Bacteria | 0.01 |
| 1858 | 1739536 | Corynebacterium sp. HMSC073D01 | Bacteria | 0.01 |
| 1859 | 54289 | Vicia cryptic virus | Viruses | 0.01 |
| 1860 | 1439319 | Citrobacter sp. MGH 55 | Bacteria | 0.01 |
| 1861 | 326202 | Vanilla distortion mosaic virus | Viruses | 0.01 |
| 1862 | 83655 | Leclercia adecarboxylata | Bacteria | 0.01 |
| 1863 | 1336 | Streptococcus equi | Bacteria | 0.01 |
| 1864 | 1520 | Clostridium beijerinckii | Bacteria | 0.01 |
| 1865 | 1673719 | Anaerococcus sp. SB3 | Bacteria | 0.01 |
| 1866 | 1051676 | Erwinia phage vB_EamM-Y2 | Viruses | 0.01 |
| 1867 | 1416026 | Yellow tailflower mild mottle virus | Viruses | 0.01 |
| 1868 | 244366 | Klebsiella variicola | Bacteria | 0.01 |

| | | | | |
|---|---|---|---|---|
| 1869 | 576789 | Enterobacteria phage JSE | Viruses | 0.01 |
| 1870 | 5082 | Penicillium roqueforti | Eukaryota | 0.01 |
| 1871 | 277944 | Human coronavirus NL63 | Viruses | 0.01 |
| 1872 | 142843 | Hop mosaic virus | Viruses | 0.01 |
| 1873 | 1477000 | Peptoniphilus sp. DNF00840 | Bacteria | 0.01 |
| 1874 | 489828 | Enterobacteria phage WA13 sensu lato | Viruses | 0.01 |
| 1875 | 1235640 | Enterobacteria phage M | Viruses | 0.01 |
| 1876 | 5755 | Acanthamoeba castellanii | Eukaryota | 0.01 |
| 1877 | 134533 | Acinetobacter parvus | Bacteria | 0.01 |
| 1878 | 12178 | Cymbidium mosaic virus | Viruses | 0.01 |
| 1879 | 633135 | Streptococcus phage Abc2 | Viruses | 0.01 |
| 1880 | 1182762 | Enterococcus sp. C1 | Bacteria | 0.01 |
| 1881 | 1813769 | Salmonella phage 64795_sal3 | Viruses | 0.01 |
| 1882 | 871203 | Caballeronia zhejiangensis | Bacteria | 0.01 |
| 1883 | 37636 | Thermus scotoductus | Bacteria | 0.01 |
| 1884 | 712538 | Selenomonas sp. oral taxon 478 | Bacteria | 0.01 |
| 1885 | 1203573 | Propionibacterium sp. KPL1844 | Bacteria | 0.01 |
| 1886 | 62059 | Shallot yellow stripe virus | Viruses | 0.01 |
| 1887 | 72000 | Kocuria rhizophila | Bacteria | 0.01 |
| 1888 | 1341 | Streptococcus ratti | Bacteria | 0.01 |
| 1889 | 1768770 | Caulobacter sp. CCH5-E12 | Bacteria | 0.01 |
| 1890 | 759620 | Weissella ceti | Bacteria | 0.01 |
| 1891 | 81464 | Anaeromusa acidaminophila | Bacteria | 0.01 |
| 1892 | 1218493 | Lactobacillus kullabergensis | Bacteria | 0.01 |
| 1893 | 1840644 | Psophocarpus tetragonolobus endornavirus | Viruses | 0.01 |
| 1894 | 566 | Escherichia vulneris | Bacteria | 0.01 |
| 1895 | 370833 | Tomato torrado virus | Viruses | 0.01 |
| 1896 | 51663 | Pediococcus damnosus | Bacteria | 0.01 |
| 1897 | 51664 | Lactobacillus dextrinicus | Bacteria | 0.01 |
| 1898 | 1923308 | Hubei toti-like virus 2 | Viruses | 0.01 |
| 1899 | 302449 | Corynebacterium tuscaniense | Bacteria | 0.01 |
| 1900 | 936594 | Lachnoanaerobaculum sp. ICM7 | Bacteria | 0.01 |
| 1901 | 1883202 | Escherichia phage Gluttony | Viruses | 0.01 |
| 1902 | 1739351 | Corynebacterium sp. HMSC074C03 | Bacteria | 0.01 |
| 1903 | 33010 | Cutibacterium avidum | Bacteria | 0.01 |
| 1904 | 33011 | Cutibacterium granulosum | Bacteria | 0.01 |
| 1905 | 37662 | Brettanomyces anomalus | Eukaryota | 0.01 |
| 1906 | 528209 | Lactobacillus kimchicus | Bacteria | 0.01 |
| 1907 | 1817674 | Geobacillus sp. 8 | Bacteria | 0.01 |
| 1908 | 1200547 | Prevotella sp. RM4 | Bacteria | 0.01 |
| 1909 | 1381464 | Blackberry vein banding associated virus | Viruses | 0.01 |
| 1910 | 577 | Raoultella terrigena | Bacteria | 0.01 |
| 1911 | 187764 | Escherichia virus K1-5 | Viruses | 0.01 |
| 1912 | 645687 | Astrovirus VA1 | Viruses | 0.01 |
| 1913 | 1581113 | Corynebacterium sp. HMSC05C01 | Bacteria | 0.01 |
| 1914 | 267633 | Lactobacillus hammesii | Bacteria | 0.01 |
| 1915 | 92444 | Acute bee paralysis virus | Viruses | 0.01 |
| 1916 | 979982 | Leuconostoc sp. C2 | Bacteria | 0.01 |

| | | | | |
|---|---|---|---|---|
| 1917 | 1513 | Clostridium tetani | Bacteria | 0.01 |
| 1918 | 1715164 | Streptococcus sp. HMSC074F05 | Bacteria | 0.01 |
| 1919 | 222805 | Mycobacterium chimaera | Bacteria | 0.01 |
| 1920 | 1462608 | Pseudomonas phage KPP25 | Viruses | 0.01 |
| 1921 | 1169321 | Escherichia sp. KTE114 | Bacteria | 0.01 |
| 1922 | 430606 | Polygonum ringspot tospovirus | Viruses | 0.01 |
| 1923 | 77775 | Salmonella phage FelixO1 | Viruses | 0.01 |
| 1924 | 1293 | Staphylococcus gallinarum | Bacteria | 0.01 |
| 1925 | 1768759 | Bradyrhizobium sp. CCH4-A6 | Bacteria | 0.01 |
| 1926 | 1922578 | Beihai picorna-like virus 35 | Viruses | 0.01 |
| 1927 | 84136 | Gemella bergeri | Bacteria | 0.01 |
| 1928 | 1406816 | Zucchini tigre mosaic virus | Viruses | 0.01 |
| 1929 | 49118 | Candidatus Arthromitus sp. SFB-mouse | Bacteria | 0.01 |
| 1930 | 1739611 | Lactobacillus phage iLp1308 | Viruses | 0.01 |
| 1931 | 473784 | Opium poppy mosaic virus | Viruses | 0.01 |
| 1932 | 255066 | Pepper veinal mottle virus | Viruses | 0.01 |
| 1933 | 412383 | Solibacillus isronensis | Bacteria | 0.01 |
| 1934 | 1639 | Listeria monocytogenes | Bacteria | 0.01 |
| 1935 | 255248 | Leuconostoc garlicum | Bacteria | 0.01 |
| 1936 | 29484 | Yersinia frederiksenii | Bacteria | 0.01 |
| 1937 | 564 | Escherichia fergusonii | Bacteria | 0.01 |
| 1938 | 168471 | Laribacter hongkongensis | Bacteria | 0.01 |
| 1939 | 1749 | Acidipropionibacterium jensenii | Bacteria | 0.01 |
| 1940 | 157268 | Helicobacter winghamensis | Bacteria | 0.01 |
| 1941 | 331679 | Pediococcus stilesii | Bacteria | 0.01 |
| 1942 | 28375 | Soil-borne wheat mosaic virus | Viruses | 0.01 |
| 1943 | 468911 | Lactobacillus hordei | Bacteria | 0.01 |
| 1944 | 151043 | Tulare apple mosaic virus | Viruses | 0.01 |
| 1945 | 390842 | Lactobacillus parafarraginis | Bacteria | 0.01 |
| 1946 | 1768792 | Erythrobacter sp. CCH5-A1 | Bacteria | 0.01 |

Table S4: All genera identified in 10,000 human stool samples

| | Taxonomy ID | Genus name | SuperKingdom | Prevalence in 10,000 samples, % |
|---|---|---|---|---|
| 1 | 1485 | Clostridium | Bacteria | 99.72 |
| 2 | 816 | Bacteroides | Bacteria | 99.57 |
| 3 | 572511 | Blautia | Bacteria | 97.62 |
| 4 | 469 | Acinetobacter | Bacteria | 97.45 |
| 5 | 1730 | Eubacterium | Bacteria | 97.16 |
| 6 | 375288 | Parabacteroides | Bacteria | 96.89 |
| 7 | 1357 | Lactococcus | Bacteria | 96.88 |
| 8 | 216851 | Faecalibacterium | Bacteria | 96.39 |
| 9 | 841 | Roseburia | Bacteria | 95.72 |
| 10 | 239759 | Alistipes | Bacteria | 95.05 |
| 11 | 1263 | Ruminococcus | Bacteria | 94.67 |
| 12 | 244127 | Anaerotruncus | Bacteria | 93.04 |
| 13 | 459786 | Oscillibacter | Bacteria | 90.88 |
| 14 | 1301 | Streptococcus | Bacteria | 90.47 |
| 15 | 189330 | Dorea | Bacteria | 88.56 |
| 16 | 1407607 | Fusicatenibacter | Bacteria | 87.22 |
| 17 | 561 | Escherichia | Bacteria | 85.92 |
| 18 | 1508657 | Ruminiclostridium | Bacteria | 84.26 |
| 19 | 1505663 | Erysipelatoclostridium | Bacteria | 83.51 |
| 20 | 33042 | Coprococcus | Bacteria | 80.66 |
| 21 | 292632 | Subdoligranulum | Bacteria | 80.59 |
| 22 | 29465 | Veillonella | Bacteria | 78.62 |
| 23 | 283168 | Odoribacter | Bacteria | 78.45 |
| 24 | 61170 | Holdemania | Bacteria | 75.80 |
| 25 | 1392389 | Intestinimonas | Bacteria | 75.67 |
| 26 | 102106 | Collinsella | Bacteria | 75.50 |
| 27 | 35832 | Bilophila | Bacteria | 74.75 |
| 28 | 12234 | Tobamovirus | Viruses | 71.58 |
| 29 | 447020 | Adlercreutzia | Bacteria | 70.39 |
| 30 | 1654 | Actinomyces | Bacteria | 69.37 |
| 31 | 577310 | Parasutterella | Bacteria | 66.65 |
| 32 | 1506553 | Lachnoclostridium | Bacteria | 65.97 |
| 33 | 644652 | Gordonibacter | Bacteria | 65.09 |
| 34 | 1501226 | Romboutsia | Bacteria | 63.54 |
| 35 | 84111 | Eggerthella | Bacteria | 61.12 |
| 36 | 239934 | Akkermansia | Bacteria | 60.64 |
| 37 | 207244 | Anaerostipes | Bacteria | 59.70 |
| 38 | 574697 | Butyricimonas | Bacteria | 59.21 |
| 39 | 195950 | Tannerella | Bacteria | 57.23 |
| 40 | 1350 | Enterococcus | Bacteria | 55.78 |
| 41 | 1678 | Bifidobacterium | Bacteria | 55.28 |
| 42 | 1378 | Gemella | Bacteria | 55.05 |
| 43 | 100883 | Coprobacillus | Bacteria | 53.50 |
| 44 | 613 | Serratia | Bacteria | 53.13 |
| 45 | 1926663 | Phocea | Bacteria | 52.64 |

| | | | | |
|---|---|---|---|---|
| 46 | 397864 | Barnesiella | Bacteria | 52.30 |
| 47 | 1432051 | Eisenbergiella | Bacteria | 50.55 |
| 48 | 28050 | Lachnospira | Bacteria | 49.17 |
| 49 | 191303 | Turicibacter | Bacteria | 47.21 |
| 50 | 838 | Prevotella | Bacteria | 46.45 |
| 51 | 946234 | Flavonifractor | Bacteria | 42.39 |
| 52 | 1506577 | Tyzzerella | Bacteria | 42.03 |
| 53 | 1870884 | Clostridioides | Bacteria | 41.79 |
| 54 | 1017280 | Pseudoflavonifractor | Bacteria | 41.18 |
| 55 | 40544 | Sutterella | Bacteria | 40.64 |
| 56 | 1649459 | Hungatella | Bacteria | 38.28 |
| 57 | 580596 | Butyricicoccus | Bacteria | 36.67 |
| 58 | 1578 | Lactobacillus | Bacteria | 36.48 |
| 59 | 1905344 | Ruthenibacterium | Bacteria | 32.05 |
| 60 | 872 | Desulfovibrio | Bacteria | 31.93 |
| 61 | 577309 | Paraprevotella | Bacteria | 30.37 |
| 62 | 1935176 | Angelakisella | Bacteria | 28.63 |
| 63 | 212742 | Morococcus | Bacteria | 28.29 |
| 64 | 1924105 | Neglecta | Bacteria | 27.48 |
| 65 | 39948 | Dialister | Bacteria | 27.40 |
| 66 | 86331 | Mogibacterium | Bacteria | 27.03 |
| 67 | 1935927 | Massilioclostridium | Bacteria | 26.41 |
| 68 | 1926556 | Emergencia | Bacteria | 26.08 |
| 69 | 264995 | Anaerofustis | Bacteria | 25.78 |
| 70 | 2172 | Methanobrevibacter | Archaea | 25.70 |
| 71 | 4930 | Saccharomyces | Eukaryota | 25.44 |
| 72 | 830 | Butyrivibrio | Bacteria | 24.25 |
| 73 | 33024 | Phascolarctobacterium | Bacteria | 23.62 |
| 74 | 420345 | Lactonifactor | Bacteria | 22.77 |
| 75 | 135858 | Catenibacterium | Bacteria | 22.04 |
| 76 | 1573535 | Holdemanella | Bacteria | 21.87 |
| 77 | 12163 | Carlavirus | Viruses | 20.89 |
| 78 | 1473205 | Senegalimassilia | Bacteria | 19.55 |
| 79 | 1380 | Atopobium | Bacteria | 19.32 |
| 80 | 724 | Haemophilus | Bacteria | 17.80 |
| 81 | 846 | Oxalobacter | Bacteria | 17.77 |
| 82 | 848 | Fusobacterium | Bacteria | 16.98 |
| 83 | 990721 | Christensenella | Bacteria | 16.65 |
| 84 | 2753 | Synergistes | Bacteria | 16.55 |
| 85 | 12316 | Ilarvirus | Viruses | 16.47 |
| 86 | 1470349 | Candidatus Stoquefichus | Bacteria | 16.10 |
| 87 | 12176 | Potexvirus | Viruses | 15.91 |
| 88 | 1243 | Leuconostoc | Bacteria | 15.85 |
| 89 | 270497 | Catabacter | Bacteria | 15.67 |
| 90 | 590 | Salmonella | Bacteria | 15.61 |
| 91 | 34072 | Variovorax | Bacteria | 15.51 |
| 92 | 1348911 | Coprobacter | Bacteria | 15.41 |
| 93 | 310748 | Endornavirus | Viruses | 15.19 |

| | | | | |
|---|---|---|---|---|
| 94 | 1940255 | Fournierella | Bacteria | 15.03 |
| 95 | 836 | Porphyromonas | Bacteria | 13.98 |
| 96 | 1924081 | Bariatricus | Bacteria | 12.65 |
| 97 | 904 | Acidaminococcus | Bacteria | 11.77 |
| 98 | 1472649 | Dielma | Bacteria | 11.73 |
| 99 | 194 | Campylobacter | Bacteria | 11.45 |
| 100 | 12967 | Blastocystis | Eukaryota | 11.37 |
| 101 | 76833 | Actinobaculum | Bacteria | 10.88 |
| 102 | 1505657 | Intestinibacter | Bacteria | 9.92 |
| 103 | 84108 | Slackia | Bacteria | 9.78 |
| 104 | 162289 | Peptoniphilus | Bacteria | 9.54 |
| 105 | 119164 | Polerovirus | Viruses | 9.33 |
| 106 | 1926672 | Prevotellamassilia | Bacteria | 8.81 |
| 107 | 248744 | Marvinbryantia | Bacteria | 8.55 |
| 108 | 906 | Megasphaera | Bacteria | 8.50 |
| 109 | 150022 | Finegoldia | Bacteria | 8.31 |
| 110 | 12195 | Potyvirus | Viruses | 8.15 |
| 111 | 1511809 | Betapartitivirus | Viruses | 8.14 |
| 112 | 1253 | Pediococcus | Bacteria | 7.61 |
| 113 | 1929045 | Traorella | Bacteria | 7.29 |
| 114 | 133925 | Olsenella | Bacteria | 7.07 |
| 115 | 1635148 | Sanguibacteroides | Bacteria | 6.59 |
| 116 | 1715798 | Levyella | Bacteria | 6.59 |
| 117 | 158846 | Megamonas | Bacteria | 6.52 |
| 118 | 165779 | Anaerococcus | Bacteria | 6.51 |
| 119 | 580024 | Enterorhabdus | Bacteria | 6.47 |
| 120 | 123375 | Solobacterium | Bacteria | 6.41 |
| 121 | 186767 | Narnavirus | Viruses | 6.19 |
| 122 | 1257 | Peptostreptococcus | Bacteria | 6.10 |
| 123 | 508459 | Cloacibacillus | Bacteria | 5.55 |
| 124 | 1573534 | Faecalitalea | Bacteria | 5.54 |
| 125 | 1080709 | Methanomassiliicoccus | Archaea | 5.25 |
| 126 | 46255 | Weissella | Bacteria | 5.13 |
| 127 | 1763 | Mycobacterium | Bacteria | 5.09 |
| 128 | 67753 | Crinivirus | Viruses | 4.87 |
| 129 | 35829 | Acetivibrio | Bacteria | 4.55 |
| 130 | 2316 | Methanosphaera | Archaea | 4.34 |
| 131 | 43996 | Catonella | Bacteria | 4.32 |
| 132 | 40276 | Trichovirus | Viruses | 4.32 |
| 133 | 12137 | Sobemovirus | Viruses | 4.29 |
| 134 | 286 | Pseudomonas | Bacteria | 4.28 |
| 135 | 1686313 | Fenollaria | Bacteria | 4.26 |
| 136 | 281915 | Curvibacter | Bacteria | 4.06 |
| 137 | 1929083 | Tidjanibacter | Bacteria | 4.05 |
| 138 | 1926659 | Mediterranea | Bacteria | 4.03 |
| 139 | 12304 | Cucumovirus | Viruses | 3.93 |
| 140 | 1935200 | Merdibacter | Bacteria | 3.87 |
| 141 | 1743 | Propionibacterium | Bacteria | 3.77 |

| | | | | |
|---|---|---|---|---|
| 142 | 129725 | Foveavirus | Viruses | 3.51 |
| 143 | 1164882 | Lachnoanaerobaculum | Bacteria | 3.45 |
| 144 | 543311 | Parvimonas | Bacteria | 3.41 |
| 145 | 1923867 | Gabonia | Bacteria | 3.26 |
| 146 | 140295 | Varicosavirus | Viruses | 2.96 |
| 147 | 28138 | Rikenella | Bacteria | 2.82 |
| 148 | 46123 | Abiotrophia | Bacteria | 2.76 |
| 149 | 12174 | Capillovirus | Viruses | 2.75 |
| 150 | 1965226 | Urmitella | Bacteria | 2.73 |
| 151 | 1716 | Corynebacterium | Bacteria | 2.66 |
| 152 | 52225 | Mitsuokella | Bacteria | 2.53 |
| 153 | 1472762 | Enorma | Bacteria | 2.41 |
| 154 | 570 | Klebsiella | Bacteria | 2.38 |
| 155 | 12141 | Tombusvirus | Viruses | 2.37 |
| 156 | 104394 | Picobirnavirus | Viruses | 2.37 |
| 157 | 2701 | Gardnerella | Bacteria | 2.35 |
| 158 | 419014 | Alloscardovia | Bacteria | 2.35 |
| 159 | 1911601 | Gammacarmovirus | Viruses | 2.31 |
| 160 | 1386 | Bacillus | Bacteria | 2.16 |
| 161 | 217160 | Ampelovirus | Viruses | 2.08 |
| 162 | 620 | Shigella | Bacteria | 2.08 |
| 163 | 12270 | Nepovirus | Viruses | 2.08 |
| 164 | 13687 | Sphingomonas | Bacteria | 2.05 |
| 165 | 5721 | Trichomonas | Eukaryota | 2.00 |
| 166 | 265975 | Oribacterium | Bacteria | 1.97 |
| 167 | 1922299 | Beduini | Bacteria | 1.96 |
| 168 | 642 | Aeromonas | Bacteria | 1.89 |
| 169 | 1511808 | Alphapartitivirus | Viruses | 1.86 |
| 170 | 1573536 | Faecalicoccus | Bacteria | 1.82 |
| 171 | 1903506 | Drancourtella | Bacteria | 1.79 |
| 172 | 1929297 | Bittarella | Bacteria | 1.68 |
| 173 | 1935188 | Libanicoccus | Bacteria | 1.66 |
| 174 | 186457 | Aureusvirus | Viruses | 1.63 |
| 175 | 48736 | Ralstonia | Bacteria | 1.58 |
| 176 | 1283313 | Alloprevotella | Bacteria | 1.46 |
| 177 | 1926651 | Culturomica | Bacteria | 1.40 |
| 178 | 11611 | Tospovirus | Viruses | 1.39 |
| 179 | 12320 | Alfamovirus | Viruses | 1.37 |
| 180 | 32207 | Rothia | Bacteria | 1.32 |
| 181 | 12300 | Bromovirus | Viruses | 1.29 |
| 182 | 156207 | Tritimovirus | Viruses | 1.28 |
| 183 | 1470353 | Candidatus Soleaferrea | Bacteria | 1.26 |
| 184 | 184869 | Varibaculum | Bacteria | 1.22 |
| 185 | 209 | Helicobacter | Bacteria | 1.16 |
| 186 | 12160 | Closterovirus | Viruses | 1.14 |
| 187 | 196081 | Scardovia | Bacteria | 1.11 |
| 188 | 1273095 | Anaerosphaera | Bacteria | 1.09 |
| 189 | 140568 | Allexivirus | Viruses | 1.04 |

| | | | | |
|---|---|---|---|---|
| 190 | 1637257 | Mageeibacillus | Bacteria | 0.94 |
| 191 | 963 | Herbaspirillum | Bacteria | 0.92 |
| 192 | 1912217 | Pseudopropionibacterium | Bacteria | 0.89 |
| 193 | 1505652 | Terrisporobacter | Bacteria | 0.87 |
| 194 | 1847725 | Lawsonella | Bacteria | 0.86 |
| 195 | 142786 | Norovirus | Viruses | 0.82 |
| 196 | 416916 | Aggregatibacter | Bacteria | 0.81 |
| 197 | 5758 | Entamoeba | Eukaryota | 0.79 |
| 198 | 2747 | Carnobacterium | Bacteria | 0.79 |
| 199 | 662 | Vibrio | Bacteria | 0.75 |
| 200 | 168808 | Sneathia | Bacteria | 0.75 |
| 201 | 13366 | Brettanomyces | Eukaryota | 0.74 |
| 202 | 156454 | Anaeroglobus | Bacteria | 0.69 |
| 203 | 2129 | Ureaplasma | Bacteria | 0.69 |
| 204 | 34104 | Streptobacillus | Bacteria | 0.69 |
| 205 | 75 | Caulobacter | Bacteria | 0.68 |
| 206 | 482 | Neisseria | Bacteria | 0.67 |
| 207 | 39760 | Idaeovirus | Viruses | 0.65 |
| 208 | 1924093 | Anaeromassilibacillus | Bacteria | 0.65 |
| 209 | 156973 | Dysgonomonas | Bacteria | 0.61 |
| 210 | 12036 | Luteovirus | Viruses | 0.61 |
| 211 | 60919 | Sanguibacter | Bacteria | 0.59 |
| 212 | 43987 | Geotrichum | Eukaryota | 0.59 |
| 213 | 194960 | Kobuvirus | Viruses | 0.58 |
| 214 | 1937664 | Criibacterium | Bacteria | 0.56 |
| 215 | 12258 | Comovirus | Viruses | 0.56 |
| 216 | 588605 | Robinsoniella | Bacteria | 0.55 |
| 217 | 1299311 | Betanecrovirus | Viruses | 0.55 |
| 218 | 117563 | Granulicatella | Bacteria | 0.54 |
| 219 | 1911312 | Gabonibacter | Bacteria | 0.54 |
| 220 | 544 | Citrobacter | Bacteria | 0.53 |
| 221 | 10639 | Caulimovirus | Viruses | 0.52 |
| 222 | 1279 | Staphylococcus | Bacteria | 0.51 |
| 223 | 249185 | Maculavirus | Viruses | 0.51 |
| 224 | 698776 | Cellulosilyticum | Bacteria | 0.50 |
| 225 | 1161127 | Murdochiella | Bacteria | 0.48 |
| 226 | 1623304 | Sfi21dt1virus | Viruses | 0.48 |
| 227 | 1582879 | Ezakiella | Bacteria | 0.46 |
| 228 | 1279388 | Kandleria | Bacteria | 0.46 |
| 229 | 40323 | Stenotrophomonas | Bacteria | 0.45 |
| 230 | 29521 | Brachyspira | Bacteria | 0.45 |
| 231 | 1291539 | Candidatus Methanomethylophilus | Archaea | 0.45 |
| 232 | 638847 | Pyramidobacter | Bacteria | 0.43 |
| 233 | 1230390 | Poacevirus | Viruses | 0.42 |
| 234 | 35823 | Arthrospira | Bacteria | 0.42 |
| 235 | 12059 | Enterovirus | Viruses | 0.41 |
| 236 | 12269 | Fabavirus | Viruses | 0.41 |
| 237 | 32067 | Leptotrichia | Bacteria | 0.41 |

| | | | | |
|---|---|---|---|---|
| 238 | 970 | Selenomonas | Bacteria | 0.40 |
| 239 | 547 | Enterobacter | Bacteria | 0.40 |
| 240 | 519427 | Sharpea | Bacteria | 0.38 |
| 241 | 11007 | Totivirus | Viruses | 0.37 |
| 242 | 2093 | Mycoplasma | Bacteria | 0.35 |
| 243 | 144051 | Cripavirus | Viruses | 0.35 |
| 244 | 95341 | Sapovirus | Viruses | 0.35 |
| 245 | 137757 | Ipomovirus | Viruses | 0.35 |
| 246 | 84162 | Cryptobacterium | Bacteria | 0.35 |
| 247 | 4919 | Pichia | Eukaryota | 0.34 |
| 248 | 1535326 | Candida | Eukaryota | 0.34 |
| 249 | 12051 | Marafivirus | Viruses | 0.32 |
| 250 | 47670 | Lautropia | Bacteria | 0.32 |
| 251 | 1213720 | Stomatobaculum | Bacteria | 0.30 |
| 252 | 39742 | Barnavirus | Viruses | 0.29 |
| 253 | 1935192 | Mobilibacterium | Bacteria | 0.29 |
| 254 | 674963 | Succinatimonas | Bacteria | 0.29 |
| 255 | 528 | Ochrobactrum | Bacteria | 0.28 |
| 256 | 129337 | Geobacillus | Bacteria | 0.28 |
| 257 | 186532 | C2virus | Viruses | 0.28 |
| 258 | 196082 | Parascardovia | Bacteria | 0.28 |
| 259 | 1945592 | Lagierella | Bacteria | 0.28 |
| 260 | 12148 | Tymovirus | Viruses | 0.27 |
| 261 | 29832 | Hanseniaspora | Eukaryota | 0.26 |
| 262 | 2050 | Mobiluncus | Bacteria | 0.25 |
| 263 | 66831 | Facklamia | Bacteria | 0.25 |
| 264 | 12293 | Tobravirus | Viruses | 0.24 |
| 265 | 83770 | Succinivibrio | Bacteria | 0.23 |
| 266 | 57493 | Kocuria | Bacteria | 0.23 |
| 267 | 1016 | Capnocytophaga | Bacteria | 0.22 |
| 268 | 39734 | Umbravirus | Viruses | 0.22 |
| 269 | 583 | Proteus | Bacteria | 0.20 |
| 270 | 1769710 | Sellimonas | Bacteria | 0.20 |
| 271 | 28263 | Arcanobacterium | Bacteria | 0.20 |
| 272 | 1472763 | Kallipyga | Bacteria | 0.19 |
| 273 | 5073 | Penicillium | Eukaryota | 0.19 |
| 274 | 581 | Morganella | Bacteria | 0.19 |
| 275 | 1849828 | Paeniclostridium | Bacteria | 0.19 |
| 276 | 1973274 | Niameybacter | Bacteria | 0.19 |
| 277 | 33882 | Microbacterium | Bacteria | 0.18 |
| 278 | 174708 | Allobaculum | Bacteria | 0.18 |
| 279 | 1111 | Porphyrobacter | Bacteria | 0.18 |
| 280 | 1375 | Aerococcus | Bacteria | 0.17 |
| 281 | 413970 | Anulavirus | Viruses | 0.17 |
| 282 | 82373 | Anaerovibrio | Bacteria | 0.16 |
| 283 | 254250 | Isoptericola | Bacteria | 0.16 |
| 284 | 5740 | Giardia | Eukaryota | 0.16 |
| 285 | 428711 | Jonquetella | Bacteria | 0.16 |

| | | | | |
|---|---|---|---|---|
| 286 | 300275 | Lachancea | Eukaryota | 0.16 |
| 287 | 44259 | Filifactor | Bacteria | 0.15 |
| 288 | 232799 | Iflavirus | Viruses | 0.15 |
| 289 | 180162 | Cetobacterium | Bacteria | 0.15 |
| 290 | 12289 | Enamovirus | Viruses | 0.15 |
| 291 | 1069494 | Trueperella | Bacteria | 0.15 |
| 292 | 71245 | Kazachstania | Eukaryota | 0.15 |
| 293 | 5340 | Agaricus | Eukaryota | 0.14 |
| 294 | 241189 | Hespellia | Bacteria | 0.13 |
| 295 | 579 | Kluyvera | Bacteria | 0.13 |
| 296 | 1696 | Brevibacterium | Bacteria | 0.12 |
| 297 | 184751 | Vitivirus | Viruses | 0.12 |
| 298 | 4958 | Debaryomyces | Eukaryota | 0.12 |
| 299 | 186886 | Pomovirus | Viruses | 0.12 |
| 300 | 150247 | Anoxybacillus | Bacteria | 0.11 |
| 301 | 1623303 | Sfi11virus | Viruses | 0.11 |
| 302 | 46254 | Oenococcus | Bacteria | 0.11 |
| 303 | 1653174 | Actinotignum | Bacteria | 0.11 |
| 304 | 113286 | Pseudoramibacter | Bacteria | 0.10 |
| 305 | 559173 | Fructobacillus | Bacteria | 0.10 |
| 306 | 374 | Bradyrhizobium | Bacteria | 0.09 |
| 307 | 604195 | Cyberlindnera | Eukaryota | 0.09 |
| 308 | 49082 | Candidatus Arthromitus | Bacteria | 0.09 |
| 309 | 46205 | Pseudobutyrivibrio | Bacteria | 0.09 |
| 310 | 39739 | Machlomovirus | Viruses | 0.08 |
| 311 | 177971 | Shuttleworthia | Bacteria | 0.08 |
| 312 | 106589 | Cupriavidus | Bacteria | 0.07 |
| 313 | 310684 | Cheravirus | Viruses | 0.07 |
| 314 | 586418 | Cosavirus | Viruses | 0.07 |
| 315 | 4910 | Kluyveromyces | Eukaryota | 0.07 |
| 316 | 29330 | Alicyclobacillus | Bacteria | 0.07 |
| 317 | 586 | Providencia | Bacteria | 0.07 |
| 318 | 43994 | Johnsonella | Bacteria | 0.07 |
| 319 | 12266 | Dianthovirus | Viruses | 0.07 |
| 320 | 54066 | Xylophilus | Bacteria | 0.07 |
| 321 | 1198140 | Felixo1virus | Viruses | 0.07 |
| 322 | 118747 | Bulleidia | Bacteria | 0.07 |
| 323 | 57499 | Kytococcus | Bacteria | 0.07 |
| 324 | 675073 | Torradovirus | Viruses | 0.07 |
| 325 | 253238 | Ethanoligenens | Bacteria | 0.07 |
| 326 | 283 | Comamonas | Bacteria | 0.07 |
| 327 | 45153 | Westerdykella | Eukaryota | 0.07 |
| 328 | 1663 | Arthrobacter | Bacteria | 0.06 |
| 329 | 186768 | Mitovirus | Viruses | 0.06 |
| 330 | 187214 | Soymovirus | Viruses | 0.06 |
| 331 | 5748 | Nannochloropsis | Eukaryota | 0.06 |
| 332 | 1911599 | Alphacarmovirus | Viruses | 0.06 |
| 333 | 12916 | Acidovorax | Bacteria | 0.06 |

| | | | | |
|---|---|---|---|---|
| 334 | 413496 | Cronobacter | Bacteria | 0.06 |
| 335 | 1511810 | Deltapartitivirus | Viruses | 0.06 |
| 336 | 203470 | Tepidiphilus | Bacteria | 0.06 |
| 337 | 1922209 | Triatovirus | Viruses | 0.06 |
| 338 | 1299309 | Alphanecrovirus | Viruses | 0.06 |
| 339 | 1562 | Desulfotomaculum | Bacteria | 0.06 |
| 340 | 28196 | Arcobacter | Bacteria | 0.06 |
| 341 | 5506 | Fusarium | Eukaryota | 0.05 |
| 342 | 688449 | Salivirus | Viruses | 0.05 |
| 343 | 1542744 | Gemycircularvirus | Viruses | 0.05 |
| 344 | 810 | Chlamydia | Bacteria | 0.05 |
| 345 | 289201 | Pontibacillus | Bacteria | 0.05 |
| 346 | 12008 | Allolevivirus | Viruses | 0.05 |
| 347 | 1912215 | Acidipropionibacterium | Bacteria | 0.05 |
| 348 | 542837 | Sp6virus | Viruses | 0.05 |
| 349 | 11990 | Levivirus | Viruses | 0.05 |
| 350 | 39725 | Circovirus | Viruses | 0.05 |
| 351 | 568 | Hafnia | Bacteria | 0.05 |
| 352 | 28105 | Sinorhizobium | Bacteria | 0.05 |
| 353 | 138954 | Parechovirus | Viruses | 0.05 |
| 354 | 39749 | Hypovirus | Viruses | 0.05 |
| 355 | 1910992 | K1gvirus | Viruses | 0.05 |
| 356 | 11305 | Cytorhabdovirus | Viruses | 0.05 |
| 357 | 675845 | Emaravirus | Viruses | 0.05 |
| 358 | 1505660 | Asaccharospora | Bacteria | 0.05 |
| 359 | 333750 | Alphapapillomavirus | Viruses | 0.05 |
| 360 | 1910954 | Phix174microvirus | Viruses | 0.05 |
| 361 | 88129 | Ophiovirus | Viruses | 0.05 |
| 362 | 114248 | Tepidimonas | Bacteria | 0.05 |
| 363 | 59732 | Chryseobacterium | Bacteria | 0.05 |
| 364 | 5475 | Candida | Eukaryota | 0.05 |
| 365 | 10814 | Begomovirus | Viruses | 0.05 |
| 366 | 1921117 | Nonagvirus | Viruses | 0.05 |
| 367 | 374468 | Nakaseomyces | Eukaryota | 0.05 |
| 368 | 140625 | Lachnobacterium | Bacteria | 0.05 |
| 369 | 157 | Treponema | Bacteria | 0.05 |
| 370 | 79603 | Denitrobacterium | Bacteria | 0.04 |
| 371 | 338 | Xanthomonas | Bacteria | 0.04 |
| 372 | 1794910 | Anaerosporomusa | Bacteria | 0.04 |
| 373 | 160674 | Raoultella | Bacteria | 0.04 |
| 374 | 3011 | Fucus | Eukaryota | 0.04 |
| 375 | 146 | Spirochaeta | Bacteria | 0.04 |
| 376 | 1940395 | Caenibacillus | Bacteria | 0.04 |
| 377 | 699166 | Citrivirus | Viruses | 0.04 |
| 378 | 653683 | Anaerosporobacter | Bacteria | 0.04 |
| 379 | 143901 | Benyvirus | Viruses | 0.04 |
| 380 | 28895 | Thermoanaerobacterium | Bacteria | 0.04 |
| 381 | 1269 | Micrococcus | Bacteria | 0.04 |

| | | | | |
|---|---|---|---|---|
| 382 | 1283209 | Alphacarmotetravirus | Viruses | 0.04 |
| 383 | 2147 | Acholeplasma | Bacteria | 0.04 |
| 384 | 1920860 | Kp36virus | Viruses | 0.04 |
| 385 | 2192 | Methanocorpusculum | Archaea | 0.04 |
| 386 | 1033 | Afipia | Bacteria | 0.04 |
| 387 | 186188 | Xylanimonas | Bacteria | 0.04 |
| 388 | 501783 | Cloacibacterium | Bacteria | 0.04 |
| 389 | 10509 | Mastadenovirus | Viruses | 0.04 |
| 390 | 10813 | Curtovirus | Viruses | 0.04 |
| 391 | 674962 | Bacillarnavirus | Viruses | 0.04 |
| 392 | 1921122 | Nonanavirus | Viruses | 0.04 |
| 393 | 538 | Eikenella | Bacteria | 0.04 |
| 394 | 31983 | Helcococcus | Bacteria | 0.04 |
| 395 | 36910 | Clavispora | Eukaryota | 0.04 |
| 396 | 256806 | Curvibasidium | Eukaryota | 0.04 |
| 397 | 1729679 | Faecalibaculum | Bacteria | 0.03 |
| 398 | 270 | Thermus | Bacteria | 0.03 |
| 399 | 13334 | Anaerobiospirillum | Bacteria | 0.03 |
| 400 | 2160 | Methanobacterium | Archaea | 0.03 |
| 401 | 12050 | Waikavirus | Viruses | 0.03 |
| 402 | 227307 | Gyrovirus | Viruses | 0.03 |
| 403 | 5352 | Lentinula | Eukaryota | 0.03 |
| 404 | 53335 | Pantoea | Bacteria | 0.03 |
| 405 | 1914852 | V5virus | Viruses | 0.03 |
| 406 | 69965 | Macrococcus | Bacteria | 0.03 |
| 407 | 32008 | Burkholderia | Bacteria | 0.03 |
| 408 | 2717 | Cardiobacterium | Bacteria | 0.03 |
| 409 | 373053 | Calditerricola | Bacteria | 0.03 |
| 410 | 28453 | Sphingobacterium | Bacteria | 0.03 |
| 411 | 378210 | Methyloversatilis | Bacteria | 0.03 |
| 412 | 909928 | Negativicoccus | Bacteria | 0.03 |
| 413 | 248038 | Mucispirillum | Bacteria | 0.03 |
| 414 | 4895 | Schizosaccharomyces | Eukaryota | 0.03 |
| 415 | 1792239 | Caviibacter | Bacteria | 0.03 |
| 416 | 1914295 | Prunevirus | Viruses | 0.03 |
| 417 | 249588 | Mamastrovirus | Viruses | 0.03 |
| 418 | 165812 | Sporanaerobacter | Bacteria | 0.03 |
| 419 | 5410 | Cystofilobasidium | Eukaryota | 0.02 |
| 420 | 187218 | T5virus | Viruses | 0.02 |
| 421 | 745 | Pasteurella | Bacteria | 0.02 |
| 422 | 10663 | T4virus | Viruses | 0.02 |
| 423 | 477967 | Phikmvvirus | Viruses | 0.02 |
| 424 | 104766 | Nanovirus | Viruses | 0.02 |
| 425 | 2289 | Desulfobacter | Bacteria | 0.02 |
| 426 | 1913651 | Rb49virus | Viruses | 0.02 |
| 427 | 4948 | Torulaspora | Eukaryota | 0.02 |
| 428 | 16 | Methylophilus | Bacteria | 0.02 |
| 429 | 1573 | Clavibacter | Bacteria | 0.02 |

| | | | | |
|---|---|---|---|---|
| 430 | 1159323 | Macellibacteroides | Bacteria | 0.02 |
| 431 | 1912216 | Cutibacterium | Bacteria | 0.02 |
| 432 | 61434 | Dehalococcoides | Bacteria | 0.02 |
| 433 | 201096 | Alicycliphilus | Bacteria | 0.02 |
| 434 | 10912 | Rotavirus | Viruses | 0.02 |
| 435 | 1513308 | Velarivirus | Viruses | 0.02 |
| 436 | 1920753 | G7cvirus | Viruses | 0.02 |
| 437 | 1913599 | Peptoanaerobacter | Bacteria | 0.02 |
| 438 | 13075 | Globicatella | Bacteria | 0.02 |
| 439 | 80865 | Delftia | Bacteria | 0.02 |
| 440 | 1827195 | Caballeronia | Bacteria | 0.02 |
| 441 | 41273 | Tissierella | Bacteria | 0.02 |
| 442 | 1922243 | Sextaecvirus | Viruses | 0.01 |
| 443 | 203133 | Propionimicrobium | Bacteria | 0.01 |
| 444 | 1611681 | Mucinivorans | Bacteria | 0.01 |
| 445 | 186536 | Ebolavirus | Viruses | 0.01 |
| 446 | 278028 | Naumovozyma | Eukaryota | 0.01 |
| 447 | 1707 | Cellulomonas | Bacteria | 0.01 |
| 448 | 1213379 | Aparavirus | Viruses | 0.01 |
| 449 | 107449 | Phaffia | Eukaryota | 0.01 |
| 450 | 83654 | Leclercia | Bacteria | 0.01 |
| 451 | 144193 | Turicella | Bacteria | 0.01 |
| 452 | 44249 | Paenibacillus | Bacteria | 0.01 |
| 453 | 1960084 | Rodentibacter | Bacteria | 0.01 |
| 454 | 400634 | Lysinibacillus | Bacteria | 0.01 |
| 455 | 1911600 | Betacarmovirus | Viruses | 0.01 |
| 456 | 227979 | Jeotgalicoccus | Bacteria | 0.01 |
| 457 | 53457 | Janibacter | Bacteria | 0.01 |
| 458 | 39744 | Rubulavirus | Viruses | 0.01 |
| 459 | 187217 | T1virus | Viruses | 0.01 |
| 460 | 110456 | T7virus | Viruses | 0.01 |
| 461 | 766728 | Meyerozyma | Eukaryota | 0.01 |
| 462 | 551 | Erwinia | Bacteria | 0.01 |
| 463 | 1914851 | Se1virus | Viruses | 0.01 |
| 464 | 171412 | Eremococcus | Bacteria | 0.01 |
| 465 | 497 | Psychrobacter | Bacteria | 0.01 |
| 466 | 675062 | Mycoflexivirus | Viruses | 0.01 |
| 467 | 1055323 | Aeribacillus | Bacteria | 0.01 |
| 468 | 1835 | Saccharopolyspora | Bacteria | 0.01 |
| 469 | 261933 | Pleomorphomonas | Bacteria | 0.01 |
| 470 | 85651 | Omegatetravirus | Viruses | 0.01 |
| 471 | 702 | Plesiomonas | Bacteria | 0.01 |
| 472 | 99479 | Tetrasphaera | Bacteria | 0.01 |
| 473 | 1920774 | Kp34virus | Viruses | 0.01 |
| 474 | 155493 | Gallibacterium | Bacteria | 0.01 |
| 475 | 186844 | Panicovirus | Viruses | 0.01 |
| 476 | 1637 | Listeria | Bacteria | 0.01 |
| 477 | 629 | Yersinia | Bacteria | 0.01 |

| | | | | |
|---|---|---|---|---|
| 478 | 5543 | Trichoderma | Eukaryota | 0.01 |
| 479 | 36739 | Dermabacter | Bacteria | 0.01 |
| 480 | 657 | Photobacterium | Bacteria | 0.01 |
| 481 | 190323 | Plantibacter | Bacteria | 0.01 |
| 482 | 1910951 | Alpha3microvirus | Viruses | 0.01 |
| 483 | 168470 | Laribacter | Bacteria | 0.01 |
| 484 | 81463 | Anaeromusa | Bacteria | 0.01 |
| 485 | 1322061 | Rhizoctonia | Eukaryota | 0.01 |
| 486 | 55506 | Schwartzia | Bacteria | 0.01 |
| 487 | 74380 | Undaria | Eukaryota | 0.01 |
| 488 | 1041 | Erythrobacter | Bacteria | 0.01 |
| 489 | 204037 | Dickeya | Bacteria | 0.01 |
| 490 | 5754 | Acanthamoeba | Eukaryota | 0.01 |
| 491 | 862 | Syntrophomonas | Bacteria | 0.01 |
| 492 | 582472 | Wohlfahrtiimonas | Bacteria | 0.01 |
| 493 | 1742989 | Glutamicibacter | Bacteria | 0.01 |
| 494 | 985001 | Piscicoccus | Bacteria | 0.01 |
| 495 | 39731 | Bymovirus | Viruses | 0.01 |
| 496 | 693996 | Alphacoronavirus | Viruses | 0.01 |
| 497 | 694002 | Betacoronavirus | Viruses | 0.01 |
| 498 | 1911929 | Bc431virus | Viruses | 0.01 |
| 499 | 1647 | Erysipelothrix | Bacteria | 0.01 |
| 500 | 10861 | Inovirus | Viruses | 0.01 |
| 501 | 1623289 | Hk578virus | Viruses | 0.01 |
| 502 | 407 | Methylobacterium | Bacteria | 0.01 |
| 503 | 2207 | Methanosarcina | Archaea | 0.01 |
| 504 | 29393 | Dolosigranulum | Bacteria | 0.01 |
| 505 | 635 | Edwardsiella | Bacteria | 0.01 |
| 506 | 94008 | Thermicanus | Bacteria | 0.01 |
| 507 | 90243 | Oligella | Bacteria | 0.01 |
| 508 | 2737 | Vagococcus | Bacteria | 0.01 |
| 509 | 33057 | Thauera | Bacteria | 0.01 |
| 510 | 5206 | Cryptococcus | Eukaryota | 0.01 |
| 511 | 150333 | Thermobrachium | Bacteria | 0.01 |
| 512 | 1279384 | Eggerthia | Bacteria | 0.01 |
| 513 | 156208 | Macluravirus | Viruses | 0.01 |
| 514 | 1930845 | Ndongobacter | Bacteria | 0.01 |
| 515 | 5010 | Taphrina | Eukaryota | 0.01 |
| 516 | 1330546 | Pluralibacter | Bacteria | 0.01 |
| 517 | 79808 | Coniochaeta | Eukaryota | 0.01 |
| 518 | 44000 | Caldicellulosiruptor | Bacteria | 0.01 |
| 519 | 371730 | N4virus | Viruses | 0.01 |
| 520 | 82802 | Trichococcus | Bacteria | 0.01 |
| 521 | 5455 | Colletotrichum | Eukaryota | 0.01 |
| 522 | 169998 | Phakopsora | Eukaryota | 0.01 |
| 523 | 150203 | Blastomonas | Bacteria | 0.01 |
| 524 | 12291 | Furovirus | Viruses | 0.01 |
| 525 | 119492 | Pecluvirus | Viruses | 0.01 |

| | | | | |
|---|---|---|---|---|
| 526 | 2422 | Fervidobacterium | Bacteria | 0.01 |
| 527 | 41275 | Brevundimonas | Bacteria | 0.01 |
| 528 | 489909 | Nosocomiicoccus | Bacteria | 0.01 |

Table S5: Top 100 KEGG functions in 10,000 human stool samples

| | **KEGG annotation** | **Prevalence in 10000 samples, %** |
|---|---|---|
| 1 | K00936 | 100.00 |
| 2 | K01190 | 99.99 |
| 3 | K03046 | 99.99 |
| 4 | K02355 | 99.99 |
| 5 | K00540 | 99.99 |
| 6 | K03695 | 99.99 |
| 7 | K03296 | 99.99 |
| 8 | K01362 | 99.98 |
| 9 | K02358 | 99.98 |
| 10 | K06950 | 99.98 |
| 11 | K03737 | 99.98 |
| 12 | K00975 | 99.98 |
| 13 | K02469 | 99.97 |
| 14 | K00754 | 99.97 |
| 15 | K03070 | 99.97 |
| 16 | K00705 | 99.97 |
| 17 | K02519 | 99.97 |
| 18 | K00599 | 99.97 |
| 19 | K03043 | 99.97 |
| 20 | K01834 | 99.96 |
| 21 | K01448 | 99.96 |
| 22 | K03654 | 99.96 |
| 23 | K04043 | 99.96 |
| 24 | K00100 | 99.96 |
| 25 | K00134 | 99.96 |
| 26 | K13993 | 99.96 |
| 27 | K03686 | 99.96 |
| 28 | K00262 | 99.96 |
| 29 | K01784 | 99.96 |
| 30 | K03088 | 99.96 |
| 31 | K02343 | 99.96 |
| 32 | K01006 | 99.96 |
| 33 | K00962 | 99.95 |
| 34 | K06400 | 99.95 |
| 35 | K07114 | 99.95 |
| 36 | K01952 | 99.95 |
| 37 | K01624 | 99.95 |
| 38 | K00688 | 99.95 |

| | | |
|---|---|---|
| 39 | K08303 | 99.95 |
| 40 | K02004 | 99.95 |
| 41 | K03530 | 99.95 |
| 42 | K07720 | 99.95 |
| 43 | K03086 | 99.95 |
| 44 | K01417 | 99.95 |
| 45 | K01238 | 99.95 |
| 46 | K04077 | 99.95 |
| 47 | K00615 | 99.95 |
| 48 | K03696 | 99.95 |
| 49 | K05808 | 99.95 |
| 50 | K02945 | 99.95 |
| 51 | K04079 | 99.95 |
| 52 | K02035 | 99.95 |
| 53 | K02529 | 99.95 |
| 54 | K03798 | 99.95 |
| 55 | K01840 | 99.94 |
| 56 | K00951 | 99.94 |
| 57 | K05349 | 99.94 |
| 58 | K01785 | 99.94 |
| 59 | K01610 | 99.94 |
| 60 | K02337 | 99.94 |
| 61 | K00845 | 99.94 |
| 62 | K00700 | 99.94 |
| 63 | K08884 | 99.93 |
| 64 | K02014 | 99.93 |
| 65 | K01951 | 99.93 |
| 66 | K02470 | 99.93 |
| 67 | K01689 | 99.93 |
| 68 | K00532 | 99.93 |
| 69 | K07335 | 99.92 |
| 70 | K07636 | 99.92 |
| 71 | K01262 | 99.92 |
| 72 | K01955 | 99.92 |
| 73 | K06147 | 99.92 |
| 74 | K02863 | 99.92 |
| 75 | K00680 | 99.92 |
| 76 | K02967 | 99.92 |
| 77 | K12373 | 99.92 |
| 78 | K00656 | 99.92 |
| 79 | K00945 | 99.92 |

| | | |
|---|---|---|
| 80 | K02886 | 99.92 |
| 81 | K01810 | 99.91 |
| 82 | K01270 | 99.91 |
| 83 | K02982 | 99.91 |
| 84 | K02111 | 99.91 |
| 85 | K01897 | 99.91 |
| 86 | K01925 | 99.91 |
| 87 | K02027 | 99.91 |
| 88 | K00927 | 99.91 |
| 89 | K03545 | 99.91 |
| 90 | K03733 | 99.91 |
| 91 | K01187 | 99.91 |
| 92 | K03076 | 99.90 |
| 93 | K03555 | 99.90 |
| 94 | K02600 | 99.90 |
| 95 | K07133 | 99.90 |
| 96 | K03531 | 99.90 |
| 97 | K02888 | 99.90 |
| 98 | K03701 | 99.90 |
| 99 | K02112 | 99.90 |
| 100 | K00850 | 99.90 |
